# Uncovering the genetic basis of agronomic traits in over 1,000 grapevine genotypes derived from a disease resistance breeding program

**DOI:** 10.1101/2025.10.05.680539

**Authors:** Clémentine Borrelli, Emilce Prado, Vincent Dumas, Guillaume Arnold, Christine Onimus, Gisèle Butterlin, Nathalie Jaegli, Sabine Wiedemann-Merdinoglu, Marie-Céline Lacombe, Marie-Annick Dorne, Aurélie Umar-Faruk, Serge Chaumonnot, Sophie Valentin, Lionel Ley, Jean-Sébastien Reynard, Jean-Laurent Spring, Éric Duchêne, Christophe Schneider, Didier Merdinoglu, Komlan Avia

## Abstract

Breeding disease-resistant grapevines that retain agronomic performance under variable climates requires loci and predictions that transfer across related hybrid families. We analyzed 1,081 genotypes from 95 crosses within the French INRAE-ResDur breeding program for 13 phenology, yield and berry-composition traits evaluated at five sites from 2006 to 2024. We integrated within-family QTL mapping, kinship- and population-adjusted multiple-population QTL mapping in 772 progeny, and structure-aware GWAS in 899-968 individuals, depending on the trait, together with cross-environment, cross-trait and local genomic estimated breeding-value analyses. Genetic and phenotypic differentiation among families accompanied extensive family specificity of detected loci. Of 79 family-QTL intervals, seven, representing six trait-region hypotheses, were locally concordant across all three mapping frameworks. The strongest recurrent evidence involved a chromosome-16 region for veraison and harvest date, where a localGEBV block at 14.69 Mb ranked first for veraison and second for harvest; chromosome-14 cluster traits and chromosome-1 compactness emerged as additional validation priorities. Cross-environment meta-analysis detected no common fixed-effect association at 5% FDR but revealed extensive heterogeneous evidence. Cross-trait analysis grouped 270 significant marker tests into 54 candidate multi-trait regions, 53 of which were retained when the derived trait mean cluster weight was excluded, without establishing biological pleiotropy. Population-adjusted localGEBV yielded a mean leave-one-population-out correlation of 0.470 between phenotypic BLUPs and genomic scores across traits. These results distinguish compact haplotype-validation targets from background- and environment-dependent signals, supporting a staged strategy that combines marker-assisted selection for validated recurrent regions with externally validated multi-trait, multi-environment genomic prediction for polygenic traits.

## Introduction

In the context of climate change and increasing constraints on pesticide use, marker-assisted selection (MAS) offers a way to accelerate grapevine breeding, provided that markers tag loci whose effects are stable across genetic backgrounds and target environments. MAS relies on molecular markers in linkage disequilibrium with the loci controlling phenotypic traits [1]. Quantitative trait locus (QTL) mapping and genome-wide association studies (GWAS) provide complementary evidence for identifying such loci: biparental QTL mapping has high power for alleles segregating within a family but limited resolution and transferability [2–4], whereas GWAS exploits broader recombination but is vulnerable to confounding by relatedness and population structure [5,6]. Mixed and multilocus approaches, including MLMM [7] and BLINK [8,9], address this confounding through kinship, population covariates, or iteratively selected marker cofactors.

Traditional GWAS captures primarily additive genetic effects, yet phenotypes can also arise from non-additive effects (dominance, recessivity, and overdominance) that shape the genetic architecture of complex traits and can reveal loci explaining little additive variance. Non-additive GWAS has proven informative across taxa, including the heterozygous perennial crop almond [10], as well as livestock [11] and human complex-disease [12] studies, so accounting for these effects complements additive models and can expose associations they would miss. Genetic determinism is further modulated by the environment, as genotype-by-environment (GxE) interactions alter gene expression and individual performance, and associations detected in one environment may not hold in another. Meta-analysis of GWAS addresses this by combining evidence across populations and environments to resolve the genetic architecture of complex traits [13]; well established in human [14,15] and animal [16,17] genetics, it is increasingly applied in plants such as maize [13], tomato [18], peach, and apricot [19], and can additionally reveal pleiotropy by identifying loci associated with multiple traits [20]. A further layer of complexity arises in connected breeding families, where population structure is not merely a statistical nuisance to be corrected but reflects genuine differences in parental allele frequencies, linkage phase, and genetic background among crosses. Consequently, a locus cannot be considered transferable simply because QTL and GWAS signals fall on the same chromosome: establishing that it is shared, and usable for MAS across families, requires evidence of physical co-location, robustness to correction for population structure, and segregation of the underlying alleles within each family [21].

*Vitis vinifera* is the principal cultivated grapevine species, prized for its yield and organoleptic qualities and propagated vegetatively to fix the traits of emblematic varietals. Its domestication began approximately 11,000 years ago through at least two independent events in Western Asia and the Caucasus [22]. Cultivated grapevine is nonetheless genetically narrow and susceptible to major diseases, whereas the roughly 70 wild *Vitis* species of North America and Asia carry valuable resistances, notably to downy and powdery mildew, but generally lack agronomic performance and sensory quality [23]. Durable-resistance breeding therefore pyramids resistance loci from wild sources through repeated crossing while recovering the fruit and wine quality of *V. vinifera*, an effort supported by a well-characterized genome of 19 chromosomes spanning about 475 Mb, with a reference assembly available since 2007 [24] and high-quality assemblies since [25,26]. Because introgression must raise disease resistance without eroding agronomic value, effective selection also requires understanding the genetic determinism of the agronomic traits themselves, and in grapevine these analytical challenges are acute: its high heterozygosity makes non-additive gene action a plausible contributor to complex traits [10–12], multi-environment cultivation makes GxE a central concern [13,18,19], and its many correlated traits make pleiotropy worth testing explicitly [27].

Regulatory restrictions on fungicide use, tightened in Europe under the EU Green Deal [23], have made durable disease resistance a central breeding objective. Public programs in several countries, including JKI and WBI in Germany, VCR, IGA and FEM in Italy, and INRAE and IFV in France, aim to combine resistance to downy mildew, powdery mildew, and black rot with the agronomic performance and organoleptic quality of *V. vinifera* [28–31]. These programs already use MAS to track introgressed resistance loci, but its extension to agronomic traits such as phenology, yield, and berry composition remains limited, even though their genetic determinism has been studied (reviewed by Tello et al. [32]). QTL analyses have mapped loci for phenology [33], berry weight [34–36], sugars and acids [37,38], cluster compactness [39,40], bunch rot [41], or yield components [42,43], and GWAS has broadened this catalogue, from the first grapevine association study in 2007 [44] to diversity panels such as Flutre et al.’s survey of 127 traits [45]. Results from *V. vinifera* panels, however, may not transfer directly to interspecific breeding material, and the associations detected can depend on the choice of response variable and analytical model [46]. Non-additive gene action, cross-environment meta-analysis, and pleiotropy nonetheless remain largely unexplored in grapevine, and few studies have jointly evaluated family differentiation, non-additive encodings, GxE, and cross-trait association.

Genomic selection (GS) complements locus mapping by predicting performance from genome-wide variation, including effects too small to detect individually. Recent grapevine studies have integrated pangenome variation with the analysis of 29 agronomic traits and genomic breeding [47], combined polygenic mapping and prediction for seedlessness [48], and coupled deep-learning phenotyping with machine-learning prediction of pest resistance [49]. Our study complements these approaches in connected hybrid breeding families by combining locus discovery with local genomic prediction and testing predictions across held-out populations, thereby examining transferability within an operational resistance-breeding program.

Here we leverage the French INRAE-ResDur program [31], initiated in the early 2000s at the INRAE center in Colmar to pyramid durable resistance to downy and powdery mildews while retaining wine-grape performance, and which has already registered 18 resistant varieties in the French national catalogue. The program has proceeded in three phases that differ in the downy mildew resistance factors pyramided (*Rpv1* and *Rpv3* in phase 1, *Rpv1* and *Rpv10* in phase 2, and *Rpv1*, *Rpv3* and *Rpv10* in phase 3), whereas the same three powdery mildew resistance factors (*Run1*, *Ren3* and *Ren9*) were used throughout. The population comprises 1,081 genotypes from 95 connected crosses, phenotyped between 2006 and 2024 at five sites in France and Switzerland. Using this operational breeding population, we ask which agronomic loci are reproducible across families, analytical models, and environments, and which signals instead remain contingent on family structure or non-additive encoding. Specifically, we (1) quantify phenotypic and genome-wide differentiation among the six largest families and test whether design imbalance explains variation in broad-sense heritability; (2) map QTLs within individual families; (3) perform multi-population QTL mapping on a connected-family composite map with kinship and family indicators; (4) compare additive GWAS models after explicit ancestry adjustment; (5) extend GWAS to ancestry-adjusted dominant, recessive, and overdominant encodings, environmental meta-analysis, and cross-trait composite tests; (6) integrate the three mapping frameworks at the physical-coordinate level, together with local linkage disequilibrium, family allele frequencies, and PN40024.v4 annotation; and (7) complement locus discovery with population-adjusted localGEBV block ranking and family-held-out prediction to assess breeding relevance for genomic selection and targeted marker-panel development.

## Results

To distinguish family-specific associations from recurrent breeding targets, we first characterized environmental and family differentiation. Within-family QTL mapping tested segregating loci in individual crosses, while population-adjusted multiple-population mapping and structure-aware GWAS examined associations across the connected panel. We then tested alternative genotype encodings and association patterns across environments and traits, before evaluating the aggregate predictive value of genomic segments with localGEBV.

### Environmental and genetic differentiation in the ResDur panel

The agronomic panel comprised 1,081 genotypes from 95 INRAE-ResDur crosses evaluated at five sites from 2006 to 2024. The population-structure visualization used a subset of 344 individuals comprising *V. vinifera* accessions, other interspecific hybrids, ResDur parents and progeny. Principal component analysis of harmonized annual climate descriptors separated the sites along gradients dominated by temperature, radiation and evapotranspiration, and by rainfall; the first two components explained 90.3% of the variation (Figure 1A-B). Thus, the panel combined contrasting climatic settings with an unbalanced distribution of genotypes among environments.

Discriminant analysis of principal components (DAPC) of the 344-individual subset, fitted with nine principal components to seven genotype-derived clusters, separated the ampelographic groups: *V. vinifera* accessions formed a compact group at positive LD1 values, whereas the other interspecific hybrids spanned a wide range of LD1 values; ResDur parents and progeny overlapped both groups rather than forming a separate cluster (Figure 1C; Supplementary File 1). The genomic relationship matrix additionally revealed family blocks and relationships among connected families (Supplementary Figure 1). Among the six largest families, pairwise Hudson FST ranged from 0.111 to 0.253, establishing substantial differentiation of the segregating allele pools (Supplementary Figure 2; Supplementary Table 1). These patterns motivated ancestry covariates in all GWAS and kinship adjustment in the mixed-model scans.

**Figure 1.**
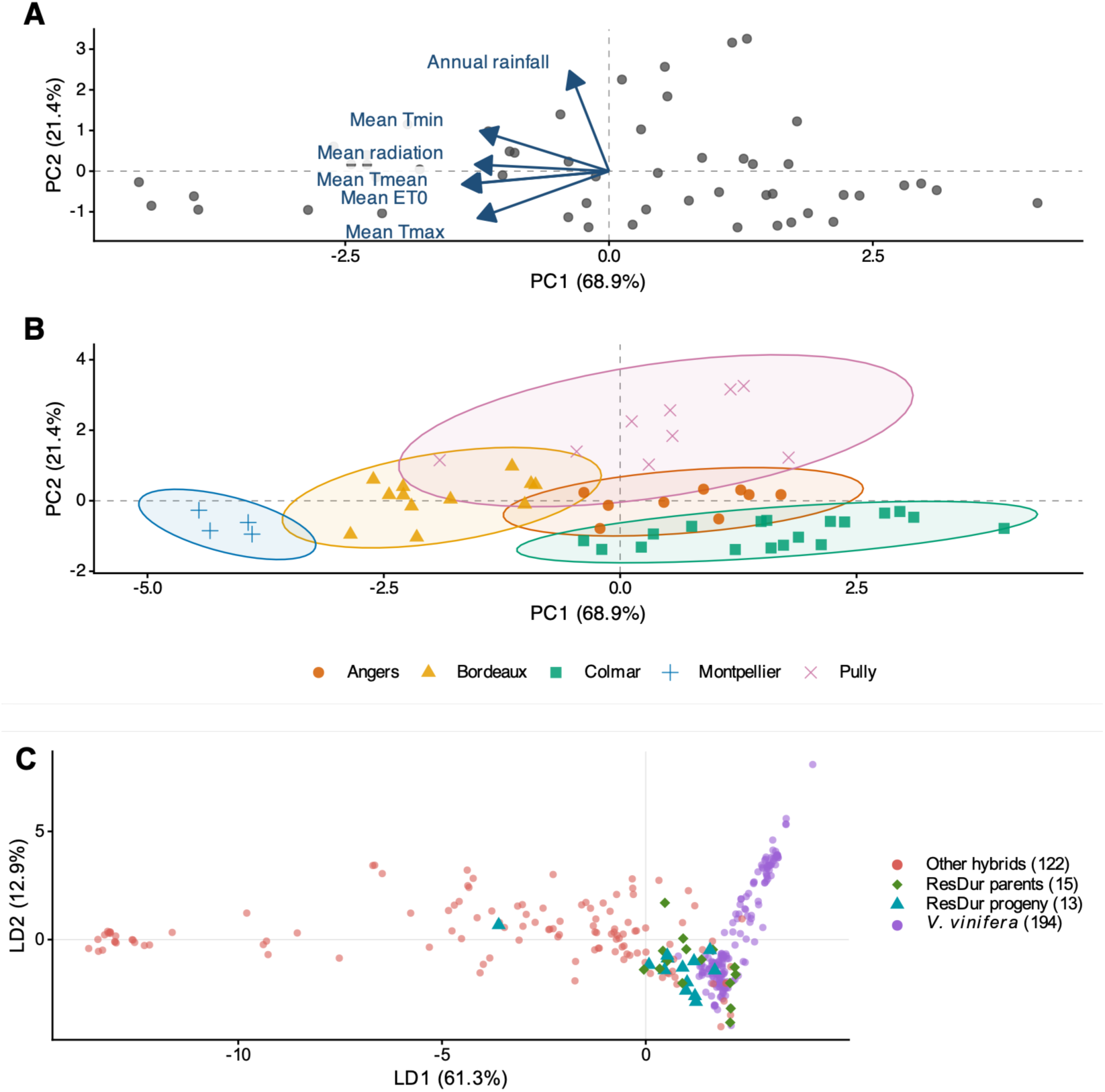
Environmental and population-structure analyses. (A) Loading biplot and (B) site x year scores from principal component analysis of annual mean minimum, maximum and mean temperature, annual rainfall, mean reference evapotranspiration and mean solar radiation. (C) DAPC of 344 individuals (194 *V. vinifera* accessions, 122 other interspecific hybrids, 15 ResDur parents and 13 progeny) fitted with nine principal components to seven genotype-derived clusters. LD1 and LD2 are the first two of the six linear discriminant functions, with their shares of the sum of discriminant eigenvalues; colors and symbols denote the biological categories. Settings and eigenvalues are given in Supplementary File 1.

### Trait variation and within-family heritability

Trait distributions differed both in shape and among families. Veraison date, berry pH and berry weight were comparatively symmetric, whereas several yield traits were skewed and compactness was discrete (Figure 2). Family effects on environment-adjusted BLUPs were substantial (η^2^ = 17.8-80.9% across traits; Supplementary Figure 3; Supplementary Table 2), including later phenology (flowering, veraison and harvest), berry composition (sugar content, titratable acidity and pH), and yield components (berry weight, cluster compactness, cluster number, total and mean cluster weight, and yield per area). These phenotypic differences accompanied the genomic differentiation, providing context for comparing QTL detection among families without identifying its causal mechanism.

**Figure 2.**
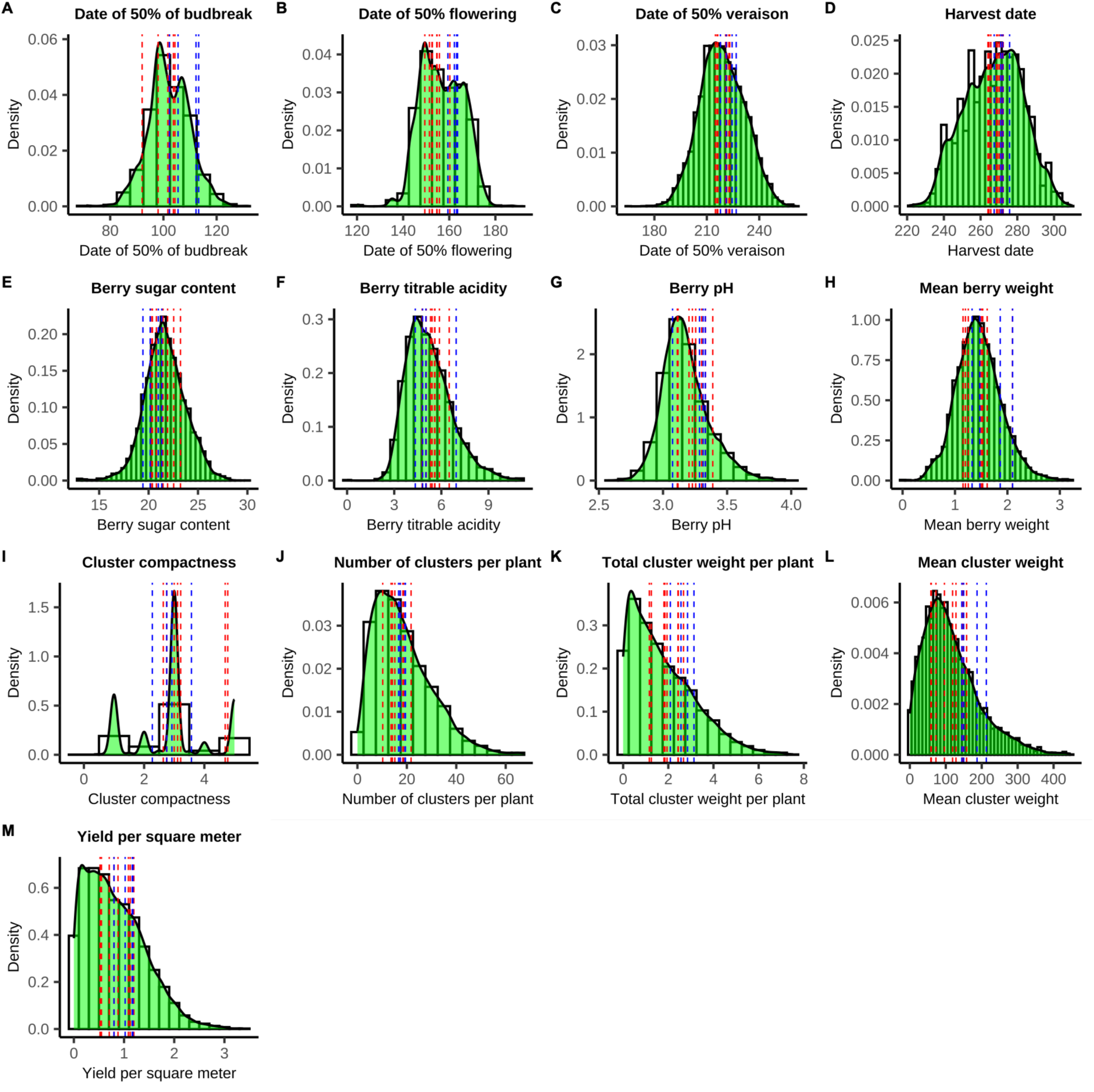
Distribution of phenotypes across the INRAE-ResDur individuals. Green curves show density; dashed lines show the means of raw observations from the intermediate-stage evaluation dataset for the four varieties registered from INRAE-ResDur phase 1 (blue: Artaban, Vidoc, Floreal and Voltis) and the eight registered from phase 2 (red: Sirano, Artys, Selenor, Opalor, Exelys, Calys, Coliris and Lilaro); the two phases differ in the resistance factors pyramided against downy mildew (see Introduction section). These means are descriptive benchmarks, not adjusted for the unequal years and sites contributing to each variety and trait (Supplementary File 1). Traits: (A) budbreak; (B) flowering; (C) veraison; (D) harvest date; (E) berry sugar content; (F) berry titratable acidity; (G) berry pH; (H) berry weight; (I) cluster compactness; (J) clusters per plant; (K) total cluster weight per plant; (L) mean cluster weight; (M) yield per m^2^.

Pearson correlations were calculated from 1,671 complete raw phenotypic observations representing 863 ResDur genotypes (Figure 3). Veraison and harvest were strongly correlated (r = 0.909; genotype-clustered bootstrap 95% interval, 0.901-0.917); this relationship persisted after averaging records within genotype (r = 0.930) and using the existing environment-adjusted BLUPs for the same genotypes (r = 0.951). Budbreak was less closely coupled to later phenology. Yield per vine remained positively associated with cluster number and mean cluster weight in all three analyses. However, the association between these two components depended on the values analyzed (r = -0.009 for raw records, 0.101 for genotype means and 0.545 for BLUPs; Supplementary File 1). Yield per vine and per area were nearly perfectly correlated (r = 0.99835, rounded to 1.00), reflecting their numerical conversion relationship rather than independent measurements. These pooled descriptive relationships motivated the subsequent cross-trait analysis; they combine environmental, family and within-genotype variation and do not establish shared causal loci.

**Figure 3.**
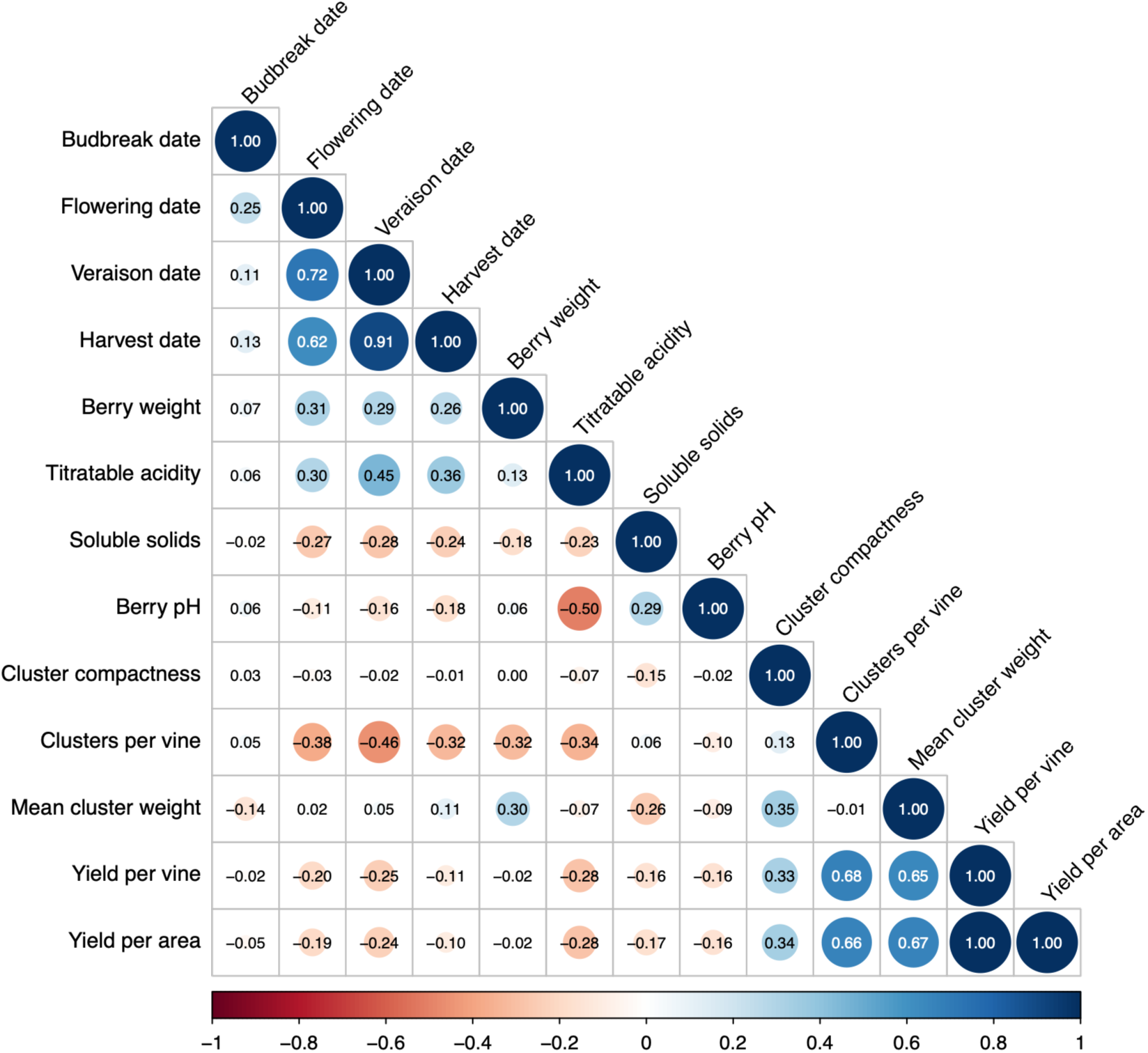
Raw phenotypic correlations among ResDur progeny. Pearson correlations use 1,671 complete records from 863 genotypes; repeated records were retained without averaging or environmental adjustment. Circle area and color indicate correlation magnitude and sign; values are rounded to two decimals. The yield-per-vine versus yield-per-area value is 0.99835, displayed as 1.00 by rounding. Genotype-mean and BLUP comparisons and genotype-clustered bootstrap intervals are reported in Supplementary File 1.

Mixed models were fitted separately by trait and analysis set to account for the unequal representation of genotypes across sites and years before obtaining BLUPs. Trait coverage and selected models are reported in Supplementary Tables 3 and 4. Within-family broad-sense heritability (Table 1) was generally high but trait dependent (median = 0.74): veraison was consistently heritable, whereas budbreak and flowering showed greater variation among families. Across estimable family-by-trait combinations, heritability was not significantly associated with the number of detected QTLs (Spearman ρ = 0.127, P = 0.276; Supplementary Tables 5 and 6). High within-family heritability therefore did not consistently translate into detectable loci in these family scans.

**Table 1.** Broad-sense heritability estimates per trait and family.

| Trait | 50025 | 42050 | 50001 | 50013 | 50015 | 50035 |
| --- | --- | --- | --- | --- | --- | --- |
| Budbreak | 0.70 | 0.25 | 0.49 | 0.75 | 0.76 | 0.68 |
| Flowering | 0.68 | 0.65 | 0.16 | 0.12 | 0.80 | 0.76 |
| Veraison | 0.93 | 0.86 | 0.86 | 0.86 | 0.92 | 0.93 |
| Mean berry weight | 0.62 | 0.91 | 0.83 | 0.88 | 0.83 | 0.87 |
| Berry sugar content | 0.70 | 0.75 | 0.57 | 0.65 | 0.71 | 0.71 |
| Berry pH | 0.46 | 0.76 | 0.84 | 0.54 | 0.66 | 0.64 |
| Berry titratable acidity | 0.72 | 0.84 | 0.67 | 0.75 | 0.84 | 0.74 |
| Harvest date | 0.74 | 0.72 | 0.78 | 0.75 | 0.84 | 0.74 |
| Cluster compactness | 0.57 | 0.76 | 0.85 | 0.85 | 0.36 | 0.62 |
| Number of clusters per plant | 0.67 | 0.81 | NA | 0.83 | 0.77 | 0.68 |
| Cluster weight per plant | 0.71 | 0.74 | NA | 0.82 | 0.61 | 0.74 |
| Mean cluster weight | 0.68 | 0.68 | NA | 0.78 | 0.54 | 0.81 |
| Yield per m<sup>2</sup> | 0.62 | 0.74 | 0.67 | 0.79 | 0.60 | 0.79 |

### Complementary mapping identifies family-specific and recurrent regions

Within-family QTL mapping tested segregating effects in six large crosses. Their maps were integrated with those of other connected families into a composite map representing 792 progeny from 12 families (Figure 4; Supplementary Figure 4; Supplementary Table 7). The family-QTL catalogue contains 79 BLUP-based intervals, with detection varying markedly among crosses. Most signals were family specific, but chromosome-16 veraison and harvest-date QTLs recurred in environment-specific scans of families 50025 and 50035, with additional BLUP-based support in family 50001 (Supplementary Tables 8 and 9).

**Figure 4.**
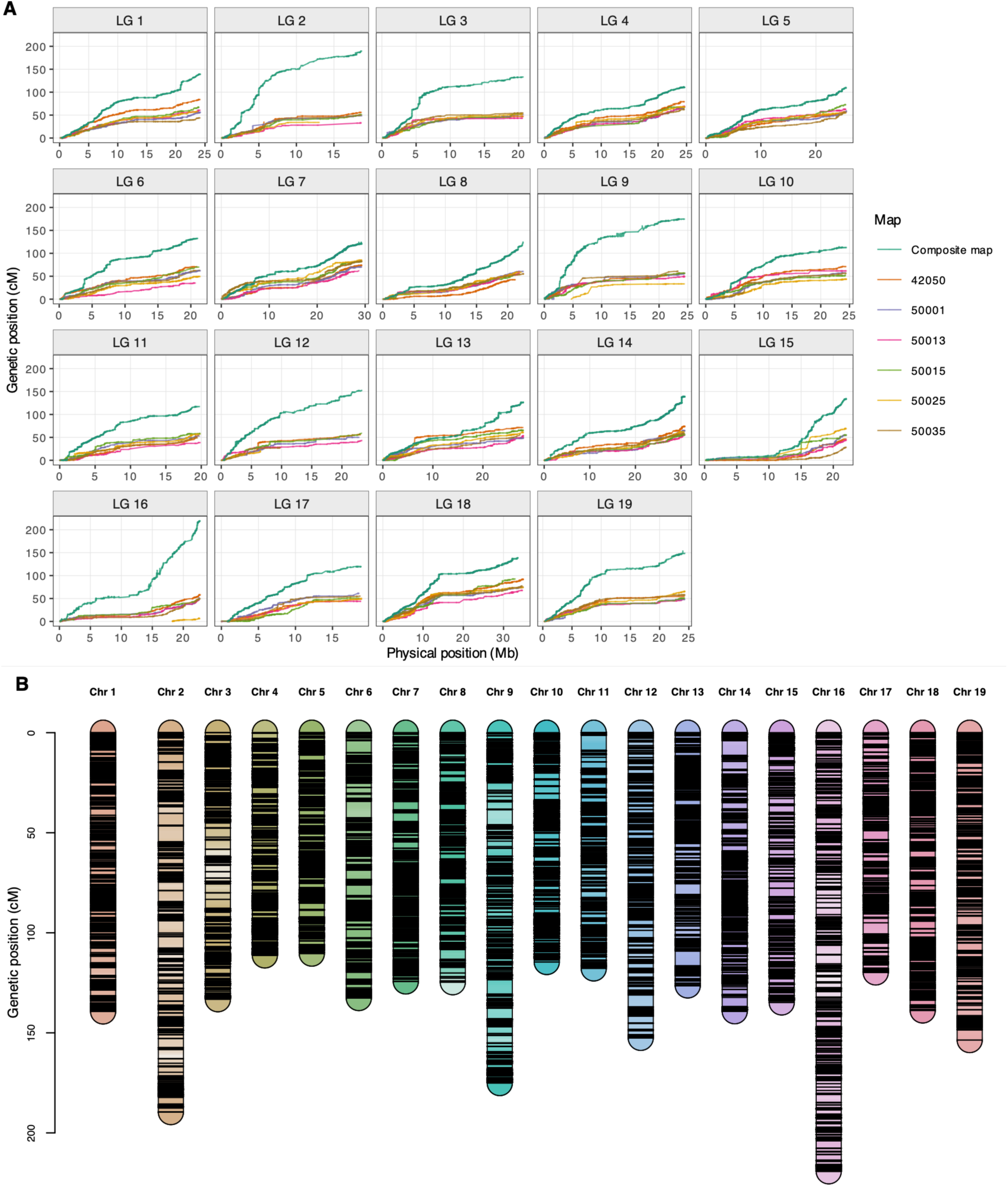
Genetic maps of biparental families and the connected-family composite panel. (A) Genetic position as a function of PN40024.v4 physical position for each chromosome and family; lines and semi-transparent points reduce overplotting. (B) LinkageMapView density map of the 19 composite linkage groups. Each chromosome has a distinct hue. Within each hue, color saturation is based on a common genome-wide transformation of local marker density calculated in 1-cM sections with a 30-cM sliding window; more saturated sections are denser. Black horizontal ticks show marker positions.

To examine pooled signals across connected crosses, multiple-population QTL mapping (mpQTL) used 772 progeny assigned to 12 population groups and 8,054 complete-genotype markers, adjusting for realized kinship and population indicators. The scan detected 74 marker-trait associations at the Li-Ji threshold, defining 45 nominal support intervals. Signals occurred for all 13 traits, including chromosome-16 phenology and chromosome-14 cluster traits (Supplementary Tables 10-12; Supplementary File 2). These pooled associations complement the family scans but do not establish that the same alleles segregate or act in every family.

Additive GWAS extended the comparison to 899-968 genotypes per trait. Twenty-eight trait-marker combinations were supported by at least two models, although total detection differed among models (Supplementary Tables 13 and 14; Supplementary Figure 5; Supplementary File 3). Different marker-selection procedures and treatment of background effects can change detection, so model-specific counts are not directly comparable. We therefore prioritized local agreement with linkage-based mapping rather than the number of significant markers from any single model.

Physical-coordinate integration identified seven family-QTL intervals, representing six trait-region hypotheses, with locally concordant support from all three mapping frameworks (Table 2). The clearest recurrence concerned chromosome-16 veraison and harvest date, with additional concordance for budbreak on chromosome 16, compactness on chromosomes 1 and 14, and mean cluster weight on chromosome 14. Thirty-eight intervals had support from two frameworks; representative examples are included alongside Tier A intervals in Table 2. One additional broad chromosome-2 berry-weight interval with suggestive family-QTL support contained all three evidence types, but the pooled-QTL and GWAS signals were spatially separated and it was not classified as locally concordant. The complete classification and positional candidate-gene evidence are reported in Supplementary Tables 15-18 and Supplementary Figure 6. Because the frameworks reuse overlapping progeny, concordance provides complementary evidence rather than independent replication.

**Table 2.** Physical-coordinate integration of single-family QTL, population-adjusted multiple-population QTL and GWAS evidence. Coordinates refer to PN40024.v4. Each row represents a trait-family interval; all representative entries from the integration analysis are shown, including two-framework evidence. Tier A requires all three frameworks and local mpQTL-GWAS concordance (overlap or a gap ≤500 kb within the family QTL): A1, at least two adjusted GWAS models; A2, one model. B1, family QTL plus mpQTL; B2, family QTL plus GWAS. GWAS LD bounds use local r^2^ ≥ 0.20 within ±500 kb of the lead. mpQTL entries are lead positions or their range, not support boundaries. Candidate genes are positional hypotheses, not causal assignments. Chr, chromosome; Mb, megabase; LD, linkage disequilibrium; —, no corresponding support or candidate listed. Full integration evidence, candidate genes, and family-specific allele frequencies and effects are provided in Supplementary Tables 15-18; mpQTL results and diagnostics are provided in Supplementary Tables 10-12 and Supplementary File 2.

| Trait | Family | Chr | Family QTL (Mb) | mpQTL lead(s) (Mb) | GWAS lead (LD bounds; Mb) | Tier | Positional candidate genes |
| --- | --- | --- | --- | --- | --- | --- | --- |
| Budbreak | 50035 | 16 | 2.00-16.80 | 15.361 | 15.108<br>(14.684-15.592) | A2 | Vitvi16g00859<br>Vitvi16g00882 |
| Budbreak | 50025 | 7 | 0.40-18.20 | 4.415 | — | B1 | — |
| Flowering | 42050 | 7 | 17.50-24.30 | 20.342 | — | B1 | — |
| Flowering | 50035 | 7 | 4.10-20.60 | 20.342 | — | B1 | — |
| Veraison | 50025 | 16 | 14.20-15.20 | 14.690 | 15.035<br>(14.550-15.178) | A1 | Vitvi16g00849<br>Vitvi16g00853 |
| Veraison | 50035 | 16 | 0.10-17.60 | 7.122-14.690 | 15.035<br>(14.550-15.488) | A1 | Vitvi16g00849<br>Vitvi16g00882 |
| Harvest date | 50025 | 16 | 14.00-15.20 | 14.690 | 15.035<br>(14.550-15.178) | A1 | Vitvi16g00849<br>Vitvi16g00853 |
| Harvest date | 50015 | 8 | 0.20-20.70 | — | 5.781<br>(5.781-5.781) | B2 | Vitvi08g00289 |
| Mean berry weight | 50013 | 7 | 4.70-28.70 | — | 23.591<br>(23.186-24.088) | B2 | Vitvi07g04654<br>Vitvi07g04655 |
| Mean berry weight | 50035 | 11 | 9.60-19.50 | — | 18.029<br>(17.558-18.508) | B2 | Vitvi11g01186<br>Vitvi11g01641 |
| Berry sugar content | 50035 | 9 | 0.10-2.70 | 2.155 | — | B1 | — |
| Berry sugar content | 50035 | 6 | 3.60-7.30 | 7.263 | — | B1 | — |
| Berry pH | 50035 | 4 | 8.10-18.90 | — | 11.935<br>(11.886-11.947) | B2 | Vitvi04g04257<br>Vitvi04g04256 |
| Berry pH | 50035 | 16 | 15.00-16.80 | — | 16.539<br>(16.192-16.793) | B2 | Vitvi16g00936<br>Vitvi16g00953 |
| Titrateable acidity | 50015 | 13 | 3.90-7.10 | — | 5.515<br>(5.026-5.991) | B2 | Vitvi13g00594<br>Vitvi13g00596<br>Vitvi13g00601 |
| Titrateable acidity | 50025 | 14 | 21.00-29.80 | — | 22.282<br>(21.879-22.761) | B2 | Vitvi14g04452<br>Vitvi14g01269 |
| Cluster compactness | 50035 | 1 | 2.00-5.80 | 5.742 | 5.671<br>(5.228-5.791) | A1 | Vitvi01g00476<br>Vitvi01g00475 |
| Cluster compactness | 50013 | 14 | 20.70-30.80 | 26.846-29.206 | 26.574<br>(26.074-27.063) | A2 | Vitvi14g01576<br>Vitvi14g01574 |
| Clusters per plant | 50035 | 7 | 1.80-5.80 | — | 2.425<br>(1.952-2.919) | B2 | Vitvi07g00213<br>Vitvi07g00209 |
| Cluster weight per plant | 50013 | 14 | 20.70-29.60 | 22.600-26.075 | — | B1 | — |
| Cluster weight per plant | 50035 | 14 | 17.40-27.40 | 22.600-26.075 | — | B1 | — |
| Mean cluster weight | 50013 | 14 | 22.70-29.60 | 22.600-26.113 | 27.029<br>(26.534-27.518) | A2 | Vitvi14g01596<br>Vitvi14g01593 |
| Mean cluster weight | 50035 | 16 | 14.40-16.70 | — | 15.543<br>(15.074-15.946) | B2 | Vitvi16g00886<br>Vitvi16g00890 |
| Yield per m<sup>2</sup> | 50013 | 14 | 19.50-29.60 | 22.600 | — | B1 | — |
| Yield per m<sup>2</sup> | 50035 | 14 | 18.00-28.40 | 22.600 | — | B1 | — |

### Alternative genotype encodings identify additional association patterns

To examine associations not adequately represented by additive dosage, we tested dominant, recessive and overdominant genotype encodings. Of 606 BLINK candidates, 74 passed the genome-wide threshold in a selected-candidate retest incorporating both kinship and DAPC axes (19 dominant, 27 recessive and 28 overdominant; Supplementary Tables 13 and 19; Supplementary Figure 7; Supplementary File 4). Of the retained non-additive candidates, 18 coincided with an exact additive signal and 25 lay within 250 kb of one, whereas others provided additional signals. The marked reduction after kinship adjustment emphasizes the importance of relatedness control when interpreting non-additive patterns in this breeding panel; these encoded contrasts do not by themselves establish a causal mode of gene action.

### Environment-specific analyses reveal heterogeneous association evidence

To examine whether associations were consistent across environments, we combined 303 site-by-year GWAS. Fixed-effect meta-analysis tested a common association, random-effect analysis allowed variation among environments, local scores assessed spatially clustered marker evidence, and meta-regression examined covariation with climatic descriptors. No marker passed the fixed-effect false-discovery-rate threshold, whereas the random-effect tests yielded 4,629 significant marker-trait results and the local-score analysis defined 30 fixed-effect and 692 random-effect zones (Supplementary Table 20; Supplementary Figures 8 and 9; Supplementary File 5). Climate-associated signals (504 marker-covariate tests passing 5% FDR) were concentrated in flowering (470 marker tests in 17 groups, particularly mean minimum temperature and annual rainfall). For berry pH, a chromosome-16 climate (rainfall and maximum-temperature)-associated marker (chr16_14486694) was in population-adjusted linkage disequilibrium (r^2^ = 0.622) with an additive GWAS signal (chr16_14447625), identifying a focused candidate for environmental follow-up. These patterns support investigation of context-dependent associations, although changing family composition prevents attributing the observed variation exclusively to climate.

### Cross-trait analysis identifies candidate shared regions

We next asked whether individual markers were associated with more than one of the correlated traits. Query Composite Hypotheses (QCH) accounted for dependence among the trait tests and selected 270 markers for association with at least two traits at 5% false discovery rate. Grouping nearby selected markers yielded 54 candidate multi-trait regions across 15 chromosomes (Figure 5; Supplementary Table 21). The exact rejection set was sensitive to the copula specification despite strong genome-wide rank agreement (Supplementary Table 22). Because mean cluster weight is calculated as cluster weight per plant divided by cluster number, and yield per area is derived from yield per vine, shared signals among these traits could partly reflect trait construction. When QCH was refitted without mean cluster weight (12 traits), 53 of the 54 regions retained at least one of their originally selected markers; without mean cluster weight and yield per area (11 traits), 38 regions were retained, the regions lost being mainly those supported by the cluster-yield traits. The three chromosome-16 markers associated with both veraison and harvest date were retained in both analyses (Supplementary File 1, section D; Supplementary Excel File). These regions identify shared association patterns for follow-up, without distinguishing pleiotropy from linkage between trait-specific variants.

**Figure 5.**
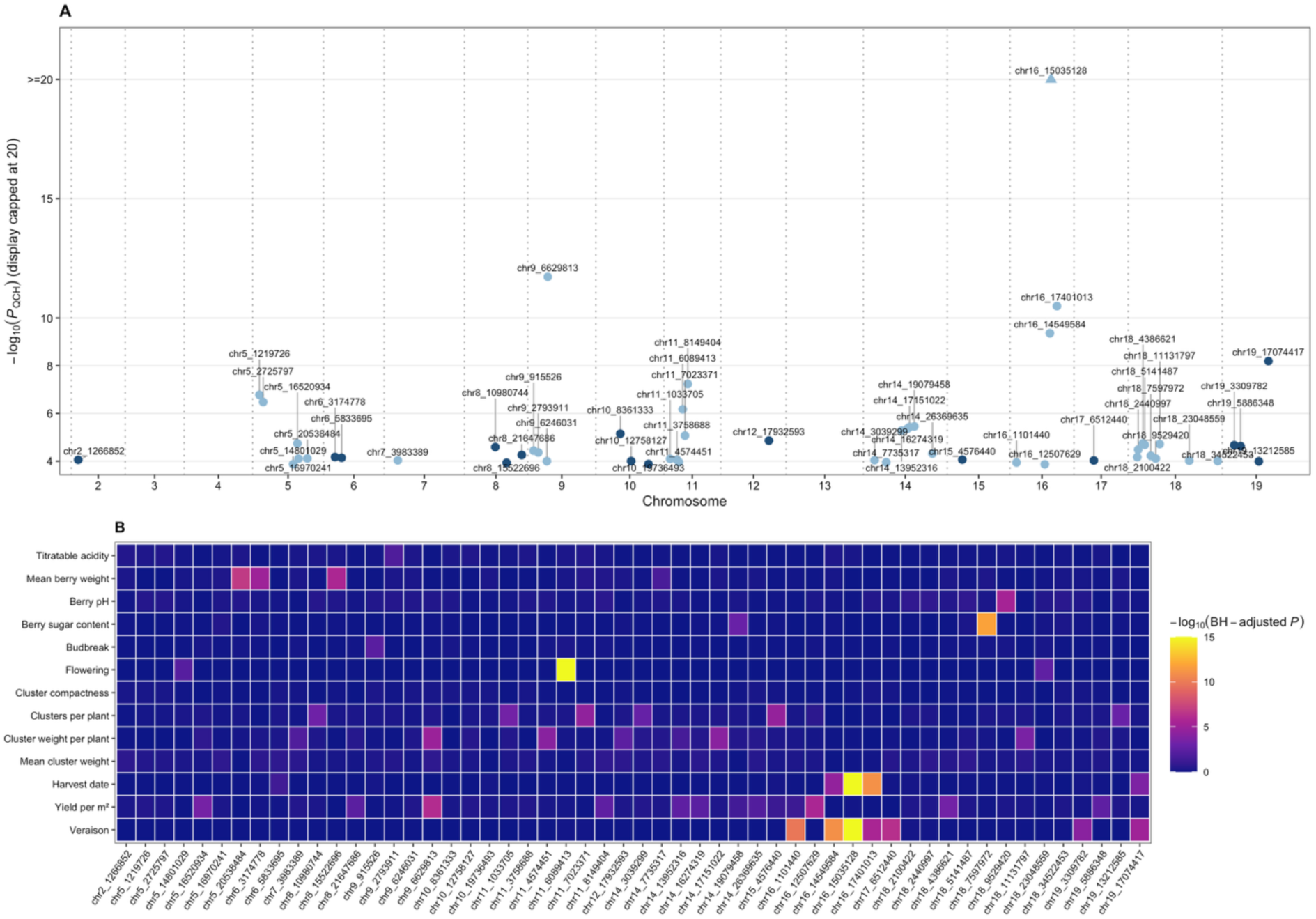
Cross-trait QCH results from the ancestry-adjusted BLINK scans. The 5% QCH FDR procedure selected 270 marker tests for association with at least two traits; adjacent selected SNPs no more than 250 kb apart were joined into 54 display loci. (A) -log10 of the raw composite QCH P value for the lead marker in each display locus. Scores above 20 are capped at 20 for legibility and shown as upward triangles; uncapped values are given in Supplementary Table 21. Chromosomes are separated by dotted lines placed in inter-marker gaps, and every plotted lead is labeled by marker identifier. (B) Descriptive heat map of -log10 Benjamini-Hochberg-adjusted P-values from the corresponding single-trait GWAS for the same lead markers. Retention of each display locus in the sensitivity analyses excluding the derived yield traits is reported in Supplementary File 1, section D.

### Local genomic scores support prediction across held-out populations

Mapping localizes statistical associations but does not evaluate their aggregate predictive value. We therefore estimated genome-wide marker effects jointly and summed their contributions within LD-defined blocks using population-adjusted localGEBV. The analysis used 698-756 phenotype-matched progeny per trait and the same 8,054 markers as mpQTL. In stratified random five-fold validation, the mean correlation was 0.832 for total predictions, including population effects, and 0.459 for genomic scores alone. When entire populations were held out, the mean correlation between genomic scores and BLUPs within the held-out populations was 0.470 across traits (range 0.382-0.580). Population adjustment improved this measure for 12 of 13 traits and raised its trait average from 0.445 to 0.470 (Supplementary Table 23; Supplementary Figure 10). Thus, the high total random-fold correlation should not be interpreted as genomic prediction accuracy for an unseen family.

The chromosome-16 block near 14.69 Mb ranked first for veraison and second for harvest and overlapped all three mapping frameworks, linking the recurrent mapping region to a predictive marker segment. Nevertheless, training-selected top-5% block subsets exceeded whole-genome scores for only two traits in nested leave-one-population-out validation; for veraison, the mean correlation increased from 0.489 to 0.521 (Supplementary Tables 24-26; Supplementary Figures 11 and 12). Local scores therefore support trait-specific investigation of compact marker panels, while whole-genome prediction remains the relevant benchmark.

## Discussion

The ResDur panel reveals agronomic architectures that combine recurrent genomic regions with substantial family specificity. Integrating linkage mapping, structure-aware association and genomic prediction shifts attention from marker counts to the regions and predictions that remain informative across analytical settings. The strongest example is the chromosome-16 region associated with veraison and harvest date, but the contrasting patterns among traits argue for different routes from discovery to breeding use.

### Genetic architecture across connected hybrid families

Within-family mapping and panel-wide association interrogate different components of the same breeding material. A QTL can be detectable in one cross and absent from another because its alleles are fixed or rare, because linkage phase or LD differs, or because family size and recombination limit detection. Background-dependent effects are another possibility, but family specificity alone does not demonstrate epistasis. The pronounced genomic and phenotypic differentiation among ResDur families makes differences in segregating alleles a plausible contributor to the observed variation in locus detection. Environment-specific recurrence, particularly for chromosome-16 phenology, identifies useful targets for testing effect stability, consistent with earlier phenology and compactness studies [33,39].

High broad-sense heritability is compatible with this family specificity. Heritability describes genetic relative to residual variance in a defined population and trial design; it does not specify the number, distribution or cross-family frequency of causal effects. Highly heritable flowering time in maize was associated with many small-effect loci [50], whereas barley combined major loci with additional predictive contributions from epistasis [51]. Wheat and rice likewise illustrate that heritable phenology can involve both recurrent and population-or environment-specific QTLs [52,53]. Thus, the ResDur results do not contradict high heritability, nor do they distinguish polygenicity from genetic-background interactions. The family-specific estimates accommodate unbalanced trials [54,55], but their interpretation remains conditional on the sampled families and environments.

Hybrid breeding adds a further dimension because resistance introgressions are recombined under selection for wine-grape performance. A recent theoretical perspective proposes that breeding constraints can restrict the accessible combinations of genotypes and phenotypes, potentially altering how interactions appear as additive or epistatic variance [56]. This is a possible interpretation of background dependence, not a mechanism demonstrated here. Allele frequency, linkage phase, LD and sampling remain more immediate explanations to test with phased parental haplotypes and common genotypes evaluated across environments.

### Recurrent regions and candidate biological processes

Comparisons with previous studies were restricted to matched or closely related traits, with coordinates harmonized by genome assembly and stable gene aliases (Supplementary Table 27). This distinction matters because chromosome-level recurrence can encompass different loci, whereas coincident refined intervals provide a more focused biological hypothesis. Candidate genes were prioritized by position and annotation; none is assigned a causal role on that basis alone.

Phenology provides the clearest within-study convergence but also illustrates the limits of chromosome-level comparison. Earlier work suggested at least two veraison-time QTLs on chromosome 16 [33]. The fine-mapped *Ver1* region contains the proposed candidate *VviERF027*, or *Vitvi16g00942* [57], at approximately 16.421 Mb in PN40024.v4.3. In contrast, the convergent ResDur signal is centred at 14.69-15.04 Mb, with LD-refined support at approximately 14.55-15.49 Mb. The broad family-50035 QTL encompasses *VviERF027*, but the compact family-50025 interval and the convergent mpQTL, GWAS and localGEBV signals do not. The association with both veraison and harvest is consistent with shared variation in ripening timing, but an adjacent or distinct locus remains plausible. *Vitvi16g00849* and *Vitvi16g00853* are therefore positional candidates for further investigation, rather than evidence that ResDur has resolved the causal basis of *Ver1*. Comparisons within family 50025 also show that QTL detection changes with the subsets, years and phenotyping protocols analyzed [58–60].

Berry and cluster architecture show a similar combination of recurrence and incomplete transfer. The ResDur chromosome-17 interval overlaps the cluster-architecture region of Richter et al. [39] and the berry-weight region of Doligez et al. [36], spanning evidence from *V. vinifera* and disease-resistant hybrid material. Previously proposed candidates include *CYP78A5*, *WRKY72*, *bHLH47* and *expansin A1*, suggesting hypotheses involving growth and cell expansion. However, the nearby localGEBV peak contains different positional candidates and does not resolve these genes causally. The chromosome-1 compactness interval and chromosome-14 cluster-trait intervals also lie near cluster-weight or cluster-length associations in other panels [61,62]. These are partial regional matches because the measured traits are related but not identical. Reports of poor quality-marker transfer across grape use types [63] reinforce the need to validate the haplotypes and trait definitions relevant to the target breeding population.

For acidity, the chromosome-13 region provides the strongest external positional match. The family-50015 titratable-acidity QTL and its refined GWAS interval overlap the pH, tartaric-acid and potassium:tartaric-acid region described by Duchêne et al. [37]. Their candidate *VIT_13s0019g04230* corresponds to *Vitvi13g00601*, encoding a V-type proton ATPase subunit G1 within the ResDur interval. This makes proton transport a testable biological hypothesis for the regional association. The same chromosome has lower-resolution support in an interspecific family [64]. On chromosome 17, broad ResDur acidity and pH intervals overlap the stable malate QTL of Reshef et al. [65], including the malate-dehydrogenase gene *Vitvi17g00607*. Chromosome-16 acidity intervals also contain the K^+^ efflux antiporter *Vitvi16g00860* associated with sugar and acid traits by Kadium et al. [66]. These observations nominate organic-acid metabolism and ion transport for functional investigation, without establishing shared causal alleles or effect directions.

Across these traits, comparisons between *V. vinifera* and hybrid backgrounds show both regional recurrence and differences in locus detection. Chromosome-13 acidity evidence spans the *V. vinifera* Riesling x Gewürztraminer cross [37], a *V. labruscana* x *V. vinifera* family [64] and ResDur. By contrast, support for chromosome-16 acidity and chromosome-17 malate regions comes predominantly from interspecific materials [65,66], while stable acidity QTLs in other crosses occur on chromosome 6 [67] or chromosomes 5 and 8 [68]. Hybrid breeding populations may therefore expose alleles or linkage configurations that are absent, fixed or poorly tagged in other germplasms. This does not establish wild-species origin of the causal variation; ancestry reconstruction and validation across vinifera and hybrid families are needed to separate introgression-specific effects from ordinary differences among segregating populations.

### Non-additive effects and associations across environments and traits

Alternative genotype encodings identified patterns not captured by additive dosage alone, extending approaches used in other perennial fruit crops [10,19] and other systems [11,69]. Such patterns can motivate hypotheses about dominance or heterozygote effects in highly heterozygous breeding material. The reduction in candidates after kinship-adjusted retesting emphasizes sensitivity to relatedness control and model specification; it cannot be attributed exclusively to confounding because the retest also changes statistical power. Retained contrasts require adequate genotype-class representation and replicated allelic directions before being interpreted as biological dominance or overdominance; the scans do not directly test epistasis.

The environmental analyses address another source of variation. Their recovery of association evidence under random-effect and local-score models, despite no significant common fixed-effect marker, motivates testing whether associations depend on environmental context. Failure to pass the fixed-effect threshold does not establish absence of a common effect. Climate-associated flowering signals and the chromosome-16 pH example provide focused hypotheses for developmental and compositional responses. However, the random-effect test jointly evaluates mean effects and heterogeneity, and neither it nor climate meta-regression is a direct variance-component test of genotype-by-environment interaction. Changing family composition and correlated annual climate descriptors can contribute to the observed associations. Balanced trials and covariates defined over relevant developmental periods are therefore needed to distinguish climatic modulation from population differences.

Cross-trait QCH results connect the observed trait correlations to shared genomic association patterns. Their concentration in phenology and yield-related traits makes the selected regions relevant to correlated breeding responses. Part of the shared signal among cluster-yield traits, however, reflects trait construction, because mean cluster weight is the ratio of cluster weight per plant to cluster number and yield per area is derived from yield per vine. The sensitivity analyses identify the regions whose multi-trait support does not depend on this construction: 53 of the 54 regions were retained without mean cluster weight and 38 without either derived trait, including the chromosome-16 phenology region. Loss of a region does not by itself indicate an artifact, because removing traits also changes the composite hypothesis and the power of the test; conversely, a common signal can arise from one variant affecting several traits or from linked variants with distinct effects, so retained regions are not proof of biological pleiotropy. Joint phenotyping, phased-haplotype analysis and functional evidence will be needed to resolve the alternatives. For breeding, the immediate implication is to evaluate favorable and unfavorable correlated responses when validating a region, rather than to assume that all associated traits can be improved independently.

### From genomic evidence to breeding decisions

The locally concordant regions provide priorities for haplotype validation. The chromosome-16 phenology region is the clearest target, followed by the compactness and cluster-weight regions on chromosomes 1 and 14. Validation should establish segregation, within-family allelic direction, linkage to favorable variants and stability across independent crosses and environments. Haplotypes with reproducible effects and acceptable linkage drag could advance to early-generation marker-assisted selection alongside resistance-gene pyramids (*Rpv*, *Ren/Run* and *Rgb* resistance loci). The present lead SNPs are starting points for this work, not immediately deployable assays.

localGEBV adds a complementary perspective by estimating marker effects jointly and aggregating them within LD-defined segments [70]. Convergence of a highly ranked block with mapping evidence increases its value as a validation target, whereas localGEBV-only blocks remain prediction-derived hypotheses. Total random-fold prediction concerns progeny from crosses already represented in training, whose population means can be estimated. By contrast, the held-out-population results assess ranking of progeny within an unseen family, not prediction of that family’s mean. Their variation among families is consistent with heterogeneous transfer of genomic information, potentially reflecting allele frequencies, linkage phase or background effects as well as sampling variation.

Selected-block prediction was competitive for some traits but not uniformly superior to whole-genome scores. Restricting the marker set can improve interpretability and suggest reduced-cost panels, while losing distributed polygenic information. Whole-genome prediction should therefore remain the benchmark. A practical strategy combines family-aware marker-assisted selection for validated recurrent regions with multi-trait genomic prediction for aggregate performance. Multi-environment models, major-locus effects or non-additive kernels should be added when they improve externally validated prediction, with climatic covariates matched to developmental periods. Selection indices can then balance phenology, yield, berry composition and resistance while monitoring correlated responses.

### Limitations and next steps

The operational breeding design combines fixed plots, unequal family sizes and incomplete overlap of genotypes among environments. Mixed models and structure-aware analyses address these constraints but cannot fully separate environmental and background effects. Cross-method agreement remains internal to overlapping material, and localGEBV uses unphased dosage blocks rather than sequence-resolved haplotypes. The next priorities are phased-haplotype validation and fine mapping of recurrent regions in independent crosses, balanced trials with common genotypes, and functional data at relevant phenophases [71]. External validation and iterative updating of genomic prediction, together with economic weighting of traits, will determine how these locus and segment priorities translate into practical breeding gains.

## Conclusion

The ResDur breeding panel combines recurrent agronomic loci with substantial family- and environment-dependent association patterns. Integrating within-family mapping, population-adjusted multiple-population mapping and GWAS identifies a focused set of regions for validation, led by chromosome-16 veraison and harvest timing. Cross-trait analyses highlight potential correlated responses, while localGEBV links genomic segments to prediction across held-out families. These complementary results support phased-haplotype validation for targeted marker-assisted selection and externally validated genomic prediction for polygenic traits. Together, they provide a practical foundation for combining durable disease resistance with appropriate phenology, yield and berry composition in future grapevine cultivars.

## Materials & methods

### Plant material

All plant material originated from step 2 of the INRAE-ResDur breeding program (Figure 6). The analyzed panel contained 1,081 genotypes from 95 crosses designed to pyramid resistance to downy and powdery mildews while recovering agronomic and sensory quality. The six largest families (42050, 50001, 50013, 50015, 50025 and 50035) were retained for family-specific QTL and heritability analyses. The pedigree is connected: Bronner is a parent of 42050 and 50013 and a grandparent of 50025 and 50035, and two additional families contain a parent selected from family 50001 (Supplementary Figure 4). For the population-structure visualization only, the analysis set included *V. vinifera* accessions and other interspecific hybrids alongside selected ResDur parents and progeny (344 individuals in total).

**Figure 6.**
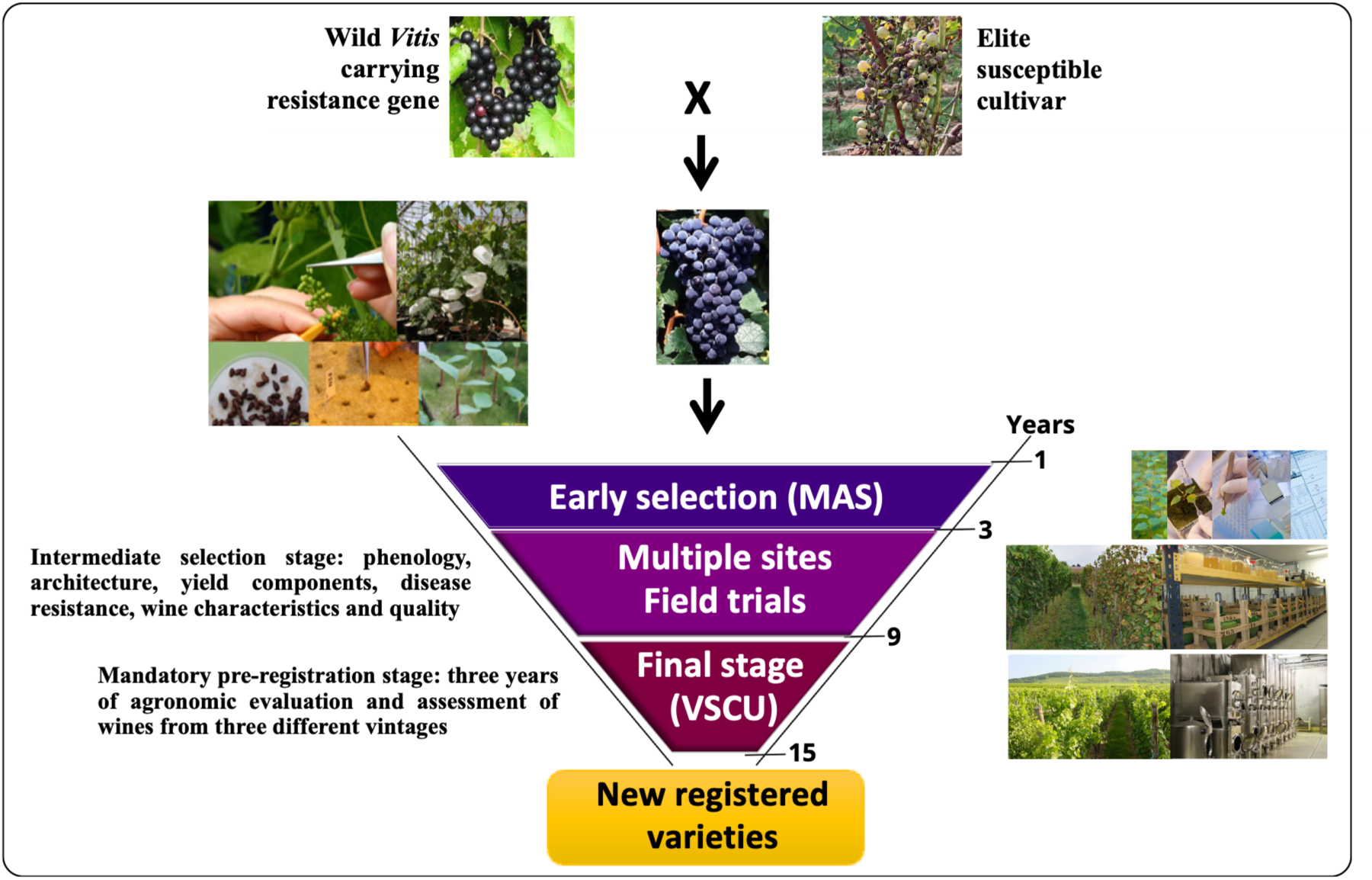
INRAE-ResDur grapevine breeding scheme. Parental genotypes are selected for disease-resistance genes and favorable agronomic and organoleptic traits. Progenies from multiple crosses undergo early greenhouse screening by marker-assisted selection (MAS) to retain seedlings carrying the targeted pyramids of resistance genes; at this stage, each genotype is represented by a single seedling. Selected genotypes subsequently enter intermediate field trials, with four to five vines per genotype evaluated across multiple sites for phenology, architecture, yield, resistance and wine quality. During the final mandatory stage for assessment of Value for Sustainable Cultivation and Use (VSCU), each candidate variety is grown at two contrasting sites, with at least 90 vines per genotype. Agronomic performance is evaluated over three years, together with wines produced from three vintages.

### Phenotypic data collection

Phenotypic observations were collected between 2006 and 2024 at Angers, Bordeaux, Colmar and Montpellier in France and Pully in Switzerland. Material entered the network in successive breeding phases, so genotypes were not evaluated in a single balanced design and replication varied among traits, years and sites. Chardonnay and Merlot were included as reference cultivars in the field trials. Trait-specific sample sizes and numbers of years, sites and site x year environments are reported in Supplementary Tables 3 and 4.

Figure 2 reference means were calculated from the raw intermediate-stage observations of the 12 varieties registered from ResDur phases 1 and 2. Variety identifiers, observation coverage and reference means are given in Supplementary File 1 and the Supplementary Excel File. These descriptive values were not used to estimate family effects, which were assessed on the environment-adjusted whole-panel BLUPs.

Dozens of traits were initially measured for each individual, but only those with limited missing data were retained for analysis. Thirteen traits were selected and grouped into three categories:

- Phenological traits: date of 50% of budbreak (BUD_DATE), date of 50% flowering (FLO_50), date of 50% veraison (VER_50), and harvest date (HARVEST_DATE).
- Yield components: number of clusters per plant (NB_CLUST_PLANT), total cluster weight per plant (YIELD_PLANT), mean cluster weight (SCLUST_W, calculated from total weight and number of clusters), mean berry weight (SBER_W_g), calculated from a standard berry sample, yield per square meter (YIELD_OIV_504), and cluster compactness (MORPHO_OIV_204, scored on a 1-5 scale).
- Berry quality traits: berry titratable acidity (BER_TA_g), berry pH (BERRY_pH), and berry sugar content (TSS, measured on a standard berry sample collected 42 days after veraison).

### Environment description

An environment was defined as a site x year combination. Daily minimum and maximum temperature, rainfall, Penman-Monteith reference evapotranspiration and global radiation were summarized annually as mean minimum temperature, mean maximum temperature, mean temperature, cumulative rainfall, mean evapotranspiration and mean radiation. The six descriptors for 52 trial environments (2006-2024) were centered and scaled before PCA.

### Genotyping

ResDur genotypes and the ampelographic collection were genotyped by *ApeKI* genotyping-by-sequencing [72] on an Illumina NovaSeq 6000 machine. Reads were demultiplexed with Stacks [73,74], aligned to PN40024.v4 [26] with BWA [75], and variants were called with GATK [76]. VCFtools and BCFtools were used for filtering [77,78]. For whole-panel analyses, we retained biallelic SNPs with depth ≥3, ≤10% missing genotypes and MAF ≥0.01; genotype calls were imputed before association analysis. LD-clumped SNP panels were used for the population-structure analyses [79]. The exact map match across populations and within-trait MAF filter yielded 137,285-140,181 SNPs in the structure-adjusted BLINK analyses. For family QTL mapping, filters were depth ≥3, ≤5% marker missingness, ≤20% sample missingness, removal of individuals with extreme Mendelian error, and removal of distorted markers at the stated FDR threshold (Supplementary Table 28). Of approximately 11.8 million raw variants called in the combined ResDur and ampelographic panels, 189,426 biallelic SNPs passed depth, missingness and minor-allele-frequency filters. The Figure 1C subset contained 62,621 LD-clumped SNPs (62,619 polymorphic), whereas the GWAS-covariate DAPC used 36,726 SNPs.

### Population structure and family differentiation

For Figure 1C, DAPC [79,80] was fitted to a subset of 344 individuals (194 *V. vinifera* accessions, 122 other interspecific hybrids, 15 ResDur parents and 13 progeny) genotyped at 62,619 polymorphic LD-clumped SNPs. The fitting groups were seven clusters inferred from the genotypes by K-means clustering on the principal components, selected by the Bayesian information criterion over 1-40 clusters. The number of principal components retained for the discriminant step (nine, explaining 31.1% of the variance) was chosen by stratified cross-validation of cluster assignment, and all six discriminant axes were retained; the four biological categories were used only to annotate the plot. Settings, eigenvalues and diagnostics are given in Supplementary File 1, section A.

The ancestry covariates used in the GWAS came from a separate DAPC of 1,389 genotypes and 36,726 LD-clumped SNPs, with 16 inferred clusters, 15 principal components selected by a-score optimization and five retained discriminant axes; its first five discriminant axes (LD1-LD5) were used as ancestry covariates in all additive, non-additive and environment-specific GWAS. Pairwise family differentiation was quantified from the un-clumped panel using the Hudson FST ratio of summed numerator and denominator terms across informative SNPs. Phenotypic family differentiation was quantified for each BLUP trait by the one-way-model η^2^ and checked with Kruskal-Wallis tests.

### Statistical analysis

Pearson correlations among the 13 agronomic traits were calculated from raw phenotypic observations with complete measurements for all traits (Figure 3). Complete-case selection from the available dataset retained 1,671 observations from 863 ResDur progeny evaluated at Angers, Bordeaux and Colmar during 2012-2022. To assess sensitivity to repeated observations, correlations were recalculated after averaging these same records within genotype, with equal weight for each of the 863 genotypes, and among their existing whole-panel BLUPs, all complete for the 13 traits. For the pooled raw correlations, 2,000 bootstrap replicates resampled genotypes with replacement and retained all observations of each sampled genotype; the 2.5th and 97.5th percentiles defined 95% intervals. This bootstrap preserves within-genotype dependence but does not remove environmental or family effects. In the complete-case records, yield per area equalled yield per vine divided by the plot-specific area per vine (2 or 2.329 m^2^); the two yield variables were therefore treated as redundant in the interpretation and in the QCH sensitivity analysis.

Best linear unbiased predictors (BLUPs) were obtained separately for the complete ResDur panel and for each of the six largest biparental populations. Within each trait and analysis set, extreme raw observations were screened with Grubbs’ test, and transformations selected with bestNormalize were applied when required [81,82]. The maximal candidate linear mixed model for the complete panel was:

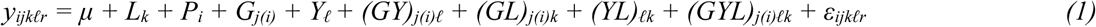

Here i indexes population, j genotype within population, k site, ℓ year and r repeated observations when present. μ is the intercept; L is the fixed site effect; P, G and Y are the population, genotype and year effects; and GY, GL, YL and GYL denote the corresponding interaction terms. Population, genotype, year and the interaction terms were fitted as random effects, each with its own variance component; ε is the residual error. Random effects and residuals were assumed normally distributed with mean zero. This equation defines the maximal candidate structure. Unsupported terms were removed within trait, and the selected trait-specific models are reported in Supplementary Table 3.

Candidate random-effect structures sharing the same fixed-effect specification were compared by Akaike information criterion (AIC). Comparisons involving the fixed site term were made by maximum likelihood; the selected model was then fitted by restricted maximum likelihood for variance-component estimation and BLUP extraction. For each genotype, the fitted population effect was added to the nested-genotype conditional mode to obtain the whole-panel genotypic value used in panel-wide analyses. Adding a constant for presentation, where applicable, does not affect QTL or GWAS tests. Because the whole-panel BLUPs came from strongly structured families, the subsequent panel-wide analyses explicitly fitted ancestry, population and/or kinship as appropriate.

For each of the six largest biparental populations, the maximal candidate model was:

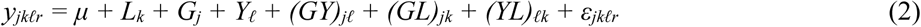

Here j indexes genotype, k site, ℓ year and r repeated observations when present. μ is the intercept; L is the fixed site effect; ε is the residual error; and G, Y, GY, GL and YL are candidate random effects. The same distributional assumptions were used as in the complete-panel model. The selected set of effects varied among trait-by-population analyses and is reported in Supplementary Table 4. For each trait and biparental population, the final model was selected by sequential elimination of unsupported effects according to AIC while retaining the genotype term required for BLUP estimation. The selected model was fitted by restricted maximum likelihood, and genotype conditional modes were extracted as BLUPs.

### Broad-sense heritability

Broad-sense heritability was estimated only within the six largest biparental populations because the complete-panel design combined distinct populations with highly unequal coverage across years and sites. For each estimable trait-by-population combination, we used the generalized Piepho-Möhring heritability for unbalanced trials [54,55]:

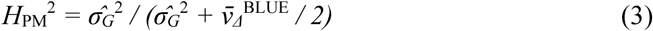

where the genetic-variance term is the estimated among-genotype variance and the BLUE-difference term is the arithmetic mean of the estimated variances of all pairwise differences between genotype BLUEs. This formulation accommodates unequal replication and unequal precision among genotype means. Calculations were performed in R using the inti implementation [83].

### QTL analyses

Genetic maps were built separately for families 42050, 50001, 50013, 50015, 50025 and 50035 with Lep-MAP3 [84], using 16,223-25,295 markers. Stacks-exported JoinMap files [85] were filtered for Mendelian error and segregation distortion with FDR control. Lep-MAP3 module “ParentCall2” assigned parental genotypes, “SeparateChromosomes2” defined linkage groups and “OrderMarkers2” ordered markers and estimated distances. Single-family QTL scans used Haley-Knott regression in R/qtl [86] and 4,000 permutations to establish a 5% genome-wide threshold. Family BLUPs were analyzed in all six families; raw environment-specific values were additionally analyzed in 42050, 50025 and 50035. For the connected-family map, VCFs from 12 families containing at least 20 progenies were filtered under the same family-level rules, merged and analyzed jointly with Lep-MAP3. This led to a composite-map of 792 progeny and 62,202 markers, whose marker density was visualized with LinkageMapView v2.1.2 [87].

### Population-adjusted multiple-population QTL mapping

Multiple-population QTL mapping used mpQTL version 0.6.2 [88] on 772 progenies with 12 population labels, 8,054 composite-map markers and the 13 whole-panel BLUP traits. Genotypes were diploid alternative-allele dosages with no missing calls in the analyzed matrix; no genotype imputation was performed in this scan. The mixed model included the realized genomic kinship matrix and 12 fixed numerical population indicators. Marker tests used the package’s approximate P3D/EMMAX implementation (approximate = TRUE), and the Li-Ji effective-test threshold was P = 3.44E-5 (1,490 effective tests). Within each chromosome, significant leads separated by more than 5 cM were retained. A nominal support interval comprised markers within ±10 cM whose -log10(P) was no more than 1.5 below the lead; these marker-defined spans are heuristic support intervals, not formal confidence intervals. Standardized dosage effects and approximate 95% Wald intervals were obtained from the marker coefficient, trait standard deviation and one-degree-of-freedom F statistic. Identical dosage vectors were grouped to avoid counting duplicate marker patterns as independent loci. Physical support spans were obtained from PN40024.v4 marker coordinates. For three-framework integration, a pooled support span and a GWAS LD interval were locally concordant when they overlapped or were separated by ≤500 kb inside the same single-family QTL interval. Population sizes, model diagnostics, complete peak and effect tables, and non-estimable tests are reported in Supplementary Tables 10-12 and 18 and Supplementary File 2.

### Population-adjusted local genomic estimated breeding values

Local genomic estimated breeding values were calculated with HapSelect version 0.1.0 and rrBLUP version 4.6.3 [70] for the same 772 progeny, 12 population labels, 8,054 complete-genotype composite-map markers and 13 whole-panel BLUP traits used in mpQTL. Trait-specific phenotype availability ranged from 698 to 756 genotypes. Marker effects were estimated by rrBLUP after centering dosages by their training-set marker means. The primary model fitted 12 population indicators as fixed effects, with population 42048 as the reference; the HapSelect package-default intercept-only solution was retained as an implementation and structure-sensitivity benchmark. Pairwise r^2^ was calculated within chromosome after subtracting population-specific marker means, thereby reducing LD attributable solely to population mean differences. The package’s pooled-panel PLINK LD was retained as a sensitivity analysis. Marker blocks were defined with the flanking algorithm and complete marker coverage. A grid of r^2^ thresholds (0.10, 0.20, 0.30 and 0.50) and flank tolerances (0-4) was evaluated for veraison; the primary setting used within-population r^2^ ≥ 0.30 with one tolerated low-LD flank, whereas pooled r^2^ ≥ 0.20 with tolerance 4 provided the manual-reference sensitivity. Blocks are analytical LD-defined marker segments, not sequence-phased haplotypes. For individual i and block b, localGEBV was the sum over markers in b of centered dosage multiplied by its fitted marker effect. Blocks were ranked by the variance of localGEBV across genotypes. Prediction was evaluated by population-stratified random five-fold cross-validation and by leave-one-population-out validation for populations with at least 20 phenotyped progenies. Random-fold results report both total predictions, including the fitted population effect, and the genomic component. Leave-one-population-out results report the genomic component because the held-out population intercept cannot be estimated from training data. Pearson correlations with observed BLUPs were calculated separately across the mixed-population individuals in each random validation fold or within each eligible held-out population, then summarized as unweighted means and standard deviations. For selected-block validation, marker effects and block-variance ranks were recomputed within each training set before selecting the top 1%, 5% or 10% of blocks; fixed marker-only block boundaries were then used to score the held-out population. Whole-genome rrBLUP provided the comparator.

### GWAS for additive effects

Additive GWAS used trait-specific whole-panel BLUPs from 899 to 968 genotypes. Genotypes were encoded as 0, 1 and 2 alternative alleles; exact PN40024.v4 map matching and within-trait MAF ≥0.01 left 137,285-140,181 SNPs (Supplementary Table 14). BLINK, MLMM and MM4LMM were fitted to the same trait-specific inputs. BLINK included LD1-LD5 as fixed ancestry covariates; MLMM and MM4LMM fitted both a trait-specific genomic kinship matrix and LD1-LD5. BLINK was implemented in GAPIT [8,89]. BLINK iteratively selected marker cofactors through its Bayesian-information and linkage-disequilibrium procedure. Significance was assessed with a trait-specific 5% Bonferroni threshold. MLMM included a trait-specific genomic kinship matrix and LD1-LD5 as unpenalized fixed covariates, then iteratively added the most significant SNP as a cofactor [7]. Forward selection was limited to 10 marker cofactors and the selected multilocus model followed the package’s Bonferroni stopping criterion. MM4LMM was fitted marker by marker with the same trait-specific kinship matrix and a fixed-effect design containing an intercept and LD1-LD5 [90]. Manhattan and Q-Q plots and complete summary statistics are provided in Supplementary File 3 and the Supplementary Excel File. For cross-method summaries, SNPs were matched by PN40024.v4 coordinate and nearby associations were grouped only when their physical positions were locally concordant.

### GWAS for non-additive effects

For exploratory non-additive scans, hard-call dosages 0/1/2 were recoded as dominant (0 versus 1/2), recessive (0/1 versus 2) or overdominant (0/2 versus 1) contrasts (Supplementary Table 29) [10,19]. Markers first had to satisfy original-allele MAF ≥0.01 and then retain at least 10 individuals in both encoded classes. BLINK was fitted with LD1-LD5 and the same trait-specific 5% Bonferroni rule used for additive scans. Because the ancestry-only scans retained residual genomic inflation for some trait-contrast combinations, every Bonferroni candidate was retested marker by marker with MM4LMM using the trait-specific genomic kinship matrix and LD1-LD5. A candidate was retained only when its K + LD1-LD5 P-value also passed the original genome-wide Bonferroni threshold calculated over all eligible markers for that trait and encoding.

### Meta-GWAS analysis

For each trait, additive BLINK GWAS including the first five discriminant axes (LD1-LD5) were fitted separately in site-by-year environments containing at least 100 matched genotypes. Replicate observations for a genotype within an environment were averaged before association analysis. Only genuine site-by-year scans were included in metaGE [13]. Summary statistics were restricted to SNPs shared across all eligible environments for the trait. For marker m in environment k, the signed association Z-score was calculated from the two-sided GWAS P value pₘₖ and estimated allelic effect β^ₘₖ as

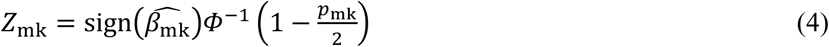

where Φ⁻¹ denotes the quantile function of the standard normal distribution. The magnitude of Zₘₖ therefore reflects the strength of association, whereas its sign reflects the direction of the estimated marker effect. Let Zₘ = (Zₘ₁, …, Zₘₖ)ᵀ denote the vector of signed Z-scores across the K eligible environments. Dependence among environment-specific GWAS arising from overlapping or related genotype panels was represented by the inter-environment correlation matrix Σ. This matrix was estimated from null-like markers using metaGE.cor, with the posterior-H_1_ filtering threshold set to 0.8. The fixed-effect model was

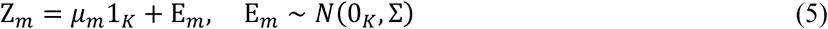

where μₘ is the common marker effect across environments and Eₘ is the vector of correlated sampling errors. The fixed-effect procedure tested H₀: μₘ = 0.

To allow marker effects to vary among environments, the random-effect model was

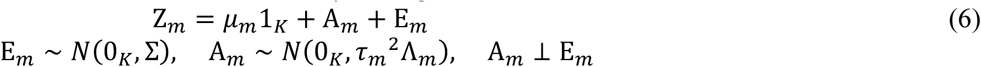

where Aₘ represents environment-specific deviations from the mean marker effect and τₘ^2^ measures between-environment heterogeneity. The matrix Λₘ is the correlation matrix of the random marker effects across environments. Following the metaGE model, Λₘ was assumed to be common to all markers (Λₘ = Λ), and Λ was set equal to the inter-environment correlation matrix Σ. Under this fitted assumption, Var(Zₘ | μₘ) = Σ + τₘ^2^Λ = (1 + τₘ^2^)Σ. The random-effect likelihood-ratio test evaluated

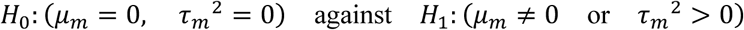

using a likelihood-ratio statistic whose null distribution was the mixture

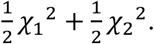

To complement marker-wise tests, spatially clustered association evidence was evaluated separately from the fixed- and random-effect meta-analysis P-values using metaGE.lscore with the local-score tuning parameter ξ = 3. This procedure accumulates evidence from neighboring markers and identifies genomic zones in which several individually modest signals jointly support association. For each centered environmental covariate vector x, metaGE tested whether the environment-specific marker effects covaried with that descriptor using

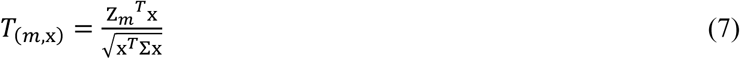

which follows a standard normal distribution under the null hypothesis of no covariance between the marker effect and the environmental descriptor. The corresponding two-sided P-value was

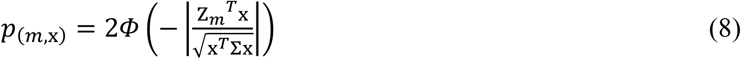

Six harmonized annual climate descriptors were tested separately. Within each trait and test, marker-wise fixed-effect, random-effect and meta-regression P-values were controlled at a 5% false discovery rate using the metaGE Benjamini-Hochberg procedure incorporating an estimated null proportion. Fixed-effect, random-effect, local-score and climate meta-regression evidence were reported separately. These analyses test association-level heterogeneity across environments. Because family composition varied among site-by-year environments, significant meta-regression results were interpreted as associations between marker-effect variation and environmental descriptors rather than as definitive causal genotype-by-environment interactions [13].

Multiple-testing control was matched to the inferential purpose. Single-family QTL mapping used 5% genome-wide permutation thresholds; single-trait additive and non-additive GWAS used 5% Bonferroni family-wise error-rate control; population-adjusted mpQTL mapping used the Li-Ji effective-test threshold; marker-wise metaGE tests used the 5% false-discovery-rate procedure described above; and the cross-trait QCH analysis used 5% false-discovery-rate control.

For the cross-trait analysis, P-values from the 13 LD1-LD5-adjusted additive BLINK scans were intersected by exact PN40024.v4 marker identifier, leaving 136,062 SNPs shared across all traits. The QCH method, implemented in the qch package [27], fitted a Gaussian copula using its memory-efficient expectation-maximization algorithm with a maximum squared parameter-change stopping threshold of 10^−4^. Dependence among trait-specific tests was thereby accounted for while testing the composite alternative that a SNP was associated with at least two traits [27]. Marginal-kernel precursor estimates from the trait-specific full GWAS are reported in Supplementary Table 30; estimates on the common SNP universe are provided in Supplementary File 1. Additional supporting documentation is provided for the phenotyping-environment inventory (Supplementary Table 31), the localGEBV LD-defined marker-block sensitivity audit (Supplementary Table 32), the rrBLUP marker-effect model summary (Supplementary Table 33), the multiple-testing and evidence-threshold guide (Supplementary Table 34), the descriptive 250-kb marker-climate proximity groups (Supplementary Table 35), and the trait-specific inputs for the 303 site-by-year ancestry-adjusted GWAS included in metaGE (Supplementary Table 36).

Sensitivity analyses retained the same 136,062 SNPs and the composite alternative of association with at least two of the retained traits. QCH was refitted with the Gaussian copula after excluding mean cluster weight (12 traits) and after additionally excluding yield per area (11 traits), with the same 5% posterior-FDR procedure. These fits used the package-default stopping threshold of 10⁻⁶ for the EM algorithm, stricter than the threshold of the primary fit; a 13-trait control fitted with the same settings reselected all 270 markers of the primary catalogue (Supplementary File 1, section D). A primary region was considered retained only if at least one of its originally selected markers was reselected; new signals within 250 kb were tabulated separately. As in the primary fit, marginal density estimation reached its iteration cap for flowering and veraison. Scenario-specific selections are listed in the Supplementary Excel File.

### Physical-coordinate evidence integration and candidate-gene annotation

Single-family QTL intervals, population-adjusted mpQTL support spans and GWAS associations were compared on PN40024.v4 physical coordinates. For each family-QTL interval, a candidate GWAS lead was chosen to maximize local support among BLINK+LD1-LD5, MLMM K+LD1-LD5 and MM4LMM K+LD1-LD5. Linkage disequilibrium was then calculated among phenotype-matched genotypes within ±500 kb of that lead after subtracting population-specific marker means. The GWAS interval comprised markers with r^2^ ≥ 0.20, clipped to the family-QTL interval; support from at least two GWAS models was called multi-model only when their significant markers belonged to the lead LD set. A family QTL, an mpQTL support span and a GWAS LD interval were locally concordant when the latter two overlapped or were separated by ≤500 kb inside the family-QTL interval. These analyses reuse overlapping individuals and were interpreted as triangulation rather than independent replication. Gene models overlapping the refined interval were obtained from the PN40024.v4.3 GFF3. Functional descriptions, Gene Ontology terms and InterPro identifiers were matched by stable gene identifier from the available PN40024.v4.1 Blast2GO table. Candidate priority combined physical overlap or distance with a prespecified trait-relevant keyword screen.

## Acknowledgements and Funding

This work was supported by funding through the IB2022_SelGen-ResDur project of the INRAE BAP division and the OIV research grant for the PhD thesis of Clémentine Borrelli. The INRAE-ResDur program was supported by funding from the “Innovation Variétale et Diversité (IVD)” strategic plant breeding supporting program of the INRAE BAP division (from 2021 to 2025), CASDAR FranceAgriMer grants for projects “ViRéVATE” and “INNOVRES” and a contribution from CTNSP. We thank all the staff of the INRAE UEAV experimental unit of Colmar for their support and the management of the vineyards. We also thank the INRAE experimental units, UE Vigne in Bordeaux (Dominique Forget and Laurent Delière), UE Horti in Angers (Gérard Barbeau), UE Pech-Rouge (Hernan Ojeda) and UE Vassal (Cécile Marchal) in Montpellier for their assistance mainly in data collection for the intermediate selection stage. We also thank IFV (Laurent Audeguin and Pascal Bloy) and the “Partenaires de la sélection Vigne” (Géraldine Uriel, Jean-Michel Desperrier, Arnaud Ritton, François Berud, Olivier Jacquet, Bernard Genevet, Didier Viguier, Olivier Yobrégat, Thierry Dufourcq, Cédric Elia, Etienne Goulet) for their involvement in the field trial network dedicated to the final selection stage of the INRAE-ResDur program. We thank Amandine Velt for her assistance in the use of the unit bioinformatics server. We thank Morgane Roth, Hélène Muranty, Vincent Segura, Loïc Le Cunff, and David Pot for their insightful guidance and constructive feedback as members of Clémentine Borrelli’s PhD committee.

## Author contributions

DM and CS designed and managed the INRAE-ResDur program; EP, GB, NJ, and AUF conducted laboratory work and marker-assisted selection; CS, VD, and GA supervised the experiments and carried out phenotypic observations; CO managed the crosses and contributed to phenotypic observations; SWM, MCL, and MAD performed disease-resistance assays; SC, SV, and LL conducted phenotypic observations; JSR and JLS coordinated the Swiss Agroscope experiments within the co-breeding agreement between INRAE and Agroscope; ED contributed to the conceptualization of the INRAE-ResDur program and its associated research; CB and KA performed data analysis and wrote the manuscript. KA supervised the overall research and PhD thesis of CB. All authors reviewed, revised, and approved the final manuscript.

## Data availability

The raw phenotypic data, genetic maps and ResDur GBS variant dataset are available from Recherche Data Gouv at https://doi.org/10.57745/GZVK6R.

## References

1. Hospital F, Moreau L, Lacoudre F, Charcosset A, Gallais A. More on the efficiency of marker-assisted selection. Theoretical and Applied Genetics. 1997;95(8):1181–1189. doi:10.1007/s001220050679

2. Paterson AH, Lander ES, Hewitt JD, Peterson S, Lincoln SE, Tanksley SD. Resolution of quantitative traits into Mendelian factors by using a complete linkage map of restriction fragment length polymorphisms. Nature. 1988;335(6192):721–726. doi:10.1038/335721a0

3. 3. Dash M, Mishra A. QTL Mapping: Principle, Approaches, and Applications in Crop Improvement. Apple Academic Press; 2024:31–60.

4. Würschum T. Mapping QTL for agronomic traits in breeding populations. Theoretical and Applied Genetics. 2012;125(2):201–210. doi:10.1007/s00122-012-1887-6

5. Ozaki K, Tanaka T. Genome-wide association study to identify SNPs conferring risk of myocardial infarction and their functional analyses. Cellular and Molecular Life Sciences. 2005;62(16):1804–1813. doi:10.1007/s00018-005-5098-z

6. Tibbs Cortes L, Zhang Z, Yu J. Status and prospects of genome-wide association studies in plants. The Plant Genome. 2021;14(1):e20077. doi:10.1002/tpg2.20077

7. Segura V, Vilhjálmsson BJ, Platt A, et al. An efficient multi-locus mixed-model approach for genome-wide association studies in structured populations. Nature Genetics. 2012;44(7):825–830. doi:10.1038/ng.2314

8. Huang M, Liu X, Zhou Y, Summers RM, Zhang Z. BLINK: a package for the next level of genome-wide association studies with both individuals and markers in the millions. GigaScience. 2019;8(2):giy154.

9. Liu X, Huang M, Fan B, Buckler ES, Zhang Z. Iterative Usage of Fixed and Random Effect Models for Powerful and Efficient Genome-Wide Association Studies. PLOS Genetics. 2016;12(2):e1005767. doi:10.1371/journal.pgen.1005767

10. Pérez de los Cobos F, Coindre E, Dlalah N, et al. Almond population genomics and non-additive GWAS reveal new insights into almond dissemination history and candidate genes for nut traits and blooming time. Horticulture Research. 2023;10(10):uhad193. doi:10.1093/hr/uhad193

11. Xue Y, Liu S, Li W, et al. Genome-Wide Association Study Reveals Additive and Non-Additive Effects on Growth Traits in Duroc Pigs. Genes. 2022;13(8):1454. doi:10.3390/genes13081454

12. Guindo-Martínez M, Amela R, Bonàs-Guarch S, et al. The impact of non-additive genetic associations on age-related complex diseases. Nature Communications. 2021/04/23 2021;12(1):2436. doi:10.1038/s41467-021-21952-4

13. De Walsche A, Vergne A, Rincent R, et al. metaGE: Investigating genotype x environment interactions through GWAS meta-analysis. PLOS Genetics. 2025;21(1):e1011553. doi:10.1371/journal.pgen.1011553

14. Wong C, Ng JY, Sio YY, Chew FT. Genetic determinants of skin ageing: a systematic review and meta-analysis of genome-wide association studies and candidate genes. Journal of Physiological Anthropology. 2025;44(1):4. doi:10.1186/s40101-025-00384-9

15. Kang J, Ahn K, Oh J, et al. Identification of Endometriosis Pathophysiologic-Related Genes Based on Meta-Analysis and Bayesian Approach. International Journal of Molecular Sciences. 2025;26(1):424. doi:10.3390/ijms26010424

16. Cáceres P, Lopéz P, Garcia B, et al. Meta-analysis of GWAS for sea lice load in Atlantic salmon. Aquaculture. 2024;584:740543. doi:10.1016/j.aquaculture.2024.740543

17. Bouwman AC, Daetwyler HD, Chamberlain AJ, et al. Meta-analysis of genome-wide association studies for cattle stature identifies common genes that regulate body size in mammals. Nature Genetics. 2018;50(3):362–367. doi:10.1038/s41588-018-0056-5

18. Zhao J, Sauvage C, Zhao J, et al. Meta-analysis of genome-wide association studies provides insights into genetic control of tomato flavor. Nature Communications. 2019;10(1):1534. doi:10.1038/s41467-019-09462-w

19. Serrie M, Segura V, Blanc A, et al. Multi-environment GWAS uncovers markers associated to biotic stress response and genotype-by-environment interactions in stone fruit trees. Horticulture Research. 2025;12(7)doi:10.1093/hr/uhaf088

20. van Rheenen W, Peyrot WJ, Schork AJ, Lee SH, Wray NR. Genetic correlations of polygenic disease traits: from theory to practice. Nature Reviews Genetics. 2019;20(10):567–581. doi:10.1038/s41576-019-0137-z

21. Scott MF, Ladejobi O, Amer S, et al. Multi-parent populations in crops: a toolbox integrating genomics and genetic mapping with breeding. Heredity. 2020/12/01 2020;125(6):396–416. doi:10.1038/s41437-020-0336-6

22. Dong Y, Duan S, Xia Q, et al. Dual domestications and origin of traits in grapevine evolution. Science. 2023;379(6635):892–901. doi:10.1126/science.add8655

23. Töpfer R, Hausmann L, Harst M, Maul E, Zyprian E, Eibach R. New horizons for grapevine breeding. *Fruit*, Vegetable and Cereal Science and Biotechnology. 2011;5(2):79–100.

24. Jaillon O, Aury J-M, Noel B, et al. The grapevine genome sequence suggests ancestral hexaploidization in major angiosperm phyla. nature. 2007;449(7161):463.

25. Shi X, Cao S, Wang X, et al. The complete reference genome for grapevine (Vitis vinifera L.) genetics and breeding. Horticulture Research. 2023;10(5):uhad061. doi:10.1093/hr/uhad061

26. Velt A, Frommer B, Blanc S, et al. An improved reference of the grapevine genome reasserts the origin of the PN40024 highly homozygous genotype. G3 Genes|Genomes|Genetics. 2023/03/26 2023:jkad067. doi:10.1093/g3journal/jkad067

27. De Walsche A, Gauthier F, Boissot N, Charcosset A, Mary-Huard T. Large-scale composite hypothesis testing procedure for omics data analyses. NAR Genomics and Bioinformatics. 2025;7(3)doi:10.1093/nargab/lqaf118

28. Töpfer R, Trapp O. A cool climate perspective on grapevine breeding: climate change and sustainability are driving forces for changing varieties in a traditional market. Theoretical and Applied Genetics. 2022;135(11):3947–3960. doi:10.1007/s00122-022-04077-0

29. Merdinoglu D, Schneider C, Prado E, Wiedemann-Merdinoglu S, Mestre P. Breeding for durable resistance to downy and powdery mildew in grapevine. OENO One. 08/02 2018;52(3):203–209. doi:10.20870/oeno-one.2018.52.3.2116

30. Trapp O, Avia K, Borrelli C, Eibach R, Merdinoglu D, Töpfer R. More sustainability in Europe’s vineyards–Using resistant grapevine varieties to reduce the input of pesticides. *Plants, People*, Planet. 2025;7(6):1621–1628.

31. Schneider C, Onimus C, Prado E, et al. INRA-ResDur: the French grapevine breeding programme for durable resistance to downy and powdery mildew. Acta Horticulturae. 2019;(1248):207–214. doi:10.17660/actahortic.2019.1248.30

32. Tello J, Ibáñez J. Review: Status and prospects of association mapping in grapevine. Plant Science. 2023;327:111539. doi:10.1016/j.plantsci.2022.111539

33. Duchêne E, Butterlin G, Dumas V, Merdinoglu D. Towards the adaptation of grapevine varieties to climate change: QTLs and candidate genes for developmental stages. Theoretical and Applied Genetics. 2012;124(4):623–635.

34. Cabezas JA, Cervera MT, Ruiz-García L, Carreño J, Martínez-Zapater JM. A genetic analysis of seed and berry weight in grapevine. Genome. 2006;49(12):1572–1585. doi:10.1139/g06-122

35. Doligez A, Bouquet A, Danglot Y, et al. Genetic mapping of grapevine (*Vitis vinifera* L.) applied to the detection of QTLs for seedlessness and berry weight. Theoretical and Applied Genetics. 2002;105(5):780–795. doi:10.1007/s00122-002-0951-z

36. Doligez A, Bertrand Y, Farnos M, et al. New stable QTLs for berry weight do not colocalize with QTLs for seed traits in cultivated grapevine (*Vitis vinifera* L.). BMC Plant Biology. 2013;13(1):217. doi:10.1186/1471-2229-13-217

37. Duchêne É, Dumas V, Butterlin G, et al. Genetic variations of acidity in grape berries are controlled by the interplay between organic acids and potassium. Theoretical and Applied Genetics. 2020;133(3):993–1008.

38. Negus KL, Chen L-L, Fresnedo-Ramírez J, et al. Identification of QTLs for berry acid and tannin in a *Vitis aestivalis*-derived ‘Norton’-based population. Fruit Research. 2021;1(1):1–11. doi:10.48130/frures-2021-0008

39. Richter R, Gabriel D, Rist F, Töpfer R, Zyprian E. Identification of co-located QTLs and genomic regions affecting grapevine cluster architecture. Theoretical and Applied Genetics. 2018;132(4):1159–1177. doi:10.1007/s00122-018-3269-1

40. Underhill A, Hirsch C, Clark M. Image-based Phenotyping Identifies Quantitative Trait Loci for Cluster Compactness in Grape. Journal of the American Society for Horticultural Science. 2020;145(6):363–373. doi:10.21273/jashs04932-20

41. Su K, Zhao W, Lin H, Jiang C, Zhao Y, Guo Y. Candidate gene discovery of *Botrytis cinerea* resistance in grapevine based on QTL mapping and RNA-seq. Frontiers in Plant Science. 2023;14doi:10.3389/fpls.2023.1127206

42. Fanizza G, Lamaj F, Costantini L, Chaabane R, Grando MS. QTL analysis for fruit yield components in table grapes (*Vitis vinifera*). Theoretical and Applied Genetics. 2005;111(4):658–664. doi:10.1007/s00122-005-2016-6

43. Viana AP, Riaz S, Walker MA. Genetic dissection of agronomic traits within a segregating population of breeding table grapes. Genetics and Molecular Research. 2013;12(2):951–964. doi:10.4238/2013.april.2.11

44. This P, Lacombe T, Cadle-Davidson M, Owens CL. Wine grape (*Vitis vinifera* L.) color associates with allelic variation in the domestication gene VvmybA1. Theoretical and Applied Genetics. 2007;114(4):723–730. doi:10.1007/s00122-006-0472-2

45. Flutre T, Le Cunff L, Fodor A, et al. A genome-wide association and prediction study in grapevine deciphers the genetic architecture of multiple traits and identifies genes under many new QTLs. G3 Genes|Genomes|Genetics. 2022;12(7)doi:10.1093/g3journal/jkac103

46. García-Abadillo J, Barba P, Carvalho T, et al. Dissecting the complex genetic basis of pre- and post-harvest traits in *Vitis vinifera* L. using genome-wide association studies. Horticulture Research. 2024;11(2):uhad283. doi:10.1093/hr/uhad283

47. Liu Z, Wang N, Su Y, et al. Grapevine pangenome facilitates trait genetics and genomic breeding. Nature Genetics. 2024;56(12):2804–2814. doi:10.1038/s41588-024-01967-5

48. Wang X, Liu Z, Zhang F, et al. Integrative genomics reveals the polygenic basis of seedlessness in grapevine. Current Biology. 2024;34(16):3763–3777.e5. doi:10.1016/j.cub.2024.07.022

49. Gan Y, Liu Z, Zhang F, et al. Deep learning empowers genomic selection of pest-resistant grapevine. Horticulture Research. 2025;12(8):uhaf128. doi:10.1093/hr/uhaf128

50. Buckler ES, Holland JB, Bradbury PJ, et al. The genetic architecture of maize flowering time. Science. 2009;325(5941):714–718. doi:10.1126/science.1174276

51. Maurer A, Draba V, Jiang Y, et al. Modelling the genetic architecture of flowering time control in barley through nested association mapping. BMC Genomics. 2015;16:290. doi:10.1186/s12864-015-1459-7

52. Benaouda S, Dadshani S, Koua P, Léon J, Ballvora A. Identification of QTLs for wheat heading time across multiple-environments. Theor Appl Genet. 2022;135(8):2833–2848. doi:10.1007/s00122-022-04152-6

53. Hori K, Nonoue Y, Ono N, et al. Genetic architecture of variation in heading date among Asian rice accessions. BMC Plant Biol. 2015;15(1):115. doi:10.1186/s12870-015-0501-x

54. Piepho H-P, Möhring J. Computing Heritability and Selection Response From Unbalanced Plant Breeding Trials. Genetics. 2007;177(3):1881–1888. doi:10.1534/genetics.107.074229

55. Schmidt P, Hartung J, Rath J, Piepho H-P. Estimating Broad-Sense Heritability with Unbalanced Data from Agricultural Cultivar Trials. Crop Science. 2019;59(2):525–536. doi:10.2135/cropsci2018.06.0376

56. Ortiz-Barrientos D, Jordan D, Cooper M. The hidden geometry of breeding constraints. Nature Plants. 2026;12:1316–1324. doi:10.1038/s41477-026-02332-6

57. Frenzke L, Röckel F, Wenke T, et al. Genotyping-by-sequencing-based high-resolution mapping reveals a single candidate gene for the grapevine veraison locus Ver1. Plant Physiology. 2024;196(1):244–260. doi:10.1093/plphys/kiae272

58. 58. Chedid E. Aptitudes agro-œnologiques des hybrides interspécifiques complexes de vigne : incidence des régions génomiques issues des espèces sauvages du genre Vitis. 2023. https://publication-theses.unistra.fr/public/theses_doctorat/2023/CHEDID_Elsa_2023_ED414.pdf

59. Chedid E, Avia K, Dumas V, et al. LiDAR Is Effective in Characterizing Vine Growth and Detecting Associated Genetic Loci. Plant Phenomics. 2023;5:0116. doi:10.34133/plantphenomics.0116

60. Chedid E, Dumas V, Avia K, Merdinoglu D, Duchêne É. Allele-based modeling to predict phenological stages of grapevine hybrids under future climatic conditions. Theoretical and Applied Genetics. 2025;138(6):110. doi:10.1007/s00122-025-04891-2

61. de Oliveira GL, Francisco FR, de Moura YA, et al. Genome-wide association study in a diverse grapevine collection provides insights into the genetic basis of berry size and cluster architecture traits. PLOS ONE. 2026;21(3):e0343491. doi:10.1371/journal.pone.0343491

62. Thorat KD, Upadhyay A, Samarth RR, et al. Genome-wide association analysis to identify genomic regions and predict candidate genes for bunch traits in grapes (Vitis vinifera L.). Scientia Horticulturae. 2024;328:112882. doi:10.1016/j.scienta.2024.112882

63. de Sousa Moreira L, Clark MD, Tabb A, et al. Identification of novel quantitative trait loci associated with table grape fruit quality characteristics in a segregating population and transferability of existing quality markers. Journal of the American Society for Horticultural Science. 2024;149(1):50–60. doi:10.21273/jashs05334-23

64. Ban Y, Mitani N, Sato A, Kono A, Hayashi T. Genetic dissection of quantitative trait loci for berry traits in interspecific hybrid grape (*Vitis labruscana* × *Vitis vinifera*). Euphytica. 2016/10/01 2016;211(3):295–310. doi:10.1007/s10681-016-1737-8

65. Reshef N, Karn A, Manns DC, et al. Stable QTL for malate levels in ripe fruit and their transferability across Vitis species. Horticulture Research. 2022;9:uhac009. doi:10.1093/hr/uhac009

66. Kadium VR, Pilli R, Svyantek A, et al. Dissecting the genetic basis of relevant fruit quality traits in interspecific grapevines (Vitis spp.). Horticulture Research. 2026;13(4)doi:10.1093/hr/uhaf353

67. Alahakoon D, Fennell A, Helget Z, et al. Berry anthocyanin, acid, and volatile trait analyses in a grapevine-interspecific F2 population using an integrated GBS and rhAmpSeq genetic map. Plants. 2022;11(5):696. doi:10.3390/plants11050696

68. Mamani M, López ME, Correa J, Ravest G, Hinrichsen P. Identification of stable quantitative trait loci and candidate genes for sweetness and acidity in tablegrape using a highly saturated single-nucleotide polymorphism-based linkage map. Australian Journal of Grape and Wine Research. 2021;27(3):308–324. doi:10.1111/ajgw.12497

69. Tsepilov YA, Shin S-Y, Soranzo N, et al. Nonadditive Effects of Genes in Human Metabolomics. Genetics. 2015;200(3):707–718. doi:10.1534/genetics.115.175760

70. Shaffer W, Papin V, Yadav S, et al. Local genomic estimates provide a powerful framework for haplotype discovery. openRxiv; 2025:2025.08.28.672830.

71. Dal Santo S, Zenoni S, Sandri M, et al. Grapevine field experiments reveal the contribution of genotype, the influence of environment and the effect of their interaction (G×E) on the berry transcriptome. The Plant Journal. 2018;93(6):1143–1159. doi:10.1111/tpj.13834

72. Elshire RJ, Glaubitz JC, Sun Q, et al. A Robust, Simple Genotyping-by-Sequencing (GBS) Approach for High Diversity Species. PLoS ONE. 2011/05/04 2011;6(5):e19379. doi:10.1371/journal.pone.0019379

73. Catchen J, Hohenlohe PA, Bassham S, Amores A, Cresko WA. Stacks: an analysis tool set for population genomics. Molecular Ecology. 2013/06 2013;22(11):3124–3140. doi:10.1111/mec.12354

74. Catchen JM, Amores A, Hohenlohe P, Cresko W, Postlethwait JH. Stacks: Building and Genotyping Loci De Novo From Short-Read Sequences. G3-Genes Genomes Genetics. Aug 1 2011;1(3):171–182. doi:10.1534/g3.111.000240

75. Li H, Durbin R. Fast and accurate short read alignment with Burrows-Wheeler transform. Bioinformatics. 2009;25(14):1754–1760. doi:10.1093/bioinformatics/btp324

76. McKenna A, Hanna M, Banks E, et al. The Genome Analysis Toolkit: A MapReduce framework for analyzing next-generation DNA sequencing data. Genome Research. 2010;20(9):1297–1303. doi:10.1101/gr.107524.110

77. Danecek P, Auton A, Abecasis G, et al. The variant call format and VCFtools. Bioinformatics. 2011;27(15):2156–2158. doi:10.1093/bioinformatics/btr330

78. Danecek P, Bonfield JK, Liddle J, et al. Twelve years of SAMtools and BCFtools. GigaScience. 2021/01/29 2021;10(2):giab008. doi:10.1093/gigascience/giab008

79. Jombart T, Devillard S, Balloux F. Discriminant analysis of principal components: a new method for the analysis of genetically structured populations. BMC Genetics. 2010;11(1):94. doi:10.1186/1471-2156-11-94

80. Miller JM, Cullingham CI, Peery RM. The influence of a priori grouping on inference of genetic clusters: simulation study and literature review of the DAPC method. Heredity. 2020;125(5):269–280. doi:10.1038/s41437-020-0348-2

81. Grubbs FE. Procedures for Detecting Outlying Observations in Samples. Technometrics. 1969;11(1):1–21. doi:10.1080/00401706.1969.10490657

82. Peterson R, A. Finding Optimal Normalizing Transformations via bestNormalize. The R Journal. 2021;13(1):310. doi:10.32614/rj-2021-041

83. Lozano-Isla F, Diaz-Saucedo Y. Data from: inti: Tools and Statistical Procedures in Plant Science. 2020. doi:10.32614/cran.package.inti

84. Rastas P. Lep-MAP3: robust linkage mapping even for low-coverage whole genome sequencing data. Bioinformatics. 2017;33(23):3726–3732. doi:10.1093/bioinformatics/btx494

85. Stam P. Construction of integrated genetic linkage maps by means of a new computer package: JoinMap. Plant J. 1993;3:739–744.

86. Broman KW, Wu H, Sen Ś, Churchill GA. R/qtl: QTL mapping in experimental crosses. Bioinformatics. 2003/05/01 2003;19(7):889–890. doi:10.1093/bioinformatics/btg112

87. Ouellette LA, Reid RW, Blanchard SG, Brouwer CR. LinkageMapView-rendering high-resolution linkage and QTL maps. Bioinformatics. Jan 15 2018;34(2):306–307. doi:10.1093/bioinformatics/btx576

88. Thérèse Navarro A, Tumino G, Voorrips RE, et al. Multiallelic models for QTL mapping in diverse polyploid populations. BMC Bioinformatics. 2022/02/14 2022;23(1):67. doi:10.1186/s12859-022-04607-z

89. Lipka AE, Tian F, Wang Q, et al. GAPIT: genome association and prediction integrated tool. Bioinformatics. 2012;28(18):2397–2399. doi:10.1093/bioinformatics/bts444

90. Laporte F, Charcosset A, Mary-Huard T. Efficient ReML inference in variance component mixed models using a Min-Max algorithm. PLOS Computational Biology. 2022;18(1):e1009659. doi:10.1371/journal.pcbi.1009659

