## Supplementary Figures and Tables for "Uncovering the genetic basis of agronomic traits in over 1,000 grapevine genotypes derived from a disease resistance breeding program"

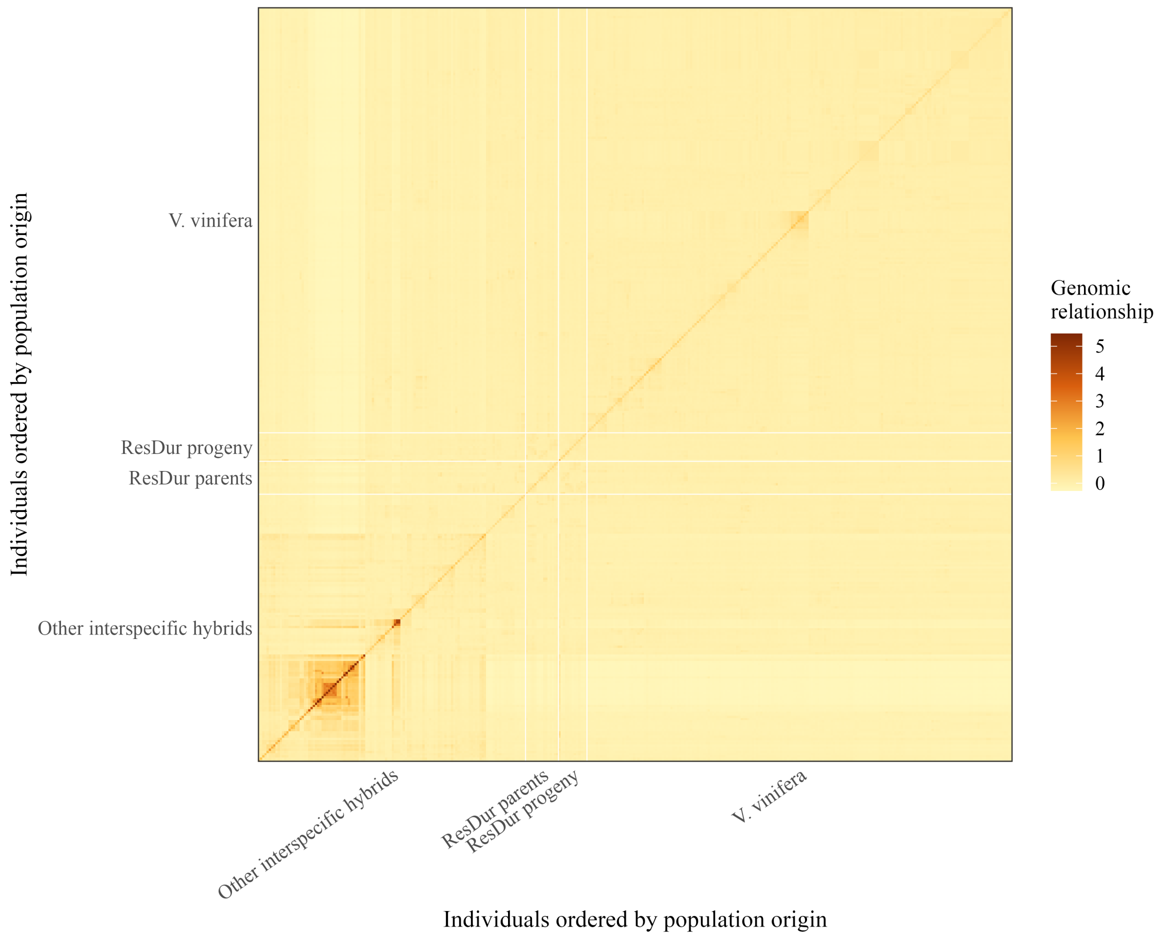


**Supplementary Figure 1.** Genomic relationship matrix for the ResDur panel and ampelographic collection. Both axes contain the same individuals ordered by origin group (other interspecific hybrids, ResDur parents, ResDur progeny and *V. vinifera* accessions); darker colors indicate higher relatedness.

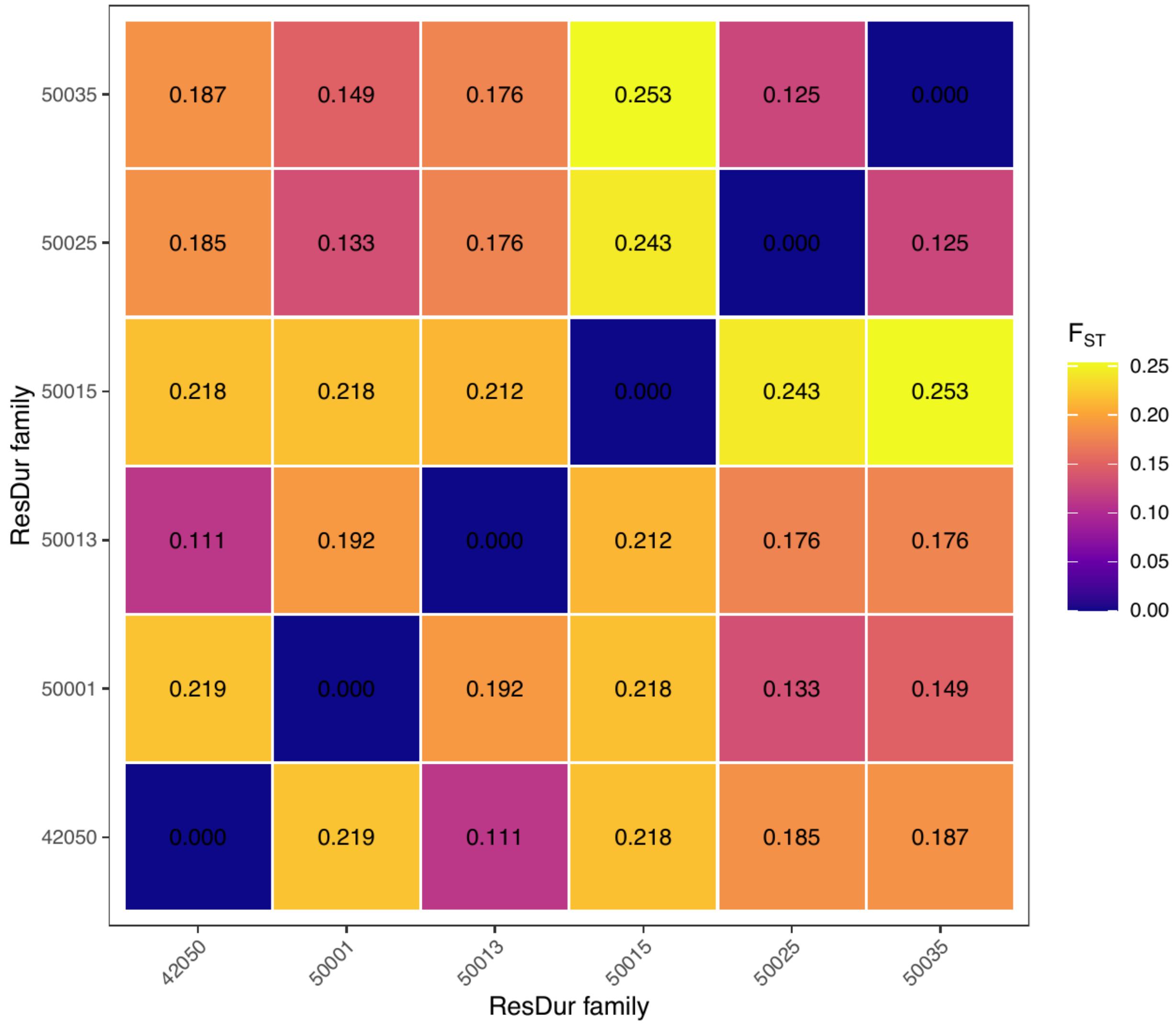


**Supplementary Figure 2.** Pairwise Hudson FST among the six largest ResDur families. Both axes identify families; each tile reports the genome-wide Hudson FST estimate from 189,426 SNPs. Abbreviations: FST, fixation index; SNP, single-nucleotide polymorphism.

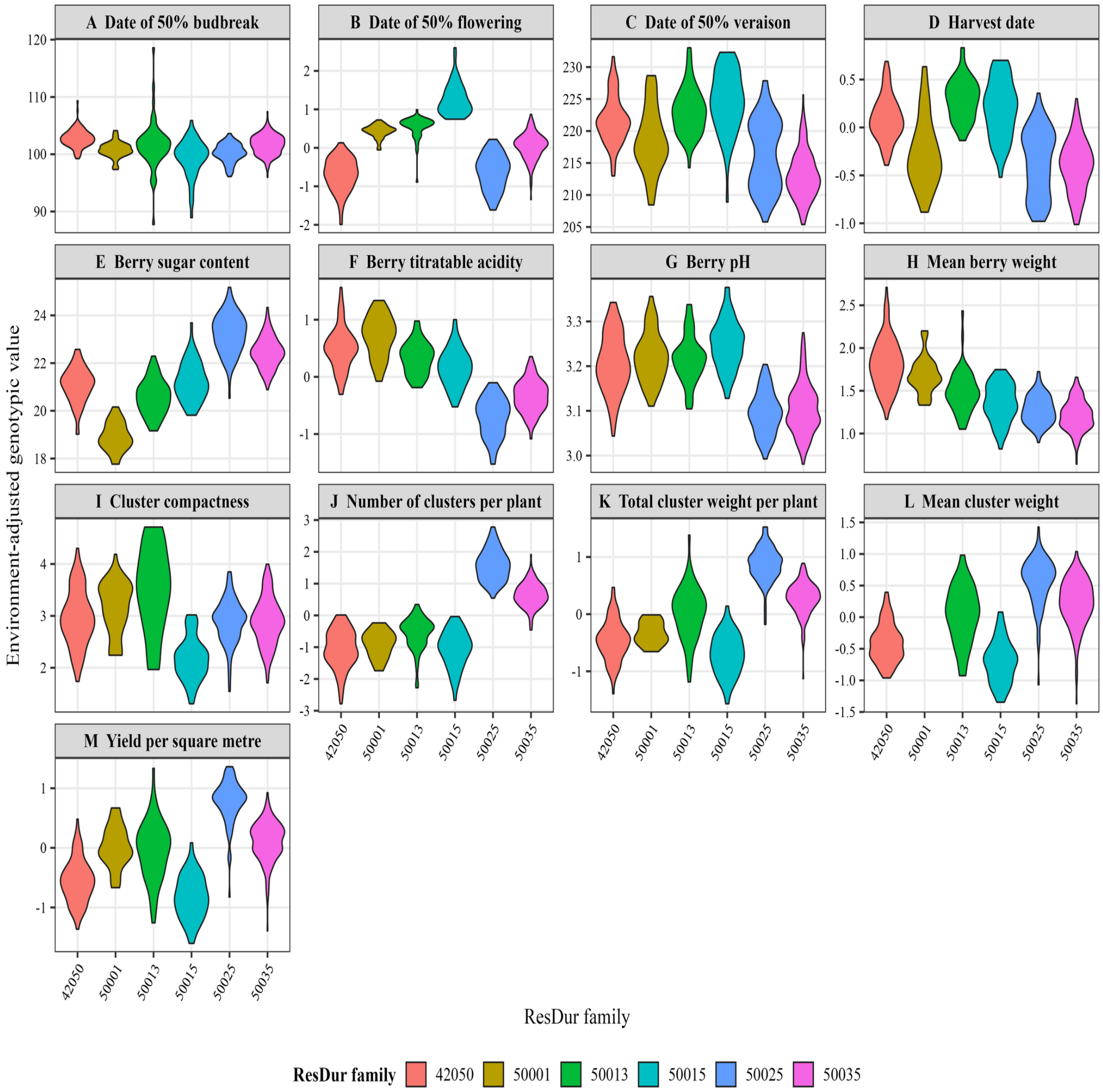


**Supplementary Figure 3.** Distribution of agronomic traits across the six largest ResDur families. Violin plots show the variability of environment-adjusted genotypic values within families: (A) budbreak, (B) flowering, (C) veraison, (D) harvest, (E) berry sugar content, (F) titratable acidity, (G) berry pH, (H) mean berry weight, (I) cluster compactness, (J) number of clusters per plant, (K) total cluster weight per plant, (L) mean cluster weight, and (M) yield per m². Colors identify the six ResDur families and are defined in the key below the panels.

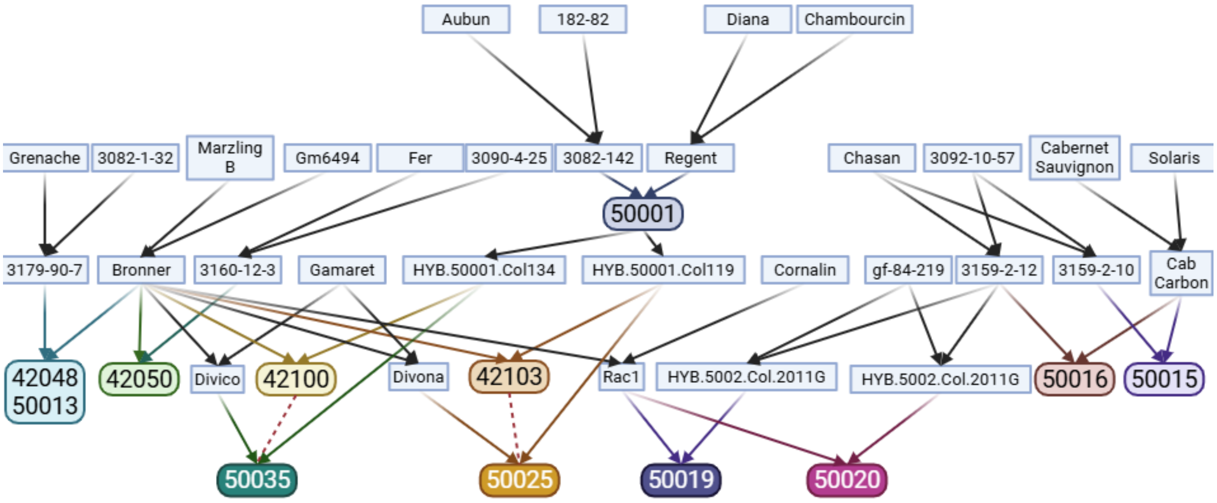


**Supplementary Figure 4.** Pedigree of the twelve largest populations of the INRAE-ResDur breeding program. Abbreviations: INRAE, French National Research Institute for Agriculture, Food and Environment.

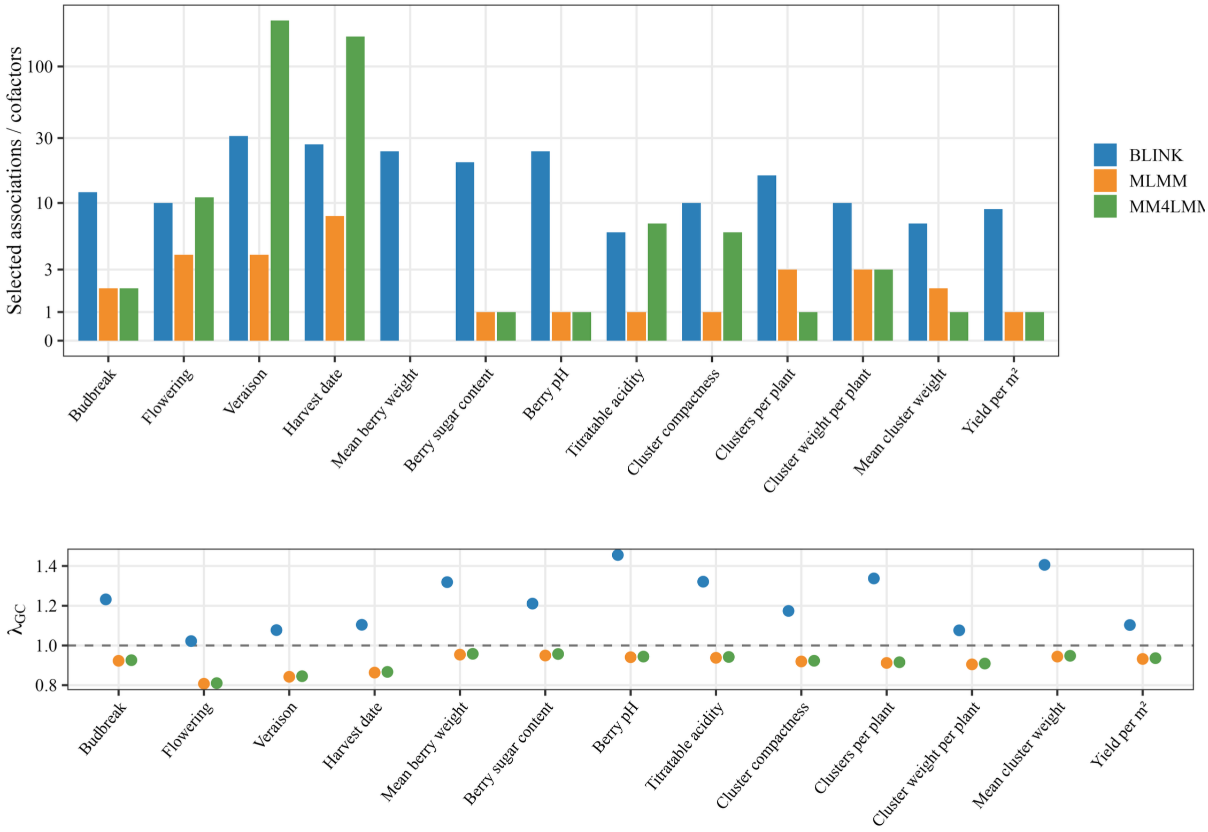


**Supplementary Figure 5. A**dditive GWAS across the 13 traits. The upper panel reports BLINK Bonferroni associations, final MLMM cofactors and MM4LMM Bonferroni associations; the lower panel reports genomic-control inflation factors. BLINK fitted LD1-LD5, whereas MLMM and MM4LMM fitted genomic kinship plus LD1-LD5. Counts are model outputs and do not represent independent validation. Abbreviations: GWAS, genome-wide association study; BLINK, Bayesian-information and Linkage-disequilibrium Iteratively Nested Keyway; MLMM, multi-locus mixed model; MM4LMM, Min-Max algorithm for large-scale mixed models; DAPC, discriminant analysis of principal components; LD1-LD5, the first five DAPC linear discriminant axes.

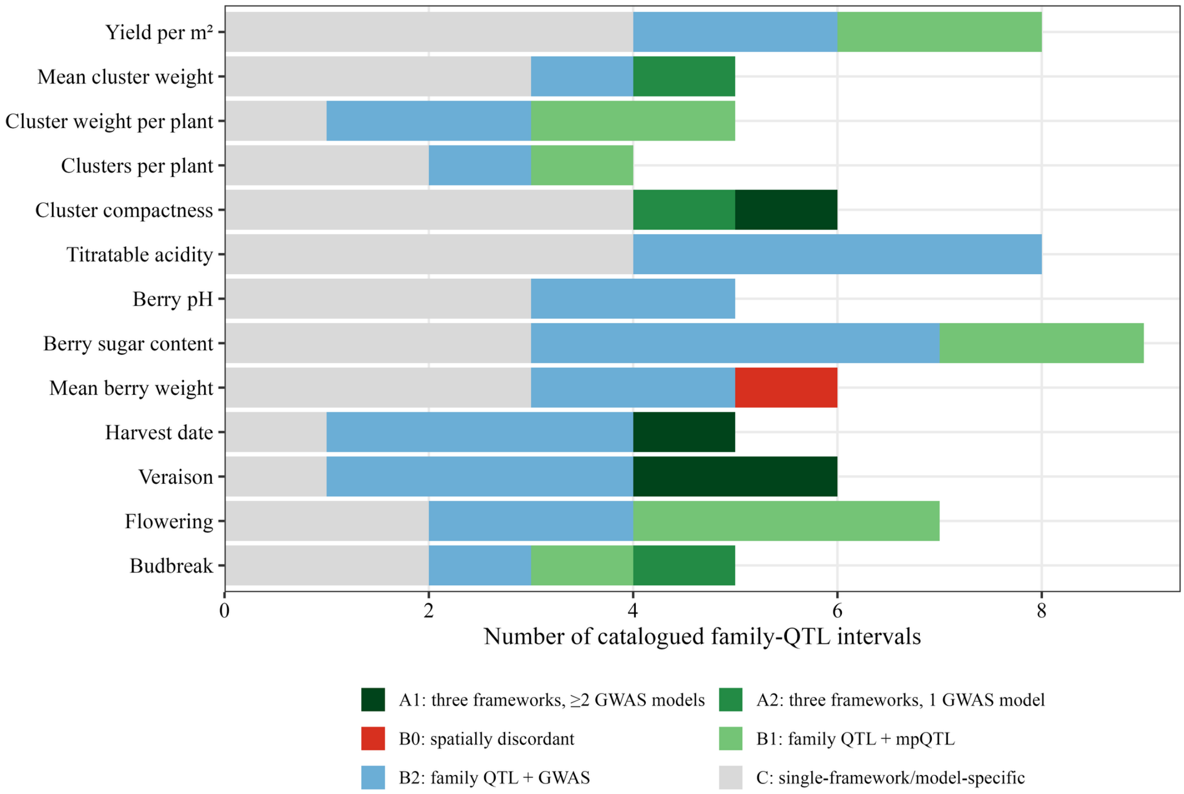


**Supplementary Figure 6.** Evidence tiers assigned to the 79 catalogued single-family QTL intervals after physical-coordinate integration with population-adjusted mpQTL and additive GWAS. Tier A requires local support from all three mapping frameworks; Tier B represents two-framework or spatially discordant evidence; Tier C remains single-framework or model-specific. Abbreviations: QTL, quantitative trait locus; mpQTL, multiple-population QTL mapping; GWAS, genome-wide association study. The catalogue includes the three suggestive family-50015 signals identified in Supplementary Table 8.

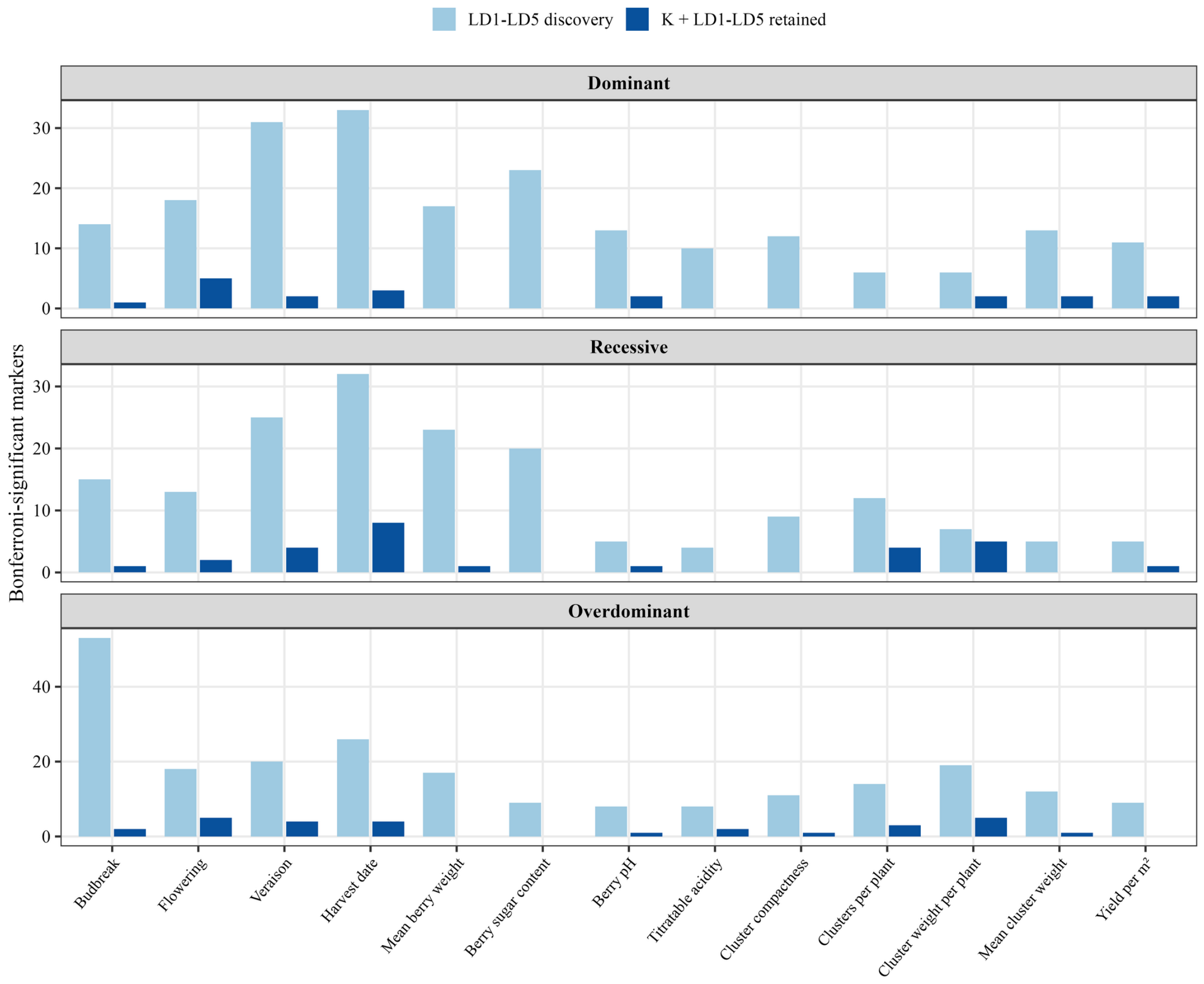


**Supplementary Figure 7.** Exploratory non-additive association sensitivity analysis. Pale bars show Bonferroni candidates from BLINK fitted with LD1-LD5; dark bars show candidates whose marker-by-marker MM4LMM sensitivity retest with genomic kinship plus LD1-LD5 also passed the original genome-wide threshold. The second-stage test is selected-candidate sensitivity analysis, not independent replication. Abbreviations: BLINK, Bayesian-information and Linkage-disequilibrium Iteratively Nested Keyway; MM4LMM, Min-Max algorithm for large-scale mixed models; DAPC, discriminant analysis of principal components; LD1-LD5, the first five DAPC linear discriminant axes.

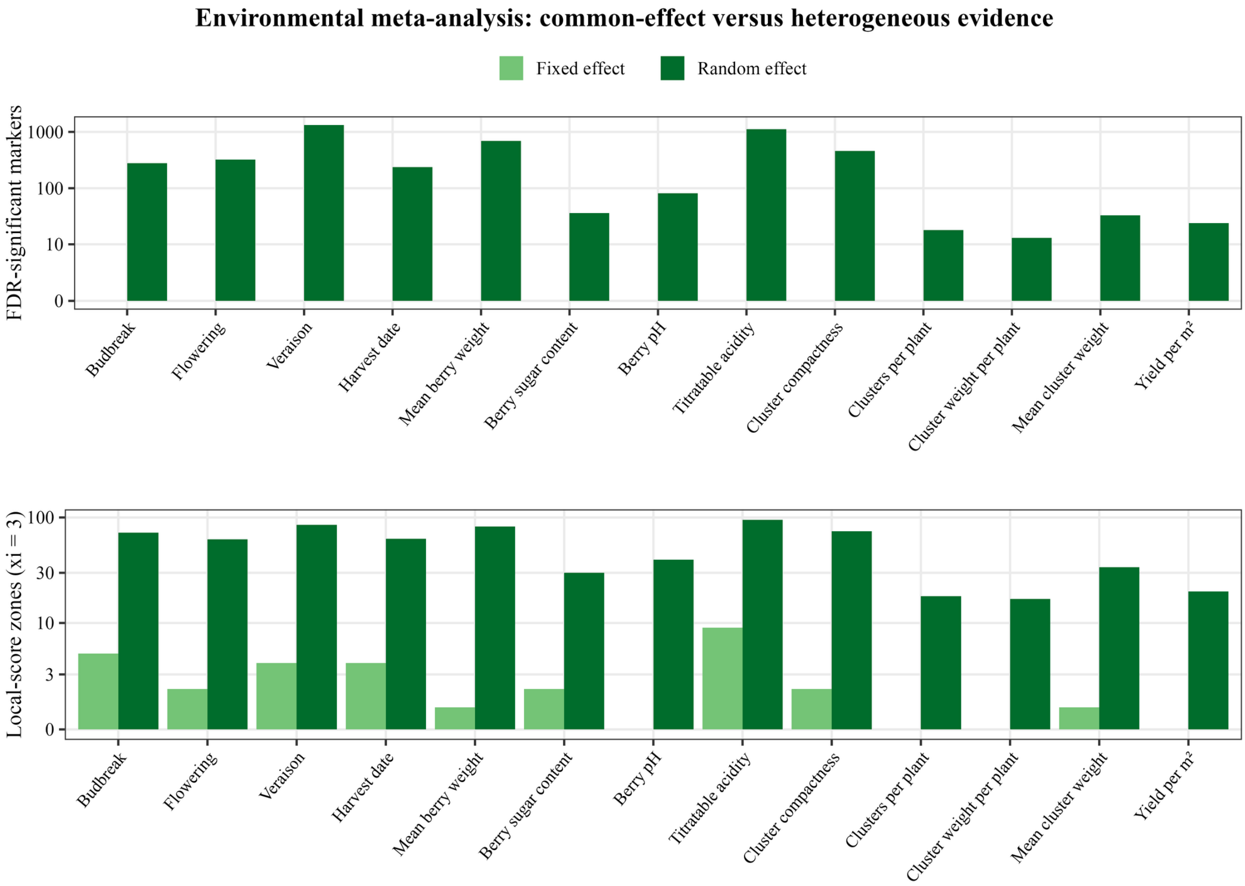


**Supplementary Figure 8.** Environment-specific GWAS meta-analysis. Marker-level FDR counts and local-score zones are shown separately for common fixed effects and heterogeneous random effects. The large excess of random-effect evidence is interpreted cautiously because genotype and family composition changed among site-by-year environments. Abbreviations: GWAS, genome-wide association study; FDR, false discovery rate.

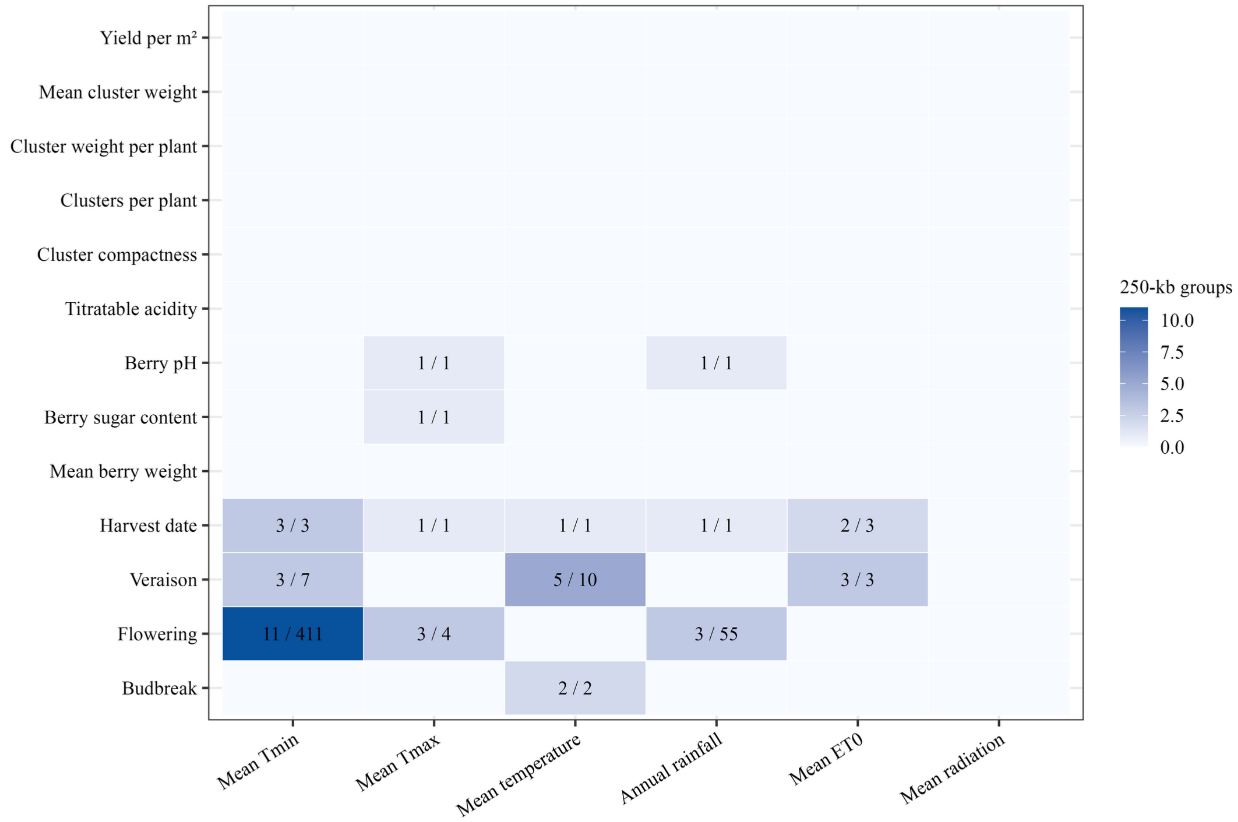


**Supplementary Figure 9.** Climate meta-regression summary. Each non-empty cell gives the number of descriptive 250-kb proximity groups followed by the number of marker-covariate tests passing 5% FDR. Climate predictors were tested in separate univariate regressions and are correlated; counts therefore do not represent independent causal climatic effects. Abbreviations: FDR, false discovery rate.

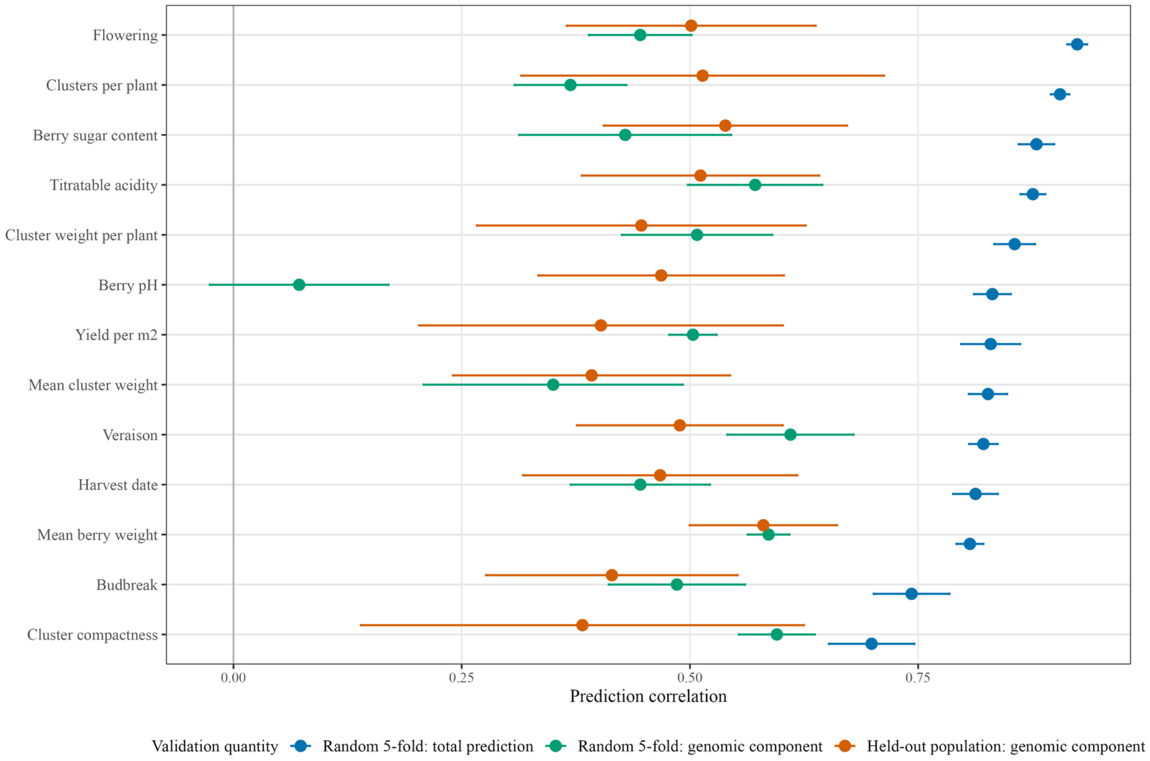


**Supplementary Figure 10.** Prediction validation across 13 traits for the population-adjusted localGEBV marker model. Points are mean Pearson correlations and horizontal bars show ±1 SD across stratified random folds or held-out populations. Blue values are total predictions from stratified random five-fold validation and include both the estimated population fixed effect and the genomic marker score. Green values are the genomic component alone in the same random folds. Orange values are the genomic component in leave-one-population-out validation; the fixed effect of a population absent from training cannot be estimated and was therefore not included. Random-fold total prediction can benefit from known population means, whereas leave-one-population-out validation more directly evaluates transfer of marker effects across genetic backgrounds. These are internal cross-validation results, not independent validation. Abbreviations: localGEBV, local genomic estimated breeding value; SD, standard deviation.

A

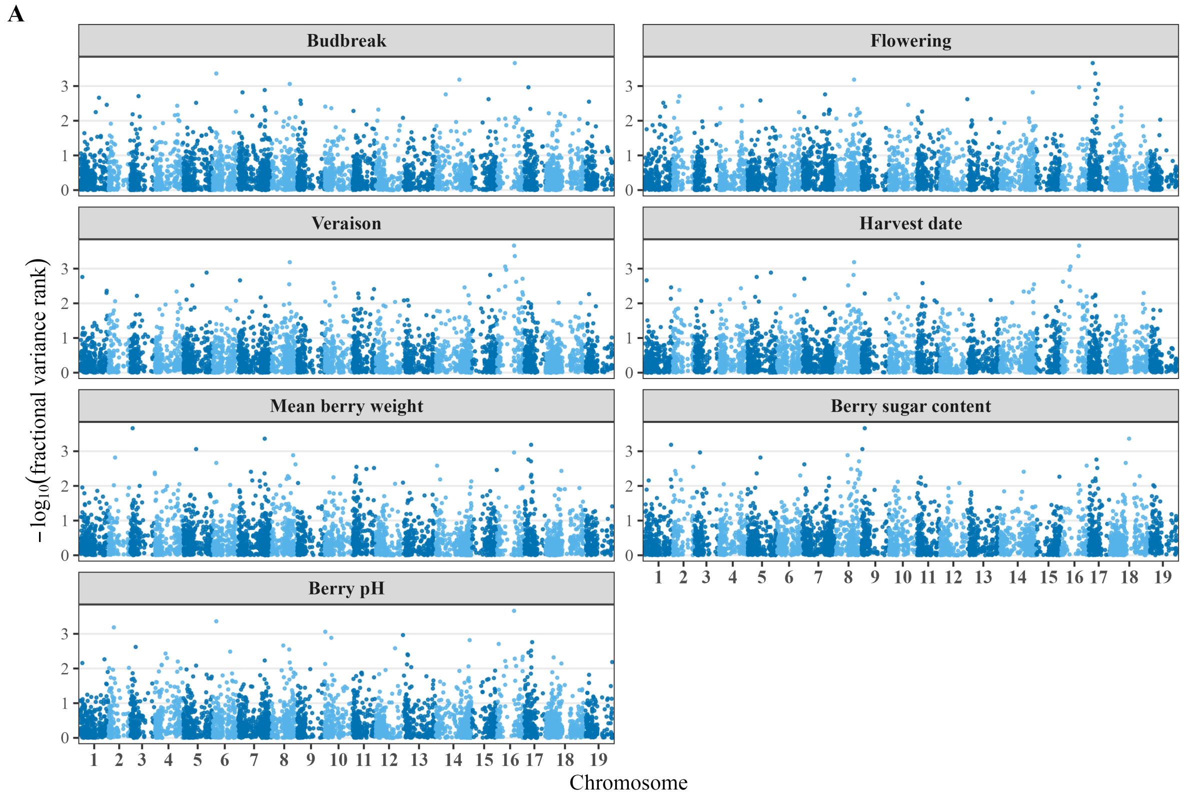


**Supplementary Figure 11, panel A.** Population-adjusted localGEBV block landscapes for budbreak, flowering, veraison, harvest date, mean berry weight, berry sugar content and berry pH. Panel B continues the figure on the following page. Abbreviations: localGEBV, local genomic estimated breeding value.

**B**

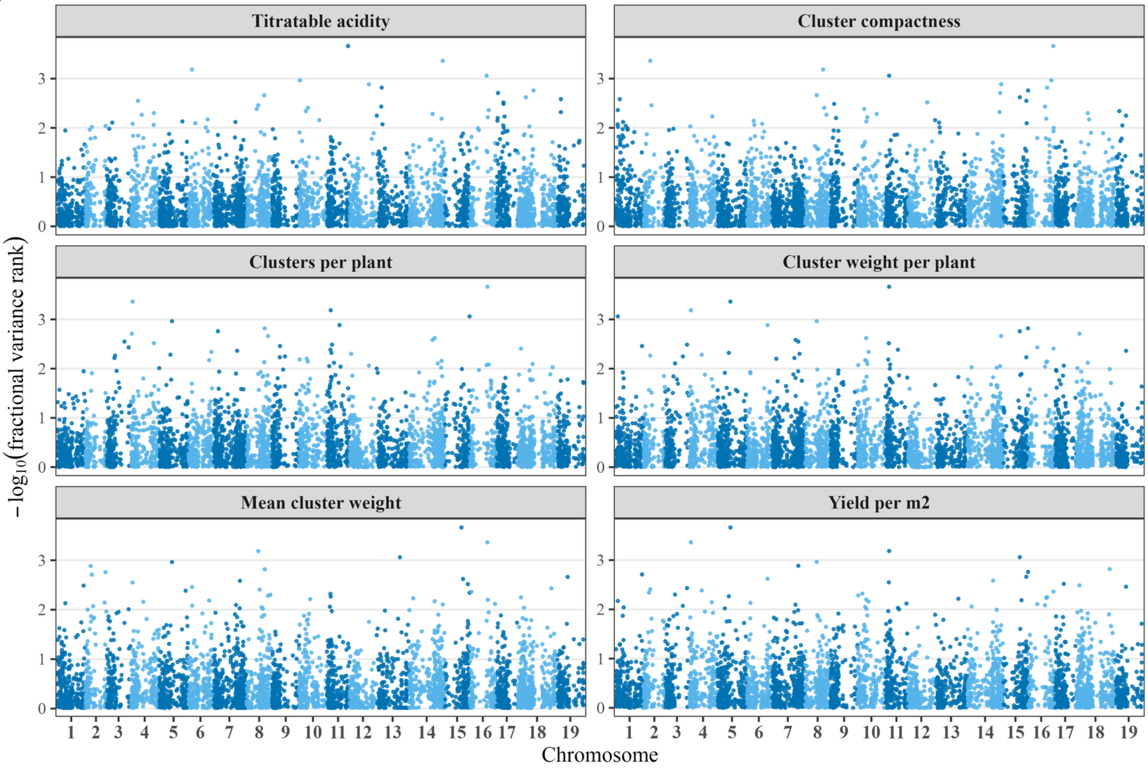


**Supplementary Figure 11 (continued), panel B.** Population-adjusted localGEBV block landscapes for all 13 traits. Each point represents one chromosome-specific marker block defined from within-population-centered pairwise linkage disequilibrium using $r^2 \geq 0.30$ and one tolerated low-LD flank. Marker effects were estimated in 772 progeny with ridge-regression BLUP and 12 fixed population indicators. Centered marker-dosage-by-effect contributions were summed within each block, and the among-genotype variance of the resulting block score was ranked within trait. Alternating blue shades distinguish chromosomes. The ordinate is −log10[r/(B+1)], where r is the block variance rank and B is the number of blocks. It is therefore a rank-based breeding-prioritization display, not an association test, P value or significance scale. Panel A shows budbreak through berry pH; panel B shows titratable acidity through yield per m². Abbreviations: localGEBV, local genomic estimated breeding value; LD, linkage disequilibrium; BLUP, best linear unbiased predictor.

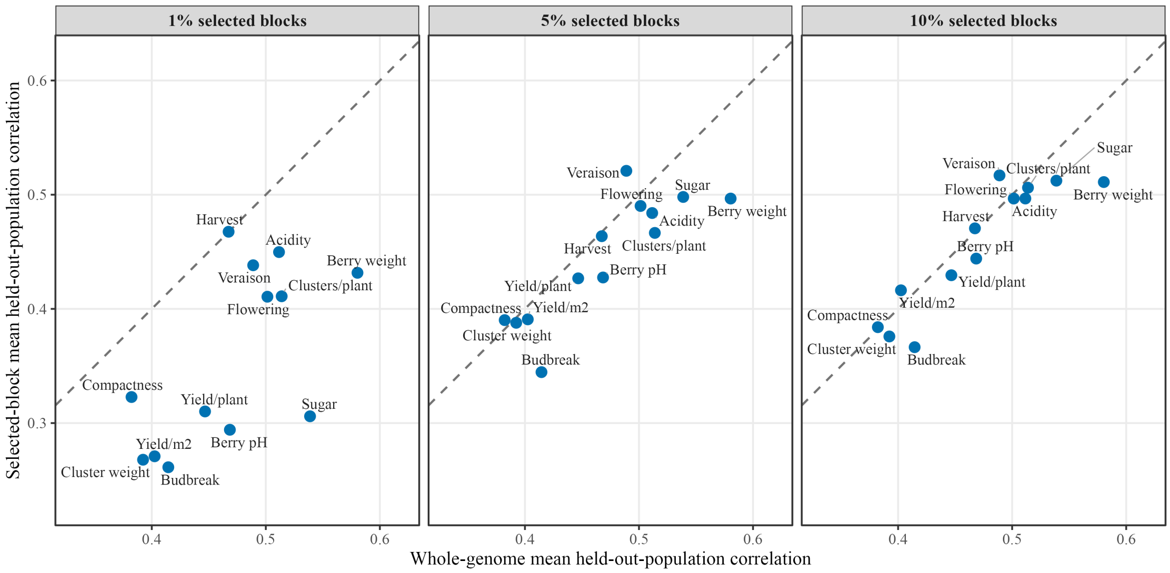


**Supplementary Figure 12.** Nested leave-one-population-out comparison of training-selected localGEBV block subsets with whole-genome prediction. For every trait and held-out population, marker centering, rrBLUP marker effects, block scores and block-variance ranking were recomputed using only the training populations while retaining the pre-defined primary LD-block boundaries. The top 1%, 5% or 10% of training-ranked blocks were then used to predict the held-out population. Each labelled point gives the mean correlation across eligible held-out populations for the whole-genome genomic score (x-axis) and the selected-block score (y-axis). The dashed diagonal denotes equal predictive correlation; points above it favor the selected subset. This nested procedure prevents phenotype information from the held-out population entering block selection, but remains internal validation rather than independent confirmation. Abbreviations: localGEBV, local genomic estimated breeding value; rrBLUP, ridge-regression best linear unbiased prediction; LD, linkage disequilibrium.

**Supplementary Table 1.** Pairwise Hudson FST estimates. Abbreviations: FST, fixation index.

| Family | 42050 | 50001 | 50013 | 50015 | 50025 | 50035 |
|---|---|---|---|---|---|---|
| 42050 | — | 0.219 | 0.111 | 0.218 | 0.185 | 0.187 |
| 50001 | 0.219 | — | 0.192 | 0.218 | 0.133 | 0.149 |
| 50013 | 0.111 | 0.192 | — | 0.212 | 0.176 | 0.176 |
| 50015 | 0.218 | 0.218 | 0.212 | — | 0.243 | 0.253 |
| 50025 | 0.185 | 0.133 | 0.176 | 0.243 | — | 0.125 |
| 50035 | 0.187 | 0.149 | 0.176 | 0.253 | 0.125 | — |

**Supplementary Table 2.** Phenotypic differentiation among the six largest families. Abbreviations: η², eta-squared effect size; ANOVA, analysis of variance.

| Trait | n | families | η² family | anova p | kruskal p |
|---|---|---|---|---|---|
| BUD_DATE | 555 | 6 | 0.178 | 9.959E-22 | 4.258E-25 |
| FLO_50 | 553 | 6 | 0.732 | 1.334E-153 | 3.674E-88 |
| VER_50 | 553 | 6 | 0.502 | 2.27E-80 | 1.713E-58 |
| HARVEST_DATE | 526 | 6 | 0.497 | 3.38E-75 | 1.525E-57 |
| TSS | 518 | 6 | 0.708 | 3.076E-134 | 4.579E-75 |
| BER_TA_g | 551 | 6 | 0.698 | 4.235E-139 | 1.416E-84 |
| BERRY_pH | 551 | 6 | 0.537 | 9.426E-89 | 2.179E-64 |
| SBER_W_g | 529 | 6 | 0.546 | 2.75E-87 | 4.76E-59 |
| MORPHO_OIV_204 | 549 | 6 | 0.265 | 2.183E-34 | 1.044E-24 |
| NB_CLUST_PLANT | 507 | 6 | 0.809 | 1.983E-177 | 8.224E-89 |
| YIELD_PLANT | 500 | 6 | 0.706 | 5.631E-129 | 1.614E-75 |
| SCLUST_W | 490 | 5 | 0.557 | 2.532E-84 | 7.806E-58 |
| YIELD_OIV_504 | 518 | 6 | 0.647 | 2.871E-113 | 3.811E-69 |

**Supplementary Table 3.** Descriptive statistics for the ResDur population, including trait means, variances, number of genotypes, years, sites, environments, and the models used to calculate BLUPs. Abbreviations: BLUP, best linear unbiased predictor.

| Trait | Mean | Standard deviation | Number of genotype | Fixe effect | Random effects | Genetic variance | Residual variance | Phenotypic variance | Number of years | Number of locations | Number of environ ments | Normalisati on |
|---|---|---|---|---|---|---|---|---|---|---|---|---|
| Berry titratable acidity | 5.22 | 1.43 | 1,049 | Location | Genotype + Year + Genotype*Year + Genotype*Location + Year*Location + Pop | 0.211 | 0.199 | 0.543 | 16 | 5 | 41 | Yes |
| Berry pH | 3.17 | 0.18 | 1,051 | Location | Genotype + Genotype*Year + Genotype*Location + Year*Location + Pop | 0.005 | 0.008 | 0.017 | 17 | 5 | 44 | No |
| Berry sugar content | 21.06 | 2.06 | 1,026 | Location | Genotype + Genotype*Location + Year*Location + Pop | 0.814 | 1.299 | 2.418 | 15 | 5 | 36 | No |
| Date of 50% budbreak | 102.12 | 7.71 | 1,072 | Location | Genotype + Genotype*Location + Year*Location + Genotype*Year*Location + Pop | 9.236 | 4.534 | 22.409 | 14 | 4 | 25 | No |
| Date of 50% veraison | 218.93 | 12.62 | 1,068 | Location | Genotype/Pop + Genotype + Year + Genotype*Year + Genotype*Location + Year*Location + Genotype*Year*Location + Pop | 2.737 | 5.234 | 15.599 | 17 | 5 | 41 | No |
| Date of 50% flowering | 156.83 | 9 | 1,072 | Location | Genotype + Year + Genotype*Location + Year*Location + Genotype*Year*Location + Pop | 0.051 | 0.037 | 0.146 | 15 | 4 | 28 | No |
| Harvest date | 267.15 | 15.61 | 1,049 | Location | Genotype/Pop + Genotype + Year + Genotype*Year + Genotype*Location + Year*Location + Genotype*Year*Location + Pop | 0.04 | 0.08 | 0.199 | 17 | 5 | 37 | No |
| Mean berry weight | 1.45 | 0.42 | 1,038 | Location | Genotype + Genotype*Location + Year*Location + Genotype*Year*Location + Pop | 0.063 | 0.029 | 0.19 | 16 | 5 | 38 | No |
| Cluster compactness | 2.95 | 1.25 | 1,041 | Location | Genotype + Genotype*Location + Year*Location + Genotype*Year*Location + Pop | 0.434 | 0.584 | 1.306 | 16 | 5 | 31 | No |
| Number of clusters per plant | 18.27 | 11.52 | 998 | Location | Genotype + Genotype*Location + Year*Location + Genotype*Year*Location + Pop | 0.18 | 0.176 | 0.455 | 14 | 4 | 27 | Yes |
| Total cluster weight per plant | 1.75 | 1.4 | 990 | Location | Genotype + Genotype*Location + Year*Location + Genotype*Year*Location + Pop | 0.201 | 0.209 | 0.591 | 14 | 4 | 27 | Yes |

| Trait | Mean | Standard deviation | Number of genotype | Fixe effect | Random effects | Genetic variance | Residual variance | Phenotypic variance | Number of years | Number of locations | Number of environ ments | Normalisati on |
|---|---|---|---|---|---|---|---|---|---|---|---|---|
| Mean cluster weight | 114.77 | 76.93 | 993 | Location | Genotype + Genotype*Location + Year*Location + Genotype*Year*Location + Pop | 0.189 | 0.186 | 0.537 | 12 | 5 | 34 | Yes |
| Yield per square meter | 0.83 | 0.58 | 1,019 | Location | Genotype + Genotype*Location + Year*Location + Genotype*Year*Location + Pop | 0.217 | 0.279 | 0.633 | 15 | 5 | 40 | Yes |

**Supplementary Table 4.** Descriptive statistics for the six largest families, including trait means, variances, heritability, and number of genotypes, years, sites, and environments. Abbreviations: $H^2$, broad-sense heritability.

| Trait | Population ID | Mean | Standard deviation | Number of genotype | Fixe effect | Random effects | Genetic variance | Residual variance | Phenotypic variance | $H^2$ | Numbe r of year | Numbe r of location | Number of environmen t | Normalisation |
|---|---|---|---|---|---|---|---|---|---|---|---|---|---|---|
| Berry titratable acidity | 42050 | 6.04 | 1.7 | 109 | Location | Genotype + Genotype*Year + Genotype*Location + Year*Location | 0.725 | 0.443 | 1.339 | 0.84 | 6 | 3 | 14 | No |
| | 50001 | 6.52 | 2.01 | 63 | Location | Genotype + Year*Location | 0.407 | 0.3 | 0.706 | 0.84 | 3 | 3 | 6 | Yes |
| | 50013 | 5.71 | 1.5 | 64 | Location | Genotype + Genotype*Location + Year*Location | 0.311 | 0.451 | 0.818 | 0.67 | 5 | 3 | 12 | No |
| | 50015 | 5.48 | 1.46 | 56 | Location | Genotype + Year + Genotype*Location + Year*Location | 0.383 | 0.55 | 1.014 | 0.75 | 5 | 3 | 10 | No |
| | 50025 | 4.23 | 0.82 | 93 | None | Genotype + Genotype*Location + Year*Location | 0.345 | 0.355 | 0.882 | 0.72 | 5 | 2 | 9 | Yes |
| | 50035 | 4.7 | 1.04 | 191 | Location | Genotype + Year + Genotype*Year + Genotype*Location + Year*Location | 0.159 | 0.198 | 0.565 | 0.74 | 5 | 2 | 9 | Yes |
| **Berry pH** | 42050 | 3.2 | 0.22 | 109 | Location | Genotype + Year + Year*Location | 0.164 | 0.321 | 0.487 | 0.76 | 6 | 3 | 14 | Yes |
| | 50001 | 3.22 | 0.17 | 62 | None | Genotype + Year | 0.017 | 0.005 | 0.022 | 0.84 | 4 | 3 | 7 | No |
| | 50013 | 3.21 | 0.2 | 64 | Location | Genotype + Genotype*Year + Genotype*Location + Year*Location | 0.101 | 0.23 | 0.447 | 0.54 | 5 | 3 | 13 | Yes |
| | 50015 | 3.25 | 0.18 | 56 | Location | Genotype + Year | 0.152 | 0.387 | 0.539 | 0.66 | 5 | 3 | 11 | Yes |
| | 50025 | 3.1 | 0.14 | 93 | Location | Genotype + Genotype*Location + Year*Location | 0.003 | 0.005 | 0.012 | 0.46 | 5 | 2 | 9 | No |
| | 50035 | 3.1 | 0.14 | 191 | None | Genotype + Genotype*Year + Genotype*Location + Year*Location | 0.006 | 0.004 | 0.016 | 0.64 | 5 | 2 | 8 | No |
| **Berry sugar content** | 42050 | 21.02 | 1.5 | 109 | None | Genotype + Genotype*Location + Year*Location | 0.721 | 1.07 | 1.915 | 0.75 | 6 | 3 | 14 | No |
| | 50001 | 17.86 | 3.29 | 29 | None | Genotype + Year*Location | 0.105 | 0.251 | 0.356 | 0.57 | 2 | 3 | 4 | Yes |
| | 50013 | 20.6 | 1.57 | 64 | Location | Genotype + Genotype*Location + Year*Location | 0.648 | 1.171 | 2.145 | 0.65 | 5 | 3 | 12 | No |
| | 50015 | 21.23 | 1.62 | 56 | None | Genotype + Genotype*Location + Year*Location | 0.889 | 1.107 | 2.399 | 0.71 | 5 | 3 | 10 | No |
| | 50025 | 23.23 | 1.82 | 92 | Location | Genotype + Genotype*Year + Genotype*Location + Year*Location | 0.892 | 1.042 | 2.495 | 0.7 | 4 | 2 | 8 | No |
| | 50035 | 22.44 | 1.69 | 191 | Location | Genotype + Year + Genotype*Year + Genotype*Location + Year*Location | 0.216 | 0.238 | 0.587 | 0.71 | 5 | 2 | 9 | Yes |

| Trait | Population ID | Mean | Standard deviation | Number of genotype | Fixe effect | Random effects | Genetic variance | Residual variance | Phenotypic variance | H² | Number of year | Number of location | Number of environment | Normalisation |
|---|---|---|---|---|---|---|---|---|---|---|---|---|---|---|
| **Date of 50% budbreak** | 42050 | 102.96 | 8.64 | 113 | Location | Gentoype + Year + Genotype*Year | 0.027 | 0.188 | 0.265 | 0.25 | 4 | 3 | 6 | Yes |
| | 50001 | 100.81 | 5.76 | 62 | None | Genotype + Year | 0.143 | 0.266 | 0.409 | 0.49 | 4 | 2 | 4 | Yes |
| | 50013 | 101.75 | 13.73 | 66 | Location | Genotype + Year | 0.334 | 0.213 | 0.547 | 0.75 | 2 | 2 | 3 | Yes |
| | 50015 | 99.5 | 12.1 | 58 | Location | Genotype + Year + Genotype*Location | 0.3 | 0.039 | 0.49 | 0.76 | 2 | 2 | 3 | Yes |
| | 50025 | 100.15 | 5.72 | 93 | Location | Genotype + Genotype*Location + Year*Location | 0.117 | 0.174 | 0.333 | 0.7 | 5 | 2 | 9 | Yes |
| | 50035 | 101.64 | 5.94 | 190 | Location | Gentoype + Gentoype*Year + Gentoype*Location + Year*Location | 0.121 | 0.154 | 0.346 | 0.68 | 5 | 2 | 8 | Yes |
| **Cluster compactness** | 42050 | 2.96 | 1.16 | 109 | None | Genotype + Year*Location | 0.4 | 0.773 | 1.173 | 0.76 | 6 | 3 | 14 | No |
| | 50001 | 3.22 | 1.6 | 61 | Location | Genotype + Year + Gentoype*Year | 0.131 | 2.2E-12 | 2.14E+0 | 0.85 | 3 | 2 | 3 | No |
| | 50013 | 3.45 | 1.39 | 63 | None | Genotype + Year*Location | 0.909 | 0.909 | 1.819 | 0.85 | 5 | 3 | 12 | No |
| | 50015 | 2.24 | 1.11 | 56 | None | Gentoype + Genotype*Location + Year*Location | 0.125 | 0.69 | 1.017 | 0.36 | 5 | 3 | 10 | No |
| | 50025 | 2.89 | 0.99 | 92 | None | Genotype + Year | 0.263 | 0.703 | 0.967 | 0.57 | 4 | 2 | 5 | No |
| | 50035 | 2.9 | 1.13 | 188 | None | Genotype*Year | 0.397 | 0.711 | 1.108 | 0.62 | 4 | 1 | 4 | No |
| **Number of clusters per plant** | 42050 | 11.69 | 7.72 | 109 | Location | Gentoype + Gentoype*Location + Year*Location | 0.256 | 0.256 | 0.559 | 0.81 | 6 | 3 | 14 | Yes |
| | 50013 | 14.31 | 7.87 | 64 | Location | Gentoype + Year*Location | 0.253 | 0.284 | 0.537 | 0.83 | 5 | 3 | 12 | Yes |
| | 50015 | 11.48 | 7.12 | 56 | Location | Gentoype + Gentoype*Location + Year*Location | 0.221 | 0.238 | 0.536 | 0.77 | 5 | 3 | 10 | Yes |
| | 50025 | 36.47 | 10.76 | 91 | None | Genotype + Year | 43.57 | 66.28 | 109.85 | 0.67 | 4 | 1 | 4 | No |
| | 50035 | 26.51 | 10.87 | 188 | None | Gentoype + Year | 0.293 | 0.418 | 0.711 | 0.68 | 4 | 1 | 4 | Yes |
| **Total cluster weight per plant** | 42050 | 1.13 | 1.07 | 109 | Location | Genotype + Genotype*Location + Year*Location | 0.228 | 0.276 | 0.585 | 0.74 | 6 | 3 | 14 | Yes |
| | 50013 | 1.73 | 1.36 | 63 | Location | Genotype + Year*Location | 0.323 | 0.405 | 0.728 | 0.82 | 5 | 3 | 12 | Yes |
| | 50015 | 0.79 | 0.73 | 56 | None | Genotype + Genotype*Year + Genotype*Location + Year*Location | 0.22 | 0.269 | 0.72 | 0.61 | 5 | 3 | 10 | Yes |

| Trait | Population ID | Mean | Standard deviation | Number of genotype | Fixe effect | Random effects | Genetic variance | Residual variance | Phenotypic variance | $H^2$ | Numbe r of year | Numbe r of location | Number of environmen t | Normalisation |
|---|---|---|---|---|---|---|---|---|---|---|---|---|---|---|
| | 50025 | 3.06 | 1.4 | 91 | None | Gentoype + Year | 0.717 | 0.885 | 1.602 | 0.71 | 4 | 1 | 4 | No |
| | 50035 | 2.02 | 1.14 | 188 | None | Genotype + Year | 0.364 | 0.377 | 0.741 | 0.74 | 4 | 1 | 4 | Yes |
| **Date of 50% flowering** | 42050 | 153.97 | 5.65 | 113 | Location | Gentoype + Year | 0.085 | 0.194 | 0.279 | 0.65 | 4 | 3 | 8 | Yes |
| | 50001 | 158.94 | 5.48 | 61 | None | Genotype + Year + Genotype*Location | 0.049 | 0.196 | 0.347 | 0.16 | 4 | 2 | 4 | Yes |
| | 50013 | 160.24 | 5.17 | 65 | Location | Genoytpe + Year + Genotype*Year + Genotype*Location | 3.95E-2 | 5.31E-5 | 4.83E-1 | 0.12 | 3 | 3 | 5 | Yes |
| | 50015 | 164.67 | 3.63 | 58 | Location | Genotype + Year | 4.788 | 2.827 | 7.615 | 0.8 | 3 | 2 | 4 | No |
| | 50025 | 154.36 | 9.43 | 93 | Location | Genotype + Year + Genotype*Location + Year*Location | 0.038 | 0.063 | 0.116 | 0.68 | 5 | 2 | 5 | Yes |
| | 50035 | 156.89 | 9.73 | 190 | None | Genotype + Year + Genotype*Location + Year*Location | 0.056 | 0.081 | 0.153 | 0.76 | 5 | 2 | 9 | Yes |
| **Date of 50% veraison** | 42050 | 222.33 | 12.73 | 110 | Location | Genotype + Year + Gentoype*Location + Year*Location | 0.122 | 0.068 | 0.204 | 0.86 | 6 | 3 | 14 | Yes |
| | 50001 | 218.04 | 7.92 | 62 | None | Gentoype + Gentoype*Year + Gentoype*Location + Year*Location | 37.404 | 3.638 | 47.275 | 0.86 | 4 | 3 | 7 | No |
| | 50013 | 223 | 13.63 | 64 | Location | Genotype + Year + Genotype*Location + Year*Location | 0.113 | 0.066 | 0.195 | 0.86 | 5 | 3 | 13 | Yes |
| | 50015 | 224.92 | 13.89 | 58 | Location | Genotype + Year + Genotype*Location + Year*Location | 29.302 | 11.469 | 43.237 | 0.92 | 5 | 3 | 11 | Yes |
| | 50025 | 216.29 | 9.44 | 93 | Location | Genotype + Year + Genotype*Location + Year*Location | 0.445 | 0.114 | 0.578 | 0.93 | 4 | 2 | 8 | Yes |
| | 50035 | 213.14 | 8.18 | 190 | None | Genotype + Year + Genotype*Year + Genotype*Location + Year*Location | 0.269 | 0.083 | 0.388 | 0.93 | 5 | 2 | 9 | Yes |
| **Mean berry weight** | 42050 | 1.82 | 0.43 | 109 | Location | Genotype + Genotype*Location + Year*Location | 0.106 | 0.041 | 0.155 | 0.91 | 6 | 3 | 14 | No |
| | 50001 | 1.71 | 0.33 | 40 | None | Genotype + Year + Genotype*Location | 0.08 | 0.014 | 0.103 | 0.83 | 3 | 2 | 4 | No |
| | 50013 | 1.46 | 0.35 | 64 | Location | Genotype + Genotype*Location + Year*Location | 0.07 | 0.029 | 0.108 | 0.88 | 5 | 3 | 12 | No |
| | 50015 | 1.39 | 0.32 | 56 | Location | Gentoype + Genotype*Location + Year*Location | 0.057 | 0.028 | 0.098 | 0.83 | 5 | 3 | 10 | No |
| | 50025 | 1.28 | 0.44 | 93 | None | Genotype + Genotype*Location + Year*Location | 0.024 | 0.046 | 0.084 | 0.62 | 5 | 2 | 7 | No |

| Trait | Population ID | Mean | Standard deviation | Number of genotype | Fixe effect | Random effects | Genetic variance | Residual variance | Phenotypic variance | H² | Number of year | Number of location | Number of environment | Normalisation |
|---|---|---|---|---|---|---|---|---|---|---|---|---|---|---|
| | 50035 | 1.21 | 0.3 | 191 | None | Genotype + Year + Genotype*Year + Genotype*Location + Year*Location | 0.038 | 0.02 | 0.064 | 0.87 | 4 | 2 | 7 | No |
| **Mean cluster weight** | 42050 | 85.48 | 54.92 | 109 | Location | Genotype + Genotype*Location + Year*Location | 0.198 | 0.35 | 0.623 | 0.68 | 6 | 3 | 14 | Yes |
| | 50013 | 114.97 | 73.89 | 63 | Location | Genotype + Genotype*Location + Year*Location | 0.29 | 0.312 | 0.661 | 0.78 | 4 | 3 | 11 | Yes |
| | 50015 | 62.64 | 47.68 | 56 | None | Genotype + Genotype*Location + Year*Location | 0.134 | 0.301 | 0.548 | 0.54 | 4 | 3 | 9 | Yes |
| | 50025 | 164.46 | 95.43 | 93 | Location | Genotype + Genotype*Location + Year*Location | 0.194 | 0.179 | 0.5 | 0.68 | 5 | 2 | 8 | Yes |
| | 50035 | 127.86 | 70.35 | 191 | Location | Genotype + Genotype*Year + Genotype*Location + Year*Location | 0.327 | 0.249 | 0.682 | 0.81 | 5 | 2 | 8 | Yes |
| **Harvest date** | 42050 | 269.59 | 15.45 | 110 | Location | Genotype + Year + Genotype*Location + Year*Location | 0.088 | 0.087 | 0.215 | 0.72 | 5 | 3 | 12 | Yes |
| | 50001 | 261.02 | 12.34 | 36 | Location | Genotype + Year + Genotype*Year + Genotype*Location | 77.23 | 11.15 | 124.41 | 0.78 | 4 | 3 | 7 | No |
| | 50013 | 271.94 | 14.46 | 64 | Location | Genotype + Year + Genotype*Location + Year*Location | 0.095 | 0.115 | 0.236 | 0.75 | 5 | 3 | 13 | Yes |
| | 50015 | 270.67 | 15.52 | 57 | Location | Genotype + Year + Genotype*Location + Year*Location | 0.157 | 0.079 | 0.277 | 0.84 | 5 | 3 | 11 | Yes |
| | 50025 | 261.94 | 13.82 | 93 | Location | Genotype + Year + Genotype*Location | 0.301 | 0.148 | 0.555 | 0.74 | 5 | 2 | 9 | Yes |
| | 50035 | 238.33 | 25.12 | 191 | Location | Genotype + Year + Genotype*Location | 0.093 | 0.186 | 0.279 | 0.74 | 5 | 2 | 9 | Yes |
| **Yield per square meter** | 42050 | 0.51 | 0.46 | 109 | Location | Genotype + Genotype*Location + Year*Location | 0.241 | 0.296 | 0.622 | 0.74 | 6 | 3 | 14 | Yes |
| | 50001 | 0.88 | 0.62 | 30 | Location | Genotype | 0.183 | 0.227 | 0.41 | 0.67 | 2 | 2 | 4 | Yes |
| | 50013 | 0.8 | 0.61 | 63 | Location | Gentoype + Genotype*Location + Year*Location | 0.326 | 0.394 | 0.761 | 0.79 | 5 | 3 | 12 | Yes |
| | 50015 | 0.37 | 0.34 | 56 | None | Gentoype + Genotype*Year + Genotype*Location + Year*Location | 0.222 | 0.268 | 0.722 | 0.6 | 5 | 3 | 10 | Yes |
| | 50025 | 1.29 | 0.52 | 93 | None | Gentoype + Genotype*Location + Year*Location | 0.067 | 0.11 | 0.226 | 0.62 | 5 | 2 | 9 | No |
| | 50035 | 0.85 | 0.45 | 191 | None | Gentoype + Genotype*Location + Year*Location | 0.059 | 0.086 | 0.152 | 0.79 | 5 | 2 | 9 | No |

**Supplementary Table 5.** Associations between family-specific $H^2$ and design variables. Abbreviations: $H^2$, broad-sense heritability; rho, Spearman rank-correlation coefficient.

| variable | rho | p value | n |
|---|---|---|---|
| Number.of.genotype | 0.035 | 0.763 | 75 |
| Number.of.year | 0.156 | 0.182 | 75 |
| Number.of.location | 0.176 | 0.131 | 75 |
| Number.of.environment | 0.26 | 0.024 | 75 |

**Supplementary Table 6.** Within-family broad-sense heritability and number of catalogued family QTLs for the 75 estimable family-by-trait combinations. H² was not associated with QTL count (Spearman rho = 0.171, P = 0.143), and the H² distributions with and without a detected QTL did not differ significantly (Wilcoxon P = 0.057). Abbreviations: H², broad-sense heritability; QTL, quantitative trait locus. Counts include the three suggestive family-50015 QTLs retained in the catalogue.

| Trait | Trait code | Population | H2 | N QTL | any QTL | N genotypes | N years | N locations | N environments |
|---|---|---|---|---|---|---|---|---|---|
| Berry pH | BERRY_pH | 42050 | 0.76 | 1 | TRUE | 109 | 6 | 3 | 14 |
| Berry pH | BERRY_pH | 50001 | 0.84 | 1 | TRUE | 62 | 4 | 3 | 7 |
| Berry pH | BERRY_pH | 50013 | 0.54 | 0 | FALSE | 64 | 5 | 3 | 13 |
| Berry pH | BERRY_pH | 50015 | 0.66 | 1 | TRUE | 56 | 5 | 3 | 11 |
| Berry pH | BERRY_pH | 50025 | 0.46 | 0 | FALSE | 93 | 5 | 2 | 9 |
| Berry pH | BERRY_pH | 50035 | 0.64 | 2 | TRUE | 191 | 5 | 2 | 8 |
| Berry titratable acidity | BER_TA_g | 42050 | 0.84 | 1 | TRUE | 109 | 6 | 3 | 14 |
| Berry titratable acidity | BER_TA_g | 50001 | 0.84 | 0 | FALSE | 63 | 3 | 3 | 6 |
| Berry titratable acidity | BER_TA_g | 50013 | 0.67 | 0 | FALSE | 64 | 5 | 3 | 12 |
| Berry titratable acidity | BER_TA_g | 50015 | 0.75 | 1 | TRUE | 56 | 5 | 3 | 10 |
| Berry titratable acidity | BER_TA_g | 50025 | 0.72 | 4 | TRUE | 93 | 5 | 2 | 9 |
| Berry titratable acidity | BER_TA_g | 50035 | 0.74 | 2 | TRUE | 191 | 5 | 2 | 9 |
| Date of 50% budbreak | BUD_DATE | 42050 | 0.25 | 0 | FALSE | 113 | 4 | 3 | 6 |
| Date of 50% budbreak | BUD_DATE | 50001 | 0.49 | 0 | FALSE | 62 | 4 | 2 | 4 |
| Date of 50% budbreak | BUD_DATE | 50013 | 0.75 | 0 | FALSE | 66 | 2 | 2 | 3 |
| Date of 50% budbreak | BUD_DATE | 50015 | 0.76 | 0 | FALSE | 58 | 2 | 2 | 3 |
| Date of 50% budbreak | BUD_DATE | 50025 | 0.7 | 2 | TRUE | 93 | 5 | 2 | 9 |
| Date of 50% budbreak | BUD_DATE | 50035 | 0.68 | 3 | TRUE | 190 | 5 | 2 | 8 |
| Date of 50% flowering | FLO_50 | 42050 | 0.65 | 3 | TRUE | 113 | 4 | 3 | 8 |
| Date of 50% flowering | FLO_50 | 50001 | 0.16 | 0 | FALSE | 61 | 4 | 2 | 4 |
| Date of 50% flowering | FLO_50 | 50013 | 0.12 | 0 | FALSE | 65 | 3 | 3 | 5 |
| Date of 50% flowering | FLO_50 | 50015 | 0.8 | 1 | TRUE | 58 | 3 | 2 | 4 |
| Date of 50% flowering | FLO_50 | 50025 | 0.68 | 1 | TRUE | 93 | 5 | 2 | 5 |
| Date of 50% flowering | FLO_50 | 50035 | 0.76 | 2 | TRUE | 190 | 5 | 2 | 9 |
| Harvest date | HARVEST_DATE | 42050 | 0.72 | 0 | FALSE | 110 | 5 | 3 | 12 |

| Trait | Trait code | Population | H2 | N QTL | any QTL | N genotypes | N years | N locations | N environments |
|---|---|---|---|---|---|---|---|---|---|
| Harvest date | HARVEST_DATE | 50001 | 0.78 | 0 | FALSE | 36 | 4 | 3 | 7 |
| Harvest date | HARVEST_DATE | 50013 | 0.75 | 0 | FALSE | 64 | 5 | 3 | 13 |
| Harvest date | HARVEST_DATE | 50015 | 0.84 | 2 | TRUE | 57 | 5 | 3 | 11 |
| Harvest date | HARVEST_DATE | 50025 | 0.74 | 1 | TRUE | 93 | 5 | 2 | 9 |
| Harvest date | HARVEST_DATE | 50035 | 0.74 | 2 | TRUE | 191 | 5 | 2 | 9 |
| Cluster compactness | MORPHO_OIV_204 | 42050 | 0.76 | 0 | FALSE | 109 | 6 | 3 | 14 |
| Cluster compactness | MORPHO_OIV_204 | 50001 | 0.85 | 0 | FALSE | 61 | 3 | 2 | 3 |
| Cluster compactness | MORPHO_OIV_204 | 50013 | 0.85 | 2 | TRUE | 63 | 5 | 3 | 12 |
| Cluster compactness | MORPHO_OIV_204 | 50015 | 0.36 | 0 | FALSE | 56 | 5 | 3 | 10 |
| Cluster compactness | MORPHO_OIV_204 | 50025 | 0.57 | 0 | FALSE | 92 | 4 | 2 | 5 |
| Cluster compactness | MORPHO_OIV_204 | 50035 | 0.62 | 4 | TRUE | 188 | 4 | 1 | 4 |
| Number of clusters per plant | NB_CLUST_PLANT | 42050 | 0.81 | 0 | FALSE | 109 | 6 | 3 | 14 |
| Number of clusters per plant | NB_CLUST_PLANT | 50013 | 0.83 | 0 | FALSE | 64 | 5 | 3 | 12 |
| Number of clusters per plant | NB_CLUST_PLANT | 50015 | 0.77 | 1 | TRUE | 56 | 5 | 3 | 10 |
| Number of clusters per plant | NB_CLUST_PLANT | 50025 | 0.67 | 0 | FALSE | 91 | 4 | 1 | 4 |
| Number of clusters per plant | NB_CLUST_PLANT | 50035 | 0.68 | 3 | TRUE | 188 | 4 | 1 | 4 |
| Mean berry weight | SBER_W_g | 42050 | 0.91 | 0 | FALSE | 109 | 6 | 3 | 14 |
| Mean berry weight | SBER_W_g | 50001 | 0.83 | 0 | FALSE | 40 | 3 | 2 | 4 |
| Mean berry weight | SBER_W_g | 50013 | 0.88 | 1 | TRUE | 64 | 5 | 3 | 12 |
| Mean berry weight | SBER_W_g | 50015 | 0.83 | 1 | TRUE | 56 | 5 | 3 | 10 |
| Mean berry weight | SBER_W_g | 50025 | 0.62 | 1 | TRUE | 93 | 5 | 2 | 7 |
| Mean berry weight | SBER_W_g | 50035 | 0.87 | 3 | TRUE | 191 | 4 | 2 | 7 |
| Mean cluster weight | SCLUST_W | 42050 | 0.68 | 1 | TRUE | 109 | 6 | 3 | 14 |
| Mean cluster weight | SCLUST_W | 50013 | 0.78 | 2 | TRUE | 63 | 4 | 3 | 11 |
| Mean cluster weight | SCLUST_W | 50015 | 0.54 | 0 | FALSE | 56 | 4 | 3 | 9 |
| Mean cluster weight | SCLUST_W | 50025 | 0.68 | 0 | FALSE | 93 | 5 | 2 | 8 |
| Mean cluster weight | SCLUST_W | 50035 | 0.81 | 2 | TRUE | 191 | 5 | 2 | 8 |
| Berry sugar content | TSS | 42050 | 0.75 | 0 | FALSE | 109 | 6 | 3 | 14 |

| Trait | Trait code | Population | H2 | N QTL | any QTL | N genotypes | N years | N locations | N environments |
|---|---|---|---|---|---|---|---|---|---|
| Berry sugar content | TSS | 50001 | 0.57 | 0 | FALSE | 29 | 2 | 3 | 4 |
| Berry sugar content | TSS | 50013 | 0.65 | 2 | TRUE | 64 | 5 | 3 | 12 |
| Berry sugar content | TSS | 50015 | 0.71 | 0 | FALSE | 56 | 5 | 3 | 10 |
| Berry sugar content | TSS | 50025 | 0.7 | 3 | TRUE | 92 | 4 | 2 | 8 |
| Berry sugar content | TSS | 50035 | 0.71 | 4 | TRUE | 191 | 5 | 2 | 9 |
| Date of 50% veraison | VER_50 | 42050 | 0.86 | 2 | TRUE | 110 | 6 | 3 | 14 |
| Date of 50% veraison | VER_50 | 50001 | 0.86 | 1 | TRUE | 62 | 4 | 3 | 7 |
| Date of 50% veraison | VER_50 | 50013 | 0.86 | 0 | FALSE | 64 | 5 | 3 | 13 |
| Date of 50% veraison | VER_50 | 50015 | 0.92 | 0 | FALSE | 58 | 5 | 3 | 11 |
| Date of 50% veraison | VER_50 | 50025 | 0.93 | 1 | TRUE | 93 | 4 | 2 | 8 |
| Date of 50% veraison | VER_50 | 50035 | 0.93 | 2 | TRUE | 190 | 5 | 2 | 9 |
| Yield per square meter | YIELD_OIV_504 | 42050 | 0.74 | 1 | TRUE | 109 | 6 | 3 | 14 |
| Yield per square meter | YIELD_OIV_504 | 50001 | 0.67 | 0 | FALSE | 30 | 2 | 2 | 4 |
| Yield per square meter | YIELD_OIV_504 | 50013 | 0.79 | 2 | TRUE | 63 | 5 | 3 | 12 |
| Yield per square meter | YIELD_OIV_504 | 50015 | 0.6 | 1 | TRUE | 56 | 5 | 3 | 10 |
| Yield per square meter | YIELD_OIV_504 | 50025 | 0.62 | 0 | FALSE | 93 | 5 | 2 | 9 |
| Yield per square meter | YIELD_OIV_504 | 50035 | 0.79 | 4 | TRUE | 191 | 5 | 2 | 9 |
| Total cluster weight per plant | YIELD_PLANT | 42050 | 0.74 | 1 | TRUE | 109 | 6 | 3 | 14 |
| Total cluster weight per plant | YIELD_PLANT | 50013 | 0.82 | 2 | TRUE | 63 | 5 | 3 | 12 |
| Total cluster weight per plant | YIELD_PLANT | 50015 | 0.61 | 1 | TRUE | 56 | 5 | 3 | 10 |
| Total cluster weight per plant | YIELD_PLANT | 50025 | 0.71 | 0 | FALSE | 91 | 4 | 1 | 4 |
| Total cluster weight per plant | YIELD_PLANT | 50035 | 0.74 | 1 | TRUE | 188 | 4 | 1 | 4 |

**Supplementary Table 7.** Characteristics of the multi-parental genetic map, including marker number, map length, and mean and maximum marker spacing per chromosome. Abbreviations: cM, centimorgan.

| Chromosome | Marker count | Genetic length (cM) | Mean spacing between two markers (cM) | Max spacing between two markers (cM) |
|---|---|---|---|---|
| 1 | 2,895 | 139.26 | 0.05 | 4.1 |
| 2 | 1,746 | 189.55 | 0.11 | 8.01 |
| 3 | 2,311 | 133.25 | 0.06 | 4.64 |
| 4 | 3,330 | 111.3 | 0.03 | 3.26 |
| 5 | 4,787 | 110.41 | 0.02 | 3.33 |
| 6 | 2,877 | 132.6 | 0.05 | 8.28 |
| 7 | 6,041 | 124.31 | 0.02 | 4.37 |
| 8 | 4,275 | 124.7 | 0.03 | 4.77 |
| 9 | 2,407 | 175.07 | 0.07 | 7.48 |
| 10 | 3,162 | 114.56 | 0.04 | 3.52 |
| 11 | 2,947 | 117.83 | 0.04 | 7.46 |
| 12 | 2,127 | 152.46 | 0.07 | 5.82 |
| 13 | 3,441 | 126.34 | 0.04 | 7.34 |
| 14 | 5,247 | 139.2 | 0.03 | 7.34 |
| 15 | 2,228 | 134.75 | 0.06 | 4.31 |
| 16 | 1,808 | 219.22 | 0.12 | 4.57 |
| 17 | 2,626 | 119.83 | 0.05 | 5.62 |
| 18 | 5,677 | 138.83 | 0.02 | 3.72 |
| 19 | 2,270 | 153.62 | 0.07 | 5.1 |
| TOTAL | 62,202 | 2,657.1 | 0.05 | 8.28 |

**Supplementary Table 8.** QTLs detected in the six largest ResDur families using BLUP-based analyses. For each trait, the table lists the associated marker, chromosome, genetic and physical positions, confidence intervals, effect sizes, and explained variance. BUD_DATE: date of 50% of budbreak, FLO_50: date of 50% flowering, VER_50: date of 50% veraison, SBER_W_g: mean berry weight, TSS: berry sugar content, BERRY_pH: berry pH, BER_TA_g: berry titratable acidity, HARVEST_DATE: harvest date, MORPHO_OIV_204: cluster compactness, NB_CLUST_PLANT: number of clusters per plant, YIELD_PLANT: total cluster weight per plant, SCLUST_W: mean cluster weight, YIELD_OIV_504: yield per square meter. Abbreviations: QTL, quantitative trait locus; BLUP, best linear unbiased predictor; Chr, chromosome; Pos, genetic position; CI, confidence interval; LOD, logarithm of odds; LOD5PC, 5% genome-wide LOD threshold; P, probability value; var, variance; AC, BC, AD and BD, estimated four-way-cross genotype effects; Mb, megabase; bp, base pair. The 79-interval catalogue retains three suggestive family-50015 QTLs: berry weight/chr2 ($P = 0.084$), cluster weight per plant/chr5 ($P = 0.091$) and yield/m²/chr5 ($P = 0.071$). They are not significant at 5%. Appended rows (QTL077-QTL079) use the physical extent of markers within 95% Bayesian credible intervals; AC-BD are observed genotype-class phenotype means. $P < 0.001$ denotes source permutation P values rounded to zero.

| Pop | Trait | Marker | Chr | Pos | ci.low | ci.high | LOD | pval | LOD5 PC | %var | AC effect | BC effect | AD effect | BD effect | Physical position (Mb) | CI physical position (Mb) | Physical position (bp) |
|---|---|---|---|---|---|---|---|---|---|---|---|---|---|---|---|---|---|
| 42050 | Flo_50 | chr2_4654231 | 2 | 20 | 14.3 | 26.7 | 4.66 | 0.04 | 4.54 | 18 | 0.05 | 0.17 | -0.16 | 0.1 | 4.7 | 0.5_4.7 | 4,654,231 |
| 42050 | Flo_50 | chr7_20484677 | 7 | 41.46 | 35.3 | 59.6 | 5.22 | 0.01 | 4.54 | 20 | 0.14 | 0 | -0.15 | -0.01 | 20.5 | 17.5_24.3 | 20,484,677 |
| 42050 | VER_50 | chr8_17943691 | 8 | 31.43 | 18.6 | 37.6 | 4.72 | 0.04 | 4.56 | 19 | -0.09 | -0.15 | 0.16 | 0.17 | 17.9 | 14.9_19.2 | 17,943,691 |
| 42050 | SCLUST_W | chr8_19121576 | 8 | 37.15 | 1.4 | 42.4 | 5.34 | 0.01 | 4.58 | 19 | -0.14 | -0.14 | 0.18 | 0.19 | 19.1 | 2.1_21.2 | 19,121,576 |
| 42050 | YIELD_PLANT | chr11_7704414 | 11 | 22.38 | 7.6 | 32.9 | 4.69 | 0.04 | 4.6 | 19 | -0.13 | -0.05 | -0.03 | 0.38 | 7.7 | 3.2_11.3 | 7,704,414 |
| 42050 | YIELD_OIV_504 | chr11_7704414 | 11 | 22.38 | 7.6 | 32.9 | 4.69 | 0.04 | 4.59 | 19 | -0.14 | -0.05 | -0.03 | 0.4 | 7.7 | 3.2_11.3 | 7,704,414 |
| 42050 | Flo_50 | chr14_21445121 | 14 | 40.5 | 31.9 | 65.3 | 4.6 | 0.04 | 4.54 | 18 | 0.06 | 0.03 | -0.19 | -0.02 | 21.4 | 11.3_29.7 | 21,445,121 |
| 42050 | VER_50 | chr14_24993381 | 14 | 47.65 | 45.3 | 63.9 | 4.79 | 0.03 | 4.58 | 21 | 0.19 | 0.01 | -0.2 | -0.1 | 25 | 24.3_28.8 | 24,993,381 |
| 42050 | BERRY_pH | chr17_6431844 | 17 | 20.95 | 19.5 | 34.8 | 5.65 | 0.01 | 4.55 | 22 | -0.26 | 0.08 | -0.07 | 0.24 | 6.4 | 5.9_9.6 | 6,431,844 |
| 42050 | BER_TA_g | chr17_12686850 | 17 | 43.34 | 21.9 | 45.2 | 5.25 | 0.01 | 4.52 | 21 | 6.66 | 5.85 | 6.02 | 5.67 | 12.7 | 6.8_16.6 | 12,686,850 |
| 50001 | VER_50 | chr16_16627115 | 16 | 22.4 | 20.2 | 29.1 | 14.27 | 0 | 4.63 | 67 | 214.61 | 214.32 | 222.28 | 224.19 | 16.6 | 15.9_18.2 | 16,627,115 |
| 50001 | BERRY_pH | chr16_17821664 | 16 | 29.13 | 27.6 | 31.4 | 7.49 | 0 | 4.67 | 44 | 3.25 | 3.31 | 3.11 | 3.17 | 17.8 | 17.1_19.1 | 17,821,664 |
| 50013 | TSS | chr7_650262 | 7 | 0 | 0 | 5.3 | 5.59 | 0.01 | 4.61 | 37 | 20.88 | 20.2 | 20.13 | 21.01 | 0.7 | 0.7_3.9 | 650,262 |
| 50013 | SBER_W_g | chr7_26566117 | 7 | 58.8 | 14 | 62.3 | 4.59 | 0.03 | 4.35 | 31 | 1.33 | 1.7 | 1.43 | 1.5 | 26.6 | 4.7_28.7 | 26,566,117 |

| Pop | Trait | Marker | Chr | Pos | ci.low | ci.high | LOD | pval | LOD5 PC | %var | AC effect | BC effect | AD effect | BD effect | Physical position (Mb) | CI physical position (Mb) | Physical position (bp) |
|---|---|---|---|---|---|---|---|---|---|---|---|---|---|---|---|---|---|
| 50013 | MORPHO_OIV_204 | chr14_22456692 | 14 | 30.74 | 25.5 | 54.4 | 4.61 | 0.05 | 4.59 | 32 | 2.89 | 2.8 | 4 | 3.67 | 22.5 | 20.7_30.8 | 22,456,692 |
| 50013 | YIELD_PLANT | chr14_24313848 | 14 | 36.88 | 24.6 | 47.4 | 6.34 | 0 | 4.54 | 41 | -0.16 | -0.55 | 0.3 | 0.26 | 24.3 | 20.7_29.6 | 24,313,848 |
| 50013 | YIELD_OIV_504 | chr14_24313848 | 14 | 36.88 | 23.7 | 47.4 | 6.15 | 0 | 4.56 | 40 | -0.17 | -0.54 | 0.3 | 0.24 | 24.3 | 19.5_29.6 | 24,313,848 |
| 50013 | SCLUST_W | chr14_29030452 | 14 | 46.53 | 33.4 | 47.4 | 5.55 | 0.01 | 4.59 | 37 | -0.12 | -0.47 | 0.37 | 0.1 | 29 | 22.7_29.6 | 29,030,452 |
| 50013 | TSS | chr17_5915174 | 17 | 16.67 | 10.5 | 17.6 | 5.26 | 0.02 | 4.61 | 35 | 20.37 | 21.44 | 20.19 | 20.67 | 5.9 | 4_6 | 5,915,174 |
| 50013 | MORPHO_OIV_204 | chr17_6904099 | 17 | 21.94 | 7.9 | 31.6 | 5.52 | 0.01 | 4.59 | 37 | 2.86 | 3.78 | 4.3 | 3.23 | 6.9 | 6.1_9.2 | 6,904,099 |
| 50013 | YIELD_PLANT | chr17_9209449 | 17 | 33.34 | 21.9 | 38.6 | 4.95 | 0.02 | 4.54 | 34 | -0.37 | -0.05 | 0.39 | -0.14 | 9.2 | 6.8_10.9 | 9,209,449 |
| 50013 | SCLUST_W | chr17_9209449 | 17 | 33.34 | 21.1 | 43.9 | 4.62 | 0.05 | 4.59 | 32 | -0.28 | -0.09 | 0.38 | -0.15 | 9.2 | 6.5_18.8 | 9,209,449 |
| 50013 | YIELD_OIV_504 | chr17_9209449 | 17 | 33.34 | 21.9 | 38.6 | 5.27 | 0.01 | 4.56 | 36 | -0.38 | -0.06 | 0.39 | -0.16 | 9.2 | 6.7_10.9 | 9,209,449 |
| 50015 | SBER_W_g | chr2_6620861 | 2 | 28.31 | 20.8 | 36.8 | 4.39 | 0.084 | 4.73 | 32 | 1.48 | 1.322 | 1.6 | 1.24 | 6.6 | 5.3_7.7 | 6,620,861 |
| 50015 | YIELD_PLANT | chr5_4624702 | 5 | 15.1 | 1.9 | 23.6 | 4.42 | 0.091 | 4.75 | 32 | -0.1 | -0.016 | -0.17 | 0.39 | 4.6 | 1.2_6.3 | 4,624,702 |
| 50015 | YIELD_OIV_504 | chr5_4624702 | 5 | 15.1 | 1.9 | 23.6 | 4.5 | 0.071 | 4.69 | 33 | -0.1 | -0.019 | -0.18 | 0.4 | 4.6 | 0.7_6.3 | 4,624,702 |
| 50015 | HARVEST_DATE | chr8_13368232 | 8 | 15.12 | 12.3 | 28.3 | 4.88 | 0.04 | 4.74 | 39 | 0.29 | 39.58 | 0.38 | -0.03 | 13.4 | 0.2_20.7 | 13,368,232 |
| 50015 | BER_TA_g | chr13_5256578 | 13 | 25.68 | 23.8 | 33.2 | 6.61 | 0 | 4.74 | 34 | 5.55 | 45.11 | 5.7 | 4.98 | 5.3 | 3.9_7.1 | 5,256,578 |
| 50015 | BERRY_pH | chr13_5407720 | 13 | 30.4 | 23.8 | 31.3 | 5.62 | 0.01 | 4.68 | 44 | -0.14 | 43.71 | -0.27 | 0.33 | 5.4 | 4.4_5.8 | 5,407,720 |
| 50025 | TSS | chr1_22.127183 | 1 | 51.04 | 49.5 | 58.3 | 6.15 | 0 | 4.51 | 27 | 23.74 | 23.49 | 22.84 | 22.85 | 22.1 | 20.4_23.7 | 22,127,183 |
| 50025 | BUD_DATE | chr4_10.837709 | 4 | 39.8 | 38.3 | 42.9 | 4.71 | 0.03 | 4.51 | 21 | -0.2 | 0.1 | -0.09 | 0.13 | 10.8 | 8.1_15.9 | 10,837,709 |
| 50025 | BER_TA_g | chr6_19.84535 | 6 | 46.91 | 36.1 | 50 | 4.98 | 0.02 | 4.51 | 22 | 0.08 | -0.44 | 0.17 | 0.12 | 19.8 | 15.4_21.2 | 19,845,350 |
| 50025 | BUD_DATE | chr7_2.329771 | 7 | 13.92 | 0 | 53.1 | 4.68 | 0.04 | 4.51 | 21 | -0.15 | 0.24 | -0.05 | 0.04 | 2.3 | 0.4_18.2 | 2,329,771 |
| 50025 | BER_TA_g | chr14_25.063969 | 14 | 43.82 | 33.5 | 61.9 | 5.41 | 0.01 | 4.51 | 24 | -0.08 | -0.3 | 0.26 | 0.18 | 25.1 | 21_29.8 | 25,063,969 |
| 50025 | Flo_50 | chr14_28.358345 | 14 | 55.68 | 27.3 | 61.9 | 5.53 | 0.01 | 4.49 | 24 | 0.03 | -0.06 | 0.12 | -0.77 | 28.4 | 17.9_29.8 | 28,358,345 |

| Pop | Trait | Marker | Chr | Pos | ci.low | ci.high | LOD | pval | LOD5 PC | %var | AC effect | BC effect | AD effect | BD effect | Physical position (Mb) | CI physical position (Mb) | Physical position (bp) |
|---|---|---|---|---|---|---|---|---|---|---|---|---|---|---|---|---|---|
| 50025 | SBER_W_g | chr17_1.67817 | 17 | 3.61 | 0 | 24.3 | 5.29 | 0.01 | 4.54 | 23 | 1.4 | 1.29 | 1.28 | 1.22 | 1.7 | 6.6_10.3 | 1,678,170 |
| 50025 | TSS | chr17_8.404173 | 17 | 27.4 | 23.8 | 38.8 | 6.03 | 0 | 4.51 | 26 | 22.76 | 22.88 | 23.31 | 23.72 | 8.4 | 1.9_4.3 | 8,404,173 |
| 50025 | BER_TA_g | chr17_9.976907 | 17 | 33.59 | 8.2 | 38.8 | 4.84 | 0.03 | 4.51 | 22 | 0.21 | -0.35 | 0.21 | 0.08 | 10 | 1.6_4.9 | 9,976,907 |
| 50025 | TSS | chr18_22.957478 | 18 | 66.61 | 58.4 | 70.7 | 4.71 | 0.04 | 4.51 | 21 | 23.41 | 22.92 | 24.11 | 23.01 | 23 | 12.3_29.9 | 22,957,478 |
| 50025 | BER_TA_g | chr22_13.941485 | 22 | 5.16 | 2.1 | 7.2 | 5.26 | 0.01 | 4.51 | 24 | -0.9 | -1.3 | NA | NA | 13.9 | 12.5_15.2 | 13,941,485 |
| 50025 | HARVEST_DATE | chr22_14.399259 | 22 | 6.19 | 5.2 | 7.2 | 8.94 | 0 | 4.5 | 36 | 0.68 | 0.78 | NA | NA | 14.4 | 14_15.2 | 14,399,259 |
| 50025 | VER_50 | chr22_14.399259 | 22 | 6.19 | 5.7 | 7.2 | 8.69 | 0 | 4.5 | 36 | 0.87 | 1.74 | NA | NA | 14.4 | 14.2_15.2 | 14,399,259 |
| 50035 | MORPHO_OIV_204 | chr1_3384952 | 1 | 12.64 | 6.5 | 21.5 | 9.32 | 0 | 4.41 | 21 | 2.65 | 2.69 | 3.08 | 3.16 | 3.4 | 2_5.8 | 3,384,952 |
| 50035 | BUD_DATE | chr2_1981206 | 2 | 6.99 | 1.9 | 19.9 | 5.79 | 0 | 4.44 | 13 | -0.13 | 0.11 | -0.06 | 0.11 | 2 | 2.4_5.7 | 1,981,206 |
| 50035 | VER_50 | chr2_4308310 | 2 | 15.59 | 9.4 | 23.1 | 5.2 | 0.01 | 4.46 | 12 | -0.22 | 0.02 | -0.03 | 0.27 | 4.3 | 2.3_7.4 | 4,308,310 |
| 50035 | HARVEST_DATE | chr2_5194530 | 2 | 19.89 | 8.3 | 23.1 | 5.54 | 0.01 | 4.51 | 13 | -0.19 | 0 | -0.08 | 0.28 | 5.2 | 0.9_5.7 | 5,194,530 |
| 50035 | MORPHO_OIV_204 | chr2_5459030 | 2 | 22.05 | 3.8 | 32 | 5.14 | 0.01 | 4.41 | 12 | 2.66 | 3.01 | 2.82 | 3.12 | 5.5 | 5.2_6.2 | 5,459,030 |
| 50035 | SCLUST_W | chr2_5735229 | 2 | 23.12 | 21 | 31.5 | 8.58 | 0 | 4.47 | 19 | -0.32 | -0.04 | 0 | 0.33 | 5.7 | 5.7_6.2 | 5,735,229 |
| 50035 | YIELD_OIV_504 | chr2_5735229 | 2 | 23.12 | 15.6 | 36.1 | 5.02 | 0.02 | 4.41 | 12 | 0.75 | 0.84 | 0.84 | 0.97 | 5.7 | 5.5_7.2 | 5,735,229 |
| 50035 | SBER_W_g | chr2_6954620 | 2 | 36.05 | 22.3 | 38.2 | 7.19 | 0 | 4.44 | 16 | 1.11 | 1.21 | 1.21 | 1.33 | 7 | 8.5_16.4 | 6,954,620 |
| 50035 | NB_CLUST_PLANT | chr3_8345848 | 3 | 47.6 | 36.8 | 54.6 | 4.67 | 0.03 | 4.42 | 11 | -0.02 | 0.16 | -0.23 | 0.08 | 8.3 | 4.9_20.7 | 8,345,848 |
| 50035 | BERRY_pH | chr4_16273785 | 4 | 40.66 | 35 | 46.8 | 7.42 | 0 | 4.46 | 17 | 3.08 | 3.09 | 3.14 | 3.11 | 16.3 | 8.1_18.9 | 16,273,785 |
| 50035 | BER_TA_g | chr4_18432101 | 4 | 45.23 | 21 | 46.8 | 7.29 | 0 | 4.37 | 17 | 0.17 | 0.01 | -0.17 | 0 | 18.4 | 5.2_18.9 | 18,432,101 |
| 50035 | TSS | chr6_5647483 | 6 | 19.36 | 11.6 | 21.8 | 6.31 | 0 | 4.4 | 15 | 0.17 | 0.07 | -0.05 | -0.22 | 5.6 | 3.6_7.3 | 5,647,483 |
| 50035 | SBER_W_g | chr7_3577589 | 7 | 27.43 | 23.9 | 61.1 | 6.18 | 0 | 4.44 | 14 | 1.25 | 1.27 | 1.1 | 1.21 | 3.6 | 2.5_22.1 | 3,577,589 |
| 50035 | NB_CLUST_PLANT | chr7_3885537 | 7 | 29.85 | 19.4 | 35.5 | 5.58 | 0.01 | 4.42 | 13 | -0.17 | -0.03 | -0.04 | 0.29 | 3.9 | 1.8_5.8 | 3,885,537 |

| Pop | Trait | Marker | Chr | Pos | ci.low | ci.high | LOD | pval | LOD5 PC | %var | AC effect | BC effect | AD effect | BD effect | Physical position (Mb) | CI physical position (Mb) | Physical position (bp) |
|---|---|---|---|---|---|---|---|---|---|---|---|---|---|---|---|---|---|
| 50035 | Flo_50 | chr7_7706452 | 7 | 38.45 | 30.9 | 46.2 | 4.95 | 0.02 | 4.4 | 12 | 0.12 | -0.01 | -0.02 | -0.09 | 7.7 | 4.1_20.6 | 7,706,452 |
| 50035 | TSS | chr8_18317705 | 8 | 46.25 | 46 | 50.6 | 5.85 | 0 | 4.4 | 14 | 0.14 | -0.24 | 0 | 0.08 | 18.3 | 17.7_20.3 | 18,317,705 |
| 50035 | BUD_DATE | chr9_2251882 | 9 | 9.95 | 4.3 | 30.9 | 7.94 | 0 | 4.44 | 18 | 0.04 | 0.16 | -0.16 | 0.12 | 2.3 | 0.9_5.5 | 2,251,882 |
| 50035 | Flo_50 | chr9_2251882 | 9 | 9.95 | 3.2 | 50.6 | 5.32 | 0.01 | 4.4 | 12 | 0 | 0.71 | -0.08 | 0.11 | 2.3 | 0.1_13.8 | 2,251,882 |
| 50035 | TSS | chr9_2720967 | 9 | 16.13 | 6.5 | 30.1 | 7.9 | 0 | 4.4 | 18 | 0.08 | -0.16 | 0.14 | -0.28 | 2.7 | 0.1_2.7 | 2,720,967 |
| 50035 | SBER_W_g | chr11_13720591 | 11 | 35.49 | 30.9 | 48.7 | 5.83 | 0 | 4.44 | 14 | 1.25 | 1.27 | 1.22 | 1.09 | 13.7 | 9.6_19.5 | 13,720,591 |
| 50035 | YIELD_OIV_504 | chr11_12722104 | 11 | 35.49 | 30.9 | 49.2 | 4.48 | 0.05 | 4.41 | 11 | 0.9 | 0.88 | 0.87 | 0.72 | 12.7 | 9.6_19.4 | 12,722,104 |
| 50035 | YIELD_PLANT | chr14_22997551 | 14 | 41.41 | 30.9 | 51.1 | 5.86 | 0 | 4.4 | 16 | 0.04 | -0.03 | 0.29 | -0.27 | 23 | 17.4_27.4 | 22,997,551 |
| 50035 | YIELD_OIV_504 | chr14_26703281 | 14 | 49.2 | 32 | 53.5 | 5.52 | 0.01 | 4.41 | 13 | 0.89 | 0.82 | 0.96 | 0.75 | 26.7 | 18_28.4 | 26,703,281 |
| 50035 | NB_CLUST_PLANT | chr14_29728423 | 14 | 56.73 | 35.5 | 57.8 | 4.71 | 0.03 | 4.42 | 11 | 0.16 | -0.08 | 0.16 | -0.18 | 29.7 | 20.6_30.3 | 29,728,423 |
| 50035 | BUD_DATE | chr16_1668279 | 16 | 4.84 | 0 | 25 | 5.73 | 0 | 4.44 | 13 | 0.1 | 0.1 | -0.1 | -0.12 | 1.7 | 2_16.8 | 1,668,279 |
| 50035 | SCLUST_W | chr16_3881623 | 16 | 6.99 | 3.8 | 25 | 4.72 | 0.03 | 4.47 | 11 | -0.02 | 0.19 | 0 | -0.23 | 3.9 | 14.4_16.7 | 3,881,623 |
| 50035 | YIELD_OIV_504 | chr16_4129661 | 16 | 6.99 | 0 | 11.6 | 4.77 | 0.03 | 4.41 | 11 | 0.78 | 0.94 | 0.84 | 0.78 | 4.1 | 0.8_16.7 | 4,129,661 |
| 50035 | BER_TA_g | chr16_16336324 | 16 | 13.44 | 10.5 | 15.3 | 7.08 | 0 | 4.37 | 16 | 0.23 | -0.09 | -0.07 | 0.01 | 16.3 | 15_17.3 | 16,336,324 |
| 50035 | BERRY_pH | chr16_16575285 | 16 | 13.44 | 2.2 | 15.3 | 5.8 | 0 | 4.46 | 13 | 3.06 | 3.11 | 3.12 | 3.11 | 16.6 | 15_16.8 | 16,575,285 |
| 50035 | HARVEST_DATE | chr16_16792671 | 16 | 14.52 | 11.3 | 16.4 | 11.66 | 0 | 4.51 | 25 | 0.45 | -0.02 | -0.23 | -0.16 | 16.8 | 15.4_16.8 | 16,792,671 |
| 50035 | VER_50 | chr16_16601209 | 16 | 14.52 | 11.3 | 15.1 | 20.77 | 0 | 4.46 | 40.4 | 0.64 | -0.1 | -0.22 | -0.16 | 16.6 | 0.1_17.6 | 16,601,209 |
| 50035 | MORPHO_OIV_204 | chr16_17154341 | 16 | 16.13 | 0 | 19.4 | 4.54 | 0.04 | 4.41 | 11 | 2.6 | 3.03 | 2.83 | 2.96 | 17.2 | 0.1_15.3 | 17,154,341 |
| 50035 | MORPHO_OIV_204 | chr18_10502733 | 18 | 42.5 | 37.7 | 46.5 | 6.22 | 0 | 4.41 | 14 | 2.69 | 2.77 | 2.98 | 3.19 | 10.5 | 7.9_34.3 | 10,502,733 |
| 50035 | TSS | chr18_25972037 | 18 | 63.21 | 53 | 64.5 | 11.21 | 0 | 4.4 | 24 | -0.14 | 0.13 | -0.13 | 0.33 | 26 | 12.7_31.1 | 25,972,037 |
| 50015 | Flo_50 | chr17_6478604 | 17 | 13.21 | 9.4 | 13.2 | 7.44 | <0.001 | 4.56 | 47.6 | 166.067 | 163.436 | 165.406 | 163.327 | 6.479 | 5.509-6.479 | 6,478,604 |

| Pop | Trait | Marker | Chr | Pos | ci.low | ci.high | LOD | pval | LOD5 PC | %var | AC effect | BC effect | AD effect | BD effect | Physical position (Mb) | CI physical position (Mb) | Physical position (bp) |
|---|---|---|---|---|---|---|---|---|---|---|---|---|---|---|---|---|---|
| 50015 | HARVEST_DATE | chr18_25972084 | 18 | 77.42 | 69.9 | 83.1 | 4.88 | 0.037 | 4.72 | 35.1 | -0.189 | -0.155 | 0.301 | 0.161 | 25.972 | 22.756-29.601 | 25,972,084 |
| 50015 | NB_CLUST_PLANT | chr17_3327363 | 17 | 0.94 | 0 | 3.8 | 5.62 | 0.007 | 4.68 | 39.2 | -0.205 | 0.127 | -0.329 | 0.412 | 3.327 | 0.004-4.479 | 3,327,363 |

**Supplementary Table 9.** Environment-specific QTLs detected in the 42050, 50025, and 50035 families. Results include trait, environment, associated marker, chromosome, confidence interval, significance, effect sizes, and explained variance. Abbreviations: QTL, quantitative trait locus; Chr, chromosome; Pos, genetic position; CI, confidence interval; LOD, logarithm of odds; LOD5PC, 5% genome-wide LOD threshold; P, probability value; var, variance; AC, BC, AD and BD, estimated four-way-cross genotype effects; Mb, megabase; bp, base pair. Trait codes are defined in Supplementary Table 8.

| Pop | Trait | Data | Marker | Chr | Pos | ci.low | ci.high | LOD | pval | %var | AC effect | BC effect | AD effect | BD effect | Physical position (Mb) | CI physical position (Mb) | Physical position (bp) |
|---|---|---|---|---|---|---|---|---|---|---|---|---|---|---|---|---|---|
| 42050 | FLO_50 | 2016 Bordeaux | chr2_4654438 | 2 | 19.5 | 14 | 26.7 | 5.24 | 0.02 | 22 | 154.75 | 153.11 | 152.22 | 155.23 | 4.7 | 3.6_5.7 | 4,654,438 |
| 42050 | TSS | 2014 Colmar | chr2_8423545 | 2 | 43 | 41 | 50.6 | 4.57 | 0.05 | 19 | 19.78 | 20.79 | 19.99 | 20.88 | 8.4 | 7.1_16.8 | 8,423,545 |
| 42050 | BER_TA_g | 2014 Bordeaux | chr2_9273485 | 2 | 43.4 | 41 | 49.1 | 5.05 | 0.02 | 23 | 5.53 | 6.84 | 5.25 | 6.1 | 9.3 | 7.1_16.3 | 9,273,485 |
| 42050 | HARVEST_DATE | 2014 Bordeaux | chr2_9273485 | 2 | 43.4 | 13 | 50.6 | 4.68 | 0.04 | 21 | 259.4 | 252.59 | 258.46 | 252.93 | 9.3 | 3.5_16.8 | 9,273,485 |
| 42050 | HARVEST_DATE | 2012 Bordeaux | chr2_9314224 | 2 | 43.9 | 44 | 50.1 | 12.1 | 0 | 50 | 269.83 | 257.45 | 267.08 | 254.78 | 9.3 | 9.3_16.4 | 9,314,224 |
| 42050 | TSS | 2011 Colmar | chr2_10156144 | 2 | 45.8 | 43 | 50.6 | 5 | 0.02 | 26 | 20.52 | 21.22 | 20.11 | 21.57 | 10.2 | 8.4_16.8 | 10,156,144 |
| 42050 | BERRY_pH | 2014 Bordeaux | chr2_15968391 | 2 | 48.7 | 41 | 55.8 | 4.63 | 0.04 | 22 | 3.46 | 3.26 | 3.43 | 3.3 | 16 | 7.1_18.6 | 15,968,391 |
| 42050 | HARVEST_DATE | 2013 Bordeaux | chr2_16362304 | 2 | 49.6 | 41 | 53 | 5.3 | 0.02 | 24 | 271.69 | 269 | 274.77 | 266.64 | 16.4 | 7.1_17.5 | 16,362,304 |
| 42050 | FLO_50 | 2014Colmar | chr2_16356757 | 2 | 50.1 | 42 | 55.8 | 5.29 | 0.01 | 18 | 154.44 | 154.97 | 156.31 | 155.67 | 16.4 | 7.6_18.6 | 16,356,757 |
| 42050 | MORPHO_OIV_204 | 2013 Colmar | chr3_5072604 | 3 | 27.2 | 24 | 44.3 | 4.69 | 0.04 | 20 | 2.67 | 3.14 | 2.2 | 2.19 | 5.1 | 4.2_13.5 | 5,072,604 |
| 42050 | VER_50 | 2011 Colmar | chr3_18751058 | 3 | 46.7 | 38 | 47.1 | 5.13 | 0.02 | 26 | 212.83 | 216.37 | 215 | 220.27 | 18.8 | 8.3_20.7 | 18,751,058 |
| 42050 | BERRY_pH | 2013 Colmar | chr6_192578 | 6 | 0 | 0 | 13.5 | 5.98 | 0 | 24 | 3.08 | 3.29 | 3.1 | 3.1 | 0.2 | 0.2_1.2 | 192,578 |
| 42050 | TSS | 2013 Bordeaux | chr7_1387152 | 7 | 0.95 | 0 | 14.3 | 4.71 | 0.04 | 23 | 20.04 | 22 | 21.11 | 21.2 | 1.4 | 1.4_6.7 | 1,387,152 |
| 42050 | BER_TA_g | 2012 Bordeaux | chr7_17625440 | 7 | 34.8 | 28 | 39.6 | 4.94 | 0.03 | 25 | 4.16 | 3.98 | 3.14 | 3.74 | 17.6 | 16_20 | 17,625,440 |
| 42050 | FLO_50 | 2015 Bordeaux | chr7_23844580 | 7 | 56.2 | 33 | 59.6 | 4.87 | 0.03 | 21 | 149.2 | 148.08 | 147.16 | 147.65 | 23.8 | 17.5_24.3 | 23,844,580 |
| 42050 | HARVEST_DATE | 2014 Colmar | chr8_17964909 | 8 | 31.4 | 29 | 39.5 | 5.95 | 0.01 | 19 | 280.87 | 278.72 | 284.93 | 284.79 | 18 | 17.5_20.1 | 17,964,909 |
| 42050 | VER_50 | 2014 Colmar | chr8_17964909 | 8 | 31.4 | 29 | 39.5 | 5.91 | 0 | 20 | 225.42 | 223.83 | 228.43 | 229.48 | 18 | 17.5_20.1 | 17,964,909 |
| 42050 | HARVEST_DATE | 2011 Colmar | chr8_18580531 | 8 | 34.8 | 28 | 36.2 | 4.66 | 0.04 | 24 | 259.04 | 260.4 | 268 | 264.44 | 18.6 | 17.2_19 | 18,580,531 |
| 42050 | VER_50 | 2011 Colmar | chr8_18591835 | 8 | 34.8 | 29 | 37.6 | 4.66 | 0.04 | 24 | 213.67 | 214.55 | 220.55 | 218.61 | 18.6 | 17.5_19.2 | 18,591,835 |
| 42050 | YIELD_PLANT | 2013 Colmar | chr11_7704414 | 11 | 22.4 | 18 | 23.3 | 5.84 | 0.01 | 24 | 0.95 | 1.15 | 1.28 | 2.31 | 7.7 | 6.6_7.7 | 7,704,414 |
| 42050 | YIELD_OIV_504 | 2013 Colmar | chr11_7704414 | 11 | 22.4 | 18 | 23.3 | 5.84 | 0 | 24 | 0.41 | 0.49 | 0.55 | 0.99 | 7.7 | 6.6_7.7 | 7,704,414 |
| 42050 | YIELD_PLANT | 2012 Colmar | chr11_7689237 | 11 | 23.3 | 22 | 31.4 | 6.39 | 0 | 26 | 1.61 | 1.56 | 1.26 | 3.29 | 7.7 | 7.7_10 | 7,689,237 |

| Pop | Trait | Data | Marker | Chr | Pos | ci.low | ci.high | LOD | pval | %var | AC effect | BC effect | AD effect | BD effect | Physical position (Mb) | CI physical position (Mb) | Physical position (bp) |
|---|---|---|---|---|---|---|---|---|---|---|---|---|---|---|---|---|---|
| 42050 | YIELD_OIV_504 | 2012 Colmar | chr11_7689237 | 11 | 23.3 | 22 | 31.4 | 6.39 | 0 | 26 | 0.69 | 0.67 | 0.54 | 1.41 | 7.7 | 7.7_10 | 7,689,237 |
| 42050 | VER_50 | 2013 Colmar | chr14_24359078 | 14 | 46.2 | 45 | 63.9 | 5.66 | 0.01 | 23 | 240.53 | 238.09 | 234.65 | 236.18 | 24.4 | 24.3_28.8 | 24,359,078 |
| 42050 | FLO_50 | 2015 Bordeaux | chr14_24894927 | 14 | 47.2 | 32 | 63.9 | 5.56 | 0.01 | 23 | 148.48 | 149.14 | 147 | 147.35 | 24.9 | 11.1_28.8 | 24,894,927 |
| 42050 | HARVEST_DATE | 2014 Colmar | chr14_25018215 | 14 | 48.6 | 45 | 63.9 | 4.69 | 0.04 | 23 | 285.34 | 282.9 | 278.47 | 279.2 | 25 | 24.3_28.8 | 25,018,215 |
| 42050 | VER_50 | 2014 Colmar | chr14_25018215 | 14 | 48.6 | 46 | 63.9 | 5.04 | 0.02 | 21 | 229.49 | 226.9 | 223.88 | 224.3 | 25 | 24.4_28.8 | 25,018,215 |
| 42050 | VER_50 | 2012 Colmar | chr14_25018215 | 14 | 48.6 | 45 | 63.9 | 5.22 | 0.02 | 22 | 231.55 | 228.86 | 225 | 226.59 | 25 | 24.3_28.8 | 25,018,215 |
| 42050 | VER_50 | 2011 Colmar | chr14_26062240 | 14 | 53.4 | 26 | 63.9 | 4.73 | 0.04 | 24 | 219.16 | 217.8 | 212.82 | 213.5 | 26.1 | 7_28.8 | 26,062,240 |
| 42050 | SCLUST_W | 2012 Colmar | chr14_26718892 | 14 | 54.8 | 55 | 65.3 | 14.7 | 0 | 50 | 44.8 | 61.7 | 127.47 | 124.05 | 26.7 | 26.1_29.6 | 26,718,892 |
| 42050 | YIELD_PLANT | 2012 Colmar | chr14_26542290 | 14 | 55.3 | 53 | 65.3 | 13.9 | 0 | 48 | 0.8 | 1.14 | 2.84 | 2.79 | 26.5 | 25.2_29.7 | 26,542,290 |
| 42050 | YIELD_OIV_504 | 2012 Colmar | chr14_26542290 | 14 | 55.3 | 53 | 65.3 | 13.9 | 0 | 48 | 0.34 | 0.49 | 1.22 | 1.2 | 26.5 | 25.2_29.7 | 26,542,290 |
| 42050 | VER_50 | 2011 Bordeaux | chr14_27414299 | 14 | 58.1 | 45 | 67.7 | 4.85 | 0.03 | 30 | 198.21 | 200.12 | 192.8 | 193.89 | 27.4 | 24.3_29.8 | 27,414,299 |
| 42050 | MORPHO_OIV_204 | 2012 Colmar | chr14_30124820 | 14 | 69.1 | 44 | 73.4 | 5.57 | 0.01 | 23 | 2.58 | 2.33 | 3.37 | 3.6 | 30.1 | 24.1_30.6 | 30,124,820 |
| 42050 | BERRY_pH | 2013 Bordeaux | chr17_6401199 | 17 | 21 | 20 | 30 | 7.44 | 0 | 34 | 3.32 | 3.49 | 3.31 | 3.57 | 6.4 | 5.9_8 | 6,401,199 |
| 42050 | SBER_W_g | 2013 Colmar | chr17_9623444 | 17 | 34.8 | 23 | 44.3 | 5.48 | 0.01 | 22 | 2 | 1.62 | 1.75 | 1.49 | 9.6 | 7.1_16.1 | 9,623,444 |
| 42050 | BER_TA_g | 2011 Colmar | chr17_12804250 | 17 | 43.8 | 34 | 44.3 | 5.76 | 0.01 | 29 | 7.15 | 5.74 | 6.03 | 5.67 | 12.8 | 9.2_16.1 | 12,804,250 |
| 42050 | YIELD_PLANT | 2011 Colmar | chr18_4207724 | 18 | 15.7 | 4.3 | 21 | 4.73 | 0.05 | 25 | 1.32 | 1.47 | 1.75 | 2.83 | 4.2 | 2.1_6.9 | 4,207,724 |
| 42050 | YIELD_OIV_504 | 2011 Colmar | chr18_4207724 | 18 | 15.7 | 4.3 | 21 | 4.73 | 0.04 | 25 | 0.57 | 0.63 | 0.75 | 1.22 | 4.2 | 2.1_6.9 | 4,207,724 |
| 50025 | BERRY_pH | 2018 Colmar | chr1_3235109 | 1 | 8.76 | 0 | 15 | 4.45 | 0.05 | 20 | 3.32 | 3.27 | 3.27 | 3.22 | 3.2 | 0.1_22 | 3,235,109 |
| 50025 | SCLUST_W | 2020 Pully | chr1_4838421 | 1 | 17.5 | 0 | 54.1 | 5.36 | 0.01 | 24 | 148.96 | 178.92 | 227.58 | 244.48 | 4.8 | 19.8_23.8 | 4,838,421 |
| 50025 | TSS | 2020 Pully | chr1_20615518 | 1 | 55.2 | 51 | 57.7 | 5.07 | 0.02 | 23 | 24.5 | 24.31 | 22.9 | 23.36 | 20.6 | 19.8_23.8 | 20,615,518 |
| 50025 | TSS | 2020 Colmar | chr1_20615518 | 1 | 55.2 | 50 | 58.3 | 5.09 | 0.02 | 23 | 24.11 | 23.11 | 21.77 | 22.6 | 20.6 | 0.1_4.2 | 20,615,518 |
| 50025 | SCLUST_W | 2017 Pully | chr2_1154614 | 2 | 34.6 | 16 | 34.6 | 4.85 | 0.03 | 22 | 217.86 | 124.12 | 217.53 | 188.25 | 11.5 | 5.2_13.1 | 11,546,140 |
| 50025 | BER_TA_g | 2021 Pully | chr5_6566337 | 5 | 25.8 | 13 | 28.9 | 4.81 | 0.03 | 22 | 4.42 | 4.01 | 5.52 | 4.51 | 6.6 | 4.8_7.6 | 6,566,337 |
| 50025 | BER_TA_g | 2020 Pully | chr6_10985359 | 6 | 33 | 25 | 36.1 | 7.58 | 0 | 32 | 4.03 | 3.76 | 4.04 | 4.81 | 11 | 7.6_15.2 | 10,985,359 |
| 50025 | BERRY_pH | 2020 Pully | chr6_1095527 | 6 | 33 | 6.7 | 37.6 | 4.55 | 0.05 | 21 | 3.1 | 3.14 | 3.12 | 3.01 | 11 | 3.1_16.9 | 10,955,270 |

| Pop | Trait | Data | Marker | Chr | Pos | ci.low | ci.high | LOD | pval | %var | AC effect | BC effect | AD effect | BD effect | Physical position (Mb) | CI physical position (Mb) | Physical position (bp) |
|---|---|---|---|---|---|---|---|---|---|---|---|---|---|---|---|---|---|
| 50025 | BER_TA_g | 2017 Pully | chr6_19146646 | 6 | 42.8 | 42 | 50 | 5.08 | 0.02 | 23 | 4.31 | 3.66 | 4.3 | 4.51 | 19.1 | 18.6_21.2 | 19,146,646 |
| 50025 | BUD_DATE | 2019 Colmar | chr7_8561107 | 7 | 35.1 | 12 | 38.2 | 4.95 | 0.03 | 22 | 106.88 | 106 | 101.88 | 107.53 | 8.6 | 1.8_10.5 | 8,561,107 |
| 50025 | NB_CLUST_PLANT | 2019 Colmar | chr11_2004494 | 11 | 1.55 | 0 | 2.1 | 6.98 | 0 | 30 | 39.82 | 34.6 | 36.98 | 26.32 | 2 | 0.7_2.1 | 2,004,494 |
| 50025 | SBER_W_g | 2019 Colmar | chr11_3886671 | 11 | 17.2 | 17 | 24.5 | 5.96 | 0 | 28 | 1.14 | 1.46 | 1.1 | 1.4 | 3.9 | 3.6_7.5 | 3,886,671 |
| 50025 | BER_TA_g | 2020 Pully | chr11_6640874 | 11 | 22.9 | 18 | 26 | 4.8 | 0.03 | 22 | 4.45 | 3.76 | 4.61 | 4.01 | 6.6 | 4.6_7.7 | 6,640,874 |
| 50025 | MORPHO_OIV_204 | 2018 Colmar | chr11_7450023 | 11 | 24.5 | 16 | 42.5 | 4.77 | 0.02 | 22 | 2.82 | 3.11 | 3.1 | 1.89 | 7.5 | 3.5_16.4 | 7,450,023 |
| 50025 | HARVEST_DATE | 2018 Pully | chr13_12299916 | 13 | 34.5 | 28 | 43.3 | 5.03 | 0.02 | 22 | 257.96 | 247.96 | 248.96 | 255.96 | 12.3 | 8.5_17.6 | 12,299,916 |
| 50025 | YIELD_OIV_504 | 2021 Pully | chr13_1676434 | 13 | 43.3 | 28 | 53.6 | 5.22 | 0.02 | 23 | 1.73 | 1.14 | 1.54 | 1.46 | 16.8 | 8.5_24.7 | 16,764,340 |
| 50025 | BER_TA_g | 2018 Pully | chr14_2260473 | 14 | 38.7 | 38 | 44.9 | 6.69 | 0 | 29 | 4.27 | 4.03 | 4.79 | 5.11 | 22.6 | 22.4_25.6 | 22,604,730 |
| 50025 | TSS | 2018 Colmar | chr14_2260473 | 14 | 38.7 | 20 | 45.9 | 4.56 | 0.04 | 21 | 23.93 | 21.87 | 24.01 | 23.13 | 22.6 | 9_25.8 | 22,604,730 |
| 50025 | FLO_50 | 2017 Pully | chr14_24426282 | 14 | 43.3 | 26 | 61.9 | 4.88 | 0.03 | 22 | 154.57 | 152.83 | 154.4 | 153.77 | 24.4 | 14.5_29.8 | 24,426,282 |
| 50025 | BERRY_pH | 2018 Pully | chr14_25063969 | 14 | 43.8 | 38 | 61.4 | 5.58 | 0.01 | 25 | 3.09 | 3.15 | 3 | 2.98 | 25.1 | 22.4_29.9 | 25,063,969 |
| 50025 | BER_TA_g | 2018 Colmar | chr14_25039998 | 14 | 43.8 | 32 | 61.9 | 5.4 | 0.01 | 24 | 3.58 | 3.33 | 4 | 3.89 | 25 | 20.8_29.8 | 25,039,998 |
| 50025 | YIELD_PLANT | 2018 Colmar | chr14_26950023 | 14 | 50 | 45 | 57.2 | 4.84 | 0.03 | 22 | 1.41 | 3.29 | 1.93 | 2.6 | 27 | 25.1_28.9 | 26,950,023 |
| 50025 | YIELD_OIV_504 | 2018 Colmar | chr14_26880052 | 14 | 50 | 45 | 57.2 | 4.84 | 0.03 | 22 | 0.61 | 1.42 | 0.83 | 1.12 | 26.9 | 25.1_28.9 | 26,880,052 |
| 50025 | NB_CLUST_PLANT | 2020 Colmar | chr14_27381773 | 14 | 51 | 38 | 54.1 | 4.55 | 0.05 | 21 | 40.24 | 40.12 | 31.05 | 35.49 | 27.4 | 22.4_28.1 | 27,381,773 |
| 50025 | SCLUST_W | 2018 Colmar | chr14_28255143 | 14 | 54.7 | 48 | 59.3 | 4.59 | 0.05 | 21 | 35.39 | 71.03 | 61.46 | 54.54 | 28.3 | 26.1_29.3 | 28,255,143 |
| 50025 | FLO_50 | 2019 Pully | chr14_28269129 | 14 | 55.2 | 29 | 61.9 | 6.31 | 0 | 27 | 169.45 | 168.44 | 170.14 | 168.92 | 28.3 | 19.3_29.8 | 28,269,129 |
| 50025 | FLO_50 | 2020 Pully | chr14_28358345 | 14 | 55.7 | 30 | 61.9 | 4.64 | 0.04 | 21 | 149.45 | 149.17 | 151.81 | 149.4 | 28.4 | 20.1_29.8 | 28,358,345 |
| 50025 | FLO_50 | 2018 Pully | chr14_28607101 | 14 | 55.7 | 53 | 61.9 | 7.5 | 0 | 32 | 150.55 | 149.47 | 152.12 | 150 | 28.6 | 27.8_29.8 | 28,607,101 |
| 50025 | SCLUST_W | 2021 Pully | chr16_18292826 | 16 | 0 | 0 | 0 | 8.35 | 0.05 | 34 | 233.6 | NA | 238.93 | NA | 18.3 | 18.3_18.3 | 18,292,826 |
| 50025 | YIELD_PLANT | 2019 Colmar | chr16_18292826 | 16 | 0 | 0 | 0 | 4.94 | 0.02 | 22 | 3.25 | NA | 2.79 | NA | 18.3 | 18.3_18.3 | 18,292,826 |
| 50025 | SBER_W_g | 2020 Colmar | chr17_1875578 | 17 | 3.61 | 3.1 | 8.8 | 6.62 | 0 | 28 | 1.42 | 1.12 | 1.09 | 1.01 | 1.9 | 6.6_10.3 | 1,875,578 |
| 50025 | SBER_W_g | 2018 Colmar | chr17_167817 | 17 | 3.61 | 3.6 | 8.8 | 7.24 | 0 | 31 | 1.7 | 1.25 | 1.25 | 1.14 | 1.7 | 1_6.8 | 1,678,170 |

| Pop | Trait | Data | Marker | Chr | Pos | ci.low | ci.high | LOD | pval | %var | AC effect | BC effect | AD effect | BD effect | Physical position (Mb) | CI physical position (Mb) | Physical position (bp) |
|---|---|---|---|---|---|---|---|---|---|---|---|---|---|---|---|---|---|
| 50025 | YIELD_PLANT | 2019 Colmar | chr17_2672246 | 17 | 6.19 | 4.1 | 8.8 | 5.56 | 0.01 | 25 | 4.28 | 2.72 | 2.51 | 3.3 | 2.7 | 1.8_4.3 | 2,672,246 |
| 50025 | YIELD_PLANT | 2020 Colmar | chr17_4088325 | 17 | 7.73 | 3.6 | 10.8 | 7.35 | 0 | 31 | 5.59 | 3.9 | 3.94 | 3.69 | 4.1 | 1.6_4.3 | 4,088,325 |
| 50025 | YIELD_OIV_504 | 2020 Colmar | chr17_4088325 | 17 | 7.73 | 3.6 | 10.8 | 7.35 | 0 | 31 | 2.4 | 1.67 | 1.26 | 1.59 | 4.1 | 4.2_10.3 | 4,088,325 |
| 50025 | TSS | 2020 Colmar | chr17_8593437 | 17 | 30 | 24 | 38.8 | 5.76 | 0 | 25 | 22.15 | 21.9 | 22.88 | 24.14 | 8.6 | 1.6_4.9 | 8,593,437 |
| 50025 | SBER_W_g | 2018 Colmar | chr18_12997484 | 18 | 59.9 | 53 | 69.7 | 5.68 | 0.01 | 25 | 1.21 | 1.48 | 0.98 | 1.2 | 13 | 11.9_29.6 | 12,997,484 |
| 50025 | BERRY_pH | 2018 Colmar | chr19_19392586 | 19 | 48 | 44 | 65.5 | 4.8 | 0.02 | 22 | 3.31 | 3.23 | 3.23 | 3.31 | 19.4 | 14_24.2 | 19,392,586 |
| 50025 | BER_TA_g | 2020 Pully | chr22_12456971 | 22 | 2.06 | 2.1 | 2.1 | 4.73 | 0.04 | 21 | 3.12 | 3.71 | NA | NA | 12.5 | 12.5_12.5 | 12,456,971 |
| 50025 | TSS | 2018 Colmar | chr22_12456971 | 22 | 2.06 | 2.1 | 6.2 | 5.4 | 0.01 | 24 | 20.79 | 22.47 | NA | NA | 12.5 | 12.5_14.4 | 12,456,971 |
| 50025 | BER_TA_g | 2021 Pully | chr22_14399259 | 22 | 6.19 | 4.6 | 6.2 | 12.3 | 0 | 46 | 5.29 | 6.01 | NA | NA | 14.4 | 13.4_14.4 | 14,399,259 |
| 50025 | BERRY_pH | 2021 Pully | chr22_14399259 | 22 | 6.19 | 5.2 | 7.2 | 11 | 0 | 43 | 2.92 | 2.72 | NA | NA | 14.4 | 14_15.2 | 14,399,259 |
| 50025 | HARVEST_DATE | 2021 Pully | chr22_14399259 | 22 | 6.19 | 5.2 | 7.2 | 6.78 | 0 | 29 | 281.1 | 275.1 | NA | NA | 14.4 | 14_15.2 | 14,399,259 |
| 50025 | VER_50 | 2020 Pully | chr22_14399259 | 22 | 6.19 | 5.2 | 7.2 | 6.58 | 0 | 28 | 223.8 | 223.84 | NA | NA | 14.4 | 14_15.2 | 14,399,259 |
| 50025 | HARVEST_DATE | 2020 Colmar | chr22_14399259 | 22 | 6.19 | 5.7 | 7.2 | 8.66 | 0 | 35 | 276.9 | 268.9 | NA | NA | 14.4 | 14.2_15.2 | 14,399,259 |
| 50025 | VER_50 | 2020 Colmar | chr22_14399259 | 22 | 6.19 | 5.7 | 7.2 | 8.93 | 0 | 36 | 218.7 | 214.7 | NA | NA | 14.4 | 14.2_15.2 | 14,399,259 |
| 50025 | BERRY_pH | 2019 Pully | chr22_14399259 | 22 | 6.19 | 2.1 | 7.2 | 4.72 | 0.03 | 21 | 2.96 | 3.06 | NA | NA | 14.4 | 12.5_15.2 | 14,399,259 |
| 50025 | HARVEST_DATE | 2019 Colmar | chr22_14399259 | 22 | 6.19 | 5.2 | 7.2 | 9.71 | 0 | 39 | 286.6 | 279.6 | NA | NA | 14.4 | 14_15.2 | 14,399,259 |
| 50025 | VER_50 | 2019 Colmar | chr22_14399259 | 22 | 6.19 | 5.7 | 7.2 | 10.5 | 0 | 41 | 227.1 | 226.12 | NA | NA | 14.4 | 14.2_15.2 | 14,399,259 |
| 50025 | HARVEST_DATE | 2018 Colmar | chr22_14399259 | 22 | 6.19 | 5.7 | 6.2 | 12.8 | 0 | 48 | 276.5 | 259.5 | NA | NA | 14.4 | 14.2_14.4 | 14,399,259 |
| 50025 | HARVEST_DATE | 2017 Pully | chr22_14399259 | 22 | 6.19 | 2.1 | 7.2 | 4.66 | 0.04 | 21 | 249.3 | 241.3 | NA | NA | 14.4 | 12.5_15.2 | 14,399,259 |
| 50025 | SBER_W_g | 2021 Pully | chr22_15204505 | 22 | 6.7 | 5.2 | 7.2 | 5.07 | 0.02 | 23 | 1.71 | 1.37 | NA | NA | 15.2 | 14_15.2 | 15,204,505 |
| 50025 | VER_50 | 2019 Pully | chr22_15204505 | 22 | 6.7 | 5.7 | 7.2 | 9.66 | 0 | 39 | 230.8 | 222.37 | NA | NA | 15.2 | 14.2_15.2 | 15,204,505 |
| 50025 | VER_50 | 2018 Colmar | chr22_15204328 | 22 | 7.22 | 5.7 | 7.2 | 7.37 | 0 | 36 | 212.3 | 201.3 | NA | NA | 15.2 | 14.2_15.2 | 15,204,328 |
| 50035 | MORPHO_OIV_204 | 2020 Colmar | chr1_3231074 | 1 | 11.6 | 6.2 | 22.3 | 5.49 | 0.01 | 13 | 2.16 | 2.47 | 3.08 | 3.3 | 3.2 | 2_6.1 | 3,231,074 |
| 50035 | MORPHO_OIV_204 | 2022 Colmar | chr1_3640606 | 1 | 14.8 | 9.4 | 22.9 | 6.28 | 0 | 14 | 2.24 | 2.26 | 2.98 | 2.69 | 3.6 | 2.9_6.3 | 3,640,606 |
| 50035 | BER_TA_g | 2020 Colmar | chr1_4838836 | 1 | 17.2 | 14 | 30.9 | 4.94 | 0.02 | 12 | 4.86 | 5.04 | 5.34 | 5.45 | 4.8 | 3.5_10 | 4,838,836 |

| Pop | Trait | Data | Marker | Chr | Pos | ci.low | ci.high | LOD | pval | %var | AC effect | BC effect | AD effect | BD effect | Physical position (Mb) | CI physical position (Mb) | Physical position (bp) |
|---|---|---|---|---|---|---|---|---|---|---|---|---|---|---|---|---|---|
| 50035 | MORPHO_OIV_204 | 2019 Colmar | chr1_4971695 | 1 | 18.3 | 0.8 | 35.2 | 5.27 | 0.01 | 14 | 3.16 | 3.13 | 3.64 | 4.05 | 5 | 0.6_12.7 | 4,971,695 |
| 50035 | BUD_DATE | 2020 Pully | chr2_1989263 | 2 | 6.99 | 0 | 23.1 | 4.95 | 0.02 | 12 | 94.57 | 96.05 | 95.12 | 96.61 | 2 | 1.1_7.4 | 1,989,263 |
| 50035 | BUD_DATE | 2022 Pully | chr2_2262696 | 2 | 8.07 | 0 | 19.9 | 4.89 | 0.02 | 12 | 99.13 | 100.79 | 97.84 | 101.05 | 2.3 | 2.1_5.7 | 2,262,696 |
| 50035 | FLO_50 | 2019 Pully | chr2_4703263 | 2 | 18 | 3 | 38.7 | 4.59 | 0.04 | 11 | 170.45 | 170.73 | 171.19 | 171.79 | 4.7 | 0.9_5.6 | 4,703,263 |
| 50035 | VER_50 | 2020 Pully | chr2_4909551 | 2 | 19.9 | 8.1 | 38.7 | 4.62 | 0.04 | 11 | 206.68 | 209.93 | 209.05 | 212.07 | 4.9 | 0.1_5.6 | 4,909,551 |
| 50035 | VER_50 | 2018 Pully | chr2_5090968 | 2 | 19.9 | 1.9 | 22.3 | 5.03 | 0.02 | 12 | 204.2 | 207.53 | 206.07 | 208.4 | 5.1 | 2.1_5.7 | 5,090,968 |
| 50035 | VER_50 | 2022 Pully | chr2_5353266 | 2 | 21.2 | 1.9 | 23.1 | 5.98 | 0 | 14 | 203.36 | 207.09 | 204.64 | 207.88 | 5.4 | 1.1_6.6 | 5,353,266 |
| 50035 | TSS | 2021 Pully | chr2_5353266 | 2 | 21.2 | 0 | 22.3 | 7.99 | 0 | 18 | 21.91 | 21.37 | 21.79 | 20.56 | 5.4 | 4.9_5.5 | 5,353,266 |
| 50035 | HARVEST_DATE | 2019 Colmar | chr2_5347333 | 2 | 21.2 | 8.3 | 23.1 | 5 | 0.02 | 13 | 268.24 | 269.97 | 266.73 | 262.21 | 5.3 | 4.9_10.8 | 5,347,333 |
| 50035 | YIELD_OIV_504 | 2020 Pully | chr2_5644020 | 2 | 22.3 | 20 | 26.9 | 8.86 | 0 | 20 | 0.75 | 0.98 | 0.91 | 1.28 | 5.6 | 4.2_7 | 5,644,020 |
| 50035 | TSS | 2022 Pully | chr2_5735229 | 2 | 23.1 | 15 | 26.9 | 4.67 | 0.03 | 11 | 23 | 22.76 | 22.73 | 21.83 | 5.7 | 4.8_18.6 | 5,735,229 |
| 50035 | SCLUST_W | 2020 Pully | chr2_5735189 | 2 | 23.1 | 20 | 26.9 | 8.63 | 0 | 19 | 129.45 | 168.23 | 163.91 | 223.85 | 5.7 | 4.9_5.5 | 5,735,189 |
| 50035 | SBER_W_g | 2020 Colmar | chr2_5735189 | 2 | 23.1 | 22 | 38.2 | 7.95 | 0 | 18 | 1 | 1.18 | 1.1 | 1.3 | 5.7 | 5.7_6.6 | 5,735,189 |
| 50035 | BERRY_pH | 2019 Pully | chr2_5735189 | 2 | 23.1 | 19 | 49.8 | 4.98 | 0.02 | 12 | 3.14 | 3.09 | 3.03 | 3.03 | 5.7 | 4.9_5.7 | 5,735,189 |
| 50035 | SCLUST_W | 2019 Pully | chr2_5735229 | 2 | 23.1 | 23 | 31.5 | 8.43 | 0 | 19 | 101.83 | 118.75 | 127.5 | 175.22 | 5.7 | 5.9_7.2 | 5,735,229 |
| 50035 | YIELD_OIV_504 | 2019 Pully | chr2_5735229 | 2 | 23.1 | 23 | 32 | 4.49 | 0.04 | 11 | 0.64 | 0.62 | 0.72 | 0.94 | 5.7 | 5.7_7.2 | 5,735,229 |
| 50035 | SCLUST_W | 2018 Pully | chr2_5735189 | 2 | 23.1 | 20 | 23.1 | 6.59 | 0 | 16 | 127.45 | 161.86 | 151.81 | 316.26 | 5.7 | 6.3_7.2 | 5,735,189 |
| 50035 | SCLUST_W | 2022 Pully | chr2_5934540 | 2 | 31.5 | 27 | 37.1 | 12.5 | 0 | 27 | 137.88 | 145.46 | 173.77 | 220.3 | 5.9 | 5.6_7.2 | 5,934,540 |
| 50035 | YIELD_OIV_504 | 2022 Pully | chr2_5934540 | 2 | 31.5 | 23 | 37.1 | 8.34 | 0 | 19 | 0.9 | 0.9 | 1.08 | 1.31 | 5.9 | 5.9_17.6 | 5,934,540 |
| 50035 | SCLUST_W | 2022 Colmar | chr2_6589967 | 2 | 32 | 18 | 38.2 | 6.9 | 0 | 16 | 57.82 | 66.49 | 73.64 | 88.87 | 6.6 | 7.1_17.4 | 6,589,967 |
| 50035 | BER_TA_g | 2022 Colmar | chr2_7732092 | 2 | 41.4 | 37 | 47.6 | 6.95 | 0 | 16 | 6.04 | 5.75 | 5.65 | 5.06 | 7.7 | 7_17.6 | 7,732,092 |
| 50035 | BER_TA_g | 2019 Colmar | chr2_12875255 | 2 | 44.4 | 38 | 47.9 | 5.94 | 0 | 15 | 6.05 | 5.53 | 5.8 | 5.17 | 12.9 | 12.9_18.6 | 12,875,255 |
| 50035 | BER_TA_g | 2020 Pully | chr2_17589512 | 2 | 47.9 | 44 | 49.8 | 7.01 | 0 | 16 | 3.67 | 3.64 | 3.74 | 4.37 | 17.6 | 0.1_5.7 | 17,589,512 |
| 50035 | BER_TA_g | 2022 Pully | chr4_5636403 | 4 | 21 | 11 | 57.9 | 5.53 | 0 | 13 | 4.54 | 4.27 | 3.93 | 3.95 | 5.6 | 2.2_23.3 | 5,636,403 |
| 50035 | BER_TA_g | 2020 Colmar | chr4_5346915 | 4 | 21.2 | 21 | 48.5 | 5 | 0.01 | 12 | 5.43 | 5.25 | 4.79 | 5.15 | 5.3 | 5.2_19.2 | 5,346,915 |

| Pop | Trait | Data | Marker | Chr | Pos | ci.low | ci.high | LOD | pval | %var | AC effect | BC effect | AD effect | BD effect | Physical position (Mb) | CI physical position (Mb) | Physical position (bp) |
|---|---|---|---|---|---|---|---|---|---|---|---|---|---|---|---|---|---|
| 50035 | SBER_W_g | 2020 Colmar | chr4_8139889 | 4 | 35 | 27 | 42.3 | 4.46 | 0.05 | 10 | 1.14 | 1.28 | 1.07 | 1.09 | 8.1 | 7.7_16.9 | 8,139,889 |
| 50035 | BER_TA_g | 2018 Pully | chr4_8871452 | 4 | 35 | 18 | 60.8 | 4.97 | 0.02 | 12 | 4.26 | 4.19 | 3.63 | 3.9 | 8.9 | 3.9_23.9 | 8,871,452 |
| 50035 | BERRY_pH | 2018 Pully | chr4_9343236 | 4 | 35.8 | 27 | 46.8 | 6.27 | 0 | 15 | 3.13 | 3.13 | 3.28 | 3.22 | 9.3 | 7.8_18.9 | 9,343,236 |
| 50035 | BERRY_pH | 2022 Pully | chr4_15722823 | 4 | 40.1 | 27 | 59.7 | 7.14 | 0 | 17 | 2.95 | 2.98 | 3.09 | 3.05 | 15.7 | 7.7_23.8 | 15,722,823 |
| 50035 | BER_TA_g | 2020 Pully | chr4_18382632 | 4 | 45.2 | 26 | 61.1 | 5.04 | 0.02 | 12 | 4.1 | 3.91 | 3.44 | 3.81 | 18.4 | 6.1_24.6 | 18,382,632 |
| 50035 | TSS | 2021 Pully | chr5_514286 | 5 | 2.42 | 0 | 12.6 | 5.42 | 0.01 | 13 | 22 | 21.3 | 21.6 | 20.82 | 0.5 | 0_5.1 | 514,286 |
| 50035 | BUD_DATE | 2021 Pully | chr6_944045 | 6 | 2.69 | 0 | 8.3 | 5.04 | 0.01 | 12 | 96.05 | 96.57 | 97.33 | 97.73 | 0.9 | 0.1_6.1 | 944,045 |
| 50035 | BUD_DATE | 2019 Pully | chr6_874344 | 6 | 2.69 | 1.6 | 17.5 | 5.14 | 0.02 | 12 | 99.69 | 100.64 | 101.52 | 101.13 | 0.9 | 2.4_6.4 | 874,344 |
| 50035 | TSS | 2021 Pully | chr6_3953615 | 6 | 12.4 | 0 | 20.4 | 4.59 | 0.03 | 11 | 21.95 | 21.5 | 21.15 | 20.93 | 4 | 3.8_6.4 | 3,953,615 |
| 50035 | SBER_W_g | 2020 Colmar | chr6_4140783 | 6 | 13.2 | 6.7 | 21.8 | 6.51 | 0 | 15 | 1.11 | 1.01 | 1.28 | 1.21 | 4.1 | 3.8_8.3 | 4,140,783 |
| 50035 | TSS | 2022 Pully | chr6_5705399 | 6 | 19.4 | 12 | 36.4 | 4.48 | 0.05 | 11 | 23.11 | 22.72 | 22.4 | 21.95 | 5.7 | 0.1_2.7 | 5,705,399 |
| 50035 | TSS | 2018 Pully | chr6_5705399 | 6 | 19.4 | 11 | 25.3 | 4.93 | 0.02 | 12 | 25.29 | 24.89 | 24.74 | 24.14 | 5.7 | 0.9_5.6 | 5,705,399 |
| 50035 | SBER_W_g | 2020 Pully | chr6_7317871 | 6 | 26.4 | 8.1 | 36.6 | 4.64 | 0.03 | 11 | 1.06 | 1 | 1.21 | 1.11 | 7.3 | 2.7_8.5 | 7,317,871 |
| 50035 | NB_CLUST_PLANT | 2019 Colmar | chr7_1831422 | 7 | 19.4 | 18 | 24.7 | 5.64 | 0 | 14 | 16.73 | 17.91 | 18.19 | 23.44 | 1.8 | 1.5_2.8 | 1,831,422 |
| 50035 | TSS | 2021 Pully | chr7_3404737 | 7 | 26.6 | 20 | 40.9 | 4.7 | 0.03 | 11 | 21.37 | 21.03 | 22.11 | 21.25 | 3.4 | 1.8_15.8 | 3,404,737 |
| 50035 | SBER_W_g | 2020 Pully | chr7_3680084 | 7 | 27.4 | 23 | 51.1 | 5.99 | 0 | 14 | 1.15 | 1.16 | 0.96 | 1.09 | 3.7 | 2.2_21.5 | 3,680,084 |
| 50035 | YIELD_OIV_504 | 2019 Pully | chr7_3656142 | 7 | 28.8 | 15 | 42.2 | 4.44 | 0.05 | 11 | 0.66 | 0.81 | 0.57 | 0.89 | 3.7 | 1.3_18.5 | 3,656,142 |
| 50035 | YIELD_PLANT | 2022 Colmar | chr7_3679950 | 7 | 29 | 6.2 | 42.8 | 4.52 | 0.04 | 11 | 1.77 | 2.08 | 1.53 | 2.3 | 3.7 | 0.4_19.8 | 3,679,950 |
| 50035 | YIELD_OIV_504 | 2022 Colmar | chr7_3679950 | 7 | 29 | 6.2 | 42.8 | 4.52 | 0.04 | 11 | 0.76 | 0.89 | 0.66 | 0.99 | 3.7 | 0.4_19.8 | 3,679,950 |
| 50035 | NB_CLUST_PLANT | 2022 Colmar | chr7_3755307 | 7 | 29.6 | 26 | 38.5 | 4.61 | 0.04 | 11 | 23.5 | 26.72 | 26.17 | 31.58 | 3.8 | 3.4_8 | 3,755,307 |
| 50035 | FLO_50 | 2021 Pully | chr7_7976537 | 7 | 38.5 | 19 | 45.2 | 5.62 | 0 | 13 | 168.8 | 167.94 | 167.6 | 167.38 | 8 | 1.8_20.1 | 7,976,537 |
| 50035 | FLO_50 | 2018 Pully | chr7_12182290 | 7 | 39.5 | 32 | 43 | 7.02 | 0 | 16 | 152.98 | 151.42 | 152.02 | 151 | 12.2 | 4.3_19.8 | 12,182,290 |
| 50035 | FLO_50 | 2019 Pully | chr7_11521337 | 7 | 40.1 | 15 | 51.6 | 4.54 | 0.04 | 11 | 171.85 | 170.98 | 171 | 170.37 | 11.5 | 1.4_21.6 | 11,521,337 |
| 50035 | TSS | 2022 Pully | chr7_15840349 | 7 | 40.9 | 14 | 42.8 | 5.38 | 0.01 | 13 | 22.86 | 22.18 | 23.21 | 22.12 | 15.8 | 1.5_19.8 | 15,840,349 |
| 50035 | YIELD_OIV_504 | 2022 Pully | chr7_18295234 | 7 | 42.2 | 0 | 61.1 | 4.59 | 0.04 | 11 | 0.93 | 1.17 | 0.9 | 1.12 | 18.3 | 0.1_22.1 | 18,295,234 |

| Pop | Trait | Data | Marker | Chr | Pos | ci.low | ci.high | LOD | pval | %var | AC effect | BC effect | AD effect | BD effect | Physical position (Mb) | CI physical position (Mb) | Physical position (bp) |
|---|---|---|---|---|---|---|---|---|---|---|---|---|---|---|---|---|---|
| 50035 | SBER_W_g | 2021 Pully | chr7_21317252 | 7 | 50.8 | 27 | 61.1 | 7.24 | 0 | 17 | 1.36 | 1.56 | 1.21 | 1.46 | 21.3 | 3.4_22.1 | 21,317,252 |
| 50035 | SBER_W_g | 2020 Colmar | chr7_21803624 | 7 | 59.5 | 22 | 61.6 | 5.47 | 0 | 13 | 1.08 | 1.28 | 1.05 | 1.17 | 21.8 | 2.2_22.2 | 21,803,624 |
| 50035 | TSS | 2018 Pully | chr8_17074098 | 8 | 42.2 | 21 | 52.7 | 6.27 | 0 | 15 | 25.53 | 24.11 | 24.76 | 24.75 | 17.1 | 9.1_21.3 | 17,074,098 |
| 50035 | TSS | 2019 Colmar | chr8_18172580 | 8 | 46 | 10 | 53.8 | 4.67 | 0.03 | 12 | 21.67 | 20.56 | 21.43 | 21.5 | 18.2 | 2.3_22.1 | 18,172,580 |
| 50035 | BUD_DATE | 2018 Pully | chr9_846095 | 9 | 4.57 | 0 | 20.4 | 5.22 | 0.01 | 12 | 107 | 107.84 | 106.41 | 107.41 | 0.8 | 1.2_5.7 | 846,095 |
| 50035 | BUD_DATE | 2021 Colmar | chr9_1216351 | 9 | 6.45 | 0 | 31.7 | 4.78 | 0.02 | 12 | 103.86 | 106.96 | 101.8 | 106.19 | 1.2 | 0.5_3.6 | 1,216,351 |
| 50035 | BUD_DATE | 2022 Pully | chr9_2251882 | 9 | 9.95 | 3.2 | 31.7 | 5.15 | 0.01 | 12 | 100.11 | 101.44 | 97.82 | 100.65 | 2.3 | 0.1_13.8 | 2,251,882 |
| 50035 | BUD_DATE | 2022 Colmar | chr9_2251882 | 9 | 9.95 | 6.5 | 31.7 | 5.17 | 0.01 | 12 | 111.82 | 112.68 | 109.8 | 112.18 | 2.3 | 0.1_4.1 | 2,251,882 |
| 50035 | FLO_50 | 2021 Colmar | chr9_2251882 | 9 | 9.95 | 7.3 | 31.7 | 6.22 | 0 | 14 | 166.39 | 167.08 | 165.98 | 167.56 | 2.3 | 0.1_5.3 | 2,251,882 |
| 50035 | TSS | 2019 Colmar | chr9_2251882 | 9 | 9.95 | 4.3 | 31.7 | 6.56 | 0 | 16 | 21.6 | 20.61 | 21.71 | 20.68 | 2.3 | 1.2_5.7 | 2,251,882 |
| 50035 | TSS | 2020 Pully | chr9_2721164 | 9 | 14 | 1.6 | 30.1 | 5.46 | 0.01 | 13 | 23.49 | 22.36 | 23.58 | 22.97 | 2.7 | 1.4_5.5 | 2,721,164 |
| 50035 | TSS | 2021 Pully | chr9_2720967 | 9 | 16.1 | 0.8 | 61.2 | 4.4 | 0.05 | 10 | 21.53 | 21.15 | 21.81 | 20.7 | 2.7 | 0.1_2.7 | 2,720,967 |
| 50035 | YIELD_PLANT | 2019 Colmar | chr9_2720967 | 9 | 16.1 | 0 | 49.6 | 5.37 | 0.01 | 14 | 1.25 | 1.5 | 1.65 | 2.06 | 2.7 | 0.7_23.3 | 2,720,967 |
| 50035 | YIELD_OIV_504 | 2019 Colmar | chr9_2720967 | 9 | 16.1 | 0 | 49.6 | 5.29 | 0.01 | 14 | 0.54 | 0.64 | 0.71 | 0.89 | 2.7 | 0.8_5.7 | 2,720,967 |
| 50035 | TSS | 2018 Pully | chr9_2720967 | 9 | 16.1 | 0 | 27.7 | 5.96 | 0 | 14 | 25.24 | 24.31 | 24.94 | 24.06 | 2.7 | 0.7_18.6 | 2,720,967 |
| 50035 | BUD_DATE | 2020 Colmar | chr9_4057268 | 9 | 27.7 | 0 | 31.2 | 5.56 | 0 | 13 | 98.59 | 99.35 | 97.14 | 99.69 | 4.1 | 0.7_5.5 | 4,057,268 |
| 50035 | TSS | 2022 Pully | chr9_4080476 | 9 | 28.2 | 6.5 | 30.9 | 8.89 | 0 | 20 | 22.81 | 21.53 | 23.11 | 21.87 | 4.1 | 1.2_5.5 | 4,080,476 |
| 50035 | TSS | 2019 Pully | chr9_4898093 | 9 | 29 | 7.3 | 31.7 | 7.4 | 0 | 18 | 23.01 | 22.07 | 23.68 | 22.03 | 4.9 | 1.4_5.5 | 4,898,093 |
| 50035 | FLO_50 | 2022 Colmar | chr9_23262974 | 9 | 61.2 | 3.2 | 61.2 | 5.53 | 0.01 | 13 | 150.05 | 151.49 | 151.06 | 153.5 | 23.3 | 0.8_5.5 | 23,262,974 |
| 50035 | SCLUST_W | 2019 Colmar | chr10_4979003 | 10 | 9.68 | 6.7 | 15.1 | 5.9 | 0 | 15 | 101.68 | 74.72 | 77.41 | 73.79 | 5 | 4.5_6.6 | 4,979,003 |
| 50035 | VER_50 | 2022 Pully | chr11_2587357 | 11 | 4.57 | 0 | 8.6 | 5.25 | 0.01 | 12 | 203.36 | 205.66 | 205.67 | 207.75 | 2.6 | 0_8 | 2,587,357 |
| 50035 | TSS | 2019 Pully | chr11_6641147 | 11 | 23.1 | 0 | 25.5 | 5.01 | 0.02 | 12 | 23.4 | 23.5 | 22.66 | 22.13 | 6.6 | 3.4_17.2 | 6,641,147 |
| 50035 | BUD_DATE | 2022 Colmar | chr11_7960248 | 11 | 25.5 | 8.6 | 41.1 | 4.58 | 0.04 | 11 | 110.32 | 110.31 | 112.35 | 112.47 | 8 | 5.2_12.3 | 7,960,248 |
| 50035 | FLO_50 | 2020 Colmar | chr11_8070627 | 11 | 27.2 | 20 | 34.4 | 6.19 | 0 | 14 | 146.4 | 142.41 | 147.19 | 145.45 | 8.1 | 9.6_18.1 | 8,070,627 |
| 50035 | SBER_W_g | 2019 Pully | chr11_10163835 | 11 | 30.9 | 31 | 44.4 | 7.4 | 0 | 17 | 1.55 | 1.55 | 1.42 | 1.28 | 10.2 | 9.6_19.1 | 10,163,835 |
| 50035 | SBER_W_g | 2021 Pully | chr11_13720591 | 11 | 35.5 | 31 | 48.9 | 5.68 | 0 | 13 | 1.48 | 1.51 | 1.39 | 1.21 | 13.7 | 8.7_18.7 | 13,720,591 |

| Pop | Trait | Data | Marker | Chr | Pos | ci.low | ci.high | LOD | pval | %var | AC effect | BC effect | AD effect | BD effect | Physical position (Mb) | CI physical position (Mb) | Physical position (bp) |
|---|---|---|---|---|---|---|---|---|---|---|---|---|---|---|---|---|---|
| 50035 | SBER_W_g | 2020 Pully | chr11_13720591 | 11 | 35.5 | 29 | 47.9 | 5.85 | 0 | 14 | 1.15 | 1.15 | 1.1 | 0.95 | 13.7 | 9.6_19.1 | 13,720,591 |
| 50035 | SBER_W_g | 2020 Colmar | chr11_13720591 | 11 | 35.5 | 31 | 48.7 | 4.87 | 0.02 | 11 | 1.2 | 1.16 | 1.19 | 0.99 | 13.7 | 6.6_18.4 | 13,720,591 |
| 50035 | BERRY_pH | 2022 Pully | chr11_15375592 | 11 | 37.1 | 23 | 46 | 4.5 | 0.04 | 11 | 2.96 | 3.04 | 2.99 | 3.08 | 15.4 | 0_3.5 | 15,375,592 |
| 50035 | FLO_50 | 2021 Pully | chr12_2047802 | 12 | 5.91 | 0 | 25.2 | 4.89 | 0.02 | 12 | 168.68 | 168.04 | 167.68 | 167.37 | 2 | 0_4.8 | 2,047,802 |
| 50035 | BERRY_pH | 2019 Pully | chr13_7746054 | 13 | 20.2 | 4.8 | 25.5 | 4.97 | 0.02 | 12 | 3.02 | 3.05 | 3.13 | 3.09 | 7.7 | 1.2_10.8 | 7,746,054 |
| 50035 | BERRY_pH | 2019 Colmar | chr13_8817373 | 13 | 23.4 | 0.8 | 34.4 | 5.28 | 0.01 | 13 | 3.06 | 3.05 | 3.12 | 3.09 | 8.8 | 0.5_17.6 | 8,817,373 |
| 50035 | BER_TA_g | 2019 Colmar | chr13_13611183 | 13 | 28.5 | 21 | 31.2 | 6.58 | 0 | 17 | 5.84 | 6.01 | 5.2 | 5.8 | 13.6 | 8_15.9 | 13,611,183 |
| 50035 | VER_50 | 2021 Pully | chr14_15897685 | 14 | 27.4 | 23 | 48.4 | 5.13 | 0.02 | 12 | 221.49 | 226.79 | 222.58 | 222.78 | 15.9 | 10_26.6 | 15,897,685 |
| 50035 | SCLUST_W | 2020 Colmar | chr14_20380301 | 14 | 34.2 | 32 | 50.8 | 6.91 | 0 | 16 | 85.01 | 83.31 | 98.56 | 62.8 | 20.4 | 18_27.3 | 20,380,301 |
| 50035 | NB_CLUST_PLANT | 2022 Colmar | chr14_22605165 | 14 | 39.5 | 36 | 57.8 | 4.76 | 0.03 | 11 | 27.75 | 25.37 | 31.35 | 23.76 | 22.6 | 20.6_30.3 | 22,605,165 |
| 50035 | BER_TA_g | 2021 Pully | chr14_22973706 | 14 | 40.3 | 27 | 48.9 | 5.2 | 0.01 | 12 | 4.3 | 5.11 | 4.23 | 4.37 | 23 | 15.8_26.7 | 22,973,706 |
| 50035 | YIELD_OIV_504 | 2021 Pully | chr14_23123588 | 14 | 41.4 | 32 | 51.6 | 4.53 | 0.05 | 11 | 1.07 | 0.96 | 1.2 | 0.81 | 23.1 | 18_27.4 | 23,123,588 |
| 50035 | NB_CLUST_PLANT | 2019 Colmar | chr14_25518125 | 14 | 47.1 | 44 | 57.5 | 4.92 | 0.02 | 13 | 23.21 | 18.32 | 19.39 | 16.14 | 25.5 | 23.3_30.3 | 25,518,125 |
| 50035 | YIELD_PLANT | 2020 Colmar | chr14_26703281 | 14 | 49.2 | 36 | 51.4 | 8.14 | 0 | 18 | 3.08 | 2.6 | 3.65 | 2.13 | 26.7 | 20.6_27.4 | 26,703,281 |
| 50035 | YIELD_OIV_504 | 2020 Colmar | chr14_26703281 | 14 | 49.2 | 36 | 51.4 | 8.14 | 0 | 18 | 1.32 | 1.12 | 1.57 | 0.91 | 26.7 | 20.6_27.4 | 26,703,281 |
| 50035 | SCLUST_W | 2021 Pully | chr14_27304921 | 14 | 50.8 | 32 | 54.3 | 5.49 | 0.01 | 13 | 186.82 | 165.95 | 217.79 | 137.35 | 27.3 | 18.1_28.8 | 27,304,921 |
| 50035 | TSS | 2022 Colmar | chr14_28522801 | 14 | 53.8 | 9.4 | 58.6 | 4.64 | 0.04 | 11 | 22.33 | 21.7 | 21.28 | 21.44 | 28.5 | 2.9_30.8 | 28,522,801 |
| 50035 | YIELD_OIV_504 | 2021 Pully | chr16_180334 | 16 | 0.27 | 0 | 10.8 | 4.85 | 0.02 | 11 | 0.85 | 1.18 | 0.81 | 0.98 | 0.2 | 0.1_15.2 | 180,334 |
| 50035 | BERRY_pH | 2019 Pully | chr16_1040224 | 16 | 3.76 | 1.6 | 14.2 | 6.24 | 0 | 15 | 2.98 | 3.08 | 3.06 | 3.12 | 1 | 0.5_14.2 | 1,040,224 |
| 50035 | BUD_DATE | 2021 Pully | chr16_1389385 | 16 | 4.84 | 0.8 | 14.5 | 4.59 | 0.04 | 11 | 97.57 | 97.32 | 96.04 | 96.24 | 1.4 | 0.1_18.6 | 1,389,385 |
| 50035 | BUD_DATE | 2018 Pully | chr16_1953673 | 16 | 5.65 | 0 | 17.5 | 4.69 | 0.02 | 11 | 107.46 | 107.38 | 106.65 | 106.41 | 2 | 11.9_17.9 | 1,953,673 |
| 50035 | SCLUST_W | 2021 Pully | chr16_3547462 | 16 | 6.99 | 0 | 18 | 5.02 | 0.02 | 12 | 158.93 | 211.59 | 160.25 | 147.78 | 3.5 | 16.2_16.7 | 3,547,462 |
| 50035 | YIELD_PLANT | 2020 Colmar | chr16_3006969 | 16 | 7.26 | 3.8 | 11.6 | 5.12 | 0.02 | 12 | 2.36 | 3.4 | 2.84 | 2.46 | 3 | 13.8_16.8 | 3,006,969 |
| 50035 | YIELD_OIV_504 | 2020 Colmar | chr16_3006969 | 16 | 7.26 | 3.8 | 11.6 | 5.12 | 0.01 | 12 | 1.01 | 1.46 | 1.22 | 1.06 | 3 | 0.3_17.3 | 3,006,969 |
| 50035 | BER_TA_g | 2019 Pully | chr16_13791365 | 16 | 10.2 | 1.6 | 10.2 | 6.11 | 0 | 15 | 5.07 | 4.32 | 4.23 | 4.27 | 13.8 | 15_16.8 | 13,791,365 |

| Pop | Trait | Data | Marker | Chr | Pos | ci.low | ci.high | LOD | pval | %var | AC effect | BC effect | AD effect | BD effect | Physical position (Mb) | CI physical position (Mb) | Physical position (bp) |
|---|---|---|---|---|---|---|---|---|---|---|---|---|---|---|---|---|---|
| 50035 | HARVEST_DATE | 2019 Colmar | chr16_15309542 | 16 | 11.6 | 5.6 | 15.1 | 7.74 | 0 | 19 | 276.24 | 267.8 | 265.75 | 269.43 | 15.3 | 14.4_16.7 | 15,309,542 |
| 50035 | SCLUST_W | 2020 Colmar | chr16_15542985 | 16 | 12.1 | 0 | 25.3 | 5.83 | 0 | 13 | 83.82 | 94.69 | 79.27 | 66.52 | 15.5 | 16.2_16.6 | 15,542,985 |
| 50035 | BER_TA_g | 2019 Colmar | chr16_15761241 | 16 | 12.4 | 9.4 | 21 | 4.44 | 0.05 | 11 | 6.13 | 5.43 | 5.52 | 5.78 | 15.8 | 1_16.8 | 15,761,241 |
| 50035 | BER_TA_g | 2021 Pully | chr16_16202746 | 16 | 13.4 | 13 | 15.3 | 19.1 | 0 | 19 | 5.92 | 4.16 | 4.21 | 4.34 | 16.2 | 2.2_16.7 | 16,202,746 |
| 50035 | TSS | 2020 Colmar | chr16_16575285 | 16 | 13.4 | 10 | 15.1 | 8.91 | 0 | 20 | 23.22 | 21.81 | 21.66 | 21.88 | 16.6 | 3_17.4 | 16,575,285 |
| 50035 | BUD_DATE | 2019 Pully | chr16_16575285 | 16 | 13.4 | 1.3 | 16.1 | 6.64 | 0 | 15 | 101.82 | 101.11 | 100.1 | 99.73 | 16.6 | 2_16.7 | 16,575,285 |
| 50035 | VER_50 | 2019 Colmar | chr16_16202746 | 16 | 13.4 | 11 | 15.1 | 14.7 | 0 | 33 | 221.27 | 215.67 | 215.14 | 216.69 | 16.2 | 16.2_16.8 | 16,202,746 |
| 50035 | BERRY_pH | 2021 Pully | chr16_16601202 | 16 | 14.3 | 11 | 15.3 | 13.8 | 0 | 29 | 2.9 | 3.17 | 3.13 | 3.18 | 16.6 | 15_16.7 | 16,601,202 |
| 50035 | VER_50 | 2021 Pully | chr16_16647186 | 16 | 14.3 | 13 | 14.2 | 42.7 | 0 | 66 | 234.31 | 221.7 | 220.17 | 221.61 | 16.6 | 16.2_16.8 | 16,647,186 |
| 50035 | VER_50 | 2022 Pully | chr16_16601209 | 16 | 14.5 | 5.6 | 15.3 | 7.29 | 0 | 17 | 209.47 | 204.98 | 204.62 | 204.41 | 16.6 | 3_17.6 | 16,601,209 |
| 50035 | HARVEST_DATE | 2022 Colmar | chr16_16792629 | 16 | 14.5 | 7 | 18 | 6.45 | 0 | 15 | 268.92 | 260 | 257.59 | 256.11 | 16.8 | 0.3_18.4 | 16,792,629 |
| 50035 | VER_50 | 2022 Colmar | chr16_16601209 | 16 | 14.5 | 6.2 | 15.3 | 7.9 | 0 | 18 | 215.92 | 211.82 | 211.93 | 211.25 | 16.6 | 17.9_18.6 | 16,601,209 |
| 50035 | VER_50 | 2020 Pully | chr16_16601209 | 16 | 14.5 | 13 | 15.1 | 24 | 0 | 45 | 217.77 | 208.76 | 208.07 | 207.73 | 16.6 | 0.5_16.6 | 16,601,209 |
| 50035 | HARVEST_DATE | 2020 Colmar | chr16_16601209 | 16 | 14.5 | 11 | 15.3 | 16.1 | 0 | 33 | 268.89 | 257.37 | 254.55 | 255.41 | 16.6 | 0.1_18.6 | 16,601,209 |
| 50035 | VER_50 | 2019 Pully | chr16_16601209 | 16 | 14.5 | 13 | 15.1 | 22.4 | 0 | 43 | 225.88 | 219.54 | 218.52 | 219.34 | 16.6 | 0.2_16.8 | 16,601,209 |
| 50035 | VER_50 | 2020 Colmar | chr16_16751263 | 16 | 15.1 | 12 | 15.1 | 22 | 0 | 42 | 214.39 | 206.06 | 204.9 | 205.28 | 16.8 | 0.1_17.4 | 16,751,263 |
| 50035 | VER_50 | 2018 Pully | chr16_16714028 | 16 | 15.1 | 13 | 16.4 | 13.3 | 0 | 29 | 211.21 | 206.29 | 204.83 | 204.84 | 16.7 | 0.1_17.4 | 16,714,028 |
| 50035 | MORPHO_OIV_204 | 2020 Colmar | chr16_17154341 | 16 | 16.1 | 7 | 19.4 | 5.85 | 0 | 14 | 1.86 | 2.97 | 2.7 | 3.09 | 17.2 | 1_19.4 | 17,154,341 |
| 50035 | BUD_DATE | 2020 Pully | chr16_18282234 | 16 | 22.6 | 1.3 | 24.5 | 7.67 | 0 | 17 | 96.77 | 96.09 | 95.65 | 94.13 | 18.3 | 1_15.3 | 18,282,234 |
| 50035 | SBER_W_g | 2022 Pully | chr16_18246770 | 16 | 23.1 | 21 | 25.3 | 4.87 | 0.02 | 11 | 0.97 | 1.01 | 1.08 | 0.89 | 18.2 | 1_15.3 | 18,246,770 |
| 50035 | BERRY_pH | 2018 Pully | chr18_2085789 | 18 | 9.68 | 5.6 | 17.2 | 5.62 | 0 | 13 | 3.15 | 3.19 | 3.15 | 3.29 | 2.1 | 3.3_7.4 | 2,085,789 |
| 50035 | VER_50 | 2020 Pully | chr18_4864745 | 18 | 17.5 | 13 | 25 | 5.17 | 0.01 | 12 | 212.09 | 208.15 | 209.19 | 207.13 | 4.9 | 2.5_12.8 | 4,864,745 |
| 50035 | VER_50 | 2019 Pully | chr18_4864745 | 18 | 17.5 | 11 | 54.1 | 4.99 | 0.02 | 12 | 222.3 | 219.31 | 220.43 | 219 | 4.9 | 8.7_10.8 | 4,864,745 |
| 50035 | MORPHO_OIV_204 | 2022 Colmar | chr18_8741174 | 18 | 37.7 | 38 | 43.8 | 6.34 | 0 | 10 | 2.24 | 2.37 | 2.87 | 2.69 | 8.7 | 8.7_11.5 | 8,741,174 |

| Pop | Trait | Data | Marker | Chr | Pos | ci.low | ci.high | LOD | pval | %var | AC effect | BC effect | AD effect | BD effect | Physical position (Mb) | CI physical position (Mb) | Physical position (bp) |
|---|---|---|---|---|---|---|---|---|---|---|---|---|---|---|---|---|---|
| 50035 | MORPHO_OIV_204 | 2020 Colmar | chr18_10359111 | 18 | 42.5 | 32 | 75.3 | 5.7 | 0.01 | 13 | 2.16 | 2.62 | 2.89 | 3.45 | 10.4 | 11.2_12.9 | 10,359,111 |
| 50035 | BER_TA_g | 2019 Pully | chr18_11865605 | 18 | 49.5 | 47 | 53 | 4.7 | 0.02 | 12 | 4.21 | 4.1 | 4.61 | 4.83 | 11.9 | 10.1_14.2 | 11,865,605 |
| 50035 | BUD_DATE | 2019 Pully | chr18_12844141 | 18 | 54.1 | 42 | 57 | 5.25 | 0.01 | 12 | 100.04 | 100.29 | 101.61 | 100.08 | 12.8 | 11.9_27.1 | 12,844,141 |
| 50035 | TSS | 2022 Pully | chr18_13479676 | 18 | 55.4 | 50 | 64.5 | 8.49 | 0 | 19 | 22.27 | 22.7 | 22.14 | 23.63 | 13.5 | 12.8_32 | 13,479,676 |
| 50035 | SBER_W_g | 2022 Pully | chr18_19118472 | 18 | 60 | 55 | 72.1 | 4.86 | 0.02 | 11 | 1.06 | 0.99 | 0.99 | 0.86 | 19.1 | 12.8_31.9 | 19,118,472 |
| 50035 | SBER_W_g | 2020 Colmar | chr18_19118457 | 18 | 60 | 54 | 72.3 | 5.24 | 0.01 | 12 | 1.24 | 1.14 | 1.18 | 0.97 | 19.1 | 12.8_27.1 | 19,118,457 |
| 50035 | TSS | 2021 Pully | chr18_26625049 | 18 | 63.2 | 53 | 71 | 6.23 | 0 | 14 | 20.96 | 21.61 | 21.2 | 22.28 | 26.6 | 11.5_27.1 | 26,625,049 |
| 50035 | TSS | 2019 Colmar | chr18_25690168 | 18 | 63.2 | 48 | 64.5 | 6.72 | 0 | 17 | 21.1 | 21.38 | 20.95 | 22.28 | 25.7 | 14.3_32 | 25,690,168 |
| 50035 | TSS | 2018 Pully | chr18_26979605 | 18 | 63.7 | 59 | 72.1 | 8.01 | 0 | 19 | 24.58 | 25.09 | 24.34 | 25.72 | 27 | 14.3_27.1 | 26,979,605 |
| 50035 | TSS | 2019 Pully | chr18_27084024 | 18 | 64.3 | 59 | 64.5 | 13.4 | 0 | 29 | 22.73 | 23.55 | 22.11 | 24.36 | 27.1 | 12.8_31.9 | 27,084,024 |
| 50035 | TSS | 2020 Pully | chr18_27084309 | 18 | 64.6 | 55 | 72.3 | 8.32 | 0 | 19 | 23.06 | 23.65 | 22.71 | 23.19 | 27.1 | 27_34.3 | 27,084,309 |
| 50035 | SCLUST_W | 2021 Pully | chr18_30766656 | 18 | 69.1 | 66 | 77.5 | 5.13 | 0.02 | 12 | 208.59 | 175.96 | 140.3 | 190.01 | 30.8 | 1.2_4.6 | 30,766,656 |
| 50035 | SBER_W_g | 2019 Pully | chr19_6840320 | 19 | 34.1 | 26 | 57.8 | 4.92 | 0.02 | 12 | 1.31 | 1.54 | 1.41 | 1.54 | 6.8 | 5.3_24.3 | 6,840,320 |
| 50035 | BUD_DATE | 2020 Colmar | chr19_7705621 | 19 | 37.4 | 30 | 51.1 | 4.45 | 0.03 | 11 | 96.95 | 98.19 | 99.15 | 98.89 | 7.7 | 6.2_17.2 | 7,705,621 |

**Supplementary Table 10.** Trait-level summary of the population-adjusted multiple-population QTL scan (772 progeny, 12 population labels, 8,054 composite-map markers; Li-Ji P threshold 3.442E-5). Abbreviations: QTL, quantitative trait locus; cM, centimorgan; GC, genomic control.

| trait | N tested | N significant LiJi | lead marker | lead chromosome | lead position cM | minimum P value | lambda GC | N peak support intervals | N distinguishable peak patterns |
|---|---|---|---|---|---|---|---|---|---|
| BUD_DATE | 8,054 | 4 | chr9_19559720 | 9 | 164.431 | 1.14E-17 | 1.111 | 4 | 3 |
| FLO_50 | 8,050 | 10 | chr1_17756488 | 1 | 95.328 | 1.162E-6 | 1.084 | 5 | 5 |
| VER_50 | 8,050 | 6 | chr5_20351051 | 5 | 81.747 | 1.193E-15 | 1.155 | 4 | 4 |
| SBER_W_g | 8,050 | 12 | chr17_6561319 | 17 | 55.83 | 3.567E-7 | 1.428 | 5 | 5 |
| TSS | 8,050 | 8 | chr6_7263263 | 6 | 73.831 | 6.671E-12 | 1.279 | 8 | 7 |
| BERRY_pH | 8,050 | 1 | chr16_14689931 | 16 | 77.639 | 1.301E-5 | 1.132 | 1 | 1 |
| BER_TA_g | 8,050 | 1 | chr5_20351051 | 5 | 81.747 | 4.49E-6 | 1.373 | 1 | 1 |
| HARVEST_DATE | 8,050 | 6 | chr5_20351051 | 5 | 81.747 | 1.604E-13 | 1.213 | 4 | 4 |
| MORPHO_OIV_204 | 8,050 | 13 | chr14_29206455 | 14 | 122.677 | 1.091E-8 | 1.075 | 5 | 5 |
| NB_CLUST_PLANT | 8,050 | 4 | chr17_1289030 | 17 | 6.17 | 4.201E-6 | 1.307 | 3 | 3 |
| YIELD_PLANT | 8,050 | 2 | chr14_22600042 | 14 | 83.091 | 1.089E-5 | 1.102 | 2 | 2 |
| SCLUST_W | 7,686 | 2 | chr14_22600042 | 14 | 83.091 | 2.579E-5 | 1.191 | 2 | 2 |
| YIELD_OIV_504 | 8,050 | 5 | chr14_22600042 | 14 | 83.091 | 1.698E-6 | 1.12 | 1 | 1 |

**Supplementary Table 11A.** Marker-defined physical support intervals, dosage-pattern diagnostics and overlap with single-family QTL and additive GWAS. Positional spans are heuristic 1.5-log10(P)-drop supports, not formal confidence intervals. Peak locations and support spans. Abbreviations: QTL, quantitative trait locus; GWAS, genome-wide association study; LD, linkage disequilibrium; cM, centimorgan; bp, base pair.

| trait | physical chr | lead marker | lead position bp | lead position cM | lead P value | support start bp | support end bp | support start cM | support end cM |
|---|---|---|---|---|---|---|---|---|---|
| BUD_DATE | 7 | chr7_4414937 | 4,414,937 | 25.262 | 2.881E-5 | 4,414,937 | 6,333,567 | 25.262 | 32.684 |
| BUD_DATE | 9 | chr9_19559720 | 19,559,720 | 164.431 | 1.14E-17 | 19,559,720 | 19,559,720 | 164.431 | 164.431 |
| BUD_DATE | 11 | chr11_12139361 | 12,139,361 | 92.633 | 1.14E-17 | 12,139,361 | 12,139,361 | 92.633 | 92.633 |
| BUD_DATE | 16 | chr16_15360686 | 15,360,686 | 92.756 | 1.063E-5 | 15,360,686 | 15,360,686 | 92.756 | 92.756 |
| FLO_50 | 1 | chr1_17756488 | 17,756,488 | 95.328 | 1.162E-6 | 17,756,488 | 17,756,488 | 95.328 | 95.328 |
| FLO_50 | 2 | chr2_18622604 | 18,622,604 | 189.551 | 2.235E-5 | 18,622,604 | 18,622,604 | 189.551 | 189.551 |
| FLO_50 | 7 | chr7_20342111 | 20,342,111 | 74.052 | 2.883E-6 | 20,342,111 | 20,586,233 | 74.052 | 76.005 |
| FLO_50 | 17 | chr17_5562930 | 5,562,930 | 45.596 | 8.97E-6 | 5,268,425 | 6,431,844 | 41.038 | 55.504 |
| FLO_50 | 17 | chr17_6431844 | 6,431,844 | 55.504 | 9.303E-6 | 5,562,930 | 6,841,649 | 45.596 | 57.197 |
| VER_50 | 5 | chr5_20351051 | 20,351,051 | 81.747 | 1.193E-15 | 20,351,051 | 20,351,051 | 81.747 | 81.747 |
| VER_50 | 16 | chr16_14689931 | 14,689,931 | 77.639 | 3.675E-14 | 14,689,860 | 14,689,931 | 77.639 | 77.77 |
| VER_50 | 16 | chr16_7122451 | 7,122,451 | 50.355 | 3.423E-8 | 7,122,451 | 7,158,029 | 50.355 | 50.55 |
| VER_50 | 17 | chr17_6431844 | 6,431,844 | 55.504 | 1.446E-5 | 6,211,854 | 6,561,310 | 53.746 | 55.83 |
| SBER_W_g | 2 | chr2_5621690 | 5,621,690 | 118.106 | 2.645E-5 | 5,278,904 | 5,643,634 | 113.092 | 118.431 |
| SBER_W_g | 2 | chr2_5278904 | 5,278,904 | 113.092 | 2.645E-5 | 5,278,904 | 5,643,634 | 113.092 | 118.431 |
| SBER_W_g | 15 | chr15_16599366 | 16,599,366 | 65.563 | 1.299E-5 | 16,177,758 | 16,856,009 | 62.699 | 72.857 |
| SBER_W_g | 15 | chr15_16856009 | 16,856,009 | 72.857 | 1.686E-5 | 16,351,809 | 16,856,009 | 63.675 | 72.857 |
| SBER_W_g | 17 | chr17_6561319 | 6,561,319 | 55.83 | 3.567E-7 | 6,431,844 | 6,561,319 | 55.504 | 55.83 |
| TSS | 6 | chr6_7263263 | 7,263,263 | 73.831 | 6.671E-12 | 7,263,263 | 7,263,263 | 73.831 | 73.831 |
| TSS | 8 | chr8_11465636 | 11,465,636 | 35.747 | 6.671E-12 | 11,465,636 | 11,465,636 | 35.747 | 35.747 |
| TSS | 9 | chr9_2155288 | 2,155,288 | 20.638 | 6.671E-12 | 2,155,288 | 2,155,288 | 20.638 | 20.638 |
| TSS | 12 | chr12_20333142 | 20,333,142 | 139.818 | 6.671E-12 | 20,333,142 | 20,333,142 | 139.818 | 139.818 |
| TSS | 12 | chr12_7746624 | 7,746,624 | 86.258 | 4.017E-7 | 7,185,776 | 7,746,624 | 79.227 | 86.258 |
| TSS | 12 | chr12_7185776 | 7,185,776 | 79.227 | 6.665E-7 | 7,185,776 | 7,746,624 | 79.227 | 86.258 |
| TSS | 13 | chr13_2515385 | 2,515,385 | 16.363 | 6.671E-12 | 2,515,385 | 2,515,385 | 16.363 | 16.363 |

| trait | physical chr | lead marker | lead position bp | lead position cM | lead P value | support start bp | support end bp | support start cM | support end cM |
|---|---|---|---|---|---|---|---|---|---|
| TSS | 14 | chr14_30192660 | 30,192,660 | 137.733 | 6.671E-12 | 30,192,660 | 30,192,660 | 137.733 | 137.733 |
| BERRY_pH | 16 | chr16_14689931 | 14,689,931 | 77.639 | 1.301E-5 | 14,689,860 | 14,689,931 | 77.639 | 77.77 |
| BER_TA_g | 5 | chr5_20351051 | 20,351,051 | 81.747 | 4.49E-6 | 20,351,051 | 20,351,051 | 81.747 | 81.747 |
| HARVEST_DATE | 5 | chr5_20351051 | 20,351,051 | 81.747 | 1.604E-13 | 20,351,051 | 20,351,051 | 81.747 | 81.747 |
| HARVEST_DATE | 16 | chr16_14689931 | 14,689,931 | 77.639 | 1.051E-11 | 14,689,860 | 14,689,931 | 77.639 | 77.77 |
| HARVEST_DATE | 16 | chr16_7158029 | 7,158,029 | 50.55 | 7.702E-8 | 7,122,451 | 7,158,029 | 50.355 | 50.55 |
| HARVEST_DATE | 17 | chr17_6431844 | 6,431,844 | 55.504 | 2.133E-5 | 5,599,964 | 6,646,471 | 45.921 | 56.285 |
| MORPHO_OIV_204 | 1 | chr1_5742125 | 5,742,125 | 36.201 | 8.466E-6 | 4,261,108 | 6,450,982 | 27.802 | 45.321 |
| MORPHO_OIV_204 | 13 | chr13_3543269 | 3,543,269 | 20.139 | 2.75E-5 | 3,543,269 | 3,543,269 | 20.139 | 20.139 |
| MORPHO_OIV_204 | 14 | chr14_29206455 | 29,206,455 | 122.677 | 1.091E-8 | 28,603,413 | 29,206,455 | 113.301 | 122.677 |
| MORPHO_OIV_204 | 14 | chr14_28603413 | 28,603,413 | 113.301 | 3.22E-7 | 27,915,149 | 29,206,455 | 108.743 | 122.677 |
| MORPHO_OIV_204 | 14 | chr14_26846468 | 26,846,468 | 101.125 | 1.368E-5 | 26,846,468 | 27,943,166 | 101.125 | 109.329 |
| NB_CLUST_PLANT | 5 | chr5_1219467 | 1,219,467 | 7.323 | 9.516E-6 | 602,088 | 3,287,131 | 4.951 | 16.571 |
| NB_CLUST_PLANT | 17 | chr17_1289030 | 1,289,030 | 6.17 | 4.201E-6 | 1,289,030 | 1,741,806 | 6.17 | 12.649 |
| NB_CLUST_PLANT | 17 | chr17_2154133 | 2,154,133 | 18.51 | 5.864E-6 | 1,612,434 | 2,154,133 | 12.254 | 18.51 |
| YIELD_PLANT | 14 | chr14_22600042 | 22,600,042 | 83.091 | 1.089E-5 | 22,456,692 | 25,504,875 | 82.374 | 92.661 |
| YIELD_PLANT | 14 | chr14_26074715 | 26,074,715 | 95.786 | 1.577E-5 | 24,642,352 | 26,812,126 | 87.583 | 101.06 |
| SCLUST_W | 14 | chr14_22600042 | 22,600,042 | 83.091 | 2.579E-5 | 22,209,049 | 25,472,132 | 79.966 | 92.661 |
| SCLUST_W | 14 | chr14_26112922 | 26,112,922 | 96.307 | 3.334E-5 | 24,562,322 | 26,113,398 | 87.583 | 96.307 |
| YIELD_OIV_504 | 14 | chr14_22600042 | 22,600,042 | 83.091 | 1.698E-6 | 22,456,692 | 24,958,639 | 82.374 | 88.69 |

**Supplementary Table 11B (continued).** Dosage-pattern diagnostics and cross-framework overlap. Abbreviations: QTL, quantitative trait locus; GWAS, genome-wide association study. Other abbreviations are defined in Supplementary Table 11A.

| trait | physical chr | lead marker | genotype pattern id | cross chromosome duplicate pattern | single family QTL overlap | overlapping families | adjusted GWAS overlap | adjusted GWAS leads | adjusted GWAS models |
|---|---|---|---|---|---|---|---|---|---|
| BUD_DATE | 7 | chr7_4414937 | GP011 | FALSE | TRUE | 50025 | FALSE | | |
| BUD_DATE | 9 | chr9_19559720 | GP018 | TRUE | FALSE | | FALSE | | |
| BUD_DATE | 11 | chr11_12139361 | GP018 | TRUE | FALSE | | FALSE | | |
| BUD_DATE | 16 | chr16_15360686 | GP047 | FALSE | TRUE | 50035 | FALSE | | |
| FLO_50 | 1 | chr1_17756488 | GP003 | FALSE | FALSE | | FALSE | | |
| FLO_50 | 2 | chr2_18622604 | GP006 | FALSE | FALSE | | FALSE | | |
| FLO_50 | 7 | chr7_20342111 | GP012 | FALSE | TRUE | 42050;50035 | FALSE | | |
| FLO_50 | 17 | chr17_5562930 | GP050 | FALSE | TRUE | 50015 | FALSE | | |
| FLO_50 | 17 | chr17_6431844 | GP052 | FALSE | TRUE | 50015 | TRUE | chr17_6535707 | MM4LMM_K_LD1-LD5 |
| VER_50 | 5 | chr5_20351051 | GP009 | FALSE | FALSE | | TRUE | chr5_20351051 | MM4LMM_K_LD1-LD5 |
| VER_50 | 16 | chr16_14689931 | GP046 | FALSE | TRUE | 50025;50035 | TRUE | chr16_14689931;chr16_14689860 | MM4LMM_K_LD1-LD5 |
| VER_50 | 16 | chr16_7122451 | GP043 | FALSE | TRUE | 50035 | TRUE | chr16_7122451;chr16_7158029;chr16_7135100;chr16_7154039 | MM4LMM_K_LD1-LD5 |
| VER_50 | 17 | chr17_6431844 | GP052 | FALSE | FALSE | | TRUE | chr17_6512440 | BLINK_LD1-LD5 |
| SBER_W_g | 2 | chr2_5621690 | GP005 | FALSE | TRUE | 50015 | FALSE | | |
| SBER_W_g | 2 | chr2_5278904 | GP004 | FALSE | TRUE | 50015 | FALSE | | |
| SBER_W_g | 15 | chr15_16599366 | GP040 | FALSE | FALSE | | FALSE | | |
| SBER_W_g | 15 | chr15_16856009 | GP042 | FALSE | FALSE | | FALSE | | |
| SBER_W_g | 17 | chr17_6561319 | GP055 | FALSE | FALSE | | FALSE | | |
| TSS | 6 | chr6_7263263 | GP010 | TRUE | TRUE | 50035 | FALSE | | |
| TSS | 8 | chr8_11465636 | GP016 | FALSE | FALSE | | FALSE | | |

| trait | physical chr | lead marker | genotype pattern id | cross chromosome duplicate pattern | single family QTL overlap | overlapping families | adjusted GWAS overlap | adjusted GWAS leads | adjusted GWAS models |
|---|---|---|---|---|---|---|---|---|---|
| TSS | 9 | chr9_2155288 | GP017 | FALSE | TRUE | 50035 | FALSE | | |
| TSS | 12 | chr12_20333142 | GP021 | FALSE | FALSE | | FALSE | | |
| TSS | 12 | chr12_7746624 | GP020 | FALSE | FALSE | | FALSE | | |
| TSS | 12 | chr12_7185776 | GP019 | FALSE | FALSE | | FALSE | | |
| TSS | 13 | chr13_2515385 | GP010 | TRUE | FALSE | | FALSE | | |
| TSS | 14 | chr14_30192660 | GP039 | FALSE | FALSE | | FALSE | | |
| BERRY_pH | 16 | chr16_14689931 | GP046 | FALSE | FALSE | | FALSE | | |
| BER_TA_g | 5 | chr5_20351051 | GP009 | FALSE | FALSE | | FALSE | | |
| HARVEST_DATE | 5 | chr5_20351051 | GP009 | FALSE | FALSE | | TRUE | chr5_20351051 | MM4LMM_K_LD1-LD5 |
| HARVEST_DATE | 16 | chr16_14689931 | GP046 | FALSE | TRUE | 50025 | TRUE | chr16_14689860;chr16_14689931 | MM4LMM_K_LD1-LD5 |
| HARVEST_DATE | 16 | chr16_7158029 | GP044 | FALSE | FALSE | | TRUE | chr16_7122451;chr16_7158029;chr16_7135100;chr16_7154039 | MM4LMM_K_LD1-LD5 |
| HARVEST_DATE | 17 | chr17_6431844 | GP052 | FALSE | FALSE | | TRUE | chr17_6401811 | BLINK_LD1-LD5 |
| MORPHO_OIV_204 | 1 | chr1_5742125 | GP002 | FALSE | TRUE | 50035 | TRUE | chr1_5670971;chr1_5000181;chr1_4926826;chr1_5551443;chr1_5467931;chr1_5009157 | BLINK_LD1-LD5;MLMM_K_LD1-LD5;MM4LMM_K_LD1-LD5 |
| MORPHO_OIV_204 | 13 | chr13_3543269 | GP022 | FALSE | FALSE | | FALSE | | |
| MORPHO_OIV_204 | 14 | chr14_29206455 | GP037 | FALSE | TRUE | 50013 | FALSE | | |
| MORPHO_OIV_204 | 14 | chr14_28603413 | GP036 | FALSE | TRUE | 50013 | FALSE | | |
| MORPHO_OIV_204 | 14 | chr14_26846468 | GP030 | FALSE | TRUE | 50013 | FALSE | | |
| NB_CLUST_PLANT | 5 | chr5_1219467 | GP007 | FALSE | FALSE | | FALSE | | |
| NB_CLUST_PLANT | 17 | chr17_1289030 | GP048 | FALSE | TRUE | 50015 | FALSE | | |
| NB_CLUST_PLANT | 17 | chr17_2154133 | GP049 | FALSE | TRUE | 50015 | FALSE | | |

| trait | physical chr | lead marker | genotype pattern id | cross chromosome duplicate pattern | single family QTL overlap | overlapping families | adjusted GWAS overlap | adjusted GWAS leads | adjusted GWAS models |
|---|---|---|---|---|---|---|---|---|---|
| YIELD_PLANT | 14 | chr14_22600042 | GP025 | FALSE | TRUE | 50013;50035 | FALSE | | |
| YIELD_PLANT | 14 | chr14_26074715 | GP028 | FALSE | TRUE | 50013;50035 | FALSE | | |
| SCLUST_W | 14 | chr14_22600042 | GP025 | FALSE | TRUE | 50013 | FALSE | | |
| SCLUST_W | 14 | chr14_26112922 | GP029 | FALSE | TRUE | 50013 | FALSE | | |
| YIELD_OIV_504 | 14 | chr14_22600042 | GP025 | FALSE | TRUE | 50013;50035 | FALSE | | |

**Supplementary Table 12A.** Standardized dosage effects and approximate 95% Wald intervals for population-adjusted multiple-population QTL markers significant at the Li-Ji threshold. Significant-marker locations and P values. Abbreviations: QTL, quantitative trait locus; cM, centimorgan.

| trait | marker | chromosome | position cM | P value |
|---|---|---|---|---|
| BUD_DATE | chr7_4414937 | 7 | 25.262 | 2.881E-5 |
| BUD_DATE | chr9_19559720 | 9 | 164.431 | 1.14E-17 |
| BUD_DATE | chr11_12139361 | 11 | 92.633 | 1.14E-17 |
| BUD_DATE | chr16_15360686 | 16 | 92.756 | 1.063E-5 |
| FLO_50 | chr1_17756488 | 1 | 95.328 | 1.162E-6 |
| FLO_50 | chr2_18622604 | 2 | 189.551 | 2.235E-5 |
| FLO_50 | chr7_20342111 | 7 | 74.052 | 2.883E-6 |
| FLO_50 | chr7_20396421 | 7 | 74.247 | 8.686E-6 |
| FLO_50 | chr7_20556531 | 7 | 74.573 | 1.834E-5 |
| FLO_50 | chr7_20537728 | 7 | 74.703 | 9.073E-6 |
| FLO_50 | chr7_20586233 | 7 | 76.005 | 1.834E-5 |
| FLO_50 | chr17_5562930 | 17 | 45.596 | 8.97E-6 |
| FLO_50 | chr17_5599964 | 17 | 45.921 | 2.447E-5 |
| FLO_50 | chr17_6431844 | 17 | 55.504 | 9.303E-6 |
| VER_50 | chr5_20351051 | 5 | 81.747 | 1.193E-15 |
| VER_50 | chr16_7122451 | 16 | 50.355 | 3.423E-8 |
| VER_50 | chr16_7158029 | 16 | 50.55 | 1.366E-7 |
| VER_50 | chr16_14689931 | 16 | 77.639 | 3.675E-14 |
| VER_50 | chr16_14689860 | 16 | 77.77 | 3.965E-14 |
| VER_50 | chr17_6431844 | 17 | 55.504 | 1.446E-5 |
| SBER_W_g | chr2_5278904 | 2 | 113.092 | 2.645E-5 |
| SBER_W_g | chr2_5621690 | 2 | 118.106 | 2.645E-5 |
| SBER_W_g | chr15_16599366 | 15 | 65.563 | 1.299E-5 |

| trait | marker | chromosome | position cM | P value |
|---|---|---|---|---|
| SBER_W_g | chr15_16829505 | 15 | 72.857 | 2.758E-5 |
| SBER_W_g | chr15_16856009 | 15 | 72.857 | 1.686E-5 |
| SBER_W_g | chr17_6431844 | 17 | 55.504 | 4.737E-6 |
| SBER_W_g | chr17_6490445 | 17 | 55.765 | 2.423E-6 |
| SBER_W_g | chr17_6561310 | 17 | 55.83 | 9.161E-7 |
| SBER_W_g | chr17_6561319 | 17 | 55.83 | 3.567E-7 |
| SBER_W_g | chr17_6830639 | 17 | 56.741 | 2.114E-5 |
| SBER_W_g | chr17_6830638 | 17 | 56.741 | 1.55E-5 |
| SBER_W_g | chr17_7025564 | 17 | 58.043 | 2.893E-5 |
| TSS | chr6_7263263 | 6 | 73.831 | 6.671E-12 |
| TSS | chr8_11465636 | 8 | 35.747 | 6.671E-12 |
| TSS | chr9_2155288 | 9 | 20.638 | 6.671E-12 |
| TSS | chr12_7185776 | 12 | 79.227 | 6.665E-7 |
| TSS | chr12_7746624 | 12 | 86.258 | 4.017E-7 |
| TSS | chr12_20333142 | 12 | 139.818 | 6.671E-12 |
| TSS | chr13_2515385 | 13 | 16.363 | 6.671E-12 |
| TSS | chr14_30192660 | 14 | 137.733 | 6.671E-12 |
| BERRY_pH | chr16_14689931 | 16 | 77.639 | 1.301E-5 |
| BER_TA_g | chr5_20351051 | 5 | 81.747 | 4.49E-6 |
| HARVEST_DATE | chr5_20351051 | 5 | 81.747 | 1.604E-13 |
| HARVEST_DATE | chr16_7122451 | 16 | 50.355 | 2.003E-7 |
| HARVEST_DATE | chr16_7158029 | 16 | 50.55 | 7.702E-8 |
| HARVEST_DATE | chr16_14689931 | 16 | 77.639 | 1.051E-11 |
| HARVEST_DATE | chr16_14689860 | 16 | 77.77 | 1.582E-11 |
| HARVEST_DATE | chr17_6431844 | 17 | 55.504 | 2.133E-5 |

| trait | marker | chromosome | position cM | P value |
|---|---|---|---|---|
| MORPHO_OIV_204 | chr1_4926883 | 1 | 32.881 | 1.301E-5 |
| MORPHO_OIV_204 | chr1_5742125 | 1 | 36.201 | 8.466E-6 |
| MORPHO_OIV_204 | chr13_3543269 | 13 | 20.139 | 2.75E-5 |
| MORPHO_OIV_204 | chr14_26846468 | 14 | 101.125 | 1.368E-5 |
| MORPHO_OIV_204 | chr14_26942921 | 14 | 102.036 | 2.864E-5 |
| MORPHO_OIV_204 | chr14_27915149 | 14 | 108.743 | 5.632E-7 |
| MORPHO_OIV_204 | chr14_27931961 | 14 | 109.329 | 2.922E-6 |
| MORPHO_OIV_204 | chr14_28500348 | 14 | 113.236 | 5.25E-7 |
| MORPHO_OIV_204 | chr14_28500407 | 14 | 113.236 | 5.25E-7 |
| MORPHO_OIV_204 | chr14_28466109 | 14 | 113.236 | 6.805E-7 |
| MORPHO_OIV_204 | chr14_28603413 | 14 | 113.301 | 3.22E-7 |
| MORPHO_OIV_204 | chr14_29206455 | 14 | 122.677 | 1.091E-8 |
| MORPHO_OIV_204 | chr14_30023508 | 14 | 124.175 | 1.118E-6 |
| NB_CLUST_PLANT | chr5_1219467 | 5 | 7.323 | 9.516E-6 |
| NB_CLUST_PLANT | chr5_1541332 | 5 | 8.43 | 1.212E-5 |
| NB_CLUST_PLANT | chr17_1289030 | 17 | 6.17 | 4.201E-6 |
| NB_CLUST_PLANT | chr17_2154133 | 17 | 18.51 | 5.864E-6 |
| YIELD_PLANT | chr14_22600042 | 14 | 83.091 | 1.089E-5 |
| YIELD_PLANT | chr14_26074715 | 14 | 95.786 | 1.577E-5 |
| SCLUST_W | chr14_22600042 | 14 | 83.091 | 2.579E-5 |
| SCLUST_W | chr14_26112922 | 14 | 96.307 | 3.334E-5 |
| YIELD_OIV_504 | chr14_22552184 | 14 | 82.374 | 2.124E-5 |
| YIELD_OIV_504 | chr14_22456692 | 14 | 82.374 | 7.257E-6 |
| YIELD_OIV_504 | chr14_22600042 | 14 | 83.091 | 1.698E-6 |
| YIELD_OIV_504 | chr14_24642352 | 14 | 87.583 | 8.971E-6 |

| trait | marker | chromosome | position cM | P value |
|---|---|---|---|---|
| YIELD_OIV_504 | chr14_24643545 | 14 | 87.583 | 1.819E-5 |

**Supplementary Table 12B (continued).** Standardized effects and approximate 95% Wald intervals. Abbreviations: SD, standard deviation; SE, standard error; CI, confidence interval.

| trait | marker | marker effect per genotype SD | effect SE | effect 95% CI lower | effect 95% CI upper |
|---|---|---|---|---|---|
| BUD_DATE | chr7_4414937 | -0.432 | 0.103 | -0.634 | -0.231 |
| BUD_DATE | chr9_19559720 | 0.612 | 0.07 | 0.476 | 0.749 |
| BUD_DATE | chr11_12139361 | 0.612 | 0.07 | 0.476 | 0.749 |
| BUD_DATE | chr16_15360686 | 0.683 | 0.154 | 0.381 | 0.984 |
| FLO_50 | chr1_17756488 | -0.1 | 0.02 | -0.14 | -0.06 |
| FLO_50 | chr2_18622604 | -0.078 | 0.018 | -0.114 | -0.042 |
| FLO_50 | chr7_20342111 | -0.109 | 0.023 | -0.155 | -0.064 |
| FLO_50 | chr7_20396421 | -0.106 | 0.024 | -0.152 | -0.059 |
| FLO_50 | chr7_20556531 | -0.103 | 0.024 | -0.149 | -0.056 |
| FLO_50 | chr7_20537728 | -0.105 | 0.023 | -0.151 | -0.059 |
| FLO_50 | chr7_20586233 | -0.103 | 0.024 | -0.149 | -0.056 |
| FLO_50 | chr17_5562930 | -0.129 | 0.029 | -0.185 | -0.072 |
| FLO_50 | chr17_5599964 | -0.106 | 0.025 | -0.156 | -0.057 |
| FLO_50 | chr17_6431844 | -0.116 | 0.026 | -0.167 | -0.065 |
| VER_50 | chr5_20351051 | -1.615 | 0.197 | -2.002 | -1.228 |
| VER_50 | chr16_7122451 | -1.613 | 0.289 | -2.179 | -1.046 |
| VER_50 | chr16_7158029 | -1.483 | 0.279 | -2.03 | -0.937 |
| VER_50 | chr16_14689931 | 2.26 | 0.293 | 1.686 | 2.833 |
| VER_50 | chr16_14689860 | 2.275 | 0.295 | 1.697 | 2.853 |
| VER_50 | chr17_6431844 | -1.205 | 0.276 | -1.747 | -0.664 |
| SBER_W_g | chr2_5278904 | -0.062 | 0.015 | -0.09 | -0.033 |
| SBER_W_g | chr2_5621690 | 0.052 | 0.012 | 0.028 | 0.076 |
| SBER_W_g | chr15_16599366 | 0.044 | 0.01 | 0.024 | 0.064 |

| trait | marker | marker effect per genotype SD | effect SE | effect 95% CI lower | effect 95% CI upper |
|---|---|---|---|---|---|
| SBER_W_g | chr15_16829505 | 0.054 | 0.013 | 0.029 | 0.079 |
| SBER_W_g | chr15_16856009 | 0.056 | 0.013 | 0.031 | 0.082 |
| SBER_W_g | chr17_6431844 | -0.065 | 0.014 | -0.093 | -0.037 |
| SBER_W_g | chr17_6490445 | -0.068 | 0.014 | -0.096 | -0.04 |
| SBER_W_g | chr17_6561310 | -0.07 | 0.014 | -0.098 | -0.042 |
| SBER_W_g | chr17_6561319 | -0.077 | 0.015 | -0.106 | -0.048 |
| SBER_W_g | chr17_6830639 | -0.057 | 0.013 | -0.083 | -0.031 |
| SBER_W_g | chr17_6830638 | -0.059 | 0.013 | -0.085 | -0.032 |
| SBER_W_g | chr17_7025564 | -0.049 | 0.012 | -0.072 | -0.026 |
| TSS | chr6_7263263 | 3.862 | 0.553 | 2.778 | 4.947 |
| TSS | chr8_11465636 | 3.862 | 0.553 | 2.778 | 4.947 |
| TSS | chr9_2155288 | 2.227 | 0.319 | 1.602 | 2.852 |
| TSS | chr12_7185776 | 0.373 | 0.074 | 0.227 | 0.519 |
| TSS | chr12_7746624 | 0.381 | 0.074 | 0.235 | 0.527 |
| TSS | chr12_20333142 | 0.669 | 0.096 | 0.481 | 0.857 |
| TSS | chr13_2515385 | 3.862 | 0.553 | 2.778 | 4.947 |
| TSS | chr14_30192660 | 2.729 | 0.391 | 1.963 | 3.496 |
| BERRY_pH | chr16_14689931 | -0.016 | 0.004 | -0.023 | -0.009 |
| BER_TA_g | chr5_20351051 | -0.069 | 0.015 | -0.098 | -0.04 |
| HARVEST_DATE | chr5_20351051 | -0.105 | 0.014 | -0.132 | -0.077 |
| HARVEST_DATE | chr16_7122451 | -0.101 | 0.019 | -0.139 | -0.064 |
| HARVEST_DATE | chr16_7158029 | -0.099 | 0.018 | -0.135 | -0.064 |
| HARVEST_DATE | chr16_14689931 | 0.14 | 0.02 | 0.101 | 0.18 |
| HARVEST_DATE | chr16_14689860 | 0.14 | 0.02 | 0.1 | 0.18 |

| trait | marker | marker effect per genotype SD | effect SE | effect 95% CI lower | effect 95% CI upper |
|---|---|---|---|---|---|
| HARVEST_DATE | chr17_6431844 | -0.079 | 0.019 | -0.115 | -0.043 |
| MORPHO_OIV_204 | chr1_4926883 | 0.119 | 0.027 | 0.066 | 0.172 |
| MORPHO_OIV_204 | chr1_5742125 | -0.121 | 0.027 | -0.174 | -0.068 |
| MORPHO_OIV_204 | chr13_3543269 | 0.081 | 0.019 | 0.043 | 0.119 |
| MORPHO_OIV_204 | chr14_26846468 | 0.13 | 0.03 | 0.072 | 0.188 |
| MORPHO_OIV_204 | chr14_26942921 | 0.124 | 0.03 | 0.066 | 0.182 |
| MORPHO_OIV_204 | chr14_27915149 | 0.14 | 0.028 | 0.086 | 0.194 |
| MORPHO_OIV_204 | chr14_27931961 | 0.137 | 0.029 | 0.08 | 0.194 |
| MORPHO_OIV_204 | chr14_28500348 | 0.147 | 0.029 | 0.09 | 0.203 |
| MORPHO_OIV_204 | chr14_28500407 | 0.147 | 0.029 | 0.09 | 0.203 |
| MORPHO_OIV_204 | chr14_28466109 | 0.142 | 0.028 | 0.087 | 0.198 |
| MORPHO_OIV_204 | chr14_28603413 | 0.2 | 0.039 | 0.124 | 0.276 |
| MORPHO_OIV_204 | chr14_29206455 | 0.218 | 0.038 | 0.144 | 0.292 |
| MORPHO_OIV_204 | chr14_30023508 | 0.18 | 0.037 | 0.108 | 0.251 |
| NB_CLUST_PLANT | chr5_1219467 | -0.112 | 0.025 | -0.161 | -0.063 |
| NB_CLUST_PLANT | chr5_1541332 | -0.113 | 0.026 | -0.163 | -0.063 |
| NB_CLUST_PLANT | chr17_1289030 | -0.125 | 0.027 | -0.178 | -0.072 |
| NB_CLUST_PLANT | chr17_2154133 | -0.123 | 0.027 | -0.176 | -0.07 |
| YIELD_PLANT | chr14_22600042 | 0.101 | 0.023 | 0.057 | 0.146 |
| YIELD_PLANT | chr14_26074715 | 0.105 | 0.024 | 0.058 | 0.152 |
| SCLUST_W | chr14_22600042 | 0.1 | 0.024 | 0.054 | 0.147 |
| SCLUST_W | chr14_26112922 | 0.074 | 0.018 | 0.039 | 0.108 |
| YIELD_OIV_504 | chr14_22552184 | 0.104 | 0.024 | 0.056 | 0.151 |
| YIELD_OIV_504 | chr14_22456692 | 0.107 | 0.024 | 0.061 | 0.154 |

| trait | marker | marker effect per genotype SD | effect SE | effect 95% CI lower | effect 95% CI upper |
|---|---|---|---|---|---|
| YIELD_OIV_504 | chr14_22600042 | 0.119 | 0.025 | 0.071 | 0.168 |
| YIELD_OIV_504 | chr14_24642352 | 0.115 | 0.026 | 0.065 | 0.166 |
| YIELD_OIV_504 | chr14_24643545 | 0.112 | 0.026 | 0.061 | 0.162 |

**Supplementary Table 12C (continued).** Dosage-pattern diagnostics.

| trait | marker | genotype pattern id | markers in pattern | chromosomes in pattern |
|---|---|---|---|---|
| BUD_DATE | chr7_4414937 | GP011 | 1 | 1 |
| BUD_DATE | chr9_19559720 | GP018 | 2 | 2 |
| BUD_DATE | chr11_12139361 | GP018 | 2 | 2 |
| BUD_DATE | chr16_15360686 | GP047 | 1 | 1 |
| FLO_50 | chr1_17756488 | GP003 | 1 | 1 |
| FLO_50 | chr2_18622604 | GP006 | 1 | 1 |
| FLO_50 | chr7_20342111 | GP012 | 1 | 1 |
| FLO_50 | chr7_20396421 | GP013 | 1 | 1 |
| FLO_50 | chr7_20556531 | GP015 | 2 | 1 |
| FLO_50 | chr7_20537728 | GP014 | 1 | 1 |
| FLO_50 | chr7_20586233 | GP015 | 2 | 1 |
| FLO_50 | chr17_5562930 | GP050 | 1 | 1 |
| FLO_50 | chr17_5599964 | GP051 | 1 | 1 |
| FLO_50 | chr17_6431844 | GP052 | 1 | 1 |
| VER_50 | chr5_20351051 | GP009 | 1 | 1 |
| VER_50 | chr16_7122451 | GP043 | 1 | 1 |
| VER_50 | chr16_7158029 | GP044 | 1 | 1 |
| VER_50 | chr16_14689931 | GP046 | 1 | 1 |
| VER_50 | chr16_14689860 | GP045 | 1 | 1 |
| VER_50 | chr17_6431844 | GP052 | 1 | 1 |
| SBER_W_g | chr2_5278904 | GP004 | 1 | 1 |
| SBER_W_g | chr2_5621690 | GP005 | 1 | 1 |
| SBER_W_g | chr15_16599366 | GP040 | 1 | 1 |

| trait | marker | genotype pattern id | markers in pattern | chromosomes in pattern |
|---|---|---|---|---|
| SBER_W_g | chr15_16829505 | GP041 | 1 | 1 |
| SBER_W_g | chr15_16856009 | GP042 | 1 | 1 |
| SBER_W_g | chr17_6431844 | GP052 | 1 | 1 |
| SBER_W_g | chr17_6490445 | GP053 | 1 | 1 |
| SBER_W_g | chr17_6561310 | GP054 | 1 | 1 |
| SBER_W_g | chr17_6561319 | GP055 | 1 | 1 |
| SBER_W_g | chr17_6830639 | GP057 | 1 | 1 |
| SBER_W_g | chr17_6830638 | GP056 | 1 | 1 |
| SBER_W_g | chr17_7025564 | GP058 | 1 | 1 |
| TSS | chr6_7263263 | GP010 | 2 | 2 |
| TSS | chr8_11465636 | GP016 | 1 | 1 |
| TSS | chr9_2155288 | GP017 | 1 | 1 |
| TSS | chr12_7185776 | GP019 | 1 | 1 |
| TSS | chr12_7746624 | GP020 | 1 | 1 |
| TSS | chr12_20333142 | GP021 | 1 | 1 |
| TSS | chr13_2515385 | GP010 | 2 | 2 |
| TSS | chr14_30192660 | GP039 | 1 | 1 |
| BERRY_pH | chr16_14689931 | GP046 | 1 | 1 |
| BER_TA_g | chr5_20351051 | GP009 | 1 | 1 |
| HARVEST_DATE | chr5_20351051 | GP009 | 1 | 1 |
| HARVEST_DATE | chr16_7122451 | GP043 | 1 | 1 |
| HARVEST_DATE | chr16_7158029 | GP044 | 1 | 1 |
| HARVEST_DATE | chr16_14689931 | GP046 | 1 | 1 |
| HARVEST_DATE | chr16_14689860 | GP045 | 1 | 1 |

| trait | marker | genotype pattern id | markers in pattern | chromosomes in pattern |
|---|---|---|---|---|
| HARVEST_DATE | chr17_6431844 | GP052 | 1 | 1 |
| MORPHO_OIV_204 | chr1_4926883 | GP001 | 1 | 1 |
| MORPHO_OIV_204 | chr1_5742125 | GP002 | 1 | 1 |
| MORPHO_OIV_204 | chr13_3543269 | GP022 | 1 | 1 |
| MORPHO_OIV_204 | chr14_26846468 | GP030 | 1 | 1 |
| MORPHO_OIV_204 | chr14_26942921 | GP031 | 1 | 1 |
| MORPHO_OIV_204 | chr14_27915149 | GP032 | 1 | 1 |
| MORPHO_OIV_204 | chr14_27931961 | GP033 | 1 | 1 |
| MORPHO_OIV_204 | chr14_28500348 | GP035 | 2 | 1 |
| MORPHO_OIV_204 | chr14_28500407 | GP035 | 2 | 1 |
| MORPHO_OIV_204 | chr14_28466109 | GP034 | 1 | 1 |
| MORPHO_OIV_204 | chr14_28603413 | GP036 | 1 | 1 |
| MORPHO_OIV_204 | chr14_29206455 | GP037 | 1 | 1 |
| MORPHO_OIV_204 | chr14_30023508 | GP038 | 1 | 1 |
| NB_CLUST_PLANT | chr5_1219467 | GP007 | 1 | 1 |
| NB_CLUST_PLANT | chr5_1541332 | GP008 | 1 | 1 |
| NB_CLUST_PLANT | chr17_1289030 | GP048 | 1 | 1 |
| NB_CLUST_PLANT | chr17_2154133 | GP049 | 1 | 1 |
| YIELD_PLANT | chr14_22600042 | GP025 | 1 | 1 |
| YIELD_PLANT | chr14_26074715 | GP028 | 1 | 1 |
| SCLUST_W | chr14_22600042 | GP025 | 1 | 1 |
| SCLUST_W | chr14_26112922 | GP029 | 1 | 1 |
| YIELD_OIV_504 | chr14_22552184 | GP024 | 1 | 1 |
| YIELD_OIV_504 | chr14_22456692 | GP023 | 1 | 1 |

| trait | marker | genotype pattern id | markers in pattern | chromosomes in pattern |
|---|---|---|---|---|
| YIELD_OIV_504 | chr14_22600042 | GP025 | 1 | 1 |
| YIELD_OIV_504 | chr14_24642352 | GP026 | 1 | 1 |
| YIELD_OIV_504 | chr14_24643545 | GP027 | 1 | 1 |

**Supplementary Table 13.** Trait-level counts from the additive and exploratory non-additive analyses. Additive entries report BLINK Bonferroni markers, final MLMM cofactors and MM4LMM Bonferroni markers. Non-additive entries report dominant (D), recessive (R) and overdominant (O) Bonferroni candidates in the LD1-LD5 discovery scan and the subset retained by the selected-candidate K+LD1-LD5 sensitivity retest. Abbreviations: GWAS, genome-wide association study; D, dominant; R, recessive; O, overdominant; K, genomic kinship; DAPC, discriminant analysis of principal components; LD1-LD5, the first five DAPC linear discriminant axes.

| Trait | Additive selected signals | Non-additive discovery | K + ancestry retained |
|---|---|---|---|
| Budbreak | BLINK 12; MLMM 2; MM4LMM 2 | D 14; R 15; O 53 | D 1; R 1; O 2 |
| Flowering | BLINK 10; MLMM 4; MM4LMM 11 | D 18; R 13; O 18 | D 5; R 2; O 5 |
| Veraison | BLINK 31; MLMM 4; MM4LMM 216 | D 31; R 25; O 20 | D 2; R 4; O 4 |
| Harvest date | BLINK 27; MLMM 8; MM4LMM 165 | D 33; R 32; O 26 | D 3; R 8; O 4 |
| Mean berry weight | BLINK 24; MLMM 0; MM4LMM 0 | D 17; R 23; O 17 | D 0; R 1; O 0 |
| Berry sugar content | BLINK 20; MLMM 1; MM4LMM 1 | D 23; R 20; O 9 | D 0; R 0; O 0 |
| Berry pH | BLINK 24; MLMM 1; MM4LMM 1 | D 13; R 5; O 8 | D 2; R 1; O 1 |
| Titratable acidity | BLINK 6; MLMM 1; MM4LMM 7 | D 10; R 4; O 8 | D 0; R 0; O 2 |
| Cluster compactness | BLINK 10; MLMM 1; MM4LMM 6 | D 12; R 9; O 11 | D 0; R 0; O 1 |
| Clusters per plant | BLINK 16; MLMM 3; MM4LMM 1 | D 6; R 12; O 14 | D 0; R 4; O 3 |
| Cluster weight per plant | BLINK 10; MLMM 3; MM4LMM 3 | D 6; R 7; O 19 | D 2; R 5; O 5 |
| Mean cluster weight | BLINK 7; MLMM 2; MM4LMM 1 | D 13; R 5; O 12 | D 2; R 0; O 1 |
| Yield per m2 | BLINK 9; MLMM 1; MM4LMM 1 | D 11; R 5; O 9 | D 2; R 1; O 0 |

**Supplementary Table 14A.** Trait-specific inputs, model specification, genomic-control diagnostics and significant-count summary for the three additive GWAS models. This table concerns pooled additive GWAS only; environmental, non-additive, mpQTL and localGEBV inputs are reported separately. Analysis inputs and fitted structure. Abbreviations: GWAS, genome-wide association study; DAPC, discriminant analysis of principal components; BLINK, Bayesian-information and Linkage-disequilibrium Iteratively Nested Keyway; MLMM, multi-locus mixed model; MM4LMM, Min-Max algorithm for large-scale mixed models.

| Trait | Trait label | Method | N individuals | N markers | N markers estimable | N markers nonestimable | DAPC axes | kinship fitted |
|---|---|---|---|---|---|---|---|---|
| BUD_DATE | Budbreak | BLINK | 965 | 140,181 | 140,181 | 0 | 5 | FALSE |
| BUD_DATE | Budbreak | MLMM | 965 | 140,181 | 140,181 | 0 | 5 | TRUE |
| BUD_DATE | Budbreak | MM4LMM | 965 | 140,181 | 140,180 | 1 | 5 | TRUE |
| FLO_50 | Flowering | BLINK | 967 | 139,738 | 139,738 | 0 | 5 | FALSE |
| FLO_50 | Flowering | MLMM | 967 | 139,738 | 139,738 | 0 | 5 | TRUE |
| FLO_50 | Flowering | MM4LMM | 967 | 139,738 | 139,737 | 1 | 5 | TRUE |
| VER_50 | Veraison | BLINK | 968 | 139,406 | 139,406 | 0 | 5 | FALSE |
| VER_50 | Veraison | MLMM | 968 | 139,406 | 139,406 | 0 | 5 | TRUE |
| VER_50 | Veraison | MM4LMM | 968 | 139,406 | 139,405 | 1 | 5 | TRUE |
| HARVEST_DATE | Harvest date | BLINK | 948 | 140,077 | 140,077 | 0 | 5 | FALSE |
| HARVEST_DATE | Harvest date | MLMM | 948 | 140,077 | 140,077 | 0 | 5 | TRUE |
| HARVEST_DATE | Harvest date | MM4LMM | 948 | 140,077 | 140,076 | 1 | 5 | TRUE |
| SBER_W_g | Mean berry weight | BLINK | 943 | 139,638 | 139,638 | 0 | 5 | FALSE |
| SBER_W_g | Mean berry weight | MLMM | 943 | 139,638 | 139,638 | 0 | 5 | TRUE |
| SBER_W_g | Mean berry weight | MM4LMM | 943 | 139,638 | 139,637 | 1 | 5 | TRUE |
| TSS | Berry sugar content | BLINK | 932 | 139,300 | 139,300 | 0 | 5 | FALSE |
| TSS | Berry sugar content | MLMM | 932 | 139,300 | 139,300 | 0 | 5 | TRUE |
| TSS | Berry sugar content | MM4LMM | 932 | 139,300 | 139,299 | 1 | 5 | TRUE |
| BERRY_pH | Berry pH | BLINK | 963 | 139,173 | 139,173 | 0 | 5 | FALSE |
| BERRY_pH | Berry pH | MLMM | 963 | 139,173 | 139,173 | 0 | 5 | TRUE |
| BERRY_pH | Berry pH | MM4LMM | 963 | 139,173 | 139,172 | 1 | 5 | TRUE |
| BER_TA_g | Titratable acidity | BLINK | 963 | 139,164 | 139,164 | 0 | 5 | FALSE |
| BER_TA_g | Titratable acidity | MLMM | 963 | 139,164 | 139,164 | 0 | 5 | TRUE |

| Trait | Trait label | Method | N individuals | N markers | N markers estimable | N markers nonestimable | DAPC axes | kinship fitted |
|---|---|---|---|---|---|---|---|---|
| BER_TA_g | Titratable acidity | MM4LMM | 963 | 139,164 | 139,163 | 1 | 5 | TRUE |
| MORPHO_OIV_204 | Cluster compactness | BLINK | 956 | 139,223 | 139,223 | 0 | 5 | FALSE |
| MORPHO_OIV_204 | Cluster compactness | MLMM | 956 | 139,223 | 139,223 | 0 | 5 | TRUE |
| MORPHO_OIV_204 | Cluster compactness | MM4LMM | 956 | 139,223 | 139,222 | 1 | 5 | TRUE |
| NB_CLUST_PLANT | Clusters per plant | BLINK | 916 | 138,294 | 138,294 | 0 | 5 | FALSE |
| NB_CLUST_PLANT | Clusters per plant | MLMM | 916 | 138,294 | 138,294 | 0 | 5 | TRUE |
| NB_CLUST_PLANT | Clusters per plant | MM4LMM | 916 | 138,294 | 138,293 | 1 | 5 | TRUE |
| YIELD_PLANT | Cluster weight per plant | BLINK | 908 | 137,392 | 137,392 | 0 | 5 | FALSE |
| YIELD_PLANT | Cluster weight per plant | MLMM | 908 | 137,392 | 137,392 | 0 | 5 | TRUE |
| YIELD_PLANT | Cluster weight per plant | MM4LMM | 908 | 137,392 | 137,391 | 1 | 5 | TRUE |
| SCLUST_W | Mean cluster weight | BLINK | 899 | 137,285 | 137,285 | 0 | 5 | FALSE |
| SCLUST_W | Mean cluster weight | MLMM | 899 | 137,285 | 137,285 | 0 | 5 | TRUE |
| SCLUST_W | Mean cluster weight | MM4LMM | 899 | 137,285 | 137,284 | 1 | 5 | TRUE |
| YIELD_OIV_504 | Yield per m² | BLINK | 927 | 139,102 | 139,102 | 0 | 5 | FALSE |
| YIELD_OIV_504 | Yield per m² | MLMM | 927 | 139,102 | 139,102 | 0 | 5 | TRUE |
| YIELD_OIV_504 | Yield per m² | MM4LMM | 927 | 139,102 | 139,101 | 1 | 5 | TRUE |

**Supplementary Table 14B (continued).** Thresholds, selected signals and genomic-control diagnostics. Abbreviations: FDR, false discovery rate; GC, genomic control. Other abbreviations are defined in Supplementary Table 14A.

| Trait | Method | Bonferroni threshold | Bonferroni hits step1 or single marker | MLMM selected cofactors | FDR 5% hits descriptive | Lambda GC |
|---|---|---|---|---|---|---|
| BUD_DATE | BLINK | 3.567E-7 | 12 | | 16 | 1.232 |
| BUD_DATE | MLMM | 3.567E-7 | 2 | 2 | 3 | 0.923 |
| BUD_DATE | MM4LMM | 3.567E-7 | 2 | | 3 | 0.926 |
| FLO_50 | BLINK | 3.578E-7 | 10 | | 20 | 1.022 |
| FLO_50 | MLMM | 3.578E-7 | 9 | 4 | 91 | 0.807 |
| FLO_50 | MM4LMM | 3.578E-7 | 11 | | 115 | 0.81 |
| VER_50 | BLINK | 3.587E-7 | 31 | | 39 | 1.077 |
| VER_50 | MLMM | 3.587E-7 | 204 | 4 | 287 | 0.842 |
| VER_50 | MM4LMM | 3.587E-7 | 216 | | 301 | 0.845 |
| HARVEST_DATE | BLINK | 3.569E-7 | 27 | | 35 | 1.104 |
| HARVEST_DATE | MLMM | 3.569E-7 | 158 | 8 | 348 | 0.863 |
| HARVEST_DATE | MM4LMM | 3.569E-7 | 165 | | 360 | 0.867 |
| SBER_W_g | BLINK | 3.581E-7 | 24 | | 32 | 1.319 |
| SBER_W_g | MLMM | 3.581E-7 | 0 | 0 | 0 | 0.954 |
| SBER_W_g | MM4LMM | 3.581E-7 | 0 | | 0 | 0.958 |
| TSS | BLINK | 3.589E-7 | 20 | | 25 | 1.211 |
| TSS | MLMM | 3.589E-7 | 1 | 1 | 7 | 0.949 |
| TSS | MM4LMM | 3.589E-7 | 1 | | 9 | 0.957 |
| BERRY_pH | BLINK | 3.593E-7 | 24 | | 30 | 1.455 |
| BERRY_pH | MLMM | 3.593E-7 | 0 | 1 | 0 | 0.941 |
| BERRY_pH | MM4LMM | 3.593E-7 | 1 | | 1 | 0.944 |
| BER_TA_g | BLINK | 3.593E-7 | 6 | | 14 | 1.321 |
| BER_TA_g | MLMM | 3.593E-7 | 4 | 1 | 24 | 0.938 |

| Trait | Method | Bonferroni threshold | Bonferroni hits step1 or single marker | MLMM selected cofactors | FDR 5% hits descriptive | Lambda GC |
|---|---|---|---|---|---|---|
| BER_TA_g | MM4LMM | 3.593E-7 | 7 | | 25 | 0.942 |
| MORPHO_OIV_204 | BLINK | 3.591E-7 | 10 | | 14 | 1.174 |
| MORPHO_OIV_204 | MLMM | 3.591E-7 | 3 | 1 | 11 | 0.919 |
| MORPHO_OIV_204 | MM4LMM | 3.591E-7 | 6 | | 12 | 0.923 |
| NB_CLUST_PLANT | BLINK | 3.615E-7 | 16 | | 20 | 1.337 |
| NB_CLUST_PLANT | MLMM | 3.615E-7 | 1 | 3 | 1 | 0.912 |
| NB_CLUST_PLANT | MM4LMM | 3.615E-7 | 1 | | 4 | 0.916 |
| YIELD_PLANT | BLINK | 3.639E-7 | 10 | | 13 | 1.076 |
| YIELD_PLANT | MLMM | 3.639E-7 | 3 | 3 | 26 | 0.905 |
| YIELD_PLANT | MM4LMM | 3.639E-7 | 3 | | 27 | 0.909 |
| SCLUST_W | BLINK | 3.642E-7 | 7 | | 19 | 1.406 |
| SCLUST_W | MLMM | 3.642E-7 | 1 | 2 | 3 | 0.944 |
| SCLUST_W | MM4LMM | 3.642E-7 | 1 | | 3 | 0.948 |
| YIELD_OIV_504 | BLINK | 3.594E-7 | 9 | | 14 | 1.103 |
| YIELD_OIV_504 | MLMM | 3.594E-7 | 1 | 1 | 13 | 0.932 |
| YIELD_OIV_504 | MM4LMM | 3.594E-7 | 1 | | 14 | 0.936 |

**Supplementary Table 14C.** Additive GWAS signals supported by at least two models. All models included five DAPC axes; MLMM and MM4LMM also included genomic kinship. Chr, chromosome; bp, base pair. Mb, megabase; NA, no signal reported as shared by two models.

| Trait | Chr | Position or span (bp unless Mb stated) | Models |
|---|---|---|---|
| Date of 50% of budbreak | 4 | 3,285,585 | MLMM and MM4LMM |
| Date of 50% of budbreak | 8 | 20,245,338 | BLINK and MLMM |
| Date of 50% of budbreak | 13 | 17.9 -18.1 Mb | BLINK and MM4LMM |
| Date of 50% of budbreak | 16 | 14.1 -15.1 Mb | BLINK and MLMM |
| Date of 50% flowering | 6 | 16,814,179 | MLMM and MM4LMM |
| Date of 50% flowering | 11 | 6,089,413 | MLMM, BLINK, and MM4LMM |
| Date of 50% flowering | 13 | 24.2 -24.5 Mb | BLINK and MM4LMM |
| Date of 50% veraison | 16 | 14.1 -16.4 Mb | BLINK and MLMM |
| Mean berry weight | NA | NA | |
| Berry sugar content | 19 | 6.3 -6.8 Mb | BLINK and MLMM |
| Berry pH | NA | NA | |
| Berry titrable acidity | 16 | 13.9 -15.2 Mb | MLMM, BLINK, and MM4LMM |
| Harvest date | 1 | 9,496,721 | BLINK and MLMM |
| Harvest date | 16 | 14.2 -15.9 Mb | BLINK and MLMM |
| Cluster compactness | 1 | 4.9 -6.4 Mb | MLMM, BLINK, and MM4LMM |
| Number of clusters per plant | 11 | 0.6 -1.7 Mb | BLINK and MLMM |
| Total cluster weight per plant | 5 | 12.6 -12.7 Mb | MLMM and MM4LMM |
| Total cluster weight per plant | 9 | 6,629,813 | MLMM, BLINK, and MM4LMM |
| Total cluster weight per plant | 12 | 17.9 -18.0 Mb | BLINK and MM4LMM |
| Total cluster weight per plant | 14 | 17,151,022 | BLINK and MM4LMM |
| Mean cluster weight | NA | NA | |
| Yield per square meter | 9 | 6,629,813 | MLMM and MM4LMM |

**Supplementary Table 15.** Exact single-family QTL-GWAS interval evidence and population-centred local-LD refinement. Tier 1 denotes LD-concordant support from at least two adjusted GWAS models; Tier 2 denotes support from one model or spatially discordant multi-model signals; No overlap denotes no exact QTL-GWAS overlap. Abbreviations: QTL, quantitative trait locus; GWAS, genome-wide association study; LD, linkage disequilibrium. QTL024-QTL026 have suggestive family-QTL support (0.05 < permutation P ≤ 0.10); framework-overlap flags do not change this significance status.

| QTL ID | Trait | Pop | PhysicalChr | QTL low | QTL high | LeadSNP | LeadMethod | EvidenceTier |
|---|---|---|---|---|---|---|---|---|
| QTL001 | FLO_50 | 42050 | 2 | 5E+5 | 4,700,000 | | | No overlap |
| QTL002 | FLO_50 | 42050 | 7 | 17,500,000 | 24,300,000 | | | No overlap |
| QTL003 | VER_50 | 42050 | 8 | 14,900,000 | 19,200,000 | | | No overlap |
| QTL004 | SCLUST_W | 42050 | 8 | 2,100,000 | 21,200,000 | | | No overlap |
| QTL005 | YIELD_PLANT | 42050 | 11 | 3,200,000 | 11,300,000 | chr11_4574451 | BLINK_LD1-LD5 | Tier 2 |
| QTL006 | YIELD_OIV_504 | 42050 | 11 | 3,200,000 | 11,300,000 | chr11_8149404 | BLINK_LD1-LD5 | Tier 2 |
| QTL007 | FLO_50 | 42050 | 14 | 11,300,000 | 29,700,000 | chr14_28852023 | BLINK_LD1-LD5 | Tier 2 |
| QTL008 | VER_50 | 42050 | 14 | 24,300,000 | 28,800,000 | chr14_27349067 | BLINK_LD1-LD5 | Tier 2 |
| QTL009 | BERRY_pH | 42050 | 17 | 5,900,000 | 9,600,000 | | | No overlap |
| QTL010 | BER_TA_g | 42050 | 17 | 6,800,000 | 16,600,000 | | | No overlap |
| QTL011 | VER_50 | 50001 | 16 | 15,900,000 | 18,200,000 | chr16_17401013 | BLINK_LD1-LD5;MM4LMM_K_LD1-LD5 | Tier 1 |
| QTL012 | BERRY_pH | 50001 | 16 | 17,100,000 | 19,100,000 | | | No overlap |
| QTL013 | TSS | 50013 | 7 | 7E+5 | 3,900,000 | | | No overlap |
| QTL014 | SBER_W_g | 50013 | 7 | 4,700,000 | 28,700,000 | chr7_23591061 | BLINK_LD1-LD5 | Tier 2 |
| QTL015 | MORPHO_OIV_204 | 50013 | 14 | 20,700,000 | 30,800,000 | chr14_26574115 | BLINK_LD1-LD5 | Tier 2 |
| QTL016 | YIELD_PLANT | 50013 | 14 | 20,700,000 | 29,600,000 | | | No overlap |
| QTL017 | YIELD_OIV_504 | 50013 | 14 | 19,500,000 | 29,600,000 | | | No overlap |
| QTL018 | SCLUST_W | 50013 | 14 | 22,700,000 | 29,600,000 | chr14_27028690 | BLINK_LD1-LD5 | Tier 2 |
| QTL019 | TSS | 50013 | 17 | 4E+6 | 6E+6 | | | No overlap |
| QTL020 | MORPHO_OIV_204 | 50013 | 17 | 6,100,000 | 9,200,000 | | | No overlap |
| QTL021 | YIELD_PLANT | 50013 | 17 | 6,800,000 | 10,900,000 | chr17_9255410 | BLINK_LD1-LD5 | Tier 2 |
| QTL022 | SCLUST_W | 50013 | 17 | 6,500,000 | 18,800,000 | | | No overlap |

| QTL ID | Trait | Pop | PhysicalChr | QTL low | QTL high | LeadSNP | LeadMethod | EvidenceTier |
|---|---|---|---|---|---|---|---|---|
| QTL023 | YIELD_OIV_504 | 50013 | 17 | 6,700,000 | 10,900,000 | | | No overlap |
| QTL024 | SBER_W_g | 50015 | 2 | 5,300,000 | 7,700,000 | chr2_7147735 | BLINK_LD1-LD5 | Tier 2 |
| QTL025 | YIELD_PLANT | 50015 | 5 | 1,200,000 | 6,300,000 | | | No overlap |
| QTL026 | YIELD_OIV_504 | 50015 | 5 | 7E+5 | 6,300,000 | | | No overlap |
| QTL027 | HARVEST_DATE | 50015 | 8 | 2E+5 | 20,700,000 | chr8_5781365 | BLINK_LD1-LD5;MLMM_K_LD1-LD5;MM4LMM_K_LD1-LD5 | Tier 1 |
| QTL028 | BER_TA_g | 50015 | 13 | 3,900,000 | 7,100,000 | chr13_5514868 | BLINK_LD1-LD5 | Tier 2 |
| QTL029 | BERRY_pH | 50015 | 13 | 4,400,000 | 5,800,000 | | | No overlap |
| QTL030 | TSS | 50025 | 1 | 20,400,000 | 23,700,000 | chr1_23358100 | BLINK_LD1-LD5 | Tier 2 |
| QTL031 | BUD_DATE | 50025 | 4 | 8,100,000 | 15,900,000 | | | No overlap |
| QTL032 | BER_TA_g | 50025 | 6 | 15,400,000 | 21,200,000 | | | No overlap |
| QTL033 | BUD_DATE | 50025 | 7 | 4E+5 | 18,200,000 | | | No overlap |
| QTL034 | BER_TA_g | 50025 | 14 | 2.1E+7 | 29,800,000 | chr14_22281893 | BLINK_LD1-LD5 | Tier 2 |
| QTL035 | FLO_50 | 50025 | 14 | 17,900,000 | 29,800,000 | chr14_28852023 | BLINK_LD1-LD5 | Tier 2 |
| QTL036 | SBER_W_g | 50025 | 17 | 6,600,000 | 10,300,000 | | | No overlap |
| QTL037 | TSS | 50025 | 17 | 1,900,000 | 4,300,000 | chr17_2082982 | BLINK_LD1-LD5 | Tier 2 |
| QTL038 | BER_TA_g | 50025 | 17 | 1,600,000 | 4,900,000 | | | No overlap |
| QTL039 | TSS | 50025 | 18 | 12,300,000 | 29,900,000 | chr18_13434839 | BLINK_LD1-LD5 | Tier 2 |
| QTL040 | BER_TA_g | 50025 | 16 | 12,500,000 | 15,200,000 | chr16_15035128 | MLMM_K_LD1-LD5;MM4LMM_K_LD1-LD5 | Tier 1 |
| QTL041 | HARVEST_DATE | 50025 | 16 | 1.4E+7 | 15,200,000 | chr16_15035128 | BLINK_LD1-LD5;MLMM_K_LD1-LD5;MM4LMM_K_LD1-LD5 | Tier 1 |
| QTL042 | VER_50 | 50025 | 16 | 14,200,000 | 15,200,000 | chr16_15035128 | BLINK_LD1-LD5;MLMM_K_LD1-LD5;MM4LMM_K_LD1-LD5 | Tier 1 |
| QTL043 | MORPHO_OIV_204 | 50035 | 1 | 2E+6 | 5,800,000 | chr1_5670971 | BLINK_LD1-LD5;MM4LMM_K_LD1-LD5 | Tier 1 |
| QTL044 | BUD_DATE | 50035 | 2 | 2,400,000 | 5,700,000 | | | No overlap |
| QTL045 | VER_50 | 50035 | 2 | 2,300,000 | 7,400,000 | chr2_2851725 | BLINK_LD1-LD5 | Tier 2 |
| QTL046 | HARVEST_DATE | 50035 | 2 | 9E+5 | 5,700,000 | | | No overlap |

| QTL ID | Trait | Pop | PhysicalChr | QTL low | QTL high | LeadSNP | LeadMethod | EvidenceTier |
|---|---|---|---|---|---|---|---|---|
| QTL047 | MORPHO_OIV_204 | 50035 | 2 | 5,200,000 | 6,200,000 | | | No overlap |
| QTL048 | SCLUST_W | 50035 | 2 | 5,700,000 | 6,200,000 | | | No overlap |
| QTL049 | YIELD_OIV_504 | 50035 | 2 | 5,500,000 | 7,200,000 | | | No overlap |
| QTL050 | SBER_W_g | 50035 | 2 | 8,500,000 | 16,400,000 | | | No overlap |
| QTL051 | NB_CLUST_PLANT | 50035 | 3 | 4,900,000 | 20,700,000 | | | No overlap |
| QTL052 | BERRY_pH | 50035 | 4 | 8,100,000 | 18,900,000 | chr4_11934711 | BLINK_LD1-LD5 | Tier 2 |
| QTL053 | BER_TA_g | 50035 | 4 | 5,200,000 | 18,900,000 | | | No overlap |
| QTL054 | TSS | 50035 | 6 | 3,600,000 | 7,300,000 | | | No overlap |
| QTL055 | SBER_W_g | 50035 | 7 | 2,500,000 | 22,100,000 | | | No overlap |
| QTL056 | NB_CLUST_PLANT | 50035 | 7 | 1,800,000 | 5,800,000 | chr7_2424949 | BLINK_LD1-LD5 | Tier 2 |
| QTL057 | FLO_50 | 50035 | 7 | 4,100,000 | 20,600,000 | | | No overlap |
| QTL058 | TSS | 50035 | 8 | 17,700,000 | 20,300,000 | | | No overlap |
| QTL059 | BUD_DATE | 50035 | 9 | 9E+5 | 5,500,000 | chr9_2360990 | BLINK_LD1-LD5 | Tier 2 |
| QTL060 | FLO_50 | 50035 | 9 | 1E+5 | 13,800,000 | | | No overlap |
| QTL061 | TSS | 50035 | 9 | 1E+5 | 2,700,000 | | | No overlap |
| QTL062 | SBER_W_g | 50035 | 11 | 9,600,000 | 19,500,000 | chr11_18029044 | BLINK_LD1-LD5 | Tier 2 |
| QTL063 | YIELD_OIV_504 | 50035 | 11 | 9,600,000 | 19,400,000 | | | No overlap |
| QTL064 | YIELD_PLANT | 50035 | 14 | 17,400,000 | 27,400,000 | | | No overlap |
| QTL065 | YIELD_OIV_504 | 50035 | 14 | 1.8E+7 | 28,400,000 | | | No overlap |
| QTL066 | NB_CLUST_PLANT | 50035 | 14 | 20,600,000 | 30,300,000 | | | No overlap |
| QTL067 | BUD_DATE | 50035 | 16 | 2E+6 | 16,800,000 | chr16_15107521 | BLINK_LD1-LD5 | Tier 2 |
| QTL068 | SCLUST_W | 50035 | 16 | 14,400,000 | 16,700,000 | chr16_15542722 | BLINK_LD1-LD5;MLMM_K_LD1-LD5 | Tier 1 |
| QTL069 | YIELD_OIV_504 | 50035 | 16 | 8E+5 | 16,700,000 | chr16_12507629 | BLINK_LD1-LD5 | Tier 2 |
| QTL070 | BER_TA_g | 50035 | 16 | 1.5E+7 | 17,300,000 | chr16_15035128 | MLMM_K_LD1-LD5;MM4LMM_K_LD1-LD5 | Tier 1 |
| QTL071 | BERRY_pH | 50035 | 16 | 1.5E+7 | 16,800,000 | chr16_16539486 | BLINK_LD1-LD5 | Tier 2 |

| QTL ID | Trait | Pop | PhysicalChr | QTL low | QTL high | LeadSNP | LeadMethod | EvidenceTier |
|---|---|---|---|---|---|---|---|---|
| QTL072 | HARVEST_DATE | 50035 | 16 | 15,400,000 | 16,800,000 | chr16_16718042 | BLINK_LD1-LD5;MM4LMM_K_LD1-LD5 | Tier 1 |
| QTL073 | VER_50 | 50035 | 16 | 1E+5 | 17,600,000 | chr16_15035128 | BLINK_LD1-LD5;MLMM_K_LD1-LD5;MM4LMM_K_LD1-LD5 | Tier 1 |
| QTL074 | MORPHO_OIV_204 | 50035 | 16 | 1E+5 | 15,300,000 | | | No overlap |
| QTL075 | MORPHO_OIV_204 | 50035 | 18 | 7,900,000 | 34,300,000 | | | No overlap |
| QTL076 | TSS | 50035 | 18 | 12,700,000 | 31,100,000 | chr18_13434839 | BLINK_LD1-LD5 | Tier 2 |
| QTL077 | FLO_50 | 50015 | 17 | 5,508,736 | 6,478,604 | | | No overlap |
| QTL078 | HARVEST_DATE | 50015 | 18 | 22,755,801 | 29,601,363 | chr18_23927297 | BLINK_LD1-LD5 | Tier 2 |
| QTL079 | NB_CLUST_PLANT | 50015 | 17 | 3,599 | 4,478,988 | | | No overlap |

**Supplementary Table 16.** Prioritized PN40024.v4 positional candidate genes in QTL-GWAS refined intervals. Tier 1 denotes LD-concordant support from at least two adjusted GWAS models; Tier 2 denotes support from one model or spatially discordant multi-model signals. Automated positional and functional ranking does not establish causality. Abbreviations: QTL, quantitative trait locus; GWAS, genome-wide association study.

| QTL ID | Trait | EvidenceTier | GeneID | Description |
|---|---|---|---|---|
| QTL005 | YIELD_PLANT | Tier 2 | Vitvi11g01435 | XP_002285155.1 probable sugar phosphate/phosphate translocator At5g25400 |
| QTL005 | YIELD_PLANT | Tier 2 | Vitvi11g00476 | XP_010656044.1 CBL-interacting protein kinase 09 isoform X1 |
| QTL005 | YIELD_PLANT | Tier 2 | Vitvi11g00451 | XP_002281941.1 adenine nucleotide transporter BT1, chloroplastic/mitochondrial |
| QTL006 | YIELD_OIV_504 | Tier 2 | Vitvi11g00710 | XP_002278056.1 signal recognition particle receptor subunit alpha homolog |
| QTL006 | YIELD_OIV_504 | Tier 2 | Vitvi11g01504 | XP_002277887.1 homoserine kinase |
| QTL006 | YIELD_OIV_504 | Tier 2 | Vitvi11g00677 | NP_001268212.1CBL-interacting serine/threonine-protein kinase 6-like |
| QTL007 | FLO_50 | Tier 2 | Vitvi14g01845 | XP_002276072.2 transcription factor MYB35 |
| QTL007 | FLO_50 | Tier 2 | Vitvi14g01867 | RVX08741.1General negative regulator of transcription subunit 4 |
| QTL007 | FLO_50 | Tier 2 | Vitvi14g01870 | XP_010660999.1 E2F transcription factor-like E2FF isoform X1 |
| QTL008 | VER_50 | Tier 2 | Vitvi14g04613 | RVW29386.1Abscisic acid receptor PYL4 |
| QTL008 | VER_50 | Tier 2 | Vitvi14g02998 | XP_010660834.1 GATA transcription factor 7 |
| QTL008 | VER_50 | Tier 2 | Vitvi14g01667 | XP_002271589.2 uridine kinase-like protein 5 isoform X1 |
| QTL011 | VER_50 | Tier 1 | Vitvi16g01015 | XP_002267853.1 transcription factor LAF1 |
| QTL011 | VER_50 | Tier 1 | Vitvi16g01017 | XP_002267986.1 transcription repressor MYB6 |
| QTL011 | VER_50 | Tier 1 | Vitvi16g01007 | XP_002263340.2 flap endonuclease GEN-like 2 |
| QTL014 | SBER_W_g | Tier 2 | Vitvi07g04654 | XP_002263151.3 LRR receptor-like serine/threonine-protein kinase HSL2 |

| QTL ID | Trait | EvidenceTier | GeneID | Description |
|---|---|---|---|---|
| QTL014 | SBER_W_g | Tier 2 | Vitvi07g04655 | RVW17919.1LRR receptor-like serine/threonine-protein kinase HSL2 |
| QTL014 | SBER_W_g | Tier 2 | Vitvi07g04652 | XP_003635561.1 leucine-rich repeat receptor-like serine/threonine-protein kinase BAM1 |
| QTL015 | MORPHO_OIV_204 | Tier 2 | Vitvi14g01576 | XP_010660787.1 probable inactive leucine-rich repeat receptor-like protein kinase At3g03770 |
| QTL015 | MORPHO_OIV_204 | Tier 2 | Vitvi14g01574 | XP_002263142.1 probable adenylate kinase 7, mitochondrial |
| QTL015 | MORPHO_OIV_204 | Tier 2 | Vitvi14g02984 | XP_003633787.1 uridine kinase-like protein 1, chloroplastic |
| QTL018 | SCLUST_W | Tier 2 | Vitvi14g01596 | XP_002265308.1 lysine histidine transporter 1 |
| QTL018 | SCLUST_W | Tier 2 | Vitvi14g01593 | XP_002264926.1 thymidylate kinase |
| QTL018 | SCLUST_W | Tier 2 | Vitvi14g01667 | XP_002271589.2 uridine kinase-like protein 5 isoform X1 |
| QTL021 | YIELD_PLANT | Tier 2 | Vitvi17g00727 | XP_002280523.1 serine/threonine-protein kinase AtPK2/AtPK19 |
| QTL021 | YIELD_PLANT | Tier 2 | Vitvi17g04218 | RVW52795.1Serine/threonine-protein kinase AtPK2/AtPK19 |
| QTL021 | YIELD_PLANT | Tier 2 | Vitvi17g00759 | XP_002283469.1 nucleobase-ascorbate transporter 6 |
| QTL024 | SBER_W_g | Tier 2 | Vitvi02g00682 | XP_002280602.1 probable serine/threonine-protein kinase At4g35230 |
| QTL024 | SBER_W_g | Tier 2 | Vitvi02g00684 | XP_002278359.2 pyruvate kinase isozyme G, chloroplastic isoform X1 |
| QTL024 | SBER_W_g | Tier 2 | Vitvi02g00653 | XP_002278917.1 expansin-like B1 |
| QTL027 | HARVEST_DATE | Tier 1 | Vitvi08g00289 | XP_002264091.1 putative deoxyribonuclease TATDN1 isoform X2 |
| QTL028 | BER_TA_g | Tier 2 | Vitvi13g00594 | XP_019079997.1 aminodeoxychorismate synthase, chloroplastic isoform X3 |

| QTL ID | Trait | EvidenceTier | GeneID | Description |
|---|---|---|---|---|
| QTL028 | BER_TA_g | Tier 2 | Vitvi13g00596 | XP_002274359.2 aminodeoxychorismate synthase, chloroplastic |
| QTL028 | BER_TA_g | Tier 2 | Vitvi13g00597 | XP_002275232.1 chorismate synthase 1, chloroplastic |
| QTL028 | BER_TA_g | Tier 2 | Vitvi13g00601 | XP_002275050.1 V-type proton ATPase subunit G1 |
| QTL030 | TSS | Tier 2 | Vitvi01g01719 | XP_002264875.1 bidirectional sugar transporter SWEET15 |
| QTL030 | TSS | Tier 2 | Vitvi01g04469 | |
| QTL030 | TSS | Tier 2 | Vitvi01g01740 | XP_010657142.1 putative pentatricopeptide repeat-containing protein At5g13230, mitochondrial |
| QTL034 | BER_TA_g | Tier 2 | Vitvi14g04452 | XP_003633754.1 vacuolar cation/proton exchanger 2-like |
| QTL034 | BER_TA_g | Tier 2 | Vitvi14g01269 | XP_003633754.1 vacuolar cation/proton exchanger 2-like |
| QTL034 | BER_TA_g | Tier 2 | Vitvi14g01307 | XP_002282329.1 probable sodium-coupled neutral amino acid transporter 6 |
| QTL035 | FLO_50 | Tier 2 | Vitvi14g01845 | XP_002276072.2 transcription factor MYB35 |
| QTL035 | FLO_50 | Tier 2 | Vitvi14g01867 | RVX08741.1General negative regulator of transcription subunit 4 |
| QTL035 | FLO_50 | Tier 2 | Vitvi14g01870 | XP_010660999.1 E2F transcription factor-like E2FF isoform X1 |
| QTL037 | TSS | Tier 2 | Vitvi17g00162 | XP_034712099.1potassium transporter 1-like isoform X1 |
| QTL037 | TSS | Tier 2 | Vitvi17g00143 | XP_002283475.1 probable aspartyl aminopeptidase |
| QTL037 | TSS | Tier 2 | Vitvi17g00142 | XP_002283508.1 adenylate kinase 4 |
| QTL039 | TSS | Tier 2 | Vitvi18g01215 | XP_002265836.1 bidirectional sugar transporter SWEET1 |
| QTL039 | TSS | Tier 2 | Vitvi18g01216 | XP_010664874.2 pentatricopeptide repeat-containing protein At1g08070, chloroplastic |
| QTL039 | TSS | Tier 2 | Vitvi18g01217 | RVW32582.1Pentatricopeptide repeat-containing protein |
| QTL040 | BER_TA_g | Tier 1 | Vitvi16g00860 | XP_010662360.1 K(+) efflux antiporter 5 isoform X2 |

| QTL ID | Trait | EvidenceTier | GeneID | Description |
| --- | --- | --- | --- | --- |
| QTL040 | BER_TA_g | Tier 1 | Vitvi16g00853 | XP_002268932.2 probable aminotransferase ACS12 isoform X1 |
| QTL040 | BER_TA_g | Tier 1 | Vitvi16g00852 | XP_034711320.1protein enabled homolog |
| QTL041 | HARVEST_DATE | Tier 1 | Vitvi16g00849 | XP_002269179.1 anthocyanidin 3-O-glucosyltransferase 5 |
| QTL041 | HARVEST_DATE | Tier 1 | Vitvi16g00853 | XP_002268932.2 probable aminotransferase ACS12 isoform X1 |
| QTL041 | HARVEST_DATE | Tier 1 | Vitvi16g00852 | XP_034711320.1protein enabled homolog |
| QTL042 | VER_50 | Tier 1 | Vitvi16g00849 | XP_002269179.1 anthocyanidin 3-O-glucosyltransferase 5 |
| QTL042 | VER_50 | Tier 1 | Vitvi16g00853 | XP_002268932.2 probable aminotransferase ACS12 isoform X1 |
| QTL042 | VER_50 | Tier 1 | Vitvi16g00852 | XP_034711320.1protein enabled homolog |
| QTL043 | MORPHO_OIV_204 | Tier 1 | Vitvi01g00476 | QCD85355.1B-cell receptor-associated protein 29/31 |
| QTL043 | MORPHO_OIV_204 | Tier 1 | Vitvi01g00475 | XP_002281181.1 probable inactive receptor kinase RLK902 |
| QTL043 | MORPHO_OIV_204 | Tier 1 | Vitvi01g00473 | XP_002284855.1 protein TIFY 6B isoform X1 |
| QTL045 | VER_50 | Tier 2 | Vitvi02g00317 | XP_002270239.2 transcription factor EGL1 |
| QTL045 | VER_50 | Tier 2 | Vitvi02g00316 | XP_019072492.1 gibberellin 3-beta-dioxygenase 4 |
| QTL045 | VER_50 | Tier 2 | Vitvi02g00347 | XP_002269359.1 nuclear transcription factor Y subunit C-3 |
| QTL052 | BERRY_pH | Tier 2 | Vitvi04g04257 | XP_002265747.1 ABSCISIC ACID-INSENSITIVE 5-like protein 2 isoform X1 |
| QTL052 | BERRY_pH | Tier 2 | Vitvi04g04256 | XP_019079568.1 protein argonaute 4A isoform X1 |
| QTL052 | BERRY_pH | Tier 2 | Vitvi04g04255 | CAN67520.1hypothetical protein VITISV_006825 |
| QTL056 | NB_CLUST_PLANT | Tier 2 | Vitvi07g00213 | XP_002272042.2 probable leucine-rich repeat receptor-like protein kinase At1g35710 |
| QTL056 | NB_CLUST_PLANT | Tier 2 | Vitvi07g00209 | XP_002270886.3 receptor-like cytosolic serine/threonine-protein kinase RBK2 |

| QTL ID | Trait | EvidenceTier | GeneID | Description |
|---|---|---|---|---|
| QTL056 | NB_CLUST_PLANT | Tier 2 | Vitvi07g00217 | XP_002271700.1 gibberellin receptor GID1B |
| QTL059 | BUD_DATE | Tier 2 | Vitvi09g00225 | XP_002276544.3 protein EARLY FLOWERING 3 |
| QTL059 | BUD_DATE | Tier 2 | Vitvi09g00227 | XP_010654570.1 transcription factor bHLH87 |
| QTL059 | BUD_DATE | Tier 2 | Vitvi09g01540 | XP_002280177.1 mediator of RNA polymerase II transcription subunit 9 |
| QTL062 | SBER_W_g | Tier 2 | Vitvi11g01186 | XP_002266059.1 auxin efflux carrier component 2 |
| QTL062 | SBER_W_g | Tier 2 | Vitvi11g01641 | XP_010656878.1 probable LRR receptor-like serine/threonine-protein kinase At3g47570 |
| QTL062 | SBER_W_g | Tier 2 | Vitvi11g04327 | XP_010656926.1 probable LRR receptor-like serine/threonine-protein kinase At3g47570 |
| QTL067 | BUD_DATE | Tier 2 | Vitvi16g00859 | XP_010662454.1 protein LATERAL ORGAN BOUNDARIES |
| QTL067 | BUD_DATE | Tier 2 | Vitvi16g00882 | XP_002271141.1 transcription factor bHLH118 |
| QTL067 | BUD_DATE | Tier 2 | Vitvi16g00883 | XP_002271172.2 transcription factor bHLH118 |
| QTL068 | SCLUST_W | Tier 1 | Vitvi16g00886 | XP_002271284.1 PTI1-like tyrosine-protein kinase At3g15890 |
| QTL068 | SCLUST_W | Tier 1 | Vitvi16g00890 | NP_001267911.1gibberellin 20-oxidase |
| QTL068 | SCLUST_W | Tier 1 | Vitvi16g00880 | XP_010662349.1 receptor-like serine/threonine-protein kinase At4g25390 |
| QTL069 | YIELD_OIV_504 | Tier 2 | Vitvi16g04288 | RVW30181.1ABC transporter C family member 10 |
| QTL069 | YIELD_OIV_504 | Tier 2 | Vitvi16g00692 | XP_002266677.1 putative glycerol-3-phosphate transporter 1 |
| QTL069 | YIELD_OIV_504 | Tier 2 | Vitvi16g00710 | XP_002275773.1 protein UNUSUAL FLORAL ORGANS |
| QTL070 | BER_TA_g | Tier 1 | Vitvi16g00860 | XP_010662360.1 K(+) efflux antiporter 5 isoform X2 |
| QTL070 | BER_TA_g | Tier 1 | Vitvi16g00853 | XP_002268932.2 probable aminotransferase ACS12 isoform X1 |
| QTL070 | BER_TA_g | Tier 1 | Vitvi16g00854 | XP_002284088.1 exosome complex component RRP43 |

| QTL ID | Trait | EvidenceTier | GeneID | Description |
|---|---|---|---|---|
| QTL071 | BERRY_pH | Tier 2 | Vitvi16g00936 | XP_019081606.1 ABC transporter B family member 28 |
| QTL071 | BERRY_pH | Tier 2 | Vitvi16g00953 | XP_002274702.1 dihydrodipicolinate reductase-like protein CRR1, chloroplastic |
| QTL071 | BERRY_pH | Tier 2 | Vitvi16g04341 | RVW28433.1hypothetical protein CK203_096180 |
| QTL072 | HARVEST_DATE | Tier 1 | Vitvi16g00974 | XP_003634081.1 homeobox-leucine zipper protein ANTHOCYANINLESS 2 isoform X1 |
| QTL072 | HARVEST_DATE | Tier 1 | Vitvi16g00956 | XP_002279763.2 probable transcriptional regulator SLK2 |
| QTL072 | HARVEST_DATE | Tier 1 | Vitvi16g04342 | RVW28435.1putative transcriptional regulator SLK2 |
| QTL073 | VER_50 | Tier 1 | Vitvi16g00849 | XP_002269179.1 anthocyanidin 3-O-glucosyltransferase 5 |
| QTL073 | VER_50 | Tier 1 | Vitvi16g00882 | XP_002271141.1 transcription factor bHLH118 |
| QTL073 | VER_50 | Tier 1 | Vitvi16g00883 | XP_002271172.2 transcription factor bHLH118 |
| QTL076 | TSS | Tier 2 | Vitvi18g01215 | XP_002265836.1 bidirectional sugar transporter SWEET1 |
| QTL076 | TSS | Tier 2 | Vitvi18g01216 | XP_010664874.2 pentatricopeptide repeat-containing protein At1g08070, chloroplastic |
| QTL076 | TSS | Tier 2 | Vitvi18g01217 | RVW32582.1Pentatricopeptide repeat-containing protein |
| QTL078 | HARVEST_DATE | Tier 2 | Vitvi18g01701 | RVW48844.1Auxin-induced protein X15 |
| QTL078 | HARVEST_DATE | Tier 2 | Vitvi18g01700 | XP_010665090.1 threonylcarbamoyladenosine tRNA methylthiotransferase |
| QTL078 | HARVEST_DATE | Tier 2 | Vitvi18g01699 | XP_002273003.1 FT-interacting protein 1 |

**Supplementary Table 17.** Family-specific allele frequencies and within-family effects at prioritized QTL-GWAS lead SNPs. Abbreviations: QTL, quantitative trait locus; GWAS, genome-wide association study; SNP, single-nucleotide polymorphism.

| QTL ID | Trait | LeadSNP | Family | N genotyped | alternative allele frequency | N phenotyped | within family additive effect | within family p value |
|---|---|---|---|---|---|---|---|---|
| QTL005 | YIELD_PLANT | chr11_4574451 | 42050 | 106 | 0.604 | 104 | -0.2 | 6.07E-5 |
| QTL005 | YIELD_PLANT | chr11_4574451 | 50001 | 71 | 0.268 | 10 | | |
| QTL005 | YIELD_PLANT | chr11_4574451 | 50013 | 58 | 0.302 | 56 | -0.003 | 0.973 |
| QTL005 | YIELD_PLANT | chr11_4574451 | 50015 | 54 | 0.528 | 53 | -0.073 | 0.594 |
| QTL005 | YIELD_PLANT | chr11_4574451 | 50025 | 97 | 0 | 91 | | |
| QTL005 | YIELD_PLANT | chr11_4574451 | 50035 | 186 | 0.487 | 186 | -0.062 | 0.029 |
| QTL006 | YIELD_OIV_504 | chr11_8149404 | 42050 | 106 | 0.307 | 104 | -0.17 | 0.019 |
| QTL006 | YIELD_OIV_504 | chr11_8149404 | 50001 | 71 | 0.303 | 28 | 0.047 | 0.677 |
| QTL006 | YIELD_OIV_504 | chr11_8149404 | 50013 | 58 | 0.534 | 56 | -0.096 | 0.322 |
| QTL006 | YIELD_OIV_504 | chr11_8149404 | 50015 | 54 | 0.25 | 53 | -0.222 | 0.009 |
| QTL006 | YIELD_OIV_504 | chr11_8149404 | 50025 | 97 | 0 | 91 | | |
| QTL006 | YIELD_OIV_504 | chr11_8149404 | 50035 | 186 | 0.468 | 186 | -0.083 | 0.011 |
| QTL007 | FLO_50 | chr14_28852023 | 42050 | 106 | 0.208 | 106 | -0.302 | 2.397E-4 |
| QTL007 | FLO_50 | chr14_28852023 | 50001 | 71 | 0.19 | 59 | -0.117 | 0.009 |
| QTL007 | FLO_50 | chr14_28852023 | 50013 | 58 | 0.276 | 57 | -0.086 | 0.259 |
| QTL007 | FLO_50 | chr14_28852023 | 50015 | 54 | 0 | 54 | | |
| QTL007 | FLO_50 | chr14_28852023 | 50025 | 97 | 0.479 | 91 | -0.183 | 0.002 |
| QTL007 | FLO_50 | chr14_28852023 | 50035 | 186 | 0.462 | 186 | -0.113 | 7.819E-4 |
| QTL008 | VER_50 | chr14_27349067 | 42050 | 106 | 0.5 | 105 | | |
| QTL008 | VER_50 | chr14_27349067 | 50001 | 71 | 0.254 | 60 | 0.832 | 0.523 |
| QTL008 | VER_50 | chr14_27349067 | 50013 | 58 | 0.491 | 57 | -0.472 | 0.9 |
| QTL008 | VER_50 | chr14_27349067 | 50015 | 54 | 0 | 54 | | |
| QTL008 | VER_50 | chr14_27349067 | 50025 | 97 | 0.284 | 91 | 2.924 | 0.009 |
| QTL008 | VER_50 | chr14_27349067 | 50035 | 186 | 0 | 186 | | |

| QTL ID | Trait | LeadSNP | Family | N genotyped | alternative allele frequency | N phenotyped | within family additive effect | within family p value |
|---|---|---|---|---|---|---|---|---|
| QTL011 | VER_50 | chr16_17401013 | 42050 | 106 | 0.717 | 105 | -1.209 | 0.102 |
| QTL011 | VER_50 | chr16_17401013 | 50001 | 71 | 0.683 | 60 | 6.652 | 3.368E-11 |
| QTL011 | VER_50 | chr16_17401013 | 50013 | 58 | 0.784 | 57 | -1.469 | 0.109 |
| QTL011 | VER_50 | chr16_17401013 | 50015 | 54 | 0.194 | 54 | -2.587 | 0.054 |
| QTL011 | VER_50 | chr16_17401013 | 50025 | 97 | 0.464 | 91 | -2.737 | 0.225 |
| QTL011 | VER_50 | chr16_17401013 | 50035 | 186 | 0.435 | 186 | 2.434 | 2.04E-11 |
| QTL014 | SBER_W_g | chr7_23591061 | 42050 | 106 | 0.33 | 104 | -0.055 | 0.113 |
| QTL014 | SBER_W_g | chr7_23591061 | 50001 | 71 | 0.521 | 38 | 0.091 | 0.031 |
| QTL014 | SBER_W_g | chr7_23591061 | 50013 | 58 | 0.664 | 57 | -0.081 | 0.035 |
| QTL014 | SBER_W_g | chr7_23591061 | 50015 | 54 | 0.537 | 53 | -0.025 | 0.559 |
| QTL014 | SBER_W_g | chr7_23591061 | 50025 | 97 | 0 | 91 | | |
| QTL014 | SBER_W_g | chr7_23591061 | 50035 | 186 | 0 | 186 | | |
| QTL015 | MORPHO_OIV_204 | chr14_26574115 | 42050 | 106 | 0.259 | 104 | -0.305 | 0.004 |
| QTL015 | MORPHO_OIV_204 | chr14_26574115 | 50001 | 71 | 0 | 59 | | |
| QTL015 | MORPHO_OIV_204 | chr14_26574115 | 50013 | 58 | 0.207 | 56 | -0.794 | 1.136E-4 |
| QTL015 | MORPHO_OIV_204 | chr14_26574115 | 50015 | 54 | 0.5 | 53 | | |
| QTL015 | MORPHO_OIV_204 | chr14_26574115 | 50025 | 97 | 0 | 91 | | |
| QTL015 | MORPHO_OIV_204 | chr14_26574115 | 50035 | 186 | 0 | 186 | | |
| QTL018 | SCLUST_W | chr14_27028690 | 42050 | 106 | 0.259 | 104 | -0.104 | 0.079 |
| QTL018 | SCLUST_W | chr14_27028690 | 50001 | 71 | 0 | 0 | | |
| QTL018 | SCLUST_W | chr14_27028690 | 50013 | 58 | 0.216 | 56 | -0.38 | 3.91E-4 |
| QTL018 | SCLUST_W | chr14_27028690 | 50015 | 54 | 0.5 | 53 | | |
| QTL018 | SCLUST_W | chr14_27028690 | 50025 | 97 | 0 | 91 | | |
| QTL018 | SCLUST_W | chr14_27028690 | 50035 | 186 | 0 | 186 | | |
| QTL021 | YIELD_PLANT | chr17_9255410 | 42050 | 106 | 0.495 | 104 | 0.037 | 0.412 |

| QTL ID | Trait | LeadSNP | Family | N genotyped | alternative allele frequency | N phenotyped | within family additive effect | within family p value |
|---|---|---|---|---|---|---|---|---|
| QTL021 | YIELD_PLANT | chr17_9255410 | 50001 | 71 | 0.007 | 10 | | |
| QTL021 | YIELD_PLANT | chr17_9255410 | 50013 | 58 | 0.534 | 56 | -0.129 | 0.179 |
| QTL021 | YIELD_PLANT | chr17_9255410 | 50015 | 54 | 0.491 | 53 | -0.403 | 0.265 |
| QTL021 | YIELD_PLANT | chr17_9255410 | 50025 | 97 | 0.17 | 91 | 0.067 | 0.287 |
| QTL021 | YIELD_PLANT | chr17_9255410 | 50035 | 186 | 0.497 | 186 | 0.256 | 0.366 |
| QTL024 | SBER_W_g | chr2_7147735 | 42050 | 106 | 0 | 104 | | |
| QTL024 | SBER_W_g | chr2_7147735 | 50001 | 71 | 0 | 38 | | |
| QTL024 | SBER_W_g | chr2_7147735 | 50013 | 58 | 0 | 57 | | |
| QTL024 | SBER_W_g | chr2_7147735 | 50015 | 54 | 0.287 | 53 | -0.228 | 8.188E-5 |
| QTL024 | SBER_W_g | chr2_7147735 | 50025 | 97 | 0 | 91 | | |
| QTL024 | SBER_W_g | chr2_7147735 | 50035 | 186 | 0 | 186 | | |
| QTL027 | HARVEST_DATE | chr8_5781365 | 42050 | 106 | 0.009 | 105 | -0.006 | 0.972 |
| QTL027 | HARVEST_DATE | chr8_5781365 | 50001 | 71 | 0 | 34 | | |
| QTL027 | HARVEST_DATE | chr8_5781365 | 50013 | 58 | 0 | 57 | | |
| QTL027 | HARVEST_DATE | chr8_5781365 | 50015 | 54 | 0.009 | 53 | -0.73 | 0.015 |
| QTL027 | HARVEST_DATE | chr8_5781365 | 50025 | 97 | 0 | 91 | | |
| QTL027 | HARVEST_DATE | chr8_5781365 | 50035 | 186 | 0.024 | 186 | 0.092 | 0.329 |
| QTL028 | BER_TA_g | chr13_5514868 | 42050 | 106 | 0.505 | 104 | 0.142 | 0.003 |
| QTL028 | BER_TA_g | chr13_5514868 | 50001 | 71 | 0.754 | 60 | 0.13 | 0.163 |
| QTL028 | BER_TA_g | chr13_5514868 | 50013 | 58 | 0.483 | 57 | 0.068 | 0.259 |
| QTL028 | BER_TA_g | chr13_5514868 | 50015 | 54 | 0.25 | 53 | 0.253 | 0.004 |
| QTL028 | BER_TA_g | chr13_5514868 | 50025 | 97 | 0.268 | 91 | 0.035 | 0.627 |
| QTL028 | BER_TA_g | chr13_5514868 | 50035 | 186 | 0.476 | 186 | 0.08 | 0.006 |
| QTL030 | TSS | chr1_23358100 | 42050 | 106 | 0.005 | 104 | -0.178 | 0.81 |
| QTL030 | TSS | chr1_23358100 | 50001 | 71 | 0.261 | 27 | 0.059 | 0.805 |
| QTL030 | TSS | chr1_23358100 | 50013 | 58 | 0.19 | 57 | 0.261 | 0.187 |

| QTL ID | Trait | LeadSNP | Family | N genotyped | alternative allele frequency | N phenotyped | within family additive effect | within family p value |
|---|---|---|---|---|---|---|---|---|
| QTL030 | TSS | chr1_23358100 | 50015 | 54 | 0.259 | 53 | 0.295 | 0.186 |
| QTL030 | TSS | chr1_23358100 | 50025 | 97 | 0 | 91 | | |
| QTL030 | TSS | chr1_23358100 | 50035 | 186 | 0 | 186 | | |
| QTL034 | BER_TA_g | chr14_22281893 | 42050 | 106 | 0.236 | 104 | -0.204 | 0.005 |
| QTL034 | BER_TA_g | chr14_22281893 | 50001 | 71 | 0.176 | 60 | 0.036 | 0.648 |
| QTL034 | BER_TA_g | chr14_22281893 | 50013 | 58 | 0.302 | 57 | -0.138 | 0.067 |
| QTL034 | BER_TA_g | chr14_22281893 | 50015 | 54 | 0 | 53 | | |
| QTL034 | BER_TA_g | chr14_22281893 | 50025 | 97 | 0.634 | 91 | -0.08 | 0.042 |
| QTL034 | BER_TA_g | chr14_22281893 | 50035 | 186 | 0.363 | 186 | -0.077 | 0.002 |
| QTL035 | FLO_50 | chr14_28852023 | 42050 | 106 | 0.208 | 106 | -0.302 | 2.397E-4 |
| QTL035 | FLO_50 | chr14_28852023 | 50001 | 71 | 0.19 | 59 | -0.117 | 0.009 |
| QTL035 | FLO_50 | chr14_28852023 | 50013 | 58 | 0.276 | 57 | -0.086 | 0.259 |
| QTL035 | FLO_50 | chr14_28852023 | 50015 | 54 | 0 | 54 | | |
| QTL035 | FLO_50 | chr14_28852023 | 50025 | 97 | 0.479 | 91 | -0.183 | 0.002 |
| QTL035 | FLO_50 | chr14_28852023 | 50035 | 186 | 0.462 | 186 | -0.113 | 7.819E-4 |
| QTL037 | TSS | chr17_2082982 | 42050 | 106 | 0.495 | 104 | 0.178 | 0.81 |
| QTL037 | TSS | chr17_2082982 | 50001 | 71 | 0.239 | 27 | 0.504 | 0.027 |
| QTL037 | TSS | chr17_2082982 | 50013 | 58 | 0.509 | 57 | -1.024 | 0.163 |
| QTL037 | TSS | chr17_2082982 | 50015 | 54 | 0.5 | 53 | | |
| QTL037 | TSS | chr17_2082982 | 50025 | 97 | 0.778 | 91 | 0.403 | 0.021 |
| QTL037 | TSS | chr17_2082982 | 50035 | 186 | 0.5 | 186 | | |
| QTL039 | TSS | chr18_13434839 | 42050 | 106 | 0.363 | 104 | -0.072 | 0.534 |
| QTL039 | TSS | chr18_13434839 | 50001 | 71 | 0.732 | 27 | -0.307 | 0.188 |
| QTL039 | TSS | chr18_13434839 | 50013 | 58 | 0.328 | 57 | -0.103 | 0.381 |
| QTL039 | TSS | chr18_13434839 | 50015 | 54 | 0.056 | 53 | 0.34 | 0.335 |
| QTL039 | TSS | chr18_13434839 | 50025 | 97 | 0.67 | 91 | -0.491 | 4.882E-5 |

| QTL ID | Trait | LeadSNP | Family | N genotyped | alternative allele frequency | N phenotyped | within family additive effect | within family p value |
|---|---|---|---|---|---|---|---|---|
| QTL039 | TSS | chr18_13434839 | 50035 | 186 | 0.702 | 186 | -0.369 | 2.181E-11 |
| QTL040 | BER_TA_g | chr16_15035128 | 42050 | 106 | 0 | 104 | | |
| QTL040 | BER_TA_g | chr16_15035128 | 50001 | 71 | 0.289 | 60 | -0.269 | 0.004 |
| QTL040 | BER_TA_g | chr16_15035128 | 50013 | 58 | 0 | 57 | | |
| QTL040 | BER_TA_g | chr16_15035128 | 50015 | 54 | 0.287 | 53 | 0.191 | 0.035 |
| QTL040 | BER_TA_g | chr16_15035128 | 50025 | 97 | 0.227 | 91 | -0.314 | 3.039E-6 |
| QTL040 | BER_TA_g | chr16_15035128 | 50035 | 186 | 0.554 | 186 | -0.083 | 0.004 |
| QTL041 | HARVEST_DATE | chr16_15035128 | 42050 | 106 | 0 | 105 | | |
| QTL041 | HARVEST_DATE | chr16_15035128 | 50001 | 71 | 0.289 | 34 | -0.55 | 6.251E-7 |
| QTL041 | HARVEST_DATE | chr16_15035128 | 50013 | 58 | 0 | 57 | | |
| QTL041 | HARVEST_DATE | chr16_15035128 | 50015 | 54 | 0.287 | 53 | 0.016 | 0.852 |
| QTL041 | HARVEST_DATE | chr16_15035128 | 50025 | 97 | 0.227 | 91 | -0.457 | 2.277E-13 |
| QTL041 | HARVEST_DATE | chr16_15035128 | 50035 | 186 | 0.554 | 186 | -0.143 | 2.451E-7 |
| QTL042 | VER_50 | chr16_15035128 | 42050 | 106 | 0 | 105 | | |
| QTL042 | VER_50 | chr16_15035128 | 50001 | 71 | 0.289 | 60 | -8.011 | 1.621E-13 |
| QTL042 | VER_50 | chr16_15035128 | 50013 | 58 | 0 | 57 | | |
| QTL042 | VER_50 | chr16_15035128 | 50015 | 54 | 0.287 | 54 | 0.055 | 0.967 |
| QTL042 | VER_50 | chr16_15035128 | 50025 | 97 | 0.227 | 91 | -7.053 | 6.867E-13 |
| QTL042 | VER_50 | chr16_15035128 | 50035 | 186 | 0.554 | 186 | -2.58 | 5.735E-13 |
| QTL043 | MORPHO_OIV_204 | chr1_5670971 | 42050 | 106 | 0.009 | 104 | 0.284 | 0.465 |
| QTL043 | MORPHO_OIV_204 | chr1_5670971 | 50001 | 71 | 0.746 | 59 | -0.005 | 0.968 |
| QTL043 | MORPHO_OIV_204 | chr1_5670971 | 50013 | 58 | 0.198 | 56 | 0.615 | 0.004 |
| QTL043 | MORPHO_OIV_204 | chr1_5670971 | 50015 | 54 | 0.731 | 53 | 0.012 | 0.92 |
| QTL043 | MORPHO_OIV_204 | chr1_5670971 | 50025 | 97 | 0.428 | 91 | 0.212 | 8.457E-5 |
| QTL043 | MORPHO_OIV_204 | chr1_5670971 | 50035 | 186 | 0.255 | 186 | 0.418 | 3.25E-11 |
| QTL045 | VER_50 | chr2_2851725 | 42050 | 106 | 0.538 | 105 | -0.96 | 0.062 |

| QTL ID | Trait | LeadSNP | Family | N genotyped | alternative allele frequency | N phenotyped | within family additive effect | within family p value |
|---|---|---|---|---|---|---|---|---|
| QTL045 | VER_50 | chr2_2851725 | 50001 | 71 | 0 | 60 | | |
| QTL045 | VER_50 | chr2_2851725 | 50013 | 58 | 0.595 | 57 | -0.523 | 0.526 |
| QTL045 | VER_50 | chr2_2851725 | 50015 | 54 | 0.287 | 54 | -0.261 | 0.835 |
| QTL045 | VER_50 | chr2_2851725 | 50025 | 97 | 0.5 | 91 | | |
| QTL045 | VER_50 | chr2_2851725 | 50035 | 186 | 0.263 | 186 | -1.854 | 3.603E-4 |
| QTL052 | BERRY_pH | chr4_11934711 | 42050 | 106 | 0.269 | 104 | -0.004 | 0.778 |
| QTL052 | BERRY_pH | chr4_11934711 | 50001 | 71 | 0.704 | 60 | 0.003 | 0.836 |
| QTL052 | BERRY_pH | chr4_11934711 | 50013 | 58 | 0.741 | 57 | 0.002 | 0.907 |
| QTL052 | BERRY_pH | chr4_11934711 | 50015 | 54 | 0.444 | 53 | 0.024 | 0.03 |
| QTL052 | BERRY_pH | chr4_11934711 | 50025 | 97 | 0.237 | 91 | 0.009 | 0.381 |
| QTL052 | BERRY_pH | chr4_11934711 | 50035 | 186 | 0.242 | 186 | 0.027 | 7.765E-4 |
| QTL056 | NB_CLUST_PLANT | chr7_2424949 | 42050 | 106 | 0.203 | 104 | 0.259 | 0.032 |
| QTL056 | NB_CLUST_PLANT | chr7_2424949 | 50001 | 71 | 0 | 16 | | |
| QTL056 | NB_CLUST_PLANT | chr7_2424949 | 50013 | 58 | 0.267 | 57 | 0.131 | 0.305 |
| QTL056 | NB_CLUST_PLANT | chr7_2424949 | 50015 | 54 | 0 | 53 | | |
| QTL056 | NB_CLUST_PLANT | chr7_2424949 | 50025 | 97 | 0.237 | 91 | 0.107 | 0.276 |
| QTL056 | NB_CLUST_PLANT | chr7_2424949 | 50035 | 186 | 0.255 | 186 | 0.192 | 4.755E-4 |
| QTL059 | BUD_DATE | chr9_2360990 | 42050 | 106 | 0 | 106 | | |
| QTL059 | BUD_DATE | chr9_2360990 | 50001 | 71 | 0.662 | 60 | -0.097 | 0.683 |
| QTL059 | BUD_DATE | chr9_2360990 | 50013 | 58 | 0.319 | 58 | 1.473 | 0.034 |
| QTL059 | BUD_DATE | chr9_2360990 | 50015 | 54 | 0 | 54 | | |
| QTL059 | BUD_DATE | chr9_2360990 | 50025 | 97 | 0.644 | 91 | 0.739 | 0.051 |
| QTL059 | BUD_DATE | chr9_2360990 | 50035 | 186 | 0.403 | 186 | 1.053 | 2.265E-8 |
| QTL062 | SBER_W_g | chr11_18029044 | 42050 | 106 | 0.321 | 104 | -0.02 | 0.713 |
| QTL062 | SBER_W_g | chr11_18029044 | 50001 | 71 | 0.345 | 38 | -0.237 | 6.938E-4 |
| QTL062 | SBER_W_g | chr11_18029044 | 50013 | 58 | 0.241 | 57 | -0.069 | 0.283 |

| QTL ID | Trait | LeadSNP | Family | N genotyped | alternative allele frequency | N phenotyped | within family additive effect | within family p value |
|---|---|---|---|---|---|---|---|---|
| QTL062 | SBER_W_g | chr11_18029044 | 50015 | 54 | 0.25 | 53 | -0.142 | 0.017 |
| QTL062 | SBER_W_g | chr11_18029044 | 50025 | 97 | 0 | 91 | | |
| QTL062 | SBER_W_g | chr11_18029044 | 50035 | 186 | 0.253 | 186 | -0.109 | 1.31E-5 |
| QTL067 | BUD_DATE | chr16_15107521 | 42050 | 106 | 0.505 | 106 | 0.838 | 0.598 |
| QTL067 | BUD_DATE | chr16_15107521 | 50001 | 71 | 0.275 | 60 | 0.523 | 0.15 |
| QTL067 | BUD_DATE | chr16_15107521 | 50013 | 58 | 0.233 | 58 | 1.228 | 0.265 |
| QTL067 | BUD_DATE | chr16_15107521 | 50015 | 54 | 0.222 | 54 | 0.296 | 0.725 |
| QTL067 | BUD_DATE | chr16_15107521 | 50025 | 97 | 0 | 91 | | |
| QTL067 | BUD_DATE | chr16_15107521 | 50035 | 186 | 0.272 | 186 | 1.093 | 1.26E-5 |
| QTL068 | SCLUST_W | chr16_15542722 | 42050 | 106 | 0.495 | 104 | 0.02 | 0.908 |
| QTL068 | SCLUST_W | chr16_15542722 | 50001 | 71 | 0.289 | 0 | | |
| QTL068 | SCLUST_W | chr16_15542722 | 50013 | 58 | 0.483 | 56 | 0.018 | 0.937 |
| QTL068 | SCLUST_W | chr16_15542722 | 50015 | 54 | 0.37 | 53 | -0.052 | 0.435 |
| QTL068 | SCLUST_W | chr16_15542722 | 50025 | 97 | 0.521 | 91 | -0.138 | 0.304 |
| QTL068 | SCLUST_W | chr16_15542722 | 50035 | 186 | 0.731 | 186 | -0.178 | 0.001 |
| QTL069 | YIELD_OIV_504 | chr16_12507629 | 42050 | 106 | 0 | 104 | | |
| QTL069 | YIELD_OIV_504 | chr16_12507629 | 50001 | 71 | 0.303 | 28 | -0.193 | 0.156 |
| QTL069 | YIELD_OIV_504 | chr16_12507629 | 50013 | 58 | 0 | 56 | | |
| QTL069 | YIELD_OIV_504 | chr16_12507629 | 50015 | 54 | 0 | 53 | | |
| QTL069 | YIELD_OIV_504 | chr16_12507629 | 50025 | 97 | 0.495 | 91 | -0.511 | 0.157 |
| QTL069 | YIELD_OIV_504 | chr16_12507629 | 50035 | 186 | 0 | 186 | | |
| QTL070 | BER_TA_g | chr16_15035128 | 42050 | 106 | 0 | 104 | | |
| QTL070 | BER_TA_g | chr16_15035128 | 50001 | 71 | 0.289 | 60 | -0.269 | 0.004 |
| QTL070 | BER_TA_g | chr16_15035128 | 50013 | 58 | 0 | 57 | | |
| QTL070 | BER_TA_g | chr16_15035128 | 50015 | 54 | 0.287 | 53 | 0.191 | 0.035 |
| QTL070 | BER_TA_g | chr16_15035128 | 50025 | 97 | 0.227 | 91 | -0.314 | 3.039E-6 |

| QTL ID | Trait | LeadSNP | Family | N genotyped | alternative allele frequency | N phenotyped | within family additive effect | within family p value |
|---|---|---|---|---|---|---|---|---|
| QTL070 | BER_TA_g | chr16_15035128 | 50035 | 186 | 0.554 | 186 | -0.083 | 0.004 |
| QTL071 | BERRY_pH | chr16_16539486 | 42050 | 106 | 0.311 | 104 | 0.011 | 0.282 |
| QTL071 | BERRY_pH | chr16_16539486 | 50001 | 71 | 0.408 | 60 | 0.038 | 1.358E-6 |
| QTL071 | BERRY_pH | chr16_16539486 | 50013 | 58 | 0.319 | 57 | 0.008 | 0.387 |
| QTL071 | BERRY_pH | chr16_16539486 | 50015 | 54 | 0.407 | 53 | -0.004 | 0.724 |
| QTL071 | BERRY_pH | chr16_16539486 | 50025 | 97 | 0.51 | 91 | -0.039 | 0.415 |
| QTL071 | BERRY_pH | chr16_16539486 | 50035 | 186 | 0.68 | 186 | 0.02 | 1.738E-4 |
| QTL072 | HARVEST_DATE | chr16_16718042 | 42050 | 106 | 0.278 | 105 | 0.107 | 0.016 |
| QTL072 | HARVEST_DATE | chr16_16718042 | 50001 | 71 | 0.275 | 34 | -0.538 | 7.929E-7 |
| QTL072 | HARVEST_DATE | chr16_16718042 | 50013 | 58 | 0.224 | 57 | 0.121 | 0.02 |
| QTL072 | HARVEST_DATE | chr16_16718042 | 50015 | 54 | 0.315 | 53 | 0.009 | 0.912 |
| QTL072 | HARVEST_DATE | chr16_16718042 | 50025 | 97 | 0.5 | 91 | | |
| QTL072 | HARVEST_DATE | chr16_16718042 | 50035 | 186 | 0.624 | 186 | -0.173 | 4.658E-12 |
| QTL073 | VER_50 | chr16_15035128 | 42050 | 106 | 0 | 105 | | |
| QTL073 | VER_50 | chr16_15035128 | 50001 | 71 | 0.289 | 60 | -8.011 | 1.621E-13 |
| QTL073 | VER_50 | chr16_15035128 | 50013 | 58 | 0 | 57 | | |
| QTL073 | VER_50 | chr16_15035128 | 50015 | 54 | 0.287 | 54 | 0.055 | 0.967 |
| QTL073 | VER_50 | chr16_15035128 | 50025 | 97 | 0.227 | 91 | -7.053 | 6.867E-13 |
| QTL073 | VER_50 | chr16_15035128 | 50035 | 186 | 0.554 | 186 | -2.58 | 5.735E-13 |
| QTL076 | TSS | chr18_13434839 | 42050 | 106 | 0.363 | 104 | -0.072 | 0.534 |
| QTL076 | TSS | chr18_13434839 | 50001 | 71 | 0.732 | 27 | -0.307 | 0.188 |
| QTL076 | TSS | chr18_13434839 | 50013 | 58 | 0.328 | 57 | -0.103 | 0.381 |
| QTL076 | TSS | chr18_13434839 | 50015 | 54 | 0.056 | 53 | 0.34 | 0.335 |
| QTL076 | TSS | chr18_13434839 | 50025 | 97 | 0.67 | 91 | -0.491 | 4.882E-5 |
| QTL076 | TSS | chr18_13434839 | 50035 | 186 | 0.702 | 186 | -0.369 | 2.181E-11 |
| QTL078 | HARVEST_DATE | chr18_23927297 | 42050 | 106 | 0 | 105 | | |

| QTL ID | Trait | LeadSNP | Family | N genotyped | alternative allele frequency | N phenotyped | within family additive effect | within family p value |
|---|---|---|---|---|---|---|---|---|
| QTL078 | HARVEST_DATE | chr18_23927297 | 50001 | 71 | 0 | 34 | | |
| QTL078 | HARVEST_DATE | chr18_23927297 | 50013 | 58 | 0 | 57 | | |
| QTL078 | HARVEST_DATE | chr18_23927297 | 50015 | 54 | 0.231 | 53 | 0.261 | 0.001 |
| QTL078 | HARVEST_DATE | chr18_23927297 | 50025 | 97 | 0.005 | 91 | -0.41 | 0.231 |
| QTL078 | HARVEST_DATE | chr18_23927297 | 50035 | 186 | 0 | 186 | | |

**Supplementary Table 18A.** Three-framework evidence classification for all single-family QTL intervals. Local concordance requires the mpQTL support span to overlap or lie within 500 kb of the GWAS LD-refined interval. A1 denotes locally concordant support from all three frameworks and at least two adjusted GWAS models; A2, all three frameworks and one GWAS; B0, broad but not locally concordant three-framework evidence; B1, single-family QTL plus mpQTL; B2, single-family QTL plus GWAS; and C, single-family QTL or model-specific association only. Single-family QTL intervals and mpQTL support. Abbreviations: QTL, quantitative trait locus; mpQTL, multiple-population QTL mapping; GWAS, genome-wide association study; LD, linkage disequilibrium.

| QTL ID | Pop | Trait | PhysicalChr | QTL low | QTL high | mpQTL overlap | mpQTL leads |
|---|---|---|---|---|---|---|---|
| QTL001 | 42050 | FLO_50 | 2 | 5E+5 | 4,700,000 | FALSE | |
| QTL002 | 42050 | FLO_50 | 7 | 17,500,000 | 24,300,000 | TRUE | chr7_20342111 |
| QTL003 | 42050 | VER_50 | 8 | 14,900,000 | 19,200,000 | FALSE | |
| QTL004 | 42050 | SCLUST_W | 8 | 2,100,000 | 21,200,000 | FALSE | |
| QTL005 | 42050 | YIELD_PLANT | 11 | 3,200,000 | 11,300,000 | FALSE | |
| QTL006 | 42050 | YIELD_OIV_504 | 11 | 3,200,000 | 11,300,000 | FALSE | |
| QTL007 | 42050 | FLO_50 | 14 | 11,300,000 | 29,700,000 | FALSE | |
| QTL008 | 42050 | VER_50 | 14 | 24,300,000 | 28,800,000 | FALSE | |
| QTL009 | 42050 | BERRY_pH | 17 | 5,900,000 | 9,600,000 | FALSE | |
| QTL010 | 42050 | BER_TA_g | 17 | 6,800,000 | 16,600,000 | FALSE | |
| QTL011 | 50001 | VER_50 | 16 | 15,900,000 | 18,200,000 | FALSE | |
| QTL012 | 50001 | BERRY_pH | 16 | 17,100,000 | 19,100,000 | FALSE | |
| QTL013 | 50013 | TSS | 7 | 7E+5 | 3,900,000 | FALSE | |
| QTL014 | 50013 | SBER_W_g | 7 | 4,700,000 | 28,700,000 | FALSE | |
| QTL015 | 50013 | MORPHO_OIV_204 | 14 | 20,700,000 | 30,800,000 | TRUE | chr14_29206455;chr14_28603413;chr14_26846468 |
| QTL016 | 50013 | YIELD_PLANT | 14 | 20,700,000 | 29,600,000 | TRUE | chr14_22600042;chr14_26074715 |
| QTL017 | 50013 | YIELD_OIV_504 | 14 | 19,500,000 | 29,600,000 | TRUE | chr14_22600042 |
| QTL018 | 50013 | SCLUST_W | 14 | 22,700,000 | 29,600,000 | TRUE | chr14_22600042;chr14_26112922 |
| QTL019 | 50013 | TSS | 17 | 4E+6 | 6E+6 | FALSE | |
| QTL020 | 50013 | MORPHO_OIV_204 | 17 | 6,100,000 | 9,200,000 | FALSE | |

| QTL ID | Pop | Trait | PhysicalChr | QTL low | QTL high | mpQTL overlap | mpQTL leads |
|---|---|---|---|---|---|---|---|
| QTL021 | 50013 | YIELD_PLANT | 17 | 6,800,000 | 10,900,000 | FALSE | |
| QTL022 | 50013 | SCLUST_W | 17 | 6,500,000 | 18,800,000 | FALSE | |
| QTL023 | 50013 | YIELD_OIV_504 | 17 | 6,700,000 | 10,900,000 | FALSE | |
| QTL024 | 50015 | SBER_W_g | 2 | 5,300,000 | 7,700,000 | TRUE | chr2_5621690;chr2_5278904 |
| QTL025 | 50015 | YIELD_PLANT | 5 | 1,200,000 | 6,300,000 | FALSE | |
| QTL026 | 50015 | YIELD_OIV_504 | 5 | 7E+5 | 6,300,000 | FALSE | |
| QTL027 | 50015 | HARVEST_DATE | 8 | 2E+5 | 20,700,000 | FALSE | |
| QTL028 | 50015 | BER_TA_g | 13 | 3,900,000 | 7,100,000 | FALSE | |
| QTL029 | 50015 | BERRY_pH | 13 | 4,400,000 | 5,800,000 | FALSE | |
| QTL030 | 50025 | TSS | 1 | 20,400,000 | 23,700,000 | FALSE | |
| QTL031 | 50025 | BUD_DATE | 4 | 8,100,000 | 15,900,000 | FALSE | |
| QTL032 | 50025 | BER_TA_g | 6 | 15,400,000 | 21,200,000 | FALSE | |
| QTL033 | 50025 | BUD_DATE | 7 | 4E+5 | 18,200,000 | TRUE | chr7_4414937 |
| QTL034 | 50025 | BER_TA_g | 14 | 2.1E+7 | 29,800,000 | FALSE | |
| QTL035 | 50025 | FLO_50 | 14 | 17,900,000 | 29,800,000 | FALSE | |
| QTL036 | 50025 | SBER_W_g | 17 | 6,600,000 | 10,300,000 | FALSE | |
| QTL037 | 50025 | TSS | 17 | 1,900,000 | 4,300,000 | FALSE | |
| QTL038 | 50025 | BER_TA_g | 17 | 1,600,000 | 4,900,000 | FALSE | |
| QTL039 | 50025 | TSS | 18 | 12,300,000 | 29,900,000 | FALSE | |
| QTL040 | 50025 | BER_TA_g | 16 | 12,500,000 | 15,200,000 | FALSE | |
| QTL041 | 50025 | HARVEST_DATE | 16 | 1.4E+7 | 15,200,000 | TRUE | chr16_14689931 |
| QTL042 | 50025 | VER_50 | 16 | 14,200,000 | 15,200,000 | TRUE | chr16_14689931 |
| QTL043 | 50035 | MORPHO_OIV_204 | 1 | 2E+6 | 5,800,000 | TRUE | chr1_5742125 |
| QTL044 | 50035 | BUD_DATE | 2 | 2,400,000 | 5,700,000 | FALSE | |

| QTL ID | Pop | Trait | PhysicalChr | QTL low | QTL high | mpQTL overlap | mpQTL leads |
|---|---|---|---|---|---|---|---|
| QTL045 | 50035 | VER_50 | 2 | 2,300,000 | 7,400,000 | FALSE | |
| QTL046 | 50035 | HARVEST_DATE | 2 | 9E+5 | 5,700,000 | FALSE | |
| QTL047 | 50035 | MORPHO_OIV_204 | 2 | 5,200,000 | 6,200,000 | FALSE | |
| QTL048 | 50035 | SCLUST_W | 2 | 5,700,000 | 6,200,000 | FALSE | |
| QTL049 | 50035 | YIELD_OIV_504 | 2 | 5,500,000 | 7,200,000 | FALSE | |
| QTL050 | 50035 | SBER_W_g | 2 | 8,500,000 | 16,400,000 | FALSE | |
| QTL051 | 50035 | NB_CLUST_PLANT | 3 | 4,900,000 | 20,700,000 | FALSE | |
| QTL052 | 50035 | BERRY_pH | 4 | 8,100,000 | 18,900,000 | FALSE | |
| QTL053 | 50035 | BER_TA_g | 4 | 5,200,000 | 18,900,000 | FALSE | |
| QTL054 | 50035 | TSS | 6 | 3,600,000 | 7,300,000 | TRUE | chr6_7263263 |
| QTL055 | 50035 | SBER_W_g | 7 | 2,500,000 | 22,100,000 | FALSE | |
| QTL056 | 50035 | NB_CLUST_PLANT | 7 | 1,800,000 | 5,800,000 | FALSE | |
| QTL057 | 50035 | FLO_50 | 7 | 4,100,000 | 20,600,000 | TRUE | chr7_20342111 |
| QTL058 | 50035 | TSS | 8 | 17,700,000 | 20,300,000 | FALSE | |
| QTL059 | 50035 | BUD_DATE | 9 | 9E+5 | 5,500,000 | FALSE | |
| QTL060 | 50035 | FLO_50 | 9 | 1E+5 | 13,800,000 | FALSE | |
| QTL061 | 50035 | TSS | 9 | 1E+5 | 2,700,000 | TRUE | chr9_2155288 |
| QTL062 | 50035 | SBER_W_g | 11 | 9,600,000 | 19,500,000 | FALSE | |
| QTL063 | 50035 | YIELD_OIV_504 | 11 | 9,600,000 | 19,400,000 | FALSE | |
| QTL064 | 50035 | YIELD_PLANT | 14 | 17,400,000 | 27,400,000 | TRUE | chr14_22600042;chr14_26074715 |
| QTL065 | 50035 | YIELD_OIV_504 | 14 | 1.8E+7 | 28,400,000 | TRUE | chr14_22600042 |
| QTL066 | 50035 | NB_CLUST_PLANT | 14 | 20,600,000 | 30,300,000 | FALSE | |
| QTL067 | 50035 | BUD_DATE | 16 | 2E+6 | 16,800,000 | TRUE | chr16_15360686 |
| QTL068 | 50035 | SCLUST_W | 16 | 14,400,000 | 16,700,000 | FALSE | |

| QTL ID | Pop | Trait | PhysicalChr | QTL low | QTL high | mpQTL overlap | mpQTL leads |
|---|---|---|---|---|---|---|---|
| QTL069 | 50035 | YIELD_OIV_504 | 16 | 8E+5 | 16,700,000 | FALSE | |
| QTL070 | 50035 | BER_TA_g | 16 | 1.5E+7 | 17,300,000 | FALSE | |
| QTL071 | 50035 | BERRY_pH | 16 | 1.5E+7 | 16,800,000 | FALSE | |
| QTL072 | 50035 | HARVEST_DATE | 16 | 15,400,000 | 16,800,000 | FALSE | |
| QTL073 | 50035 | VER_50 | 16 | 1E+5 | 17,600,000 | TRUE | chr16_14689931;chr16_7122451 |
| QTL074 | 50035 | MORPHO_OIV_204 | 16 | 1E+5 | 15,300,000 | FALSE | |
| QTL075 | 50035 | MORPHO_OIV_204 | 18 | 7,900,000 | 34,300,000 | FALSE | |
| QTL076 | 50035 | TSS | 18 | 12,700,000 | 31,100,000 | FALSE | |
| QTL077 | 50015 | FLO_50 | 17 | 5,508,736 | 6,478,604 | TRUE | chr17_5562930;chr17_6431844 |
| QTL078 | 50015 | HARVEST_DATE | 18 | 22,755,801 | 29,601,363 | FALSE | |
| QTL079 | 50015 | NB_CLUST_PLANT | 17 | 3,599 | 4,478,988 | TRUE | chr17_1289030;chr17_2154133 |

**Supplementary Table 18B (continued).** GWAS refinement, local concordance and evidence tier. Abbreviations: QTL, quantitative trait locus; mpQTL, multiple-population QTL mapping; GWAS, genome-wide association study; LD, linkage disequilibrium; bp, base pair.

| QTL ID | Trait | GWAS | LeadSNP | Refined low | Refined high | mpQTL GWAS min gap bp | mpQTL GWAS local concordance | ThreeFrameworkTier |
|---|---|---|---|---|---|---|---|---|
| QTL001 | FLO_50 | FALSE | | | | | FALSE | C |
| QTL002 | FLO_50 | FALSE | | | | | FALSE | B1 |
| QTL003 | VER_50 | FALSE | | | | | FALSE | C |
| QTL004 | SCLUST_W | FALSE | | | | | FALSE | C |
| QTL005 | YIELD_PLANT | TRUE | chr11_4574451 | 4,092,231 | 5,054,770 | | FALSE | B2 |
| QTL006 | YIELD_OIV_504 | TRUE | chr11_8149404 | 7,661,383 | 8,539,030 | | FALSE | B2 |
| QTL007 | FLO_50 | TRUE | chr14_28852023 | 28,355,477 | 29,331,810 | | FALSE | B2 |
| QTL008 | VER_50 | TRUE | chr14_27349067 | 27,037,658 | 27,771,157 | | FALSE | B2 |
| QTL009 | BERRY_pH | FALSE | | | | | FALSE | C |
| QTL010 | BER_TA_g | FALSE | | | | | FALSE | C |
| QTL011 | VER_50 | TRUE | chr16_17401013 | 17,105,497 | 17,883,583 | | FALSE | B2 |
| QTL012 | BERRY_pH | FALSE | | | | | FALSE | C |
| QTL013 | TSS | FALSE | | | | | FALSE | C |
| QTL014 | SBER_W_g | TRUE | chr7_23591061 | 23,186,443 | 24,087,794 | | FALSE | B2 |
| QTL015 | MORPHO_OIV_204 | TRUE | chr14_26574115 | 26,074,252 | 27,062,744 | 0 | TRUE | A2 |
| QTL016 | YIELD_PLANT | FALSE | | | | | FALSE | B1 |
| QTL017 | YIELD_OIV_504 | FALSE | | | | | FALSE | B1 |
| QTL018 | SCLUST_W | TRUE | chr14_27028690 | 26,533,841 | 27,517,943 | 420,443 | TRUE | A2 |
| QTL019 | TSS | FALSE | | | | | FALSE | C |
| QTL020 | MORPHO_OIV_204 | FALSE | | | | | FALSE | C |
| QTL021 | YIELD_PLANT | TRUE | chr17_9255410 | 8,826,475 | 9,748,810 | | FALSE | B2 |
| QTL022 | SCLUST_W | FALSE | | | | | FALSE | C |

| QTL ID | Trait | GWAS | LeadSNP | Refined low | Refined high | mpQTL GWAS min gap bp | mpQTL GWAS local concordance | ThreeFrameworkTier |
|---|---|---|---|---|---|---|---|---|
| QTL023 | YIELD_OIV_504 | FALSE | | | | | FALSE | C |
| QTL024 | SBER_W_g | TRUE | chr2_7147735 | 6,664,706 | 7,621,984 | 1,021,072 | FALSE | B0 |
| QTL025 | YIELD_PLANT | FALSE | | | | | FALSE | C |
| QTL026 | YIELD_OIV_504 | FALSE | | | | | FALSE | C |
| QTL027 | HARVEST_DATE | TRUE | chr8_5781365 | 5,781,221 | 5,781,365 | | FALSE | B2 |
| QTL028 | BER_TA_g | TRUE | chr13_5514868 | 5,025,554 | 5,991,396 | | FALSE | B2 |
| QTL029 | BERRY_pH | FALSE | | | | | FALSE | C |
| QTL030 | TSS | TRUE | chr1_23358100 | 22,906,272 | 23,667,235 | | FALSE | B2 |
| QTL031 | BUD_DATE | FALSE | | | | | FALSE | C |
| QTL032 | BER_TA_g | FALSE | | | | | FALSE | C |
| QTL033 | BUD_DATE | FALSE | | | | | FALSE | B1 |
| QTL034 | BER_TA_g | TRUE | chr14_22281893 | 21,878,626 | 22,760,949 | | FALSE | B2 |
| QTL035 | FLO_50 | TRUE | chr14_28852023 | 28,355,477 | 29,331,810 | | FALSE | B2 |
| QTL036 | SBER_W_g | FALSE | | | | | FALSE | C |
| QTL037 | TSS | TRUE | chr17_2082982 | 1,902,744 | 2,563,496 | | FALSE | B2 |
| QTL038 | BER_TA_g | FALSE | | | | | FALSE | C |
| QTL039 | TSS | TRUE | chr18_13434839 | 12,940,963 | 13,933,924 | | FALSE | B2 |
| QTL040 | BER_TA_g | TRUE | chr16_15035128 | 14,550,221 | 15,178,341 | | FALSE | B2 |
| QTL041 | HARVEST_DATE | TRUE | chr16_15035128 | 14,550,221 | 15,178,341 | 0 | TRUE | A1 |
| QTL042 | VER_50 | TRUE | chr16_15035128 | 14,550,221 | 15,178,341 | 0 | TRUE | A1 |
| QTL043 | MORPHO_OIV_204 | TRUE | chr1_5670971 | 5,228,215 | 5,791,102 | 0 | TRUE | A1 |
| QTL044 | BUD_DATE | FALSE | | | | | FALSE | C |
| QTL045 | VER_50 | TRUE | chr2_2851725 | 2,365,224 | 3,323,554 | | FALSE | B2 |
| QTL046 | HARVEST_DATE | FALSE | | | | | FALSE | C |

| QTL ID | Trait | GWAS | LeadSNP | Refined low | Refined high | mpQTL GWAS min gap bp | mpQTL GWAS local concordance | ThreeFrameworkTier |
|---|---|---|---|---|---|---|---|---|
| QTL047 | MORPHO_OIV_204 | FALSE | | | | | FALSE | C |
| QTL048 | SCLUST_W | FALSE | | | | | FALSE | C |
| QTL049 | YIELD_OIV_504 | FALSE | | | | | FALSE | C |
| QTL050 | SBER_W_g | FALSE | | | | | FALSE | C |
| QTL051 | NB_CLUST_PLANT | FALSE | | | | | FALSE | C |
| QTL052 | BERRY_pH | TRUE | chr4_11934711 | 11,886,067 | 11,947,044 | | FALSE | B2 |
| QTL053 | BER_TA_g | FALSE | | | | | FALSE | C |
| QTL054 | TSS | FALSE | | | | | FALSE | B1 |
| QTL055 | SBER_W_g | FALSE | | | | | FALSE | C |
| QTL056 | NB_CLUST_PLANT | TRUE | chr7_2424949 | 1,951,931 | 2,919,289 | | FALSE | B2 |
| QTL057 | FLO_50 | FALSE | | | | | FALSE | B1 |
| QTL058 | TSS | FALSE | | | | | FALSE | C |
| QTL059 | BUD_DATE | TRUE | chr9_2360990 | 1,865,209 | 2,793,943 | | FALSE | B2 |
| QTL060 | FLO_50 | FALSE | | | | | FALSE | C |
| QTL061 | TSS | FALSE | | | | | FALSE | B1 |
| QTL062 | SBER_W_g | TRUE | chr11_18029044 | 17,557,869 | 18,508,471 | | FALSE | B2 |
| QTL063 | YIELD_OIV_504 | FALSE | | | | | FALSE | C |
| QTL064 | YIELD_PLANT | FALSE | | | | | FALSE | B1 |
| QTL065 | YIELD_OIV_504 | FALSE | | | | | FALSE | B1 |
| QTL066 | NB_CLUST_PLANT | FALSE | | | | | FALSE | C |
| QTL067 | BUD_DATE | TRUE | chr16_15107521 | 14,684,010 | 15,591,633 | 0 | TRUE | A2 |
| QTL068 | SCLUST_W | TRUE | chr16_15542722 | 15,073,646 | 15,946,453 | | FALSE | B2 |
| QTL069 | YIELD_OIV_504 | TRUE | chr16_12507629 | 12,231,464 | 12,885,210 | | FALSE | B2 |
| QTL070 | BER_TA_g | TRUE | chr16_15035128 | 15,034,226 | 15,487,571 | | FALSE | B2 |

| QTL ID | Trait | GWAS | LeadSNP | Refined low | Refined high | mpQTL GWAS min gap bp | mpQTL GWAS local concordance | ThreeFrameworkTier |
|---|---|---|---|---|---|---|---|---|
| QTL071 | BERRY_pH | TRUE | chr16_16539486 | 16,191,685 | 16,792,671 | | FALSE | B2 |
| QTL072 | HARVEST_DATE | TRUE | chr16_16718042 | 16,221,989 | 16,772,451 | | FALSE | B2 |
| QTL073 | VER_50 | TRUE | chr16_15035128 | 14,550,221 | 15,487,571 | 0 | TRUE | A1 |
| QTL074 | MORPHO_OIV_204 | FALSE | | | | | FALSE | C |
| QTL075 | MORPHO_OIV_204 | FALSE | | | | | FALSE | C |
| QTL076 | TSS | TRUE | chr18_13434839 | 12,940,963 | 13,933,924 | | FALSE | B2 |
| QTL077 | FLO_50 | FALSE | | | | | FALSE | B1 |
| QTL078 | HARVEST_DATE | TRUE | chr18_23927297 | 23,784,266 | 24,403,024 | | FALSE | B2 |
| QTL079 | NB_CLUST_PLANT | FALSE | | | | | FALSE | B1 |

**Supplementary Table 19.** Trait-by-contrast summary of exploratory non-additive discovery signals, K+LD1-LD5 sensitivity-retained candidates, and overlap with additive GWAS. Abbreviations: K, genomic kinship; DAPC, discriminant analysis of principal components; LD1-LD5, the first five DAPC linear discriminant axes; GWAS, genome-wide association study.

| Trait | Contrast | K DAPC retest hits | exact additive overlap hits | local250kb additive overlap hits | unique chromosomes | ancestry only Bonferroni hits |
|---|---|---|---|---|---|---|
| BERRY_pH | dominant | 2 | 0 | 1 | 2 | 13 |
| BERRY_pH | overdominant | 1 | 0 | 0 | 1 | 8 |
| BERRY_pH | recessive | 1 | 0 | 0 | 1 | 5 |
| BER_TA_g | dominant | 0 | 0 | 0 | 0 | 10 |
| BER_TA_g | overdominant | 2 | 0 | 1 | 2 | 8 |
| BER_TA_g | recessive | 0 | 0 | 0 | 0 | 4 |
| BUD_DATE | dominant | 1 | 0 | 0 | 1 | 14 |
| BUD_DATE | overdominant | 2 | 1 | 1 | 2 | 53 |
| BUD_DATE | recessive | 1 | 0 | 0 | 1 | 15 |
| FLO_50 | dominant | 5 | 1 | 1 | 4 | 18 |
| FLO_50 | overdominant | 5 | 1 | 1 | 4 | 18 |
| FLO_50 | recessive | 2 | 0 | 0 | 2 | 13 |
| HARVEST_DATE | dominant | 3 | 3 | 3 | 2 | 33 |
| HARVEST_DATE | overdominant | 4 | 4 | 4 | 2 | 26 |
| HARVEST_DATE | recessive | 8 | 2 | 3 | 6 | 32 |
| MORPHO_OIV_204 | dominant | 0 | 0 | 0 | 0 | 12 |
| MORPHO_OIV_204 | overdominant | 1 | 0 | 1 | 1 | 11 |
| MORPHO_OIV_204 | recessive | 0 | 0 | 0 | 0 | 9 |
| NB_CLUST_PLANT | dominant | 0 | 0 | 0 | 0 | 6 |
| NB_CLUST_PLANT | overdominant | 3 | 0 | 0 | 3 | 14 |
| NB_CLUST_PLANT | recessive | 4 | 0 | 0 | 3 | 12 |
| SBER_W_g | dominant | 0 | 0 | 0 | 0 | 17 |
| SBER_W_g | overdominant | 0 | 0 | 0 | 0 | 17 |
| SBER_W_g | recessive | 1 | 0 | 0 | 1 | 23 |

| Trait | Contrast | K DAPC retest hits | exact additive overlap hits | local250kb additive overlap hits | unique chromosomes | ancestry only Bonferroni hits |
|---|---|---|---|---|---|---|
| SCLUST_W | dominant | 2 | 0 | 0 | 2 | 13 |
| SCLUST_W | overdominant | 1 | 0 | 0 | 1 | 12 |
| SCLUST_W | recessive | 0 | 0 | 0 | 0 | 5 |
| TSS | dominant | 0 | 0 | 0 | 0 | 23 |
| TSS | overdominant | 0 | 0 | 0 | 0 | 9 |
| TSS | recessive | 0 | 0 | 0 | 0 | 20 |
| VER_50 | dominant | 2 | 2 | 2 | 1 | 31 |
| VER_50 | overdominant | 4 | 2 | 3 | 2 | 20 |
| VER_50 | recessive | 4 | 2 | 4 | 1 | 25 |
| YIELD_OIV_504 | dominant | 2 | 0 | 0 | 2 | 11 |
| YIELD_OIV_504 | overdominant | 0 | 0 | 0 | 0 | 9 |
| YIELD_OIV_504 | recessive | 1 | 0 | 0 | 1 | 5 |
| YIELD_PLANT | dominant | 2 | 0 | 0 | 2 | 6 |
| YIELD_PLANT | overdominant | 5 | 0 | 0 | 5 | 19 |
| YIELD_PLANT | recessive | 5 | 0 | 0 | 3 | 7 |

**Supplementary Table 20A.** Trait-level environment-GWAS and metaGE summary. Fixed-effect, random-effect, local-score and climate-regression evidence are reported separately. Meta-analysis inputs and fixed- versus random-effect evidence. Abbreviations: GWAS, genome-wide association study; metaGE, meta-analysis of genotype-environment association results; FDR, false discovery rate. Local-score analyses used ξ = 3 for all traits.

| Trait | N environments | N common markers | fixed effect FDR 5% markers | random effect FDR 5% markers | fixed effect local score zones | random effect local score zones | correlation regularization threshold |
|---|---|---|---|---|---|---|---|
| BER_TA_g | 26 | 44,040 | 0 | 1,115 | 9 | 95 | 0.8 |
| BUD_DATE | 21 | 42,521 | 0 | 279 | 5 | 72 | 0.8 |
| FLO_50 | 20 | 42,801 | 0 | 324 | 2 | 62 | 0.8 |
| MORPHO_OIV_204 | 20 | 66,755 | 0 | 457 | 2 | 74 | 0.8 |
| SCLUST_W | 26 | 43,837 | 0 | 33 | 1 | 34 | 0.8 |
| VER_50 | 28 | 45,305 | 0 | 1,321 | 4 | 85 | 0.8 |
| YIELD_PLANT | 20 | 65,922 | 0 | 13 | 0 | 17 | 0.8 |
| BERRY_pH | 25 | 44,021 | 0 | 81 | 0 | 40 | 0.8 |
| HARVEST_DATE | 24 | 44,184 | 0 | 238 | 4 | 63 | 0.8 |
| NB_CLUST_PLANT | 20 | 67,401 | 0 | 18 | 0 | 18 | 0.8 |
| SBER_W_g | 24 | 44,346 | 0 | 690 | 1 | 82 | 0.8 |
| TSS | 25 | 45,306 | 0 | 36 | 2 | 30 | 0.8 |
| YIELD_OIV_504 | 24 | 43,767 | 0 | 24 | 0 | 20 | 0.8 |

**Supplementary Table 20B (continued).** Marker-climate tests passing 5% FDR in separate univariate regressions. Abbreviations: FDR, false discovery rate.

| Trait | climate FDR 5% mean Tmin C | climate FDR 5% mean Tmax C | climate FDR 5% mean Tmean C | climate FDR 5% annual rainfall mm | climate FDR 5% mean ET0 mm d | climate FDR 5% mean radiation MJ m2 d |
|---|---|---|---|---|---|---|
| BER_TA_g | 0 | 0 | 0 | 0 | 0 | 0 |
| BUD_DATE | 0 | 0 | 2 | 0 | 0 | 0 |
| FLO_50 | 411 | 4 | 0 | 55 | 0 | 0 |
| MORPHO_OIV_204 | 0 | 0 | 0 | 0 | 0 | 0 |
| SCLUST_W | 0 | 0 | 0 | 0 | 0 | 0 |
| VER_50 | 7 | 0 | 10 | 0 | 3 | 0 |

| Trait | climate FDR 5% mean Tmin C | climate FDR 5% mean Tmax C | climate FDR 5% mean Tmean C | climate FDR 5% annual rainfall mm | climate FDR 5% mean ET0 mm d | climate FDR 5% mean radiation MJ m2 d |
|---|---|---|---|---|---|---|
| YIELD_PLANT | 0 | 0 | 0 | 0 | 0 | 0 |
| BERRY_pH | 0 | 1 | 0 | 1 | 0 | 0 |
| HARVEST_DATE | 3 | 1 | 1 | 1 | 3 | 0 |
| NB_CLUST_PLANT | 0 | 0 | 0 | 0 | 0 | 0 |
| SBER_W_g | 0 | 0 | 0 | 0 | 0 | 0 |
| TSS | 0 | 1 | 0 | 0 | 0 | 0 |
| YIELD_OIV_504 | 0 | 0 | 0 | 0 | 0 | 0 |

**Supplementary Table 21.** Cross-trait marker tests selected by the QCH composite alternative of association with at least two traits using the 5% QCH FDR procedure, which orders posterior local-FDR estimates. Trait-level entries are descriptive Benjamini-Hochberg-adjusted P values from the corresponding ancestry-adjusted BLINK scans and did not select the QCH markers. Abbreviations: QCH, Query Composite Hypotheses; FDR, false discovery rate; BH, Benjamini-Hochberg; BLINK, Bayesian-information and Linkage-disequilibrium Iteratively Nested Keyway.

| Chr | Position (bp) | Marker | QCH P | Local FDR | Trait-level FDR count | Traits with BH <= 0.05 |
|---|---|---|---|---|---|---|
| 2 | 1,266,852 | chr2_1266852 | 8.791E-5 | 6.943E-2 | 0 | |
| 5 | 53,402 | chr5_53402 | 5.678E-5 | 5.395E-2 | 0 | |
| 5 | 61,614 | chr5_61614 | 8.511E-5 | 6.884E-2 | 0 | |
| 5 | 148,010 | chr5_148010 | 6.017E-5 | 5.604E-2 | 0 | |
| 5 | 148,076 | chr5_148076 | 8.441E-5 | 6.867E-2 | 0 | |
| 5 | 148,089 | chr5_148089 | 4.125E-5 | 4.793E-2 | 0 | |
| 5 | 148,098 | chr5_148098 | 6.017E-5 | 5.604E-2 | 0 | |
| 5 | 206,661 | chr5_206661 | 8.511E-5 | 6.884E-2 | 0 | |
| 5 | 310,550 | chr5_310550 | 3.839E-5 | 4.597E-2 | 0 | |
| 5 | 354,794 | chr5_354794 | 4.722E-5 | 5.059E-2 | 0 | |
| 5 | 430,004 | chr5_430004 | 1.957E-5 | 3.568E-2 | 0 | |
| 5 | 432,535 | chr5_432535 | 5.246E-5 | 5.254E-2 | 0 | |
| 5 | 469,112 | chr5_469112 | 6.889E-5 | 5.913E-2 | 0 | |
| 5 | 472,735 | chr5_472735 | 1.638E-5 | 3.43E-2 | 0 | |
| 5 | 505,872 | chr5_505872 | 4.369E-5 | 4.812E-2 | 0 | |
| 5 | 510,952 | chr5_510952 | 1.029E-4 | 7.77E-2 | 0 | |
| 5 | 511,140 | chr5_511140 | 4.418E-5 | 4.858E-2 | 0 | |
| 5 | 514,500 | chr5_514500 | 4.982E-5 | 5.147E-2 | 0 | |
| 5 | 514,529 | chr5_514529 | 4.982E-5 | 5.147E-2 | 0 | |
| 5 | 514,586 | chr5_514586 | 4.982E-5 | 5.147E-2 | 0 | |
| 5 | 525,403 | chr5_525403 | 1.152E-4 | 8.401E-2 | 0 | |
| 5 | 532,830 | chr5_532830 | 3.207E-5 | 4.064E-2 | 0 | |
| 5 | 532,869 | chr5_532869 | 1.222E-4 | 8.674E-2 | 0 | |
| 5 | 596,214 | chr5_596214 | 4.59E-6 | 2.197E-2 | 0 | |
| 5 | 602,088 | chr5_602088 | 9.077E-5 | 7.083E-2 | 0 | |
| 5 | 731,098 | chr5_731098 | 2.101E-6 | 1.82E-2 | 0 | |
| 5 | 745,008 | chr5_745008 | 6.949E-5 | 5.94E-2 | 0 | |
| 5 | 763,393 | chr5_763393 | 3.337E-5 | 4.315E-2 | 0 | |
| 5 | 770,693 | chr5_770693 | 9.005E-5 | 7.037E-2 | 0 | |

| Chr | Position (bp) | Marker | QCH P | Local FDR | Trait-level FDR count | Traits with BH <= 0.05 |
|---|---|---|---|---|---|---|
| 5 | 782,507 | chr5_782507 | 4.174E-5 | 4.802E-2 | 0 | |
| 5 | 793,081 | chr5_793081 | 4.619E-5 | 5.004E-2 | 0 | |
| 5 | 813,871 | chr5_813871 | 2.178E-5 | 3.708E-2 | 0 | |
| 5 | 816,270 | chr5_816270 | 9.088E-6 | 2.966E-2 | 0 | |
| 5 | 816,309 | chr5_816309 | 4.174E-5 | 4.802E-2 | 0 | |
| 5 | 816,377 | chr5_816377 | 2.288E-6 | 1.844E-2 | 0 | |
| 5 | 824,702 | chr5_824702 | 1.501E-5 | 3.337E-2 | 0 | |
| 5 | 834,136 | chr5_834136 | 1.501E-5 | 3.337E-2 | 0 | |
| 5 | 837,242 | chr5_837242 | 5.407E-5 | 5.308E-2 | 0 | |
| 5 | 861,653 | chr5_861653 | 3.654E-5 | 4.508E-2 | 0 | |
| 5 | 861,662 | chr5_861662 | 6.361E-5 | 5.707E-2 | 0 | |
| 5 | 896,765 | chr5_896765 | 1.673E-5 | 3.434E-2 | 0 | |
| 5 | 913,164 | chr5_913164 | 7.382E-5 | 6.156E-2 | 0 | |
| 5 | 913,222 | chr5_913222 | 6.527E-6 | 2.521E-2 | 0 | |
| 5 | 927,437 | chr5_927437 | 1.161E-4 | 8.418E-2 | 0 | |
| 5 | 927,520 | chr5_927520 | 4.671E-5 | 5.049E-2 | 0 | |
| 5 | 970,011 | chr5_970011 | 3.427E-5 | 4.451E-2 | 0 | |
| 5 | 978,913 | chr5_978913 | 3.125E-5 | 4.021E-2 | 0 | |
| 5 | 984,394 | chr5_984394 | 2.762E-5 | 3.926E-2 | 0 | |
| 5 | 986,631 | chr5_986631 | 9.696E-6 | 3.017E-2 | 0 | |
| 5 | 986,691 | chr5_986691 | 9.696E-6 | 3.017E-2 | 0 | |
| 5 | 1,017,384 | chr5_1017384 | 3.929E-6 | 2.155E-2 | 0 | |
| 5 | 1,058,484 | chr5_1058484 | 7.572E-5 | 6.279E-2 | 0 | |
| 5 | 1,060,525 | chr5_1060525 | 3.929E-6 | 2.155E-2 | 0 | |
| 5 | 1,079,665 | chr5_1079665 | 9.696E-6 | 3.017E-2 | 0 | |
| 5 | 1,097,913 | chr5_1097913 | 5.766E-6 | 2.427E-2 | 0 | |
| 5 | 1,119,566 | chr5_1119566 | 8.166E-5 | 6.631E-2 | 0 | |
| 5 | 1,120,981 | chr5_1120981 | 6.8E-6 | 2.691E-2 | 0 | |
| 5 | 1,143,112 | chr5_1143112 | 7.832E-5 | 6.526E-2 | 0 | |
| 5 | 1,143,565 | chr5_1143565 | 2.643E-5 | 3.878E-2 | 0 | |
| 5 | 1,143,630 | chr5_1143630 | 2.216E-5 | 3.727E-2 | 0 | |
| 5 | 1,215,091 | chr5_1215091 | 9.222E-5 | 7.171E-2 | 0 | |
| 5 | 1,217,906 | chr5_1217906 | 9.696E-6 | 3.017E-2 | 0 | |
| 5 | 1,219,467 | chr5_1219467 | 9.696E-6 | 3.017E-2 | 0 | |

| Chr | Position (bp) | Marker | QCH P | Local FDR | Trait-level FDR count | Traits with BH <= 0.05 |
|---|---|---|---|---|---|---|
| 5 | 1,219,726 | chr5_1219726 | 1.663E-7 | 1.066E-2 | 0 | |
| 5 | 1,219,801 | chr5_1219801 | 6.13E-5 | 5.61E-2 | 0 | |
| 5 | 1,225,256 | chr5_1225256 | 1.077E-4 | 8.014E-2 | 0 | |
| 5 | 1,280,592 | chr5_1280592 | 7.071E-5 | 6.063E-2 | 0 | |
| 5 | 1,281,872 | chr5_1281872 | 5.353E-5 | 5.299E-2 | 0 | |
| 5 | 1,281,893 | chr5_1281893 | 6.303E-5 | 5.678E-2 | 0 | |
| 5 | 1,422,159 | chr5_1422159 | 2.876E-6 | 1.969E-2 | 0 | |
| 5 | 1,436,402 | chr5_1436402 | 1.603E-5 | 3.383E-2 | 0 | |
| 5 | 1,480,443 | chr5_1480443 | 4.774E-5 | 5.111E-2 | 0 | |
| 5 | 1,487,918 | chr5_1487918 | 1.435E-5 | 3.209E-2 | 0 | |
| 5 | 1,487,923 | chr5_1487923 | 2.408E-5 | 3.853E-2 | 0 | |
| 5 | 1,539,175 | chr5_1539175 | 5.569E-5 | 5.344E-2 | 0 | |
| 5 | 1,539,245 | chr5_1539245 | 5.14E-5 | 5.199E-2 | 0 | |
| 5 | 1,539,277 | chr5_1539277 | 5.903E-5 | 5.547E-2 | 0 | |
| 5 | 1,540,475 | chr5_1540475 | 7.92E-6 | 2.798E-2 | 0 | |
| 5 | 1,540,477 | chr5_1540477 | 7.92E-6 | 2.798E-2 | 0 | |
| 5 | 1,540,573 | chr5_1540573 | 3.002E-5 | 4.004E-2 | 0 | |
| 5 | 1,540,582 | chr5_1540582 | 2.762E-5 | 3.926E-2 | 0 | |
| 5 | 1,540,772 | chr5_1540772 | 2.254E-5 | 3.728E-2 | 0 | |
| 5 | 1,540,829 | chr5_1540829 | 2.447E-5 | 3.855E-2 | 0 | |
| 5 | 1,540,838 | chr5_1540838 | 2.447E-5 | 3.855E-2 | 0 | |
| 5 | 1,540,892 | chr5_1540892 | 2.762E-5 | 3.926E-2 | 0 | |
| 5 | 1,540,937 | chr5_1540937 | 1.708E-5 | 3.44E-2 | 0 | |
| 5 | 1,541,071 | chr5_1541071 | 1.993E-5 | 3.569E-2 | 0 | |
| 5 | 1,541,172 | chr5_1541172 | 4.569E-5 | 5.001E-2 | 0 | |
| 5 | 1,541,332 | chr5_1541332 | 8.432E-7 | 1.724E-2 | 0 | |
| 5 | 1,543,492 | chr5_1543492 | 2.525E-5 | 3.864E-2 | 0 | |
| 5 | 1,564,043 | chr5_1564043 | 1.069E-4 | 7.994E-2 | 0 | |
| 5 | 1,637,232 | chr5_1637232 | 7.965E-5 | 6.56E-2 | 0 | |
| 5 | 1,637,756 | chr5_1637756 | 6.419E-5 | 5.719E-2 | 0 | |
| 5 | 1,713,024 | chr5_1713024 | 6.535E-5 | 5.744E-2 | 0 | |
| 5 | 1,713,278 | chr5_1713278 | 1.053E-4 | 7.864E-2 | 0 | |
| 5 | 1,952,034 | chr5_1952034 | 1.338E-4 | 9.024E-2 | 0 | |
| 5 | 1,952,452 | chr5_1952452 | 1.339E-5 | 3.109E-2 | 0 | |

| Chr | Position (bp) | Marker | QCH P | Local FDR | Trait-level FDR count | Traits with BH <= 0.05 |
|---|---|---|---|---|---|---|
| 5 | 1,984,057 | chr5_1984057 | 2.103E-5 | 3.642E-2 | 0 | |
| 5 | 1,990,972 | chr5_1990972 | 2.722E-5 | 3.904E-2 | 0 | |
| 5 | 2,005,655 | chr5_2005655 | 9.696E-6 | 3.017E-2 | 0 | |
| 5 | 2,033,510 | chr5_2033510 | 8.787E-6 | 2.893E-2 | 0 | |
| 5 | 2,033,601 | chr5_2033601 | 9.671E-5 | 7.491E-2 | 0 | |
| 5 | 2,041,299 | chr5_2041299 | 1.045E-4 | 7.85E-2 | 0 | |
| 5 | 2,081,872 | chr5_2081872 | 5.193E-5 | 5.221E-2 | 0 | |
| 5 | 2,091,324 | chr5_2091324 | 1.307E-5 | 3.093E-2 | 0 | |
| 5 | 2,110,797 | chr5_2110797 | 1.085E-4 | 8.031E-2 | 0 | |
| 5 | 2,151,197 | chr5_2151197 | 1.743E-5 | 3.454E-2 | 0 | |
| 5 | 2,187,784 | chr5_2187784 | 2.03E-5 | 3.595E-2 | 0 | |
| 5 | 2,191,323 | chr5_2191323 | 3.427E-5 | 4.451E-2 | 0 | |
| 5 | 2,191,341 | chr5_2191341 | 7.319E-5 | 6.15E-2 | 0 | |
| 5 | 2,192,309 | chr5_2192309 | 3.427E-5 | 4.451E-2 | 0 | |
| 5 | 2,482,576 | chr5_2482576 | 1.467E-5 | 3.217E-2 | 0 | |
| 5 | 2,490,197 | chr5_2490197 | 1.266E-4 | 8.823E-2 | 0 | |
| 5 | 2,490,307 | chr5_2490307 | 8.934E-5 | 7.026E-2 | 0 | |
| 5 | 2,493,746 | chr5_2493746 | 7.133E-5 | 6.075E-2 | 0 | |
| 5 | 2,495,913 | chr5_2495913 | 3.084E-5 | 4.012E-2 | 0 | |
| 5 | 2,496,812 | chr5_2496812 | 2.962E-5 | 3.959E-2 | 0 | |
| 5 | 2,496,854 | chr5_2496854 | 9.824E-5 | 7.524E-2 | 0 | |
| 5 | 2,497,079 | chr5_2497079 | 1.178E-4 | 8.5E-2 | 0 | |
| 5 | 2,497,189 | chr5_2497189 | 3.382E-5 | 4.433E-2 | 0 | |
| 5 | 2,497,761 | chr5_2497761 | 2.683E-5 | 3.901E-2 | 0 | |
| 5 | 2,497,793 | chr5_2497793 | 6.272E-6 | 2.504E-2 | 0 | |
| 5 | 2,497,822 | chr5_2497822 | 9.444E-5 | 7.399E-2 | 0 | |
| 5 | 2,501,423 | chr5_2501423 | 1.195E-4 | 8.6E-2 | 0 | |
| 5 | 2,501,481 | chr5_2501481 | 1.293E-4 | 8.853E-2 | 0 | |
| 5 | 2,506,715 | chr5_2506715 | 7.01E-5 | 5.974E-2 | 0 | |
| 5 | 2,506,976 | chr5_2506976 | 9.748E-5 | 7.518E-2 | 0 | |
| 5 | 2,522,382 | chr5_2522382 | 3.7E-5 | 4.541E-2 | 0 | |
| 5 | 2,522,874 | chr5_2522874 | 3.077E-6 | 1.983E-2 | 0 | |
| 5 | 2,522,898 | chr5_2522898 | 3.886E-5 | 4.613E-2 | 0 | |
| 5 | 2,522,923 | chr5_2522923 | 3.746E-5 | 4.563E-2 | 0 | |

| Chr | Position (bp) | Marker | QCH P | Local FDR | Trait-level FDR count | Traits with BH <= 0.05 |
|---|---|---|---|---|---|---|
| 5 | 2,548,438 | chr5_2548438 | 4.815E-6 | 2.222E-2 | 0 | |
| 5 | 2,556,540 | chr5_2556540 | 2.676E-6 | 1.947E-2 | 0 | |
| 5 | 2,561,676 | chr5_2561676 | 1.127E-4 | 8.22E-2 | 0 | |
| 5 | 2,616,134 | chr5_2616134 | 9.696E-6 | 3.017E-2 | 0 | |
| 5 | 2,616,182 | chr5_2616182 | 8.302E-5 | 6.742E-2 | 0 | |
| 5 | 2,626,033 | chr5_2626033 | 6.477E-5 | 5.721E-2 | 0 | |
| 5 | 2,651,789 | chr5_2651789 | 5.624E-5 | 5.387E-2 | 0 | |
| 5 | 2,654,861 | chr5_2654861 | 3.166E-5 | 4.035E-2 | 0 | |
| 5 | 2,654,971 | chr5_2654971 | 1.32E-4 | 8.998E-2 | 0 | |
| 5 | 2,656,188 | chr5_2656188 | 5.299E-5 | 5.255E-2 | 0 | |
| 5 | 2,660,364 | chr5_2660364 | 1.885E-5 | 3.561E-2 | 0 | |
| 5 | 2,665,314 | chr5_2665314 | 7.444E-5 | 6.17E-2 | 0 | |
| 5 | 2,684,082 | chr5_2684082 | 3.427E-5 | 4.451E-2 | 0 | |
| 5 | 2,691,605 | chr5_2691605 | 1.569E-5 | 3.343E-2 | 0 | |
| 5 | 2,691,716 | chr5_2691716 | 1.248E-4 | 8.76E-2 | 0 | |
| 5 | 2,691,870 | chr5_2691870 | 4.947E-7 | 1.623E-2 | 0 | |
| 5 | 2,696,389 | chr5_2696389 | 6.245E-5 | 5.656E-2 | 0 | |
| 5 | 2,696,565 | chr5_2696565 | 1.204E-4 | 8.651E-2 | 0 | |
| 5 | 2,723,856 | chr5_2723856 | 5.791E-5 | 5.547E-2 | 0 | |
| 5 | 2,723,861 | chr5_2723861 | 5.791E-5 | 5.547E-2 | 0 | |
| 5 | 2,724,755 | chr5_2724755 | 7.636E-5 | 6.32E-2 | 0 | |
| 5 | 2,725,797 | chr5_2725797 | 3.301E-7 | 1.615E-2 | 0 | |
| 5 | 2,726,160 | chr5_2726160 | 1.239E-4 | 8.738E-2 | 0 | |
| 5 | 2,730,064 | chr5_2730064 | 1.102E-4 | 8.139E-2 | 0 | |
| 5 | 2,738,297 | chr5_2738297 | 4.468E-5 | 4.865E-2 | 0 | |
| 5 | 2,738,321 | chr5_2738321 | 3.792E-5 | 4.566E-2 | 0 | |
| 5 | 2,742,074 | chr5_2742074 | 2.292E-5 | 3.743E-2 | 0 | |
| 5 | 2,748,431 | chr5_2748431 | 5.461E-5 | 5.322E-2 | 0 | |
| 5 | 2,762,620 | chr5_2762620 | 3.981E-5 | 4.691E-2 | 0 | |
| 5 | 2,762,662 | chr5_2762662 | 9.52E-5 | 7.423E-2 | 0 | |
| 5 | 2,770,379 | chr5_2770379 | 9.696E-6 | 3.017E-2 | 0 | |
| 5 | 2,863,018 | chr5_2863018 | 8.099E-5 | 6.606E-2 | 0 | |
| 5 | 2,864,486 | chr5_2864486 | 8.372E-5 | 6.847E-2 | 0 | |
| 5 | 2,900,476 | chr5_2900476 | 1.186E-4 | 8.568E-2 | 0 | |

| Chr | Position (bp) | Marker | QCH P | Local FDR | Trait-level FDR count | Traits with BH <= 0.05 |
|---|---|---|---|---|---|---|
| 5 | 2,903,454 | chr5_2903454 | 4.271E-5 | 4.805E-2 | 0 | |
| 5 | 2,934,173 | chr5_2934173 | 3.25E-5 | 4.222E-2 | 0 | |
| 5 | 2,934,208 | chr5_2934208 | 4.028E-5 | 4.721E-2 | 0 | |
| 5 | 2,940,215 | chr5_2940215 | 9.696E-6 | 3.017E-2 | 0 | |
| 5 | 2,946,624 | chr5_2946624 | 5.735E-5 | 5.539E-2 | 0 | |
| 5 | 2,961,815 | chr5_2961815 | 4.878E-5 | 5.147E-2 | 0 | |
| 5 | 2,961,843 | chr5_2961843 | 4.878E-5 | 5.147E-2 | 0 | |
| 5 | 3,000,999 | chr5_3000999 | 9.696E-6 | 3.017E-2 | 0 | |
| 5 | 3,027,554 | chr5_3027554 | 5.52E-6 | 2.365E-2 | 0 | |
| 5 | 3,085,438 | chr5_3085438 | 6.829E-5 | 5.881E-2 | 0 | |
| 5 | 3,104,308 | chr5_3104308 | 4.077E-5 | 4.738E-2 | 0 | |
| 5 | 3,104,324 | chr5_3104324 | 2.762E-5 | 3.926E-2 | 0 | |
| 5 | 3,104,607 | chr5_3104607 | 3.043E-5 | 4.007E-2 | 0 | |
| 5 | 3,104,661 | chr5_3104661 | 1.849E-5 | 3.483E-2 | 0 | |
| 5 | 3,107,961 | chr5_3107961 | 1.022E-6 | 1.758E-2 | 0 | |
| 5 | 3,119,699 | chr5_3119699 | 6.017E-6 | 2.479E-2 | 0 | |
| 5 | 3,149,984 | chr5_3149984 | 6.593E-5 | 5.75E-2 | 0 | |
| 5 | 3,158,245 | chr5_3158245 | 1.402E-5 | 3.134E-2 | 0 | |
| 5 | 3,161,774 | chr5_3161774 | 2.762E-5 | 3.926E-2 | 0 | |
| 5 | 3,176,000 | chr5_3176000 | 8.862E-5 | 7.005E-2 | 0 | |
| 5 | 3,287,131 | chr5_3287131 | 3.287E-6 | 2.077E-2 | 0 | |
| 5 | 3,487,511 | chr5_3487511 | 7.508E-5 | 6.274E-2 | 0 | |
| 5 | 3,527,559 | chr5_3527559 | 2.604E-5 | 3.875E-2 | 0 | |
| 5 | 3,546,226 | chr5_3546226 | 1.736E-6 | 1.774E-2 | 0 | |
| 5 | 3,546,240 | chr5_3546240 | 1.736E-6 | 1.774E-2 | 0 | |
| 5 | 3,566,968 | chr5_3566968 | 7.257E-5 | 6.112E-2 | 0 | |
| 5 | 3,591,657 | chr5_3591657 | 9.696E-6 | 3.017E-2 | 0 | |
| 5 | 3,608,708 | chr5_3608708 | 1.275E-4 | 8.844E-2 | 0 | |
| 5 | 3,608,711 | chr5_3608711 | 1.275E-4 | 8.844E-2 | 0 | |
| 5 | 3,622,274 | chr5_3622274 | 2.33E-5 | 3.808E-2 | 0 | |
| 5 | 3,626,149 | chr5_3626149 | 9.39E-6 | 2.974E-2 | 0 | |
| 5 | 3,632,850 | chr5_3632850 | 1.257E-4 | 8.785E-2 | 0 | |
| 5 | 3,637,348 | chr5_3637348 | 1.022E-6 | 1.758E-2 | 0 | |
| 5 | 3,720,576 | chr5_3720576 | 6.652E-5 | 5.765E-2 | 0 | |

| Chr | Position (bp) | Marker | QCH P | Local FDR | Trait-level FDR count | Traits with BH <= 0.05 |
|---|---|---|---|---|---|---|
| 5 | 3,720,579 | chr5_3720579 | 2.478E-6 | 1.875E-2 | 0 | |
| 5 | 3,720,591 | chr5_3720591 | 7.636E-6 | 2.796E-2 | 0 | |
| 5 | 3,749,253 | chr5_3749253 | 7.353E-6 | 2.725E-2 | 0 | |
| 5 | 3,800,927 | chr5_3800927 | 5.043E-6 | 2.251E-2 | 0 | |
| 5 | 3,801,500 | chr5_3801500 | 1.022E-6 | 1.758E-2 | 0 | |
| 5 | 3,801,590 | chr5_3801590 | 1.022E-6 | 1.758E-2 | 0 | |
| 5 | 3,801,733 | chr5_3801733 | 4.518E-5 | 4.969E-2 | 0 | |
| 5 | 3,813,891 | chr5_3813891 | 1.213E-4 | 8.653E-2 | 0 | |
| 5 | 3,814,052 | chr5_3814052 | 3.933E-5 | 4.658E-2 | 0 | |
| 5 | 14,801,029 | chr5_14801029 | 1.329E-4 | 9.021E-2 | 1 | FLO_50 |
| 5 | 16,520,934 | chr5_16520934 | 1.813E-5 | 3.474E-2 | 1 | YIELD_OIV_504 |
| 5 | 16,970,241 | chr5_16970241 | 7.899E-5 | 6.55E-2 | 0 | |
| 5 | 20,538,484 | chr5_20538484 | 7.7E-5 | 6.345E-2 | 1 | SBER_W_g |
| 6 | 3,174,778 | chr6_3174778 | 6.711E-5 | 5.783E-2 | 1 | SBER_W_g |
| 6 | 5,833,646 | chr6_5833646 | 1.144E-4 | 8.295E-2 | 0 | |
| 6 | 5,833,647 | chr6_5833647 | 8.234E-5 | 6.653E-2 | 0 | |
| 6 | 5,833,695 | chr6_5833695 | 7.195E-5 | 6.106E-2 | 1 | HARVEST_DATE |
| 7 | 3,983,389 | chr7_3983389 | 9.296E-5 | 7.259E-2 | 1 | YIELD_PLANT |
| 8 | 10,980,744 | chr8_10980744 | 2.565E-5 | 3.866E-2 | 1 | NB_CLUST_PLANT |
| 8 | 15,522,696 | chr8_15522696 | 1.169E-4 | 8.454E-2 | 1 | SBER_W_g |
| 8 | 21,647,686 | chr8_21647686 | 5.515E-5 | 5.327E-2 | 1 | YIELD_OIV_504 |
| 9 | 915,526 | chr9_915526 | 3.608E-5 | 4.457E-2 | 1 | BUD_DATE |
| 9 | 2,793,911 | chr9_2793911 | 4.32E-5 | 4.809E-2 | 1 | BER_TA_g |
| 9 | 6,246,031 | chr9_6246031 | 1.006E-4 | 7.696E-2 | 0 | |
| 9 | 6,246,067 | chr9_6246067 | 1.006E-4 | 7.696E-2 | 0 | |
| 9 | 6,629,813 | chr9_6629813 | 1.879E-12 | 1.852E-7 | 2 | YIELD_PLANT; YIELD_OIV_504 |
| 10 | 8,361,332 | chr10_8361332 | 6.188E-5 | 5.656E-2 | 0 | |
| 10 | 8,361,333 | chr10_8361333 | 7.076E-6 | 2.722E-2 | 0 | |
| 10 | 12,758,127 | chr10_12758127 | 9.977E-5 | 7.575E-2 | 0 | |
| 10 | 19,736,493 | chr10_19736493 | 1.311E-4 | 8.876E-2 | 0 | |
| 11 | 1,033,705 | chr11_1033705 | 8.032E-5 | 6.578E-2 | 1 | NB_CLUST_PLANT |
| 11 | 3,758,688 | chr11_3758688 | 8.721E-5 | 6.901E-2 | 0 | |
| 11 | 3,824,935 | chr11_3824935 | 1.302E-4 | 8.873E-2 | 1 | YIELD_PLANT |
| 11 | 4,574,451 | chr11_4574451 | 1.093E-4 | 8.059E-2 | 1 | YIELD_PLANT |
| 11 | 6,089,413 | chr11_6089413 | 6.683E-7 | 1.711E-2 | 1 | FLO_50 |

| Chr | Position (bp) | Marker | QCH P | Local FDR | Trait-level FDR count | Traits with BH <= 0.05 |
|---|---|---|---|---|---|---|
| 11 | 7,023,371 | chr11_7023371 | 8.494E-6 | 2.854E-2 | 1 | NB_CLUST_PLANT |
| 11 | 8,070,948 | chr11_8070948 | 1.357E-4 | 9.056E-2 | 0 | |
| 11 | 8,070,955 | chr11_8070955 | 1.357E-4 | 9.056E-2 | 0 | |
| 11 | 8,125,007 | chr11_8125007 | 1.23E-4 | 8.683E-2 | 0 | |
| 11 | 8,141,640 | chr11_8141640 | 1.11E-4 | 8.179E-2 | 0 | |
| 11 | 8,149,351 | chr11_8149351 | 1.037E-4 | 7.794E-2 | 0 | |
| 11 | 8,149,404 | chr11_8149404 | 5.816E-8 | 5.101E-3 | 1 | YIELD_OIV_504 |
| 12 | 17,932,593 | chr12_17932593 | 1.37E-5 | 3.131E-2 | 1 | YIELD_PLANT |
| 12 | 18,014,166 | chr12_18014166 | 1.061E-4 | 7.875E-2 | 1 | TSS |
| 14 | 3,039,299 | chr14_3039299 | 9.149E-5 | 7.121E-2 | 1 | NB_CLUST_PLANT |
| 14 | 7,735,317 | chr14_7735317 | 1.118E-4 | 8.207E-2 | 1 | SBER_W_g |
| 14 | 13,952,316 | chr14_13952316 | 5.28E-6 | 2.332E-2 | 2 | YIELD_PLANT; YIELD_OIV_504 |
| 14 | 16,274,319 | chr14_16274319 | 4.367E-6 | 2.156E-2 | 1 | YIELD_OIV_504 |
| 14 | 17,151,022 | chr14_17151022 | 3.711E-6 | 2.093E-2 | 1 | YIELD_PLANT |
| 14 | 19,079,458 | chr14_19079458 | 3.498E-6 | 2.08E-2 | 2 | TSS; YIELD_OIV_504 |
| 14 | 26,369,635 | chr14_26369635 | 4.826E-5 | 5.118E-2 | 1 | YIELD_OIV_504 |
| 15 | 4,576,440 | chr15_4576440 | 8.651E-5 | 6.893E-2 | 1 | NB_CLUST_PLANT |
| 16 | 1,101,440 | chr16_1101440 | 1.135E-4 | 8.251E-2 | 1 | VER_50 |
| 16 | 12,507,629 | chr16_12507629 | 1.348E-4 | 9.054E-2 | 1 | YIELD_OIV_504 |
| 16 | 14,549,584 | chr16_14549584 | 4.333E-10 | 3.958E-5 | 2 | VER_50; HARVEST_DATE |
| 16 | 15,035,128 | chr16_15035128 | 0 | 0 | 2 | VER_50; HARVEST_DATE |
| 16 | 17,401,013 | chr16_17401013 | 3.18E-11 | 2.95E-6 | 2 | VER_50; HARVEST_DATE |
| 17 | 6,512,440 | chr17_6512440 | 9.369E-5 | 7.278E-2 | 1 | VER_50 |
| 18 | 2,100,422 | chr18_2100422 | 6.769E-5 | 5.798E-2 | 0 | |
| 18 | 2,440,997 | chr18_2440997 | 3.293E-5 | 4.306E-2 | 0 | |
| 18 | 4,386,621 | chr18_4386621 | 1.778E-5 | 3.458E-2 | 1 | YIELD_OIV_504 |
| 18 | 5,141,487 | chr18_5141487 | 2.066E-5 | 3.607E-2 | 0 | |
| 18 | 7,597,972 | chr18_7597972 | 5.96E-5 | 5.552E-2 | 1 | TSS |
| 18 | 9,529,420 | chr18_9529420 | 7.766E-5 | 6.482E-2 | 1 | BERRY_pH |
| 18 | 11,131,797 | chr18_11131797 | 1.921E-5 | 3.562E-2 | 1 | YIELD_PLANT |
| 18 | 23,048,559 | chr18_23048559 | 9.595E-5 | 7.457E-2 | 1 | FLO_50 |
| 18 | 34,522,453 | chr18_34522453 | 9.901E-5 | 7.552E-2 | 0 | |
| 19 | 3,309,782 | chr19_3309782 | 2.14E-5 | 3.665E-2 | 1 | VER_50 |
| 19 | 5,886,348 | chr19_5886348 | 2.369E-5 | 3.814E-2 | 1 | YIELD_OIV_504 |

| Chr | Position (bp) | Marker | QCH P | Local FDR | Trait-level FDR count | Traits with BH <= 0.05 |
|---|---|---|---|---|---|---|
| 19 | 13,212,585 | chr19_13212585 | 1.021E-4 | 7.738E-2 | 1 | NB_CLUST_PLANT |
| 19 | 17,074,417 | chr19_17074417 | 6.407E-9 | 5.888E-4 | 2 | VER_50; HARVEST_DATE |

**Supplementary Table 22.** Sensitivity of the cross-trait QCH analysis to the copula model. The Gaussian copula was primary; the independence copula was a sensitivity analysis. Rank correlations use all 136,062 intersected markers, whereas set overlap compares marker tests selected by the 5% QCH FDR procedure. Abbreviations: QCH, Query Composite Hypotheses; FDR, false discovery rate.

| N common markers | gaussian selected marker tests | independence selected marker tests | selected intersection | selected union | selected jaccard | spearman QCH P | spearman QCH local FDR | primary model | sensitivity model |
|---|---|---|---|---|---|---|---|---|---|
| 136,062 | 270 | 182 | 114 | 338 | 0.337 | 0.989 | 0.989 | gaussian | independence |

**Supplementary Table 23.** Prediction validation for package-default and population-adjusted rrBLUP marker models. Random-fold rows report both total prediction and the genomic component; leave-one-population-out rows report the genomic component because the held-out population intercept is not estimable from training data. Abbreviations: rrBLUP, ridge-regression best linear unbiased prediction.

| trait | model | validation | prediction component | folds or populations | mean correlation | median correlation | sd correlation | min correlation | max correlation |
|---|---|---|---|---|---|---|---|---|---|
| BUD_DATE | package_default_rrBLUP | stratified_random_5fold | total_including_population | 5 | 0.755 | 0.791 | 0.077 | 0.629 | 0.812 |
| BUD_DATE | population_adjusted_rrBLUP | stratified_random_5fold | total_including_population | 5 | 0.743 | 0.762 | 0.042 | 0.682 | 0.786 |
| FLO_50 | package_default_rrBLUP | stratified_random_5fold | total_including_population | 5 | 0.845 | 0.851 | 0.023 | 0.812 | 0.867 |
| FLO_50 | population_adjusted_rrBLUP | stratified_random_5fold | total_including_population | 5 | 0.924 | 0.925 | 0.011 | 0.911 | 0.935 |
| VER_50 | package_default_rrBLUP | stratified_random_5fold | total_including_population | 5 | 0.832 | 0.837 | 0.011 | 0.816 | 0.841 |
| VER_50 | population_adjusted_rrBLUP | stratified_random_5fold | total_including_population | 5 | 0.821 | 0.821 | 0.016 | 0.798 | 0.836 |
| SBER_W_g | package_default_rrBLUP | stratified_random_5fold | total_including_population | 5 | 0.811 | 0.815 | 0.017 | 0.79 | 0.833 |
| SBER_W_g | population_adjusted_rrBLUP | stratified_random_5fold | total_including_population | 5 | 0.807 | 0.812 | 0.015 | 0.785 | 0.823 |
| TSS | package_default_rrBLUP | stratified_random_5fold | total_including_population | 5 | 0.868 | 0.874 | 0.015 | 0.844 | 0.882 |
| TSS | population_adjusted_rrBLUP | stratified_random_5fold | total_including_population | 5 | 0.88 | 0.882 | 0.019 | 0.855 | 0.905 |
| BERRY_pH | package_default_rrBLUP | stratified_random_5fold | total_including_population | 5 | 0.819 | 0.825 | 0.027 | 0.781 | 0.853 |
| BERRY_pH | population_adjusted_rrBLUP | stratified_random_5fold | total_including_population | 5 | 0.831 | 0.832 | 0.02 | 0.804 | 0.857 |
| BER_TA_g | package_default_rrBLUP | stratified_random_5fold | total_including_population | 5 | 0.868 | 0.873 | 0.013 | 0.851 | 0.882 |
| BER_TA_g | population_adjusted_rrBLUP | stratified_random_5fold | total_including_population | 5 | 0.876 | 0.878 | 0.014 | 0.86 | 0.891 |
| HARVEST_DATE | package_default_rrBLUP | stratified_random_5fold | total_including_population | 5 | 0.82 | 0.813 | 0.024 | 0.796 | 0.848 |
| HARVEST_DATE | population_adjusted_rrBLUP | stratified_random_5fold | total_including_population | 5 | 0.813 | 0.821 | 0.024 | 0.786 | 0.837 |
| MORPHO_OIV_204 | package_default_rrBLUP | stratified_random_5fold | total_including_population | 5 | 0.713 | 0.731 | 0.04 | 0.667 | 0.752 |
| MORPHO_OIV_204 | population_adjusted_rrBLUP | stratified_random_5fold | total_including_population | 5 | 0.699 | 0.722 | 0.047 | 0.63 | 0.736 |
| NB_CLUST_PLANT | package_default_rrBLUP | stratified_random_5fold | total_including_population | 5 | 0.899 | 0.893 | 0.011 | 0.89 | 0.916 |
| NB_CLUST_PLANT | population_adjusted_rrBLUP | stratified_random_5fold | total_including_population | 5 | 0.905 | 0.905 | 0.01 | 0.893 | 0.92 |

| trait | model | validation | prediction component | folds or populations | mean correlation | median correlation | sd correlation | min correlation | max correlation |
|---|---|---|---|---|---|---|---|---|---|
| YIELD_PLANT | package_default_rrBLUP | stratified_random_5fold | total_including_population | 5 | 0.844 | 0.848 | 0.026 | 0.805 | 0.877 |
| YIELD_PLANT | population_adjusted_rrBLUP | stratified_random_5fold | total_including_population | 5 | 0.856 | 0.858 | 0.022 | 0.828 | 0.878 |
| SCLUST_W | package_default_rrBLUP | stratified_random_5fold | total_including_population | 5 | 0.813 | 0.816 | 0.03 | 0.772 | 0.84 |
| SCLUST_W | population_adjusted_rrBLUP | stratified_random_5fold | total_including_population | 5 | 0.826 | 0.827 | 0.021 | 0.797 | 0.85 |
| YIELD_OIV_504 | package_default_rrBLUP | stratified_random_5fold | total_including_population | 5 | 0.828 | 0.821 | 0.023 | 0.803 | 0.862 |
| YIELD_OIV_504 | population_adjusted_rrBLUP | stratified_random_5fold | total_including_population | 5 | 0.829 | 0.822 | 0.032 | 0.793 | 0.881 |
| BUD_DATE | package_default_rrBLUP | stratified_random_5fold | genomic_component | 5 | 0.755 | 0.791 | 0.077 | 0.629 | 0.812 |
| BUD_DATE | population_adjusted_rrBLUP | stratified_random_5fold | genomic_component | 5 | 0.486 | 0.449 | 0.075 | 0.408 | 0.568 |
| FLO_50 | package_default_rrBLUP | stratified_random_5fold | genomic_component | 5 | 0.845 | 0.851 | 0.023 | 0.812 | 0.867 |
| FLO_50 | population_adjusted_rrBLUP | stratified_random_5fold | genomic_component | 5 | 0.446 | 0.459 | 0.056 | 0.38 | 0.519 |
| VER_50 | package_default_rrBLUP | stratified_random_5fold | genomic_component | 5 | 0.832 | 0.837 | 0.011 | 0.816 | 0.841 |
| VER_50 | population_adjusted_rrBLUP | stratified_random_5fold | genomic_component | 5 | 0.61 | 0.595 | 0.069 | 0.517 | 0.7 |
| SBER_W_g | package_default_rrBLUP | stratified_random_5fold | genomic_component | 5 | 0.811 | 0.815 | 0.017 | 0.79 | 0.833 |
| SBER_W_g | population_adjusted_rrBLUP | stratified_random_5fold | genomic_component | 5 | 0.586 | 0.592 | 0.023 | 0.549 | 0.61 |
| TSS | package_default_rrBLUP | stratified_random_5fold | genomic_component | 5 | 0.868 | 0.874 | 0.015 | 0.844 | 0.882 |
| TSS | population_adjusted_rrBLUP | stratified_random_5fold | genomic_component | 5 | 0.429 | 0.476 | 0.116 | 0.3 | 0.536 |
| BERRY_pH | package_default_rrBLUP | stratified_random_5fold | genomic_component | 5 | 0.819 | 0.825 | 0.027 | 0.781 | 0.853 |
| BERRY_pH | population_adjusted_rrBLUP | stratified_random_5fold | genomic_component | 5 | 0.072 | 0.021 | 0.098 | -0.006 | 0.226 |
| BER_TA_g | package_default_rrBLUP | stratified_random_5fold | genomic_component | 5 | 0.868 | 0.873 | 0.013 | 0.851 | 0.882 |
| BER_TA_g | population_adjusted_rrBLUP | stratified_random_5fold | genomic_component | 5 | 0.571 | 0.577 | 0.074 | 0.496 | 0.652 |
| HARVEST_DATE | package_default_rrBLUP | stratified_random_5fold | genomic_component | 5 | 0.82 | 0.813 | 0.024 | 0.796 | 0.848 |
| HARVEST_DATE | population_adjusted_rrBLUP | stratified_random_5fold | genomic_component | 5 | 0.446 | 0.445 | 0.076 | 0.338 | 0.52 |
| MORPHO_OIV_204 | package_default_rrBLUP | stratified_random_5fold | genomic_component | 5 | 0.713 | 0.731 | 0.04 | 0.667 | 0.752 |
| MORPHO_OIV_204 | population_adjusted_rrBLUP | stratified_random_5fold | genomic_component | 5 | 0.595 | 0.58 | 0.042 | 0.552 | 0.652 |

| trait | model | validation | prediction component | folds or populations | mean correlation | median correlation | sd correlation | min correlation | max correlation |
|---|---|---|---|---|---|---|---|---|---|
| NB_CLUST_PLANT | package_default_rrBLUP | stratified_random_5fold | genomic_component | 5 | 0.899 | 0.893 | 0.011 | 0.89 | 0.916 |
| NB_CLUST_PLANT | population_adjusted_rrBLUP | stratified_random_5fold | genomic_component | 5 | 0.369 | 0.382 | 0.061 | 0.271 | 0.433 |
| YIELD_PLANT | package_default_rrBLUP | stratified_random_5fold | genomic_component | 5 | 0.844 | 0.848 | 0.026 | 0.805 | 0.877 |
| YIELD_PLANT | population_adjusted_rrBLUP | stratified_random_5fold | genomic_component | 5 | 0.508 | 0.544 | 0.082 | 0.363 | 0.561 |
| SCLUST_W | package_default_rrBLUP | stratified_random_5fold | genomic_component | 5 | 0.813 | 0.816 | 0.03 | 0.772 | 0.84 |
| SCLUST_W | population_adjusted_rrBLUP | stratified_random_5fold | genomic_component | 5 | 0.35 | 0.411 | 0.142 | 0.172 | 0.513 |
| YIELD_OIV_504 | package_default_rrBLUP | stratified_random_5fold | genomic_component | 5 | 0.828 | 0.821 | 0.023 | 0.803 | 0.862 |
| YIELD_OIV_504 | population_adjusted_rrBLUP | stratified_random_5fold | genomic_component | 5 | 0.503 | 0.497 | 0.026 | 0.475 | 0.545 |
| BUD_DATE | package_default_rrBLUP | leave_one_population_out | genomic_component | 12 | 0.417 | 0.426 | 0.144 | 0.1 | 0.685 |
| BUD_DATE | population_adjusted_rrBLUP | leave_one_population_out | genomic_component | 12 | 0.415 | 0.386 | 0.138 | 0.152 | 0.664 |
| FLO_50 | package_default_rrBLUP | leave_one_population_out | genomic_component | 12 | 0.409 | 0.422 | 0.166 | 0.204 | 0.656 |
| FLO_50 | population_adjusted_rrBLUP | leave_one_population_out | genomic_component | 12 | 0.501 | 0.51 | 0.136 | 0.269 | 0.733 |
| VER_50 | package_default_rrBLUP | leave_one_population_out | genomic_component | 12 | 0.468 | 0.478 | 0.124 | 0.272 | 0.667 |
| VER_50 | population_adjusted_rrBLUP | leave_one_population_out | genomic_component | 12 | 0.489 | 0.483 | 0.113 | 0.324 | 0.661 |
| SBER_W_g | package_default_rrBLUP | leave_one_population_out | genomic_component | 12 | 0.574 | 0.563 | 0.084 | 0.455 | 0.685 |
| SBER_W_g | population_adjusted_rrBLUP | leave_one_population_out | genomic_component | 12 | 0.58 | 0.56 | 0.081 | 0.475 | 0.691 |
| TSS | package_default_rrBLUP | leave_one_population_out | genomic_component | 12 | 0.522 | 0.516 | 0.143 | 0.352 | 0.864 |
| TSS | population_adjusted_rrBLUP | leave_one_population_out | genomic_component | 12 | 0.539 | 0.566 | 0.133 | 0.308 | 0.813 |
| BERRY_pH | package_default_rrBLUP | leave_one_population_out | genomic_component | 12 | 0.439 | 0.425 | 0.131 | 0.229 | 0.67 |
| BERRY_pH | population_adjusted_rrBLUP | leave_one_population_out | genomic_component | 12 | 0.468 | 0.486 | 0.135 | 0.248 | 0.682 |
| BER_TA_g | package_default_rrBLUP | leave_one_population_out | genomic_component | 12 | 0.495 | 0.519 | 0.134 | 0.203 | 0.681 |
| BER_TA_g | population_adjusted_rrBLUP | leave_one_population_out | genomic_component | 12 | 0.512 | 0.524 | 0.13 | 0.26 | 0.699 |
| HARVEST_DATE | package_default_rrBLUP | leave_one_population_out | genomic_component | 12 | 0.437 | 0.418 | 0.132 | 0.229 | 0.629 |
| HARVEST_DATE | population_adjusted_rrBLUP | leave_one_population_out | genomic_component | 12 | 0.467 | 0.44 | 0.15 | 0.251 | 0.698 |

| trait | model | validation | prediction component | folds or populations | mean correlation | median correlation | sd correlation | min correlation | max correlation |
|---|---|---|---|---|---|---|---|---|---|
| MORPHO_OIV_204 | package_default_rrBLUP | leave_one_population_out | genomic_component | 12 | 0.381 | 0.374 | 0.237 | 0.016 | 0.688 |
| MORPHO_OIV_204 | population_adjusted_rrBLUP | leave_one_population_out | genomic_component | 12 | 0.382 | 0.395 | 0.243 | -0.032 | 0.68 |
| NB_CLUST_PLANT | package_default_rrBLUP | leave_one_population_out | genomic_component | 11 | 0.485 | 0.515 | 0.178 | 0.126 | 0.657 |
| NB_CLUST_PLANT | population_adjusted_rrBLUP | leave_one_population_out | genomic_component | 11 | 0.514 | 0.58 | 0.199 | 0.034 | 0.718 |
| YIELD_PLANT | package_default_rrBLUP | leave_one_population_out | genomic_component | 11 | 0.414 | 0.445 | 0.184 | 0.099 | 0.611 |
| YIELD_PLANT | population_adjusted_rrBLUP | leave_one_population_out | genomic_component | 11 | 0.447 | 0.525 | 0.18 | 0.067 | 0.621 |
| SCLUST_W | package_default_rrBLUP | leave_one_population_out | genomic_component | 11 | 0.373 | 0.403 | 0.148 | 0.12 | 0.579 |
| SCLUST_W | population_adjusted_rrBLUP | leave_one_population_out | genomic_component | 11 | 0.392 | 0.435 | 0.152 | 0.09 | 0.562 |
| YIELD_OIV_504 | package_default_rrBLUP | leave_one_population_out | genomic_component | 12 | 0.369 | 0.404 | 0.223 | -0.056 | 0.616 |
| YIELD_OIV_504 | population_adjusted_rrBLUP | leave_one_population_out | genomic_component | 12 | 0.402 | 0.443 | 0.199 | 0.073 | 0.607 |

**Supplementary Table 24.** Five highest-ranked population-adjusted localGEBV blocks per trait and overlap with the three mapping frameworks. LocalGEBV-only blocks remain breeding hypotheses rather than validated QTL. Abbreviations: localGEBV, local genomic estimated breeding value; QTL, quantitative trait locus; mpQTL, multiple-population QTL mapping; GWAS, genome-wide association study; SNP, single-nucleotide polymorphism.

| trait | variance rank | Chrom | Start Pos | End Pos | Num SNP | variance fraction | mpQTL overlap | single population QTL overlap | GWAS overlap | evidence class |
|---|---|---|---|---|---|---|---|---|---|---|
| BUD_DATE | 1 | 16 | 15,309,523 | 15,360,686 | 6 | 0.069 | TRUE | TRUE | TRUE | LocalGEBV + single-population QTL + mpQTL + structure-aware GWAS |
| BUD_DATE | 2 | 6 | 3,140,534 | 3,243,475 | 16 | 0.02 | FALSE | FALSE | FALSE | LocalGEBV only |
| BUD_DATE | 3 | 14 | 20,144,868 | 20,919,992 | 15 | 0.015 | FALSE | FALSE | FALSE | LocalGEBV only |
| BUD_DATE | 4 | 8 | 15,730,895 | 16,056,425 | 12 | 0.015 | FALSE | FALSE | FALSE | LocalGEBV only |
| BUD_DATE | 5 | 17 | 4,216,948 | 4,420,933 | 10 | 0.011 | FALSE | FALSE | FALSE | LocalGEBV only |
| FLO_50 | 1 | 17 | 4,216,948 | 4,420,933 | 10 | 0.027 | FALSE | FALSE | FALSE | LocalGEBV only |
| FLO_50 | 2 | 17 | 6,478,484 | 6,490,445 | 9 | 0.019 | TRUE | TRUE | FALSE | LocalGEBV + two mapping frameworks |
| FLO_50 | 3 | 8 | 15,730,895 | 16,056,425 | 12 | 0.017 | FALSE | FALSE | FALSE | LocalGEBV only |
| FLO_50 | 4 | 17 | 9,214,634 | 9,217,879 | 3 | 0.015 | FALSE | FALSE | FALSE | LocalGEBV only |
| FLO_50 | 5 | 16 | 15,309,523 | 15,360,686 | 6 | 0.015 | FALSE | FALSE | FALSE | LocalGEBV only |
| VER_50 | 1 | 16 | 14,689,860 | 14,689,931 | 2 | 0.086 | TRUE | TRUE | TRUE | LocalGEBV + single-population QTL + mpQTL + structure-aware GWAS |
| VER_50 | 2 | 16 | 15,309,523 | 15,360,686 | 6 | 0.073 | FALSE | TRUE | TRUE | LocalGEBV + two mapping frameworks |
| VER_50 | 3 | 8 | 15,730,895 | 16,056,425 | 12 | 0.028 | FALSE | TRUE | FALSE | LocalGEBV + one mapping framework |
| VER_50 | 4 | 16 | 7,122,451 | 7,122,451 | 1 | 0.021 | TRUE | TRUE | TRUE | LocalGEBV + single-population QTL + mpQTL + structure-aware GWAS |
| VER_50 | 5 | 16 | 7,157,963 | 8,800,534 | 6 | 0.02 | TRUE | TRUE | TRUE | LocalGEBV + single-population QTL + mpQTL + structure-aware GWAS |
| SBER_W_g | 1 | 3 | 2,146,767 | 2,254,192 | 9 | 0.034 | FALSE | FALSE | TRUE | LocalGEBV + one mapping framework |
| SBER_W_g | 2 | 7 | 23,742,284 | 23,789,464 | 8 | 0.022 | FALSE | TRUE | TRUE | LocalGEBV + two mapping frameworks |
| SBER_W_g | 3 | 17 | 6,561,274 | 6,646,471 | 6 | 0.019 | TRUE | TRUE | FALSE | LocalGEBV + two mapping frameworks |
| SBER_W_g | 4 | 5 | 10,639,856 | 12,106,899 | 15 | 0.013 | FALSE | FALSE | FALSE | LocalGEBV only |
| SBER_W_g | 5 | 16 | 14,689,860 | 14,689,931 | 2 | 0.013 | FALSE | FALSE | FALSE | LocalGEBV only |

| trait | variance rank | Chrom | Start Pos | End Pos | Num SNP | variance fraction | mpQTL overlap | single population QTL overlap | GWAS overlap | evidence class |
|---|---|---|---|---|---|---|---|---|---|---|
| TSS | 1 | 9 | 2,720,930 | 2,720,995 | 6 | 0.015 | FALSE | FALSE | FALSE | LocalGEBV only |
| TSS | 2 | 18 | 14,880,737 | 18,353,648 | 7 | 0.013 | FALSE | TRUE | FALSE | LocalGEBV + one mapping framework |
| TSS | 3 | 1 | 22,660,003 | 22,965,397 | 4 | 0.013 | FALSE | TRUE | TRUE | LocalGEBV + two mapping frameworks |
| TSS | 4 | 9 | 661,851 | 700,608 | 3 | 0.012 | FALSE | TRUE | FALSE | LocalGEBV + one mapping framework |
| TSS | 5 | 3 | 4,682,774 | 4,893,805 | 5 | 0.01 | FALSE | FALSE | FALSE | LocalGEBV only |
| BERRY_pH | 1 | 16 | 14,689,860 | 14,689,931 | 2 | 0.024 | TRUE | FALSE | FALSE | LocalGEBV + one mapping framework |
| BERRY_pH | 2 | 6 | 3,140,534 | 3,243,475 | 16 | 0.018 | FALSE | FALSE | FALSE | LocalGEBV only |
| BERRY_pH | 3 | 2 | 4,703,137 | 4,724,473 | 5 | 0.016 | FALSE | FALSE | FALSE | LocalGEBV only |
| BERRY_pH | 4 | 10 | 446,952 | 488,984 | 14 | 0.013 | FALSE | FALSE | FALSE | LocalGEBV only |
| BERRY_pH | 5 | 13 | 10,810 | 651,373 | 10 | 0.012 | FALSE | FALSE | TRUE | LocalGEBV + one mapping framework |
| BER_TA_g | 1 | 11 | 18,500,842 | 18,646,234 | 6 | 0.018 | FALSE | FALSE | FALSE | LocalGEBV only |
| BER_TA_g | 2 | 14 | 29,206,449 | 29,454,105 | 13 | 0.018 | FALSE | TRUE | FALSE | LocalGEBV + one mapping framework |
| BER_TA_g | 3 | 6 | 3,140,534 | 3,243,475 | 16 | 0.017 | FALSE | FALSE | FALSE | LocalGEBV only |
| BER_TA_g | 4 | 16 | 14,689,860 | 14,689,931 | 2 | 0.017 | FALSE | TRUE | TRUE | LocalGEBV + two mapping frameworks |
| BER_TA_g | 5 | 10 | 446,952 | 488,984 | 14 | 0.014 | FALSE | FALSE | FALSE | LocalGEBV only |
| HARVEST_DATE | 1 | 16 | 15,309,523 | 15,360,686 | 6 | 0.071 | FALSE | FALSE | TRUE | LocalGEBV + one mapping framework |
| HARVEST_DATE | 2 | 16 | 14,689,860 | 14,689,931 | 2 | 0.058 | TRUE | TRUE | TRUE | LocalGEBV + single-population QTL + mpQTL + structure-aware GWAS |
| HARVEST_DATE | 3 | 8 | 15,730,895 | 16,056,425 | 12 | 0.023 | FALSE | TRUE | FALSE | LocalGEBV + one mapping framework |
| HARVEST_DATE | 4 | 16 | 7,157,963 | 8,800,534 | 6 | 0.021 | TRUE | FALSE | TRUE | LocalGEBV + two mapping frameworks |
| HARVEST_DATE | 5 | 16 | 7,122,451 | 7,122,451 | 1 | 0.018 | TRUE | FALSE | TRUE | LocalGEBV + two mapping frameworks |
| MORPHO_OIV_204 | 1 | 16 | 21,788,659 | 21,788,710 | 5 | 0.021 | FALSE | FALSE | FALSE | LocalGEBV only |
| MORPHO_OIV_204 | 2 | 2 | 5,130,512 | 6,438,783 | 18 | 0.02 | FALSE | TRUE | FALSE | LocalGEBV + one mapping framework |
| MORPHO_OIV_204 | 3 | 8 | 15,730,895 | 16,056,425 | 12 | 0.014 | FALSE | FALSE | FALSE | LocalGEBV only |
| MORPHO_OIV_204 | 4 | 11 | 3,359,297 | 3,716,224 | 14 | 0.014 | FALSE | FALSE | FALSE | LocalGEBV only |
| MORPHO_OIV_204 | 5 | 16 | 20,056,962 | 20,257,035 | 4 | 0.013 | FALSE | FALSE | FALSE | LocalGEBV only |

| trait | variance rank | Chrom | Start Pos | End Pos | Num SNP | variance fraction | mpQTL overlap | single population QTL overlap | GWAS overlap | evidence class |
|---|---|---|---|---|---|---|---|---|---|---|
| NB_CLUST_PLANT | 1 | 16 | 15,309,523 | 15,360,686 | 6 | 0.024 | FALSE | FALSE | FALSE | LocalGEBV only |
| NB_CLUST_PLANT | 2 | 4 | 1,820,946 | 1,821,002 | 4 | 0.016 | FALSE | FALSE | FALSE | LocalGEBV only |
| NB_CLUST_PLANT | 3 | 11 | 3,359,297 | 3,716,224 | 14 | 0.016 | FALSE | FALSE | FALSE | LocalGEBV only |
| NB_CLUST_PLANT | 4 | 15 | 21,733,428 | 21,733,484 | 3 | 0.015 | FALSE | FALSE | FALSE | LocalGEBV only |
| NB_CLUST_PLANT | 5 | 5 | 10,639,856 | 12,106,899 | 15 | 0.011 | FALSE | FALSE | FALSE | LocalGEBV only |
| YIELD_PLANT | 1 | 11 | 3,359,297 | 3,716,224 | 14 | 0.018 | FALSE | TRUE | FALSE | LocalGEBV + one mapping framework |
| YIELD_PLANT | 2 | 5 | 10,639,856 | 12,106,899 | 15 | 0.016 | FALSE | FALSE | FALSE | LocalGEBV only |
| YIELD_PLANT | 3 | 4 | 1,820,946 | 1,821,002 | 4 | 0.013 | FALSE | FALSE | FALSE | LocalGEBV only |
| YIELD_PLANT | 4 | 1 | 1,891,840 | 1,904,413 | 5 | 0.013 | FALSE | FALSE | FALSE | LocalGEBV only |
| YIELD_PLANT | 5 | 8 | 9,870,913 | 10,768,579 | 14 | 0.011 | FALSE | FALSE | FALSE | LocalGEBV only |
| SCLUST_W | 1 | 15 | 14,560,035 | 14,763,239 | 11 | 0.026 | FALSE | FALSE | FALSE | LocalGEBV only |
| SCLUST_W | 2 | 16 | 15,309,523 | 15,360,686 | 6 | 0.026 | FALSE | TRUE | TRUE | LocalGEBV + two mapping frameworks |
| SCLUST_W | 3 | 8 | 9,870,913 | 10,768,579 | 14 | 0.014 | FALSE | TRUE | FALSE | LocalGEBV + one mapping framework |
| SCLUST_W | 4 | 13 | 20,347,313 | 20,387,370 | 4 | 0.014 | FALSE | FALSE | FALSE | LocalGEBV only |
| SCLUST_W | 5 | 5 | 10,639,856 | 12,106,899 | 15 | 0.013 | FALSE | FALSE | FALSE | LocalGEBV only |
| YIELD_OIV_504 | 1 | 5 | 10,639,856 | 12,106,899 | 15 | 0.025 | FALSE | FALSE | FALSE | LocalGEBV only |
| YIELD_OIV_504 | 2 | 4 | 1,820,946 | 1,821,002 | 4 | 0.017 | FALSE | FALSE | FALSE | LocalGEBV only |
| YIELD_OIV_504 | 3 | 11 | 3,359,297 | 3,716,224 | 14 | 0.015 | FALSE | TRUE | FALSE | LocalGEBV + one mapping framework |
| YIELD_OIV_504 | 4 | 15 | 14,560,035 | 14,763,239 | 11 | 0.012 | FALSE | FALSE | FALSE | LocalGEBV only |
| YIELD_OIV_504 | 5 | 8 | 9,870,913 | 10,768,579 | 14 | 0.011 | FALSE | FALSE | FALSE | LocalGEBV only |

**Supplementary Table 25.** Mapping-framework support among the top 1% of population-adjusted localGEBV blocks for each trait. Abbreviations: localGEBV, local genomic estimated breeding value; QTL, quantitative trait locus; mpQTL, multiple-population QTL mapping; GWAS, genome-wide association study.

| trait | top blocks | localGEBV only | overlap one mapping framework | overlap two mapping frameworks | overlap all three mapping frameworks | fraction with mapping support | top block variance fraction | top 1pct variance fraction |
|---|---|---|---|---|---|---|---|---|
| BUD_DATE | 46 | 39 | 3 | 3 | 1 | 0.152 | 0.069 | 0.342 |
| FLO_50 | 46 | 32 | 10 | 4 | 0 | 0.304 | 0.027 | 0.333 |
| VER_50 | 46 | 33 | 6 | 4 | 3 | 0.283 | 0.086 | 0.469 |
| SBER_W_g | 46 | 30 | 11 | 5 | 0 | 0.348 | 0.034 | 0.326 |
| TSS | 46 | 34 | 10 | 2 | 0 | 0.261 | 0.015 | 0.301 |
| BERRY_pH | 46 | 35 | 10 | 1 | 0 | 0.239 | 0.024 | 0.33 |
| BER_TA_g | 46 | 31 | 13 | 2 | 0 | 0.326 | 0.018 | 0.353 |
| HARVEST_DATE | 46 | 32 | 9 | 4 | 1 | 0.304 | 0.071 | 0.418 |
| MORPHO_OIV_204 | 46 | 37 | 6 | 3 | 0 | 0.196 | 0.021 | 0.321 |
| NB_CLUST_PLANT | 46 | 36 | 10 | 0 | 0 | 0.217 | 0.024 | 0.315 |
| YIELD_PLANT | 46 | 37 | 6 | 3 | 0 | 0.196 | 0.018 | 0.313 |
| SCLUST_W | 46 | 34 | 10 | 2 | 0 | 0.261 | 0.026 | 0.341 |
| YIELD_OIV_504 | 46 | 34 | 12 | 0 | 0 | 0.261 | 0.025 | 0.313 |

**Supplementary Table 26.** Nested leave-one-population-out validation of training-selected localGEBV block subsets. Block ranking was recomputed within each training set before prediction of the held-out population. Abbreviations: localGEBV, local genomic estimated breeding value.

| trait | selected block fraction | selected blocks | populations | mean correlation | median correlation | sd correlation | mean whole genome correlation | populations better than whole genome | mean delta vs whole genome |
|---|---|---|---|---|---|---|---|---|---|
| BUD_DATE | 0.01 | 46 | 12 | 0.261 | 0.242 | 0.144 | 0.415 | 1 | -0.153 |
| BUD_DATE | 0.05 | 230 | 12 | 0.345 | 0.313 | 0.144 | 0.415 | 0 | -0.07 |
| BUD_DATE | 0.1 | 460 | 12 | 0.366 | 0.345 | 0.142 | 0.415 | 1 | -0.048 |
| BUD_DATE | 1 | 4,600 | 12 | 0.415 | 0.386 | 0.138 | 0.415 | 5 | 0 |
| FLO_50 | 0.01 | 46 | 12 | 0.411 | 0.402 | 0.12 | 0.501 | 3 | -0.091 |
| FLO_50 | 0.05 | 230 | 12 | 0.49 | 0.492 | 0.126 | 0.501 | 5 | -0.011 |
| FLO_50 | 0.1 | 460 | 12 | 0.497 | 0.495 | 0.139 | 0.501 | 6 | -0.005 |
| FLO_50 | 1 | 4,600 | 12 | 0.501 | 0.51 | 0.136 | 0.501 | 6 | 0 |
| VER_50 | 0.01 | 46 | 12 | 0.438 | 0.485 | 0.25 | 0.489 | 5 | -0.051 |
| VER_50 | 0.05 | 230 | 12 | 0.521 | 0.521 | 0.154 | 0.489 | 6 | 0.032 |
| VER_50 | 0.1 | 460 | 12 | 0.517 | 0.499 | 0.158 | 0.489 | 9 | 0.028 |
| VER_50 | 1 | 4,600 | 12 | 0.489 | 0.483 | 0.113 | 0.489 | 7 | 0 |
| SBER_W_g | 0.01 | 46 | 12 | 0.432 | 0.46 | 0.129 | 0.58 | 2 | -0.149 |
| SBER_W_g | 0.05 | 230 | 12 | 0.497 | 0.485 | 0.054 | 0.58 | 2 | -0.084 |
| SBER_W_g | 0.1 | 460 | 12 | 0.511 | 0.504 | 0.06 | 0.58 | 2 | -0.069 |
| SBER_W_g | 1 | 4,600 | 12 | 0.58 | 0.56 | 0.081 | 0.58 | 9 | 0 |
| TSS | 0.01 | 46 | 12 | 0.306 | 0.31 | 0.102 | 0.539 | 0 | -0.233 |
| TSS | 0.05 | 230 | 12 | 0.498 | 0.464 | 0.136 | 0.539 | 3 | -0.041 |
| TSS | 0.1 | 460 | 12 | 0.512 | 0.498 | 0.141 | 0.539 | 4 | -0.026 |
| TSS | 1 | 4,600 | 12 | 0.539 | 0.566 | 0.133 | 0.539 | 3 | 0 |
| BERRY_pH | 0.01 | 46 | 12 | 0.294 | 0.328 | 0.143 | 0.468 | 0 | -0.174 |
| BERRY_pH | 0.05 | 230 | 12 | 0.428 | 0.46 | 0.154 | 0.468 | 3 | -0.041 |
| BERRY_pH | 0.1 | 460 | 12 | 0.444 | 0.483 | 0.163 | 0.468 | 3 | -0.024 |
| BERRY_pH | 1 | 4,600 | 12 | 0.468 | 0.486 | 0.135 | 0.468 | 4 | 0 |
| BER_TA_g | 0.01 | 46 | 12 | 0.45 | 0.467 | 0.129 | 0.512 | 1 | -0.062 |

| trait | selected block fraction | selected blocks | populations | mean correlation | median correlation | sd correlation | mean whole genome correlation | populations better than whole genome | mean delta vs whole genome |
|---|---|---|---|---|---|---|---|---|---|
| BER_TA_g | 0.05 | 230 | 12 | 0.484 | 0.484 | 0.154 | 0.512 | 4 | -0.028 |
| BER_TA_g | 0.1 | 460 | 12 | 0.497 | 0.499 | 0.144 | 0.512 | 5 | -0.015 |
| BER_TA_g | 1 | 4,600 | 12 | 0.512 | 0.524 | 0.13 | 0.512 | 5 | 0 |
| HARVEST_DATE | 0.01 | 46 | 12 | 0.468 | 0.506 | 0.304 | 0.467 | 6 | 0 |
| HARVEST_DATE | 0.05 | 230 | 12 | 0.464 | 0.473 | 0.226 | 0.467 | 7 | -0.004 |
| HARVEST_DATE | 0.1 | 460 | 12 | 0.471 | 0.444 | 0.208 | 0.467 | 7 | 0.003 |
| HARVEST_DATE | 1 | 4,600 | 12 | 0.467 | 0.44 | 0.15 | 0.467 | 5 | 0 |
| MORPHO_OIV_204 | 0.01 | 46 | 12 | 0.323 | 0.281 | 0.193 | 0.382 | 4 | -0.059 |
| MORPHO_OIV_204 | 0.05 | 230 | 12 | 0.39 | 0.397 | 0.237 | 0.382 | 6 | 0.008 |
| MORPHO_OIV_204 | 0.1 | 460 | 12 | 0.384 | 0.399 | 0.249 | 0.382 | 5 | 0.002 |
| MORPHO_OIV_204 | 1 | 4,600 | 12 | 0.382 | 0.395 | 0.243 | 0.382 | 6 | 0 |
| NB_CLUST_PLANT | 0.01 | 46 | 11 | 0.411 | 0.503 | 0.203 | 0.514 | 1 | -0.103 |
| NB_CLUST_PLANT | 0.05 | 230 | 11 | 0.467 | 0.516 | 0.21 | 0.514 | 2 | -0.047 |
| NB_CLUST_PLANT | 0.1 | 460 | 11 | 0.506 | 0.542 | 0.196 | 0.514 | 4 | -0.008 |
| NB_CLUST_PLANT | 1 | 4,600 | 11 | 0.514 | 0.58 | 0.199 | 0.514 | 5 | 0 |
| YIELD_PLANT | 0.01 | 46 | 11 | 0.31 | 0.309 | 0.17 | 0.447 | 1 | -0.137 |
| YIELD_PLANT | 0.05 | 230 | 11 | 0.427 | 0.476 | 0.157 | 0.447 | 4 | -0.02 |
| YIELD_PLANT | 0.1 | 460 | 11 | 0.429 | 0.484 | 0.163 | 0.447 | 4 | -0.017 |
| YIELD_PLANT | 1 | 4,600 | 11 | 0.447 | 0.525 | 0.18 | 0.447 | 3 | 0 |
| SCLUST_W | 0.01 | 46 | 11 | 0.268 | 0.26 | 0.166 | 0.392 | 2 | -0.124 |
| SCLUST_W | 0.05 | 230 | 11 | 0.388 | 0.424 | 0.151 | 0.392 | 5 | -0.005 |
| SCLUST_W | 0.1 | 460 | 11 | 0.376 | 0.414 | 0.151 | 0.392 | 4 | -0.016 |
| SCLUST_W | 1 | 4,600 | 11 | 0.392 | 0.435 | 0.152 | 0.392 | 6 | 0 |
| YIELD_OIV_504 | 0.01 | 46 | 12 | 0.271 | 0.323 | 0.172 | 0.402 | 1 | -0.132 |
| YIELD_OIV_504 | 0.05 | 230 | 12 | 0.391 | 0.452 | 0.155 | 0.402 | 4 | -0.012 |
| YIELD_OIV_504 | 0.1 | 460 | 12 | 0.416 | 0.461 | 0.166 | 0.402 | 7 | 0.014 |

| trait | selected block fraction | selected blocks | populations | mean correlation | median correlation | sd correlation | mean whole genome correlation | populations better than whole genome | mean delta vs whole genome |
|---|---|---|---|---|---|---|---|---|---|
| YIELD_OIV_504 | 1 | 4,600 | 12 | 0.402 | 0.443 | 0.199 | 0.402 | 7 | 0 |

**Supplementary Table 27.** Trait-matched, assembly-harmonized comparison of prioritized ResDur regions with previous grapevine studies. Same-chromosome evidence alone was not treated as replication. Coordinates are PN40024.v4.3 unless stated otherwise; stable aliases were resolved with the PN40024.v4.3 GFF3. References are identified by author, publication year and journal. Abbreviations: GWAS, genome-wide association study; K, potassium; LD, linkage disequilibrium; QTL, quantitative trait locus; TA, titratable acidity; TSS, total soluble solids.

| Current ResDur evidence | External study and trait | Population background | Harmonized external locus or candidate | Concordance class | Interpretation and caveat |
|---|---|---|---|---|---|
| Chr13 TA, family 50015; 3.90-7.10 Mb; GWAS LD 5.026-5.991 Mb | Duchêne et al. 2020, Theoretical and Applied Genetics; pH, tartaric acid and K:tartaric acid | *V. vinifera*: Riesling x Gewürztraminer | 12X.v2 VIT_13s0019g04230 = v4.3 Vitvi13g00601, 5.650-5.653 Mb | Exact gene and overlapping interval | Strong external support for the region and a transporter candidate; causal allele and direction remain untested. |
| Chr13 TA, family 50015; 3.90-7.10 Mb | Ban et al. 2016, Euphytica; stable TA | Interspecific: *V. labruscana* x *V. vinifera* | LG13 near VVS1/VVIC51; physical interval unavailable | Chromosome-level only | Supports LG13 for TA in an interspecific family but cannot establish positional identity. |
| Chr17 pH 5.90-9.60 Mb; TA 6.80-16.60 Mb | Reshef et al. 2022, Horticulture Research; ripe-fruit malate | Complex interspecific families | 12X.v2 6.75-8.93 Mb; VIT_17s0000g06270 = Vitvi17g00607, 7.578-7.581 Mb | Overlapping interval and exact gene alias | Trait-related support only: ResDur measured pH and total TA, and the current loci remain family-specific. |
| Chr16 TA 12.50-17.30 Mb; pH GWAS LD 16.192-16.793 Mb | Kadium et al. 2026, Horticulture Research; sugars, organic acids and pH | Multiparental interspecific hybrid population | Vitvi16g00860, 15.175-15.208 Mb; pH SNP at ~16.738 Mb | Exact gene/coordinate within broad regions | Independent candidate and regional support; shared allele and effect direction are unknown. |
| Chr6 TA 15.40-21.20 Mb | Alahakoon et al. 2022, Plants; TA and malate | Interspecific F2: *V. riparia* x Seyval blanc ancestry | Broad chr6 acid intervals ~0.28-18.52 Mb; candidates ~10.9 Mb | Partial broad-interval contact | Reported candidate genes do not fall inside the ResDur TA interval; not a direct match. |
| Principal ResDur acid regions on chrs13, 16 and 17 | Mamani et al. 2021, Australian Journal of Grape and Wine Research; stable TA and tartaric acid | *V. vinifera*: Ruby Seedless x Sultanina | LG5 and LG8 | Non-concordant | Stable acid QTLs in another cross occur elsewhere, supporting background-dependent architecture. |
| Chr17 berry/cluster QTLs, approximately 6.6-10.3 Mb | Doligez et al. 2013, BMC Plant Biology; berry weight; Richter et al. 2019, Theoretical and Applied Genetics; cluster architecture | *V. vinifera* crosses (Doligez et al. 2013); disease-resistant hybrid family (Richter et al. 2019) | Prior region ~6.59-9.61 Mb; candidates Vitvi17g00484, Vitvi17g00556, Vitvi17g00578 and Vitvi17g00616 | Overlapping regional evidence | Supports a recurrent architecture/berry hub; pleiotropy versus linked loci is unresolved. |
| Chr1 compactness GWAS LD 5.228-5.791 Mb | de Oliveira et al. 2026, PLOS ONE; cluster weight | Diverse *Vitis* panel: *vinifera*, interspecific hybrids and non-*vinifera* spp. | PN40024.v4 SNP at 5.917 Mb | Near, correlated trait | ~0.126 Mb gap and a different cluster subtrait; partial regional concordance. |
| Chr14 compactness and cluster-weight GWAS LD 26.074-27.518 Mb | Thorat et al. 2024, Scientia Horticulturae and de Oliveira et al. | *V. vinifera* panel (Thorat et al. 2024); diverse mixed-Vitis | Remapped region approximately 25.895-26.124 Mb | Near/partly overlapping, correlated trait | Supports a cluster-architecture region, not replication of an identical phenotype. |

| Current ResDur evidence | External study and trait | Population background | Harmonized external locus or candidate | Concordance class | Interpretation and caveat |
|---|---|---|---|---|---|
| | 2026, PLOS ONE; cluster length | panel (de Oliveira et al. 2026) | | | |
| Chr17 TSS QTLs 1.90-6.00 Mb | Kadium et al. 2026, Horticulture Research; TSS | Multiparental interspecific hybrid population | Reported SNPs at ~4.242 and 6.082 Mb | Overlapping/edge contact | Same trait and regional support, but the ResDur signals are broad and not all frameworks concur. |
| Chr6 TSS QTL 3.60-7.30 Mb | Kadium et al. 2026, Horticulture Research; TSS | Multiparental interspecific hybrid population | Reported TSS SNPs ~6.435-9.365 Mb | Partial overlap | Same trait and partial regional recurrence; fine-scale identity is unresolved. |
| Chr16 veraison/harvest LD 14.55-15.49 Mb | Frenzke et al. 2024, Plant Physiology; Ver1 | Fungus-resistant breeding F1: Calardis Musqué x Villard Blanc | Vitvi16g00942 (VviERF027), ~16.421 Mb | Distinct refined positions | The broad family QTL spans Ver1, but the compact ResDur multi-framework signal does not. |
| Cross-family marker transfer | de Sousa Moreira et al. 2024, Journal of the American Society for Horticultural Science; multiple fruit-quality traits | Interspecific mapping family; diverse validation panel | Existing markers tested across grape use types | Conceptual transferability evidence | Low overall transfer supports independent-cross validation before marker-assisted deployment. |

**Supplementary Table 28.** Summary of filtered VCF files and genetic maps. Abbreviations: VCF, variant call format; SNP, single-nucleotide polymorphism.

| Population | Number of individuals | Number of SNPs in the VCF | Number of SNPs in the genetic map |
|---|---|---|---|
| ResDur (GWAS) | 1,081 | 189,426 | |
| Multi-parental population | 792 | 1,783,985 | 62,203 |
| 42050 | 108 | 1,158,120 | 25,297 |
| 50001 | 71 | 875,601 | 17,714 |
| 50013 | 61 | 868,981 | 16,223 |
| 50015 | 58 | 884,775 | 16,619 |
| 50025 | 103 | 973,593 | 19,268 |
| 50035 | 189 | 1,227,968 | 16,533 |

**Supplementary Table 29.** Coding scheme of the genotypic matrix for allelic combinations. Genotypic codes (0, 1, 2) used for testing additive and non-additive effects (dominant, recessive, overdominant). Aref = reference allele; Aalt = alternative allele.

| Allelic combination | Additive effects | Dominant effects | Recessive effects | Overdominant effects |
|---|---|---|---|---|
| Aref-Aref | 0 | 0 | 0 | 0 |
| Aref-Aalt | 1 | 2 | 0 | 1 |
| Aalt-Aalt | 2 | 2 | 2 | 0 |

**Supplementary Table 30.** QCH marginal-kernel parameter estimates for the 13 ancestry-adjusted BLINK P-value distributions. p0 is the estimated proportion of null marker tests. Abbreviations: QCH, Query Composite Hypotheses; BLINK, Bayesian-information and Linkage-disequilibrium Iteratively Nested Keyway; p0, estimated proportion of null marker tests.

| Trait | N markers | p0 initial | p0 kernel |
|---|---|---|---|
| BUD_DATE | 140,181 | 0.901 | 0.918 |
| FLO_50 | 139,738 | 0.989 | 0.999 |
| VER_50 | 139,406 | 0.968 | 0.98 |
| HARVEST_DATE | 140,077 | 0.957 | 0.969 |
| SBER_W_g | 139,638 | 0.871 | 0.869 |
| TSS | 139,300 | 0.92 | 0.915 |
| BERRY_pH | 139,173 | 0.823 | 0.823 |
| BER_TA_g | 139,164 | 0.868 | 0.862 |
| MORPHO_OIV_204 | 139,223 | 0.932 | 0.941 |
| NB_CLUST_PLANT | 138,294 | 0.869 | 0.869 |
| YIELD_PLANT | 137,392 | 0.967 | 0.965 |
| SCLUST_W | 137,285 | 0.84 | 0.835 |
| YIELD_OIV_504 | 139,102 | 0.952 | 0.956 |

**Supplementary Table 31.** Summary of phenotyping environments by population. Overview of the environments (location x year combinations) in which each population was phenotyped. "ResDur" refers to all individuals from step 2 of the breeding program.

| Population ID | Angers | Bordeaux | Colmar | Montpellier | Pully |
|---|---|---|---|---|---|
| 42050 | 2011, 2012 | 2011, 2012, 2013, 2014, 2015, 2016 | 2011, 2012, 2013, 2014 | | |
| 50001 | 2006, 2007, 2008, 2009 | 2006, 2007, 2008, 2009 | 2007, 2008, 2009 | | |
| 50013 | 2010, 2011, 2012, 2013, 2014 | 2011, 2012, 2013, 2014 | 2010, 2011, 2012, 2013, 2014 | | |
| 50015 | 2010, 2011, 2012, 2013 | 2011, 2012, 2013, 2014 | 2010, 2011, 2012, 2013 | | |
| 50025 | | | 2018, 2019, 2020 | | 2017, 2018, 2019, 2020, 2021 |
| 50035 | | | 2019, 2020, 2021, 2022 | | 2018, 2019, 2020, 2021, 2022 |
| ResDur | From 2006 to 2014 | From 2006 to 2017 | From 2007 to 2024 | 2015, 2016 | From 2014 to 2022 |

**Supplementary Table 32.** LD-defined marker-block sensitivity audit for localGEBV. The primary setting was within-population-centered LD with r² = 0.30 and tolerance = 1; pooled LD and alternative thresholds are sensitivity analyses. Abbreviations: LD, linkage disequilibrium; localGEBV, local genomic estimated breeding value; SNP, single-nucleotide polymorphism.

| ld source | setting | threshold | tolerance | N blocks | Mean SNP per Block | Max SNP per Block | Percent Singleton Blocks | Mean Block Size | Max Block Size | marker coverage ok |
|---|---|---|---|---|---|---|---|---|---|---|
| pooled | r2_0.1_tol_0 | 0.1 | 0 | 5,050 | 1.59 | 15 | 68 | 88,649 | 2,168,279 | TRUE |
| pooled | r2_0.1_tol_1 | 0.1 | 1 | 3,355 | 2.4 | 30 | 62 | 183,998 | 6,565,899 | TRUE |
| pooled | r2_0.1_tol_2 | 0.1 | 2 | 2,194 | 3.67 | 57 | 57.2 | 318,704 | 7,104,293 | TRUE |
| pooled | r2_0.1_tol_3 | 0.1 | 3 | 1,517 | 5.31 | 84 | 53.8 | 492,779 | 7,670,887 | TRUE |
| pooled | r2_0.3_tol_0 | 0.3 | 0 | 6,028 | 1.34 | 9 | 77.3 | 61,869 | 1,917,361 | TRUE |
| pooled | r2_0.3_tol_1 | 0.3 | 1 | 4,846 | 1.66 | 20 | 74.7 | 113,438 | 3,472,911 | TRUE |
| pooled | r2_0.3_tol_2 | 0.3 | 2 | 3,779 | 2.13 | 40 | 72 | 189,738 | 6,208,590 | TRUE |
| pooled | r2_0.3_tol_3 | 0.3 | 3 | 3,027 | 2.66 | 84 | 70.6 | 280,620 | 6,992,666 | TRUE |
| pooled | r2_0.5_tol_0 | 0.5 | 0 | 6,388 | 1.26 | 9 | 81.2 | 52,001 | 1,642,428 | TRUE |
| pooled | r2_0.5_tol_1 | 0.5 | 1 | 5,499 | 1.46 | 19 | 79.9 | 88,022 | 2,083,038 | TRUE |
| pooled | r2_0.5_tol_2 | 0.5 | 2 | 4,590 | 1.75 | 40 | 78.4 | 148,022 | 3,575,062 | TRUE |
| pooled | r2_0.5_tol_3 | 0.5 | 3 | 3,881 | 2.08 | 84 | 77.4 | 217,959 | 5,180,570 | TRUE |
| pooled | r2_0.2_tol_4 | 0.2 | 4 | 1,835 | 4.39 | 104 | 62.2 | 470,586 | 9,060,520 | TRUE |
| within_population | r2_0.1_tol_0 | 0.1 | 0 | 4,855 | 1.66 | 13 | 65.7 | 89,567 | 2,168,279 | TRUE |
| within_population | r2_0.1_tol_1 | 0.1 | 1 | 3,041 | 2.65 | 35 | 58.6 | 197,075 | 8,170,314 | TRUE |
| within_population | r2_0.1_tol_2 | 0.1 | 2 | 1,886 | 4.27 | 53 | 52.6 | 365,580 | 9,364,429 | TRUE |
| within_population | r2_0.1_tol_3 | 0.1 | 3 | 1,236 | 6.52 | 97 | 46.8 | 555,488 | 9,364,429 | TRUE |
| within_population | r2_0.3_tol_0 | 0.3 | 0 | 5,856 | 1.38 | 9 | 75.5 | 65,179 | 1,917,388 | TRUE |
| within_population | r2_0.3_tol_1 | 0.3 | 1 | 4,600 | 1.75 | 21 | 72.5 | 122,747 | 4,898,618 | TRUE |
| within_population | r2_0.3_tol_2 | 0.3 | 2 | 3,406 | 2.36 | 40 | 69 | 208,500 | 6,572,482 | TRUE |
| within_population | r2_0.3_tol_3 | 0.3 | 3 | 2,599 | 3.1 | 84 | 66.3 | 309,336 | 7,396,671 | TRUE |
| within_population | r2_0.5_tol_0 | 0.5 | 0 | 6,320 | 1.27 | 9 | 80.5 | 52,792 | 1,917,361 | TRUE |
| within_population | r2_0.5_tol_1 | 0.5 | 1 | 5,414 | 1.49 | 19 | 79 | 87,144 | 1,917,361 | TRUE |
| within_population | r2_0.5_tol_2 | 0.5 | 2 | 4,458 | 1.81 | 40 | 77.2 | 149,771 | 3,575,062 | TRUE |
| within_population | r2_0.5_tol_3 | 0.5 | 3 | 3,736 | 2.16 | 84 | 75.9 | 218,996 | 5,808,397 | TRUE |
| within_population | r2_0.2_tol_4 | 0.2 | 4 | 1,423 | 5.66 | 90 | 55.7 | 549,139 | 12,247,804 | TRUE |

**Supplementary Table 33.** rrBLUP marker-effect model summary for the 13-trait localGEBV analysis. The reported $h^2$ uses the package's VanRaden scaling and is a model diagnostic, not the family-specific broad-sense $H^2$ in Supplementary Table 4. Abbreviations: rrBLUP, ridge-regression best linear unbiased prediction; localGEBV, local genomic estimated breeding value; $h^2$, marker-model heritability.

| trait | model | n | marker variance | residual variance | additive variance VanRaden | h2 package scaling |
|---|---|---|---|---|---|---|
| BUD_DATE | package_default_rrBLUP | 755 | 4.492E-3 | 2.585E+0 | 7.337E+0 | 0.739 |
| BUD_DATE | population_adjusted_rrBLUP | 755 | 3.371E-3 | 2.752E+0 | 5.505E+0 | 0.667 |
| FLO_50 | package_default_rrBLUP | 754 | 2.001E-4 | 1.816E-1 | 3.27E-1 | 0.643 |
| FLO_50 | population_adjusted_rrBLUP | 754 | 8.62E-5 | 9.777E-2 | 1.409E-1 | 0.59 |
| VER_50 | package_default_rrBLUP | 756 | 2.146E-2 | 6.405E+0 | 3.508E+1 | 0.846 |
| VER_50 | population_adjusted_rrBLUP | 756 | 1.989E-2 | 6.355E+0 | 3.252E+1 | 0.837 |
| SBER_W_g | package_default_rrBLUP | 733 | 4.544E-5 | 1.492E-2 | 7.4E-2 | 0.832 |
| SBER_W_g | population_adjusted_rrBLUP | 733 | 4.053E-5 | 1.544E-2 | 6.599E-2 | 0.81 |
| TSS | package_default_rrBLUP | 722 | 7.389E-4 | 1.843E-1 | 1.2E+0 | 0.867 |
| TSS | population_adjusted_rrBLUP | 722 | 4.302E-4 | 2.009E-1 | 6.988E-1 | 0.777 |
| BERRY_pH | package_default_rrBLUP | 754 | 2.569E-6 | 1.348E-3 | 4.199E-3 | 0.757 |
| BERRY_pH | population_adjusted_rrBLUP | 754 | 1.792E-6 | 1.32E-3 | 2.93E-3 | 0.689 |
| BER_TA_g | package_default_rrBLUP | 754 | 8.459E-5 | 3.907E-2 | 1.383E-1 | 0.78 |
| BER_TA_g | population_adjusted_rrBLUP | 754 | 6.167E-5 | 4.108E-2 | 1.008E-1 | 0.71 |
| HARVEST_DATE | package_default_rrBLUP | 730 | 8.179E-5 | 3.323E-2 | 1.331E-1 | 0.8 |
| HARVEST_DATE | population_adjusted_rrBLUP | 730 | 7.176E-5 | 3.351E-2 | 1.167E-1 | 0.777 |
| MORPHO_OIV_204 | package_default_rrBLUP | 750 | 1.864E-4 | 1.267E-1 | 3.045E-1 | 0.706 |
| MORPHO_OIV_204 | population_adjusted_rrBLUP | 750 | 1.824E-4 | 1.243E-1 | 2.98E-1 | 0.706 |
| NB_CLUST_PLANT | package_default_rrBLUP | 711 | 2.427E-4 | 1.159E-1 | 3.933E-1 | 0.772 |
| NB_CLUST_PLANT | population_adjusted_rrBLUP | 711 | 1.468E-4 | 1.179E-1 | 2.379E-1 | 0.669 |
| YIELD_PLANT | package_default_rrBLUP | 704 | 8.384E-5 | 6.74E-2 | 1.356E-1 | 0.668 |
| YIELD_PLANT | population_adjusted_rrBLUP | 704 | 6.126E-5 | 6.7E-2 | 9.911E-2 | 0.597 |
| SCLUST_W | package_default_rrBLUP | 698 | 1.049E-4 | 6.438E-2 | 1.695E-1 | 0.725 |
| SCLUST_W | population_adjusted_rrBLUP | 698 | 8.496E-5 | 6.463E-2 | 1.373E-1 | 0.68 |

| trait | model | n | marker variance | residual variance | additive variance VanRaden | h2 package scaling |
|---|---|---|---|---|---|---|
| YIELD_OIV_504 | package_default_rrBLUP | 722 | 9.504E-5 | 7.679E-2 | 1.544E-1 | 0.668 |
| YIELD_OIV_504 | population_adjusted_rrBLUP | 722 | 7.61E-5 | 7.806E-2 | 1.237E-1 | 0.613 |

**Supplementary Table 34.** Multiple-testing and evidence-threshold guide. Thresholds were matched to the inferential purpose of each analysis; descriptive FDR values were not substituted for primary family-wise thresholds. Abbreviations: FDR, false discovery rate.

| Analysis | Primary inferential target | Structure adjustment | Primary threshold | Role in evidence hierarchy |
|---|---|---|---|---|
| Single-family QTL | Genome-wide QTL within one biparental family | Family-specific linkage analysis | 5% genome-wide threshold from 4,000 permutations; three family-50015 signals retained at 5-10% as suggestive | Family-specific discovery |
| Pooled BLINK GWAS | Single-trait additive association across the ResDur panel | LD1-LD5 fixed ancestry covariates | 5% trait-specific Bonferroni family-wise error rate | Panel-wide association |
| Pooled MLMM GWAS | Selected multilocus additive cofactors across the ResDur panel | Trait-specific genomic kinship plus LD1-LD5 | Package Bonferroni stopping criterion | Panel-wide association |
| Pooled MM4LMM GWAS | Single-marker additive association across the ResDur panel | Trait-specific genomic kinship plus LD1-LD5 | 5% trait-specific Bonferroni family-wise error rate | Panel-wide association |
| Non-additive BLINK | Single-trait dominant/recessive/overdominant model association | LD1-LD5 fixed ancestry covariates | 5% trait-specific Bonferroni family-wise error rate | Exploratory only |
| Environment-specific BLINK | Single-trait association within a site-year environment | LD1-LD5 fixed ancestry covariates | Inputs to metaGE | Environment-specific input |
| metaGE fixed/random effects | Marker evidence combined across genuine site-year scans | LD1-LD5 in every contributing scan; environment correlation matrix | 5% BH FDR with estimated null proportion | Exploratory environmental evidence |
| metaGE local score | Contiguous regions accumulating meta-analysis evidence | As above | Local-score tuning parameter $\xi$=3 | Exploratory regional evidence |
| metaGE climate regression | Marker-effect covariation with one annual climate descriptor | As above | 5% BH FDR with estimated null proportion | Exploratory environmental evidence |
| mpQTL | Population-adjusted dosage association on the connected-family composite map | Realized kinship plus 12 population indicators | Li-Ji effective-test threshold | Second QTL mapping framework |
| QCH | Composite alternative of association with at least two traits | Uses P values from the LD1-LD5-adjusted BLINK scans | 5% QCH FDR by ordered posterior local-FDR estimates | Cross-trait pleiotropy hypothesis |

| Analysis | Primary inferential target | Structure adjustment | Primary threshold | Role in evidence hierarchy |
|---|---|---|---|---|
| Figure 5B trait values | Descriptive trait-level strength at QCH-selected loci | Uses P values from the LD1-LD5-adjusted BLINK scans | Benjamini-Hochberg values shown descriptively | Not used to select QCH loci |
| LocalGEBV | LD-block breeding-value ranking and prediction | Population fixed effects; within-population-centered LD; family-held-out validation | No association-significance threshold | Breeding-prioritization layer |

**Supplementary Table 35.** Descriptive 250-kb proximity groups of marker-climate meta-regression tests passing 5% FDR. Climate covariates were tested separately and are correlated; groups are not independent causal effects. Abbreviations: FDR, false discovery rate.

| Trait | Climate covariate | CHR | Start | End | N markers | Lead marker | Lead FDR |
|---|---|---|---|---|---|---|---|
| BERRY_pH | annual_rainfall_mm | 16 | 14,486,694 | 14,486,694 | 1 | chr16_14486694 | 6.494E-20 |
| BERRY_pH | mean_Tmax_C | 16 | 14,486,694 | 14,486,694 | 1 | chr16_14486694 | 4.267E-10 |
| BUD_DATE | mean_Tmean_C | 14 | 5,148,380 | 5,148,380 | 1 | chr14_5148380 | 3.343E-2 |
| BUD_DATE | mean_Tmean_C | 14 | 27,574,173 | 27,574,173 | 1 | chr14_27574173 | 1.121E-2 |
| FLO_50 | annual_rainfall_mm | 2 | 14,734,609 | 14,734,609 | 1 | chr2_14734609 | 4.001E-2 |
| FLO_50 | annual_rainfall_mm | 12 | 15,810,422 | 15,810,422 | 1 | chr12_15810422 | 4.95E-2 |
| FLO_50 | annual_rainfall_mm | 13 | 621,827 | 2,437,579 | 53 | chr13_621827 | 4.001E-2 |
| FLO_50 | mean_Tmax_C | 13 | 926,764 | 926,773 | 2 | chr13_926764 | 1.972E-2 |
| FLO_50 | mean_Tmax_C | 15 | 3,107,296 | 3,107,296 | 1 | chr15_3107296 | 2.571E-2 |
| FLO_50 | mean_Tmax_C | 17 | 5,603,726 | 5,603,726 | 1 | chr17_5603726 | 2.571E-2 |
| FLO_50 | mean_Tmin_C | 2 | 4,100,872 | 4,100,872 | 1 | chr2_4100872 | 3.56E-2 |
| FLO_50 | mean_Tmin_C | 3 | 4,121,601 | 4,121,601 | 1 | chr3_4121601 | 3.624E-2 |
| FLO_50 | mean_Tmin_C | 7 | 3,577,578 | 3,577,589 | 2 | chr7_3577589 | 3.913E-2 |
| FLO_50 | mean_Tmin_C | 7 | 5,224,824 | 5,224,824 | 1 | chr7_5224824 | 2.94E-2 |
| FLO_50 | mean_Tmin_C | 11 | 790,317 | 1,033,669 | 4 | chr11_886579 | 1.389E-2 |
| FLO_50 | mean_Tmin_C | 12 | 9,974,787 | 9,974,787 | 1 | chr12_9974787 | 3.976E-2 |
| FLO_50 | mean_Tmin_C | 13 | 10,752 | 4,566,106 | 397 | chr13_2437560 | 8.651E-3 |
| FLO_50 | mean_Tmin_C | 13 | 5,329,816 | 5,329,816 | 1 | chr13_5329816 | 2.761E-2 |
| FLO_50 | mean_Tmin_C | 13 | 22,324,134 | 22,324,134 | 1 | chr13_22324134 | 4.772E-2 |
| FLO_50 | mean_Tmin_C | 17 | 7,355,018 | 7,355,018 | 1 | chr17_7355018 | 4.886E-2 |
| FLO_50 | mean_Tmin_C | 18 | 4,523,957 | 4,523,957 | 1 | chr18_4523957 | 4.757E-2 |
| HARVEST_DATE | annual_rainfall_mm | 12 | 14,431,827 | 14,431,827 | 1 | chr12_14431827 | 1.689E-7 |
| HARVEST_DATE | mean_ET0_mm_d | 14 | 24,420,726 | 24,420,726 | 1 | chr14_24420726 | 1.377E-2 |
| HARVEST_DATE | mean_ET0_mm_d | 14 | 26,168,835 | 26,168,858 | 2 | chr14_26168835 | 1.377E-2 |
| HARVEST_DATE | mean_Tmax_C | 17 | 7,915,699 | 7,915,699 | 1 | chr17_7915699 | 2.673E-2 |

| Trait | Climate covariate | CHR | Start | End | N markers | Lead marker | Lead FDR |
|---|---|---|---|---|---|---|---|
| HARVEST_DATE | mean_Tmean_C | 14 | 24,420,726 | 24,420,726 | 1 | chr14_24420726 | 1.626E-2 |
| HARVEST_DATE | mean_Tmin_C | 14 | 24,420,726 | 24,420,726 | 1 | chr14_24420726 | 5.482E-5 |
| HARVEST_DATE | mean_Tmin_C | 14 | 26,168,858 | 26,168,858 | 1 | chr14_26168858 | 3.667E-2 |
| HARVEST_DATE | mean_Tmin_C | 14 | 26,922,503 | 26,922,503 | 1 | chr14_26922503 | 3.667E-2 |
| TSS | mean_Tmax_C | 5 | 506,800 | 506,800 | 1 | chr5_506800 | 2.31E-2 |
| VER_50 | mean_ET0_mm_d | 11 | 8,950,582 | 8,950,582 | 1 | chr11_8950582 | 1.084E-2 |
| VER_50 | mean_ET0_mm_d | 11 | 11,191,818 | 11,191,818 | 1 | chr11_11191818 | 3.266E-2 |
| VER_50 | mean_ET0_mm_d | 16 | 14,752,071 | 14,752,071 | 1 | chr16_14752071 | 1.084E-2 |
| VER_50 | mean_Tmean_C | 2 | 15,363,642 | 15,363,642 | 1 | chr2_15363642 | 4.694E-2 |
| VER_50 | mean_Tmean_C | 2 | 16,362,141 | 16,362,141 | 1 | chr2_16362141 | 3.082E-2 |
| VER_50 | mean_Tmean_C | 11 | 8,273,804 | 8,418,794 | 4 | chr11_8273804 | 3.082E-2 |
| VER_50 | mean_Tmean_C | 11 | 8,950,012 | 8,986,415 | 3 | chr11_8950582 | 5.211E-3 |
| VER_50 | mean_Tmean_C | 16 | 14,752,071 | 14,752,071 | 1 | chr16_14752071 | 3.082E-2 |
| VER_50 | mean_Tmin_C | 11 | 8,273,804 | 8,418,794 | 4 | chr11_8273804 | 3.712E-2 |
| VER_50 | mean_Tmin_C | 11 | 8,950,012 | 8,950,582 | 2 | chr11_8950582 | 7.918E-3 |
| VER_50 | mean_Tmin_C | 11 | 11,191,818 | 11,191,818 | 1 | chr11_11191818 | 4.708E-2 |

**Supplementary Table 36.** Trait-specific input summary for the 303 genuine site-by-year ancestry-adjusted GWAS included in metaGE. Abbreviations: GWAS, genome-wide association study; metaGE, meta-analysis of genotype-environment association results; DAPC, discriminant analysis of principal components.

| Trait | N environments | first year | last year | N sites | sites | N individuals min | N individuals max | N markers min | N markers max | DAPC axes |
|---|---|---|---|---|---|---|---|---|---|---|
| BERRY_pH | 25 | 2011 | 2024 | 4 | Angers;Bordeaux;Colmar;Pully | 123 | 391 | 60,271 | 132,644 | 5 |
| BER_TA_g | 26 | 2011 | 2024 | 4 | Angers;Bordeaux;Colmar;Pully | 123 | 391 | 60,271 | 132,644 | 5 |
| BUD_DATE | 21 | 2012 | 2024 | 4 | Angers;Bordeaux;Colmar;Pully | 100 | 349 | 60,273 | 132,352 | 5 |
| FLO_50 | 20 | 2010 | 2022 | 4 | Angers;Bordeaux;Colmar;Pully | 112 | 344 | 60,273 | 137,721 | 5 |
| HARVEST_DATE | 24 | 2011 | 2024 | 4 | Angers;Bordeaux;Colmar;Pully | 158 | 391 | 60,273 | 132,644 | 5 |
| MORPHO_OIV_204 | 20 | 2011 | 2024 | 3 | Angers;Bordeaux;Colmar | 123 | 374 | 107,634 | 132,642 | 5 |
| NB_CLUST_PLANT | 20 | 2011 | 2024 | 3 | Angers;Bordeaux;Colmar | 123 | 380 | 107,875 | 133,984 | 5 |
| SBER_W_g | 24 | 2011 | 2024 | 4 | Angers;Bordeaux;Colmar;Pully | 123 | 382 | 60,273 | 133,188 | 5 |
| SCLUST_W | 26 | 2011 | 2024 | 4 | Angers;Bordeaux;Colmar;Pully | 123 | 379 | 60,273 | 133,163 | 5 |
| TSS | 25 | 2011 | 2024 | 4 | Angers;Bordeaux;Colmar;Pully | 123 | 388 | 60,271 | 134,000 | 5 |
| VER_50 | 28 | 2010 | 2024 | 4 | Angers;Bordeaux;Colmar;Pully | 110 | 359 | 60,204 | 148,108 | 5 |
| YIELD_OIV_504 | 24 | 2011 | 2022 | 4 | Angers;Bordeaux;Colmar;Pully | 123 | 426 | 60,273 | 133,984 | 5 |
| YIELD_PLANT | 20 | 2011 | 2024 | 3 | Angers;Bordeaux;Colmar | 123 | 426 | 104,206 | 133,984 | 5 |
