## Supplementary File 1 for "Uncovering the genetic basis of agronomic traits in over 1,000 grapevine genotypes derived from a disease resistance breeding program"

Borrelli et al.: *Uncovering the genetic basis of agronomic traits in over 1,000 grapevine genotypes derived from a disease resistance breeding program*

DAPC settings for Figure 1C, Figure 2 reference varieties, robustness of the phenotypic correlations and QCH sensitivity analyses

### A. DAPC settings and eigenvalues (Figure 1C)

The DAPC shown in Figure 1C was fitted to 344 individuals (194 *V. vinifera* accessions, 122 other interspecific hybrids, 15 ResDur parents and 13 progeny) genotyped at 62,619 polymorphic LD-clumped SNPs (alternate-allele dosages, marker-centred). Fitting groups were seven clusters inferred from the genotypes by K-means on the principal components and selected by the Bayesian information criterion (K = 1-40). The number of principal components (PCs) retained for the discriminant step was chosen by stratified cross-validation of cluster assignment (100 resamples, two-thirds of the individuals in training): the smallest number of PCs within one standard error of the maximum mean assignment success was nine, explaining 31.1% of the genotypic variance. All six discriminant axes were retained; LD1 and LD2 carry 61.3% and 12.9% of the sum of the discriminant eigenvalues. The four biological categories are plotting annotations and were not used to fit the model. The ancestry covariates used in the GWAS come from a separate DAPC (1,389 genotypes, 36,726 SNPs, 16 inferred clusters, 15 PCs, five discriminant axes) and were not modified. Software: R 4.5.1, adegenet 2.1.11.

**Table S5-1. Eigenvalues of the nine retained principal components.**

| PC | Eigenvalue | Variance (%) | Cumulative (%) |
|---|---|---|---|
| 1 | 1,400.6 | 14.19 | 14.19 |
| 2 | 326.3 | 3.31 | 17.50 |
| 3 | 265.5 | 2.69 | 20.19 |
| 4 | 233.0 | 2.36 | 22.55 |
| 5 | 211.5 | 2.14 | 24.69 |
| 6 | 187.2 | 1.90 | 26.59 |
| 7 | 163.0 | 1.65 | 28.24 |
| 8 | 148.6 | 1.51 | 29.74 |
| 9 | 133.6 | 1.35 | 31.10 |

**Table S5-2. Eigenvalues of the six discriminant axes.**

| Axis | DA eigenvalue | Share of DA eigenvalue sum (%) |
|---|---|---|
| 1 | 867.4 | 61.3 |
| 2 | 182.2 | 12.9 |
| 3 | 144.2 | 10.2 |
| 4 | 97.3 | 6.9 |
| 5 | 75.2 | 5.3 |
| 6 | 48.6 | 3.4 |

### B. Reference varieties shown in Figure 2

The INRAE-ResDur programme comprises successive phases that differ in the resistance factors pyramided. The dashed reference lines in Figure 2 are the means of the raw observations recorded, during the intermediate field-trial step of the programme, for the four varieties registered from phase 1 (2006-2022; Angers, Bordeaux and Colmar) and the eight varieties registered from phase 2 (2010-2016; Angers, Bordeaux, Colmar and Montpellier). The years and sites contributing to each variety and trait vary, so these means are descriptive benchmarks rather than environmentally matched comparisons; the family-effect analysis of the manuscript used the environment-adjusted whole-panel BLUPs instead. As in the original analysis, four low-sugar and two high-yield observations were excluded from the phase-1 values. Trait-specific means, numbers of observations, years and sites are listed in the Supplementary Excel File (worksheet "Figure2 reference means").

**Table S5-3. Registered varieties used as reference in Figure 2.**

| ResDur phase | Variety |
|---|---|
| 1 | Artaban |
| 1 | Floreal |
| 1 | Vidoc |
| 1 | Voltis |
| 2 | Artys |
| 2 | Calys |
| 2 | Coliris |
| 2 | Exelys |
| 2 | Lilaro |
| 2 | Opalor |
| 2 | Selenor |
| 2 | Sirano |

### C. Robustness of the phenotypic correlations (Figure 3)

Pearson correlations in Figure 3 were calculated from 1,671 complete raw records of 863 ResDur genotypes, so that genotypes evaluated in several environments contributed repeated records (one to seven per genotype). The same records were used to recalculate the correlations (i) among genotype means, giving each genotype equal weight, and (ii) among the environment-adjusted whole-panel BLUPs of the same 863 genotypes; (iii) 95% percentile intervals of the raw correlations were obtained from 2,000 bootstrap resamples drawn by genotype, each resample retaining all records of the sampled genotypes. The BLUP correlations are associations among predicted genotypic values, not genetic correlations estimated by a multivariate model. Yield per area equals yield per vine divided by the area per vine of the plot (2 or 2.329 $m^2$), which explains the correlation of 0.998 displayed as 1.00 in Figure 3.

**Table S5-4. Pairwise correlations among the 13 traits from raw records (with 95% genotype-bootstrap interval), genotype means and BLUPs.**

| Trait 1 | Trait 2 | Raw r | 95% interval | Genotype-mean r | BLUP r |
|---|---|---|---|---|---|
| Budbreak | Flowering | 0.249 | 0.204 to 0.290 | 0.303 | 0.434 |
| Budbreak | Veraison | 0.111 | 0.063 to 0.156 | 0.186 | 0.298 |
| Flowering | Veraison | 0.720 | 0.694 to 0.746 | 0.706 | 0.441 |
| Budbreak | Harvest date | 0.129 | 0.085 to 0.171 | 0.211 | 0.302 |
| Flowering | Harvest date | 0.621 | 0.589 to 0.651 | 0.620 | 0.404 |
| Veraison | Harvest date | 0.909 | 0.901 to 0.917 | 0.930 | 0.951 |
| Budbreak | Berry weight | 0.070 | 0.031 to 0.107 | 0.102 | 0.267 |
| Flowering | Berry weight | 0.313 | 0.217 to 0.427 | 0.385 | 0.068 |
| Veraison | Berry weight | 0.294 | 0.194 to 0.419 | 0.332 | 0.224 |
| Harvest date | Berry weight | 0.264 | 0.174 to 0.380 | 0.330 | 0.231 |
| Budbreak | Titratable acidity | 0.057 | 0.013 to 0.106 | 0.143 | 0.010 |
| Flowering | Titratable acidity | 0.300 | 0.265 to 0.335 | 0.283 | -0.035 |
| Veraison | Titratable acidity | 0.453 | 0.410 to 0.494 | 0.458 | 0.364 |
| Harvest date | Titratable acidity | 0.363 | 0.317 to 0.409 | 0.367 | 0.395 |
| Berry weight | Titratable acidity | 0.135 | 0.084 to 0.208 | 0.169 | 0.241 |
| Budbreak | Sugar content | -0.018 | -0.067 to 0.028 | -0.094 | -0.055 |
| Flowering | Sugar content | -0.268 | -0.318 to -0.218 | -0.281 | 0.008 |
| Veraison | Sugar content | -0.276 | -0.322 to -0.223 | -0.275 | -0.218 |
| Harvest date | Sugar content | -0.245 | -0.293 to -0.194 | -0.237 | -0.291 |
| Berry weight | Sugar content | -0.178 | -0.259 to -0.127 | -0.202 | -0.365 |
| Titratable acidity | Sugar content | -0.227 | -0.279 to -0.171 | -0.210 | -0.343 |
| Budbreak | Berry pH | 0.062 | 0.006 to 0.119 | 0.030 | 0.055 |
| Flowering | Berry pH | -0.105 | -0.156 to -0.057 | 0.108 | 0.098 |
| Veraison | Berry pH | -0.157 | -0.215 to -0.100 | -0.014 | 0.159 |
| Harvest date | Berry pH | -0.178 | -0.236 to -0.122 | -0.043 | 0.164 |
| Berry weight | Berry pH | 0.060 | 0.007 to 0.135 | 0.073 | 0.232 |
| Titratable acidity | Berry pH | -0.504 | -0.547 to -0.458 | -0.411 | -0.054 |
| Sugar content | Berry pH | 0.291 | 0.244 to 0.338 | 0.211 | -0.415 |
| Budbreak | Compactness | 0.026 | -0.021 to 0.079 | 0.058 | 0.075 |
| Flowering | Compactness | -0.035 | -0.086 to 0.013 | -0.034 | -0.179 |
| Veraison | Compactness | -0.023 | -0.081 to 0.034 | -0.019 | -0.048 |
| Harvest date | Compactness | -0.011 | -0.063 to 0.042 | 0.015 | 0.013 |
| Berry weight | Compactness | -0.005 | -0.050 to 0.052 | 0.009 | 0.075 |
| Titratable acidity | Compactness | -0.069 | -0.121 to -0.020 | -0.065 | 0.054 |
| Sugar content | Compactness | -0.149 | -0.202 to -0.094 | -0.170 | -0.179 |
| Berry pH | Compactness | -0.024 | -0.076 to 0.028 | -0.114 | -0.065 |
| Budbreak | Clusters per plant | 0.048 | -0.001 to 0.098 | -0.108 | -0.121 |
| Flowering | Clusters per plant | -0.377 | -0.423 to -0.328 | -0.483 | -0.229 |
| Veraison | Clusters per plant | -0.459 | -0.502 to -0.414 | -0.544 | -0.424 |
| Harvest date | Clusters per plant | -0.325 | -0.372 to -0.273 | -0.436 | -0.410 |
| Berry weight | Clusters per plant | -0.318 | -0.420 to -0.243 | -0.358 | -0.384 |
| Titratable acidity | Clusters per plant | -0.344 | -0.391 to -0.299 | -0.403 | -0.600 |

| | | | | | |
|---|---|---|---|---|---|
| Sugar content | Clusters per plant | 0.056 | 0.003 to 0.114 | 0.088 | 0.465 |
| Berry pH | Clusters per plant | -0.099 | -0.151 to -0.044 | -0.178 | -0.477 |
| Compactness | Clusters per plant | 0.129 | 0.080 to 0.176 | 0.174 | 0.079 |
| Budbreak | Mean cluster weight | -0.142 | -0.196 to -0.089 | -0.079 | 0.178 |
| Flowering | Mean cluster weight | 0.025 | -0.026 to 0.072 | 0.035 | 0.038 |
| Veraison | Mean cluster weight | 0.048 | -0.008 to 0.102 | 0.014 | -0.111 |
| Harvest date | Mean cluster weight | 0.106 | 0.053 to 0.159 | 0.107 | -0.075 |
| Berry weight | Mean cluster weight | 0.298 | 0.232 to 0.394 | 0.371 | 0.145 |
| Titratable acidity | Mean cluster weight | -0.070 | -0.119 to -0.021 | -0.121 | -0.421 |
| Sugar content | Mean cluster weight | -0.259 | -0.311 to -0.206 | -0.247 | 0.204 |
| Berry pH | Mean cluster weight | -0.092 | -0.148 to -0.036 | -0.105 | -0.390 |
| Compactness | Mean cluster weight | 0.350 | 0.303 to 0.399 | 0.403 | 0.316 |
| Clusters per plant | Mean cluster weight | -0.009 | -0.059 to 0.046 | 0.101 | 0.545 |
| Budbreak | Yield per vine | -0.020 | -0.065 to 0.027 | -0.091 | -0.005 |
| Flowering | Yield per vine | -0.198 | -0.249 to -0.144 | -0.246 | -0.222 |
| Veraison | Yield per vine | -0.246 | -0.293 to -0.196 | -0.321 | -0.321 |
| Harvest date | Yield per vine | -0.106 | -0.153 to -0.056 | -0.181 | -0.276 |
| Berry weight | Yield per vine | -0.022 | -0.070 to 0.022 | 0.008 | -0.072 |
| Titratable acidity | Yield per vine | -0.284 | -0.326 to -0.241 | -0.362 | -0.486 |
| Sugar content | Yield per vine | -0.160 | -0.210 to -0.107 | -0.130 | 0.268 |
| Berry pH | Yield per vine | -0.155 | -0.197 to -0.110 | -0.192 | -0.416 |
| Compactness | Yield per vine | 0.328 | 0.286 to 0.368 | 0.386 | 0.318 |
| Clusters per plant | Yield per vine | 0.675 | 0.643 to 0.706 | 0.705 | 0.830 |
| Mean cluster weight | Yield per vine | 0.646 | 0.610 to 0.681 | 0.706 | 0.835 |
| Budbreak | Yield per $m^2$ | -0.048 | -0.093 to -0.001 | -0.105 | 0.004 |
| Flowering | Yield per $m^2$ | -0.194 | -0.244 to -0.140 | -0.239 | -0.217 |
| Veraison | Yield per $m^2$ | -0.237 | -0.286 to -0.187 | -0.311 | -0.284 |
| Harvest date | Yield per $m^2$ | -0.103 | -0.150 to -0.053 | -0.174 | -0.238 |
| Berry weight | Yield per $m^2$ | -0.016 | -0.062 to 0.030 | 0.013 | -0.014 |
| Titratable acidity | Yield per $m^2$ | -0.281 | -0.324 to -0.237 | -0.357 | -0.451 |
| Sugar content | Yield per $m^2$ | -0.169 | -0.220 to -0.116 | -0.138 | 0.225 |
| Berry pH | Yield per $m^2$ | -0.156 | -0.199 to -0.109 | -0.191 | -0.399 |
| Compactness | Yield per $m^2$ | 0.338 | 0.297 to 0.379 | 0.391 | 0.329 |
| Clusters per plant | Yield per $m^2$ | 0.660 | 0.627 to 0.692 | 0.696 | 0.784 |
| Mean cluster weight | Yield per $m^2$ | 0.666 | 0.632 to 0.699 | 0.716 | 0.866 |
| Yield per vine | Yield per $m^2$ | 0.998 | 0.998 to 0.999 | 0.999 | 0.973 |

### D. QCH sensitivity analyses

The cross-trait analysis selected 270 markers grouped into 54 display regions (Figure 5; Supplementary Table 21). Because mean cluster weight is the ratio of cluster weight per plant to cluster number, and yield per area is derived from yield per vine, QCH was refitted on the same 136,062 SNPs after excluding mean cluster weight (S1, 12 traits) and after additionally excluding yield per area (S2, 11 traits), with the Gaussian copula, the alternative of association with at least two of the retained traits and the 5% posterior-FDR procedure (qch 2.1.3). These fits used the package-default stopping threshold of the EM algorithm ($10^{-6}$) instead of the $10^{-4}$ threshold of the primary fit; a 13-trait fit with the same settings (S0) served as control and reselected all 270 markers of the primary catalogue. A primary region is counted as retained when at least one of its originally selected markers is reselected; new signals within 250 kb are not counted. Marker-level results of the three fits are provided in the Supplementary Excel File (worksheet “QCH selected markers”).

**Table S5-5. Summary of the QCH fits.**

| Fit | Traits analysed | n traits | Selected markers | Display loci | Primary markers reselected | Primary regions retained |
|---|---|---|---|---|---|---|
| S0 | All 13 traits (control) | 13 | 4,223 | 412 | 270 / 270 | 54 / 54 |
| S1 | Without mean cluster weight | 12 | 2,940 | 337 | 258 / 270 | 53 / 54 |
| S2 | Without mean cluster weight and yield per area | 11 | 1,236 | 229 | 77 / 270 | 38 / 54 |

**Table S5-6. Retention of the 54 primary QCH regions (Supplementary Table 21) in the sensitivity analyses. Entries are the numbers of originally selected markers reselected in each fit (0 = region not retained).**

| Region | Chr | Position (bp) | Markers | Lead marker | S0 | S1 | S2 |
|---|---|---|---|---|---|---|---|
| 1 | 2 | 1,266,852 | 1 | chr2_1266852 | 1 | 1 | 0 |
| 2 | 5 | 53,402-2,192,309 | 110 | chr5_1219726 | 110 | 107 | 18 |
| 3 | 5 | 2,482,576-3,814,052 | 97 | chr5_2725797 | 97 | 95 | 20 |
| 4 | 5 | 14,801,029 | 1 | chr5_14801029 | 1 | 1 | 1 |
| 5 | 5 | 16,520,934 | 1 | chr5_16520934 | 1 | 1 | 0 |
| 6 | 5 | 16,970,241 | 1 | chr5_16970241 | 1 | 1 | 1 |
| 7 | 5 | 20,538,484 | 1 | chr5_20538484 | 1 | 1 | 1 |
| 8 | 6 | 3,174,778 | 1 | chr6_3174778 | 1 | 1 | 1 |
| 9 | 6 | 5,833,646-5,833,695 | 3 | chr6_5833695 | 3 | 3 | 3 |

| | | | | | | | |
|---|---|---|---|---|---|---|---|
| 10 | 7 | 3,983,389 | 1 | chr7_3983389 | 1 | 1 | 1 |
| 11 | 8 | 10,980,744 | 1 | chr8_10980744 | 1 | 1 | 1 |
| 12 | 8 | 15,522,696 | 1 | chr8_15522696 | 1 | 0 | 0 |
| 13 | 8 | 21,647,686 | 1 | chr8_21647686 | 1 | 1 | 0 |
| 14 | 9 | 915,526 | 1 | chr9_915526 | 1 | 1 | 1 |
| 15 | 9 | 2,793,911 | 1 | chr9_2793911 | 1 | 1 | 1 |
| 16 | 9 | 6,246,031-6,246,067 | 2 | chr9_6246031 | 2 | 2 | 2 |
| 17 | 9 | 6,629,813 | 1 | chr9_6629813 | 1 | 1 | 1 |
| 18 | 10 | 8,361,332-8,361,333 | 2 | chr10_8361333 | 2 | 2 | 1 |
| 19 | 10 | 12,758,127 | 1 | chr10_12758127 | 1 | 1 | 0 |
| 20 | 10 | 19,736,493 | 1 | chr10_19736493 | 1 | 1 | 1 |
| 21 | 11 | 1,033,705 | 1 | chr11_1033705 | 1 | 1 | 0 |
| 22 | 11 | 3,758,688-3,824,935 | 2 | chr11_3758688 | 2 | 2 | 1 |
| 23 | 11 | 4,574,451 | 1 | chr11_4574451 | 1 | 1 | 1 |
| 24 | 11 | 6,089,413 | 1 | chr11_6089413 | 1 | 1 | 1 |
| 25 | 11 | 7,023,371 | 1 | chr11_7023371 | 1 | 1 | 1 |
| 26 | 11 | 8,070,948-8,149,404 | 6 | chr11_8149404 | 6 | 1 | 0 |
| 27 | 12 | 17,932,593-18,014,166 | 2 | chr12_17932593 | 2 | 1 | 1 |
| 28 | 14 | 3,039,299 | 1 | chr14_3039299 | 1 | 1 | 0 |
| 29 | 14 | 7,735,317 | 1 | chr14_7735317 | 1 | 1 | 1 |
| 30 | 14 | 13,952,316 | 1 | chr14_13952316 | 1 | 1 | 0 |
| 31 | 14 | 16,274,319 | 1 | chr14_16274319 | 1 | 1 | 0 |
| 32 | 14 | 17,151,022 | 1 | chr14_17151022 | 1 | 1 | 1 |
| 33 | 14 | 19,079,458 | 1 | chr14_19079458 | 1 | 1 | 1 |
| 34 | 14 | 26,369,635 | 1 | chr14_26369635 | 1 | 1 | 0 |
| 35 | 15 | 4,576,440 | 1 | chr15_4576440 | 1 | 1 | 1 |
| 36 | 16 | 1,101,440 | 1 | chr16_1101440 | 1 | 1 | 1 |
| 37 | 16 | 12,507,629 | 1 | chr16_12507629 | 1 | 1 | 0 |
| 38 | 16 | 14,549,584 | 1 | chr16_14549584 | 1 | 1 | 1 |
| 39 | 16 | 15,035,128 | 1 | chr16_15035128 | 1 | 1 | 1 |
| 40 | 16 | 17,401,013 | 1 | chr16_17401013 | 1 | 1 | 1 |
| 41 | 17 | 6,512,440 | 1 | chr17_6512440 | 1 | 1 | 1 |
| 42 | 18 | 2,100,422 | 1 | chr18_2100422 | 1 | 1 | 1 |
| 43 | 18 | 2,440,997 | 1 | chr18_2440997 | 1 | 1 | 0 |
| 44 | 18 | 4,386,621 | 1 | chr18_4386621 | 1 | 1 | 0 |
| 45 | 18 | 5,141,487 | 1 | chr18_5141487 | 1 | 1 | 1 |
| 46 | 18 | 7,597,972 | 1 | chr18_7597972 | 1 | 1 | 1 |
| 47 | 18 | 9,529,420 | 1 | chr18_9529420 | 1 | 1 | 0 |
| 48 | 18 | 11,131,797 | 1 | chr18_11131797 | 1 | 1 | 1 |
| 49 | 18 | 23,048,559 | 1 | chr18_23048559 | 1 | 1 | 1 |
| 50 | 18 | 34,522,453 | 1 | chr18_34522453 | 1 | 1 | 1 |
| 51 | 19 | 3,309,782 | 1 | chr19_3309782 | 1 | 1 | 1 |
| 52 | 19 | 5,886,348 | 1 | chr19_5886348 | 1 | 1 | 0 |
| 53 | 19 | 13,212,585 | 1 | chr19_13212585 | 1 | 1 | 1 |
| 54 | 19 | 17,074,417 | 1 | chr19_17074417 | 1 | 1 | 1 |
