## Supplementary File 5 for "Uncovering the genetic basis of agronomic traits in over 1,000 grapevine genotypes derived from a disease resistance breeding program"

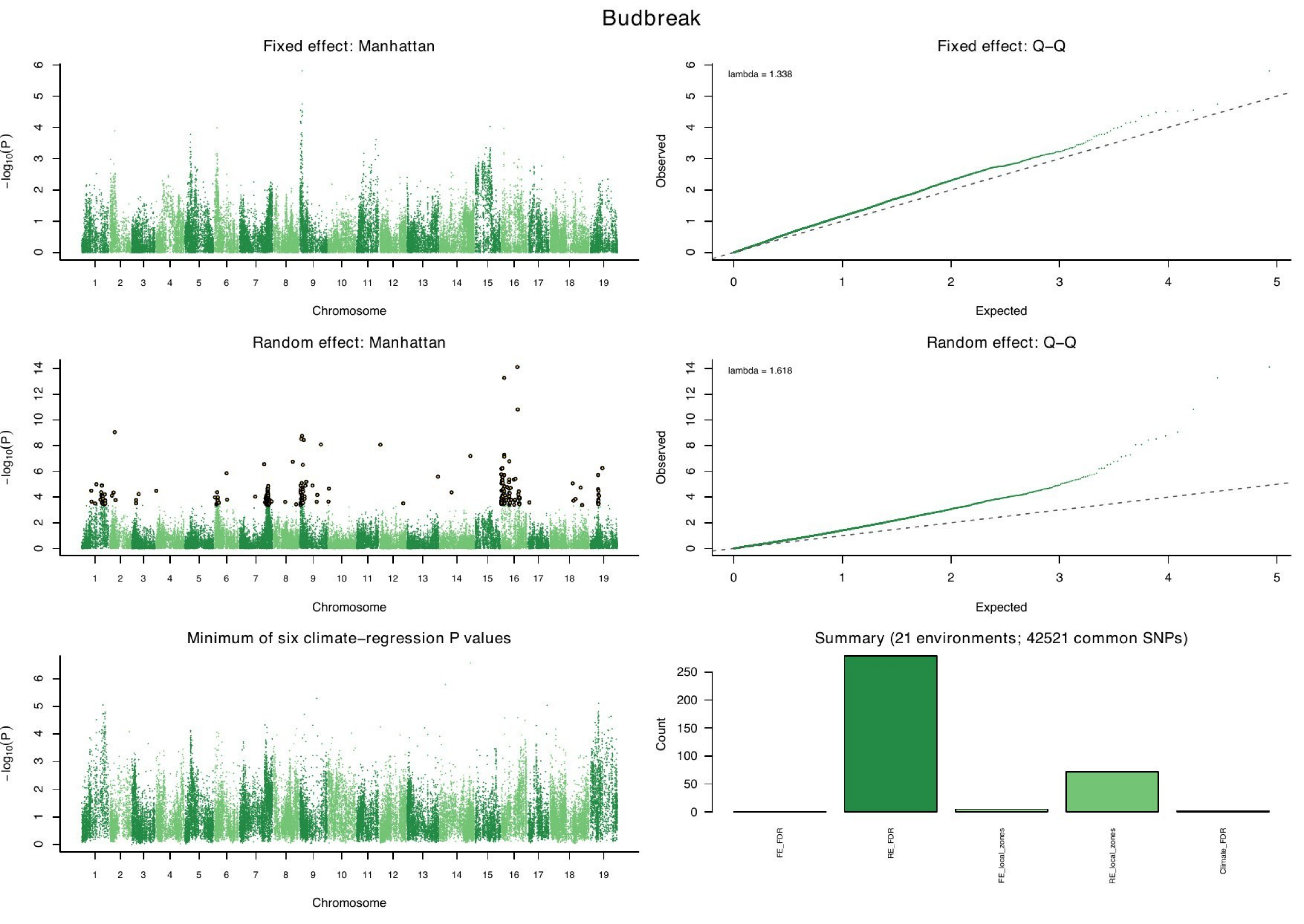

Budbreak
Fixed effect: Manhattan
$-\log_{10}(P)$
Chromosome
1 2 3 4 5 6 7 8 9 10 11 12 13 14 15 16 17 18 19
Fixed effect: Q–Q
lambda = 1.338
Observed
Expected
Random effect: Manhattan
$-\log_{10}(P)$
Chromosome
Random effect: Q–Q
lambda = 1.618
Observed
Expected
Minimum of six climate–regression P values
$-\log_{10}(P)$
Chromosome
Summary (21 environments; 42521 common SNPs)
Count
FE_FDR
RE_FDR
FE_local_zones
RE_local_zones
Climate_FDR

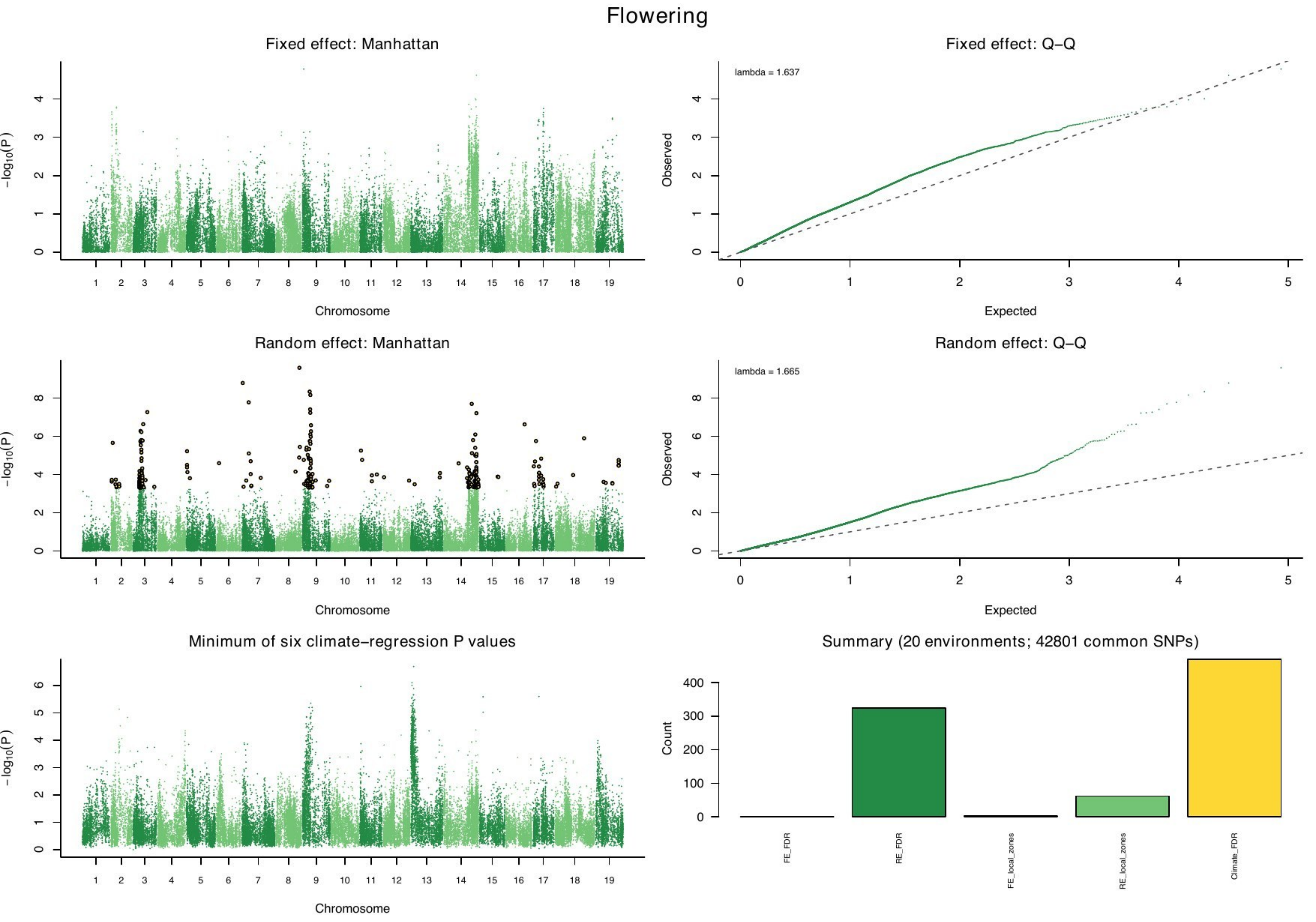

Flowering
Fixed effect: Manhattan
Fixed effect: Q–Q
lambda = 1.637
Random effect: Manhattan
Random effect: Q–Q
lambda = 1.665
Minimum of six climate–regression P values
Summary (20 environments; 42801 common SNPs)
−log10(P)
Chromosome
Observed
Expected
Count
FE_FDR
RE_FDR
FE_local_zones
RE_local_zones
Climate_FDR

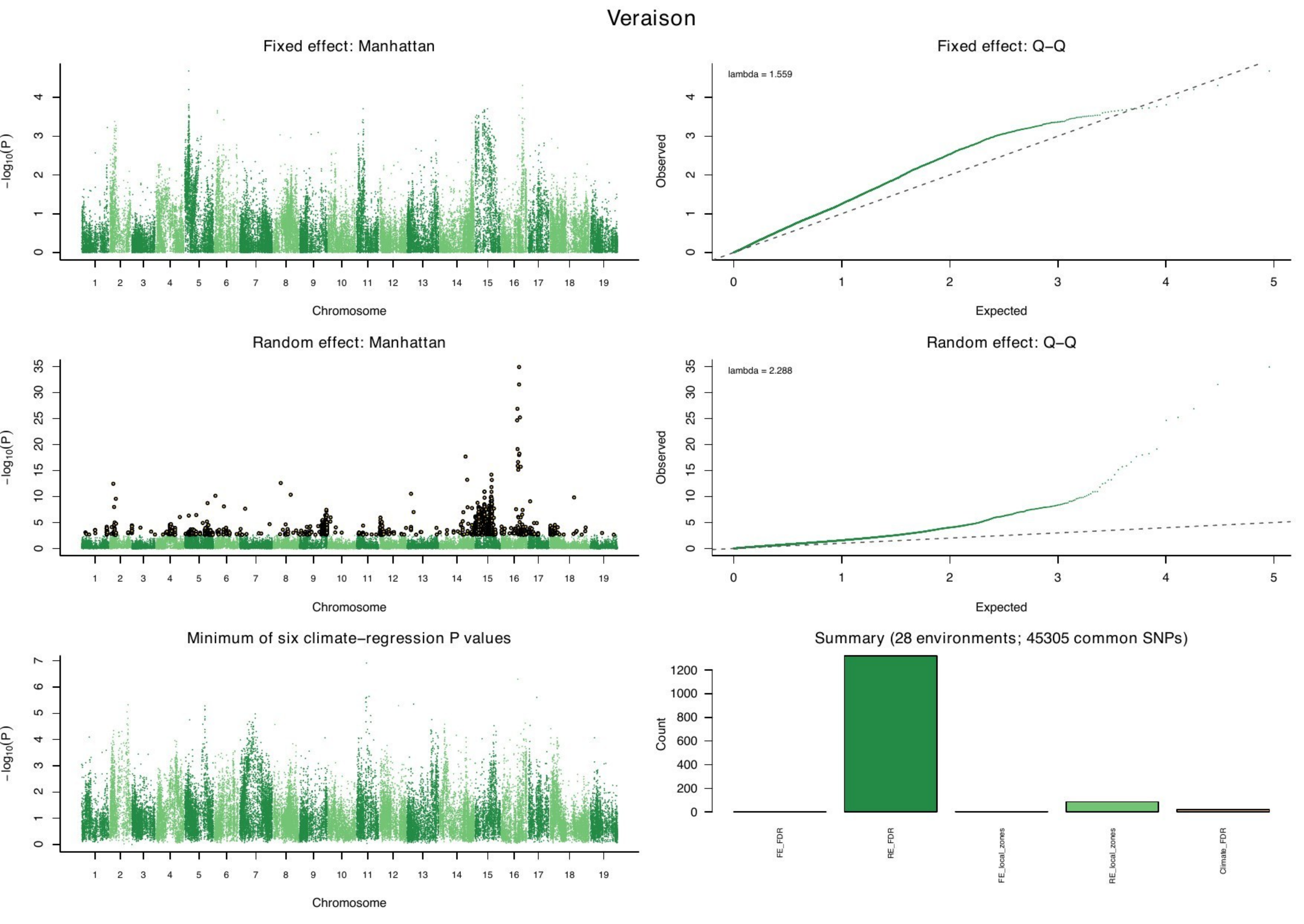
Veraison
Fixed effect: Manhattan
Fixed effect: Q–Q
lambda = 1.559
Random effect: Manhattan
Random effect: Q–Q
lambda = 2.288
Minimum of six climate–regression P values
Summary (28 environments; 45305 common SNPs)
−log10(P)
Chromosome
Observed
Expected
Count
FE_FDR
RE_FDR
FE_local_zones
RE_local_zones
Climate_FDR

### Harvest date

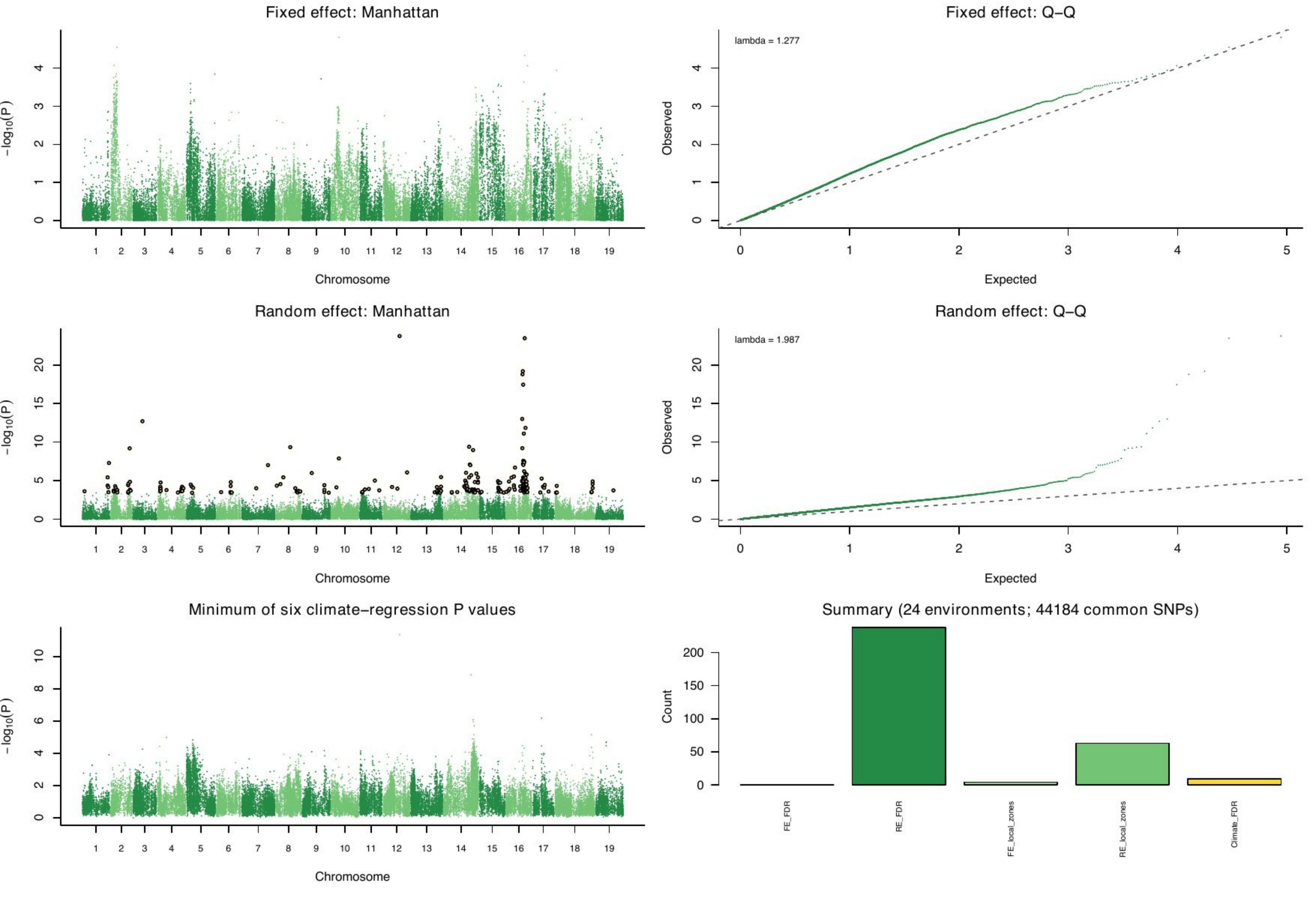

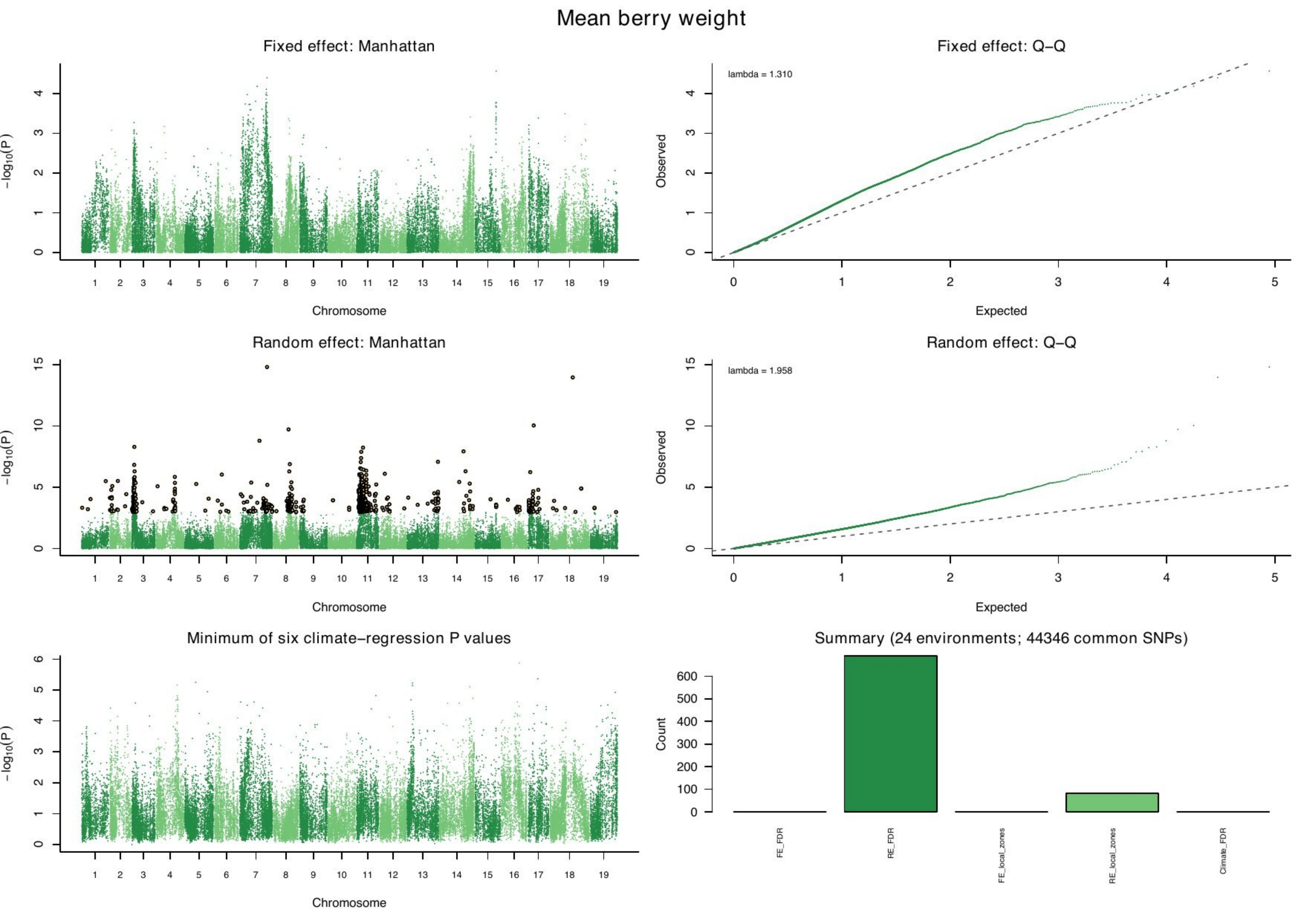
Mean berry weight
Fixed effect: Manhattan
Fixed effect: Q–Q
lambda = 1.310
Random effect: Manhattan
Random effect: Q–Q
lambda = 1.958
Minimum of six climate–regression P values
Summary (24 environments; 44346 common SNPs)
−log10(P)
Chromosome
Observed
Expected
Count
FE_FDR
RE_FDR
FE_local_zones
RE_local_zones
Climate_FDR

Berry sugar content

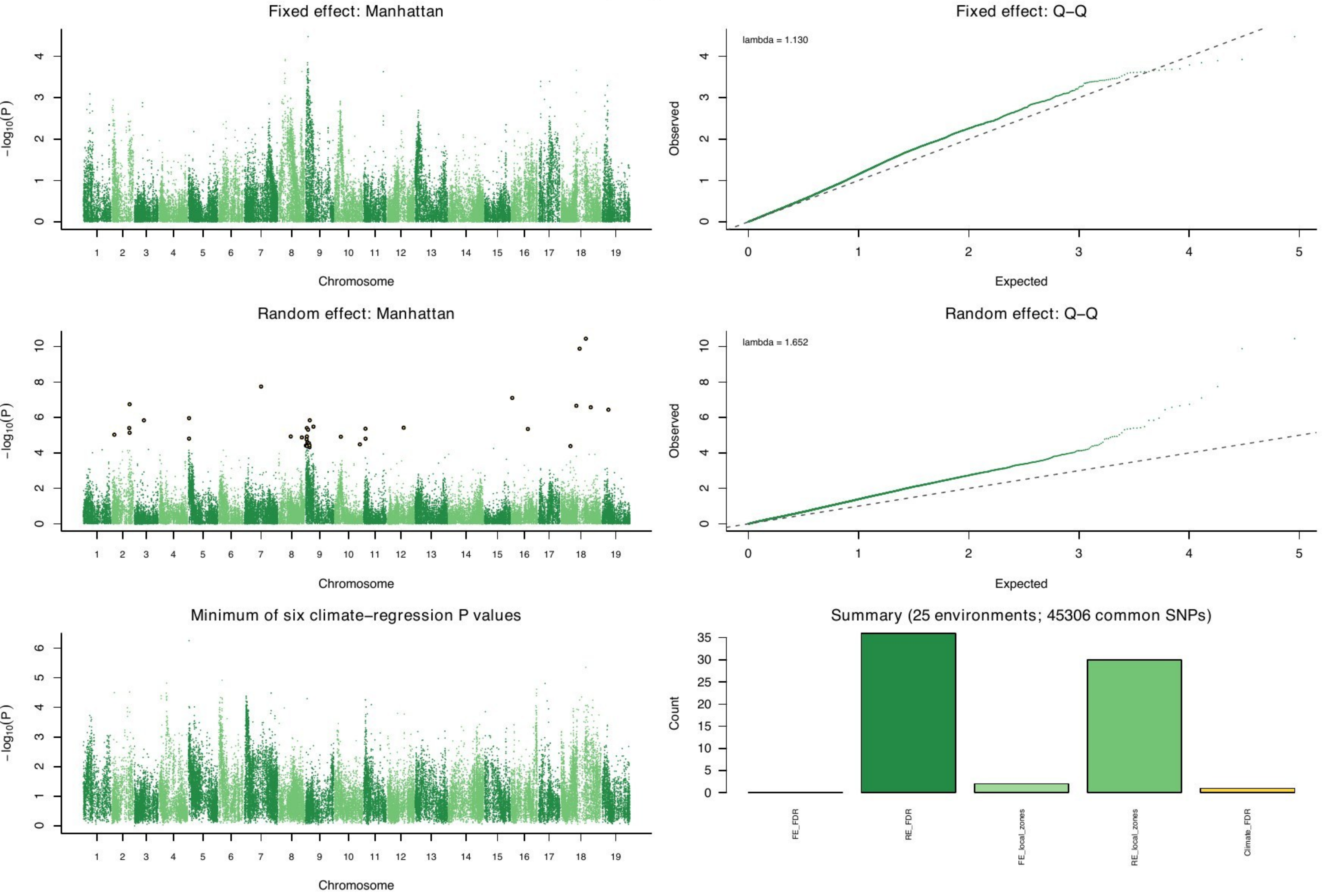

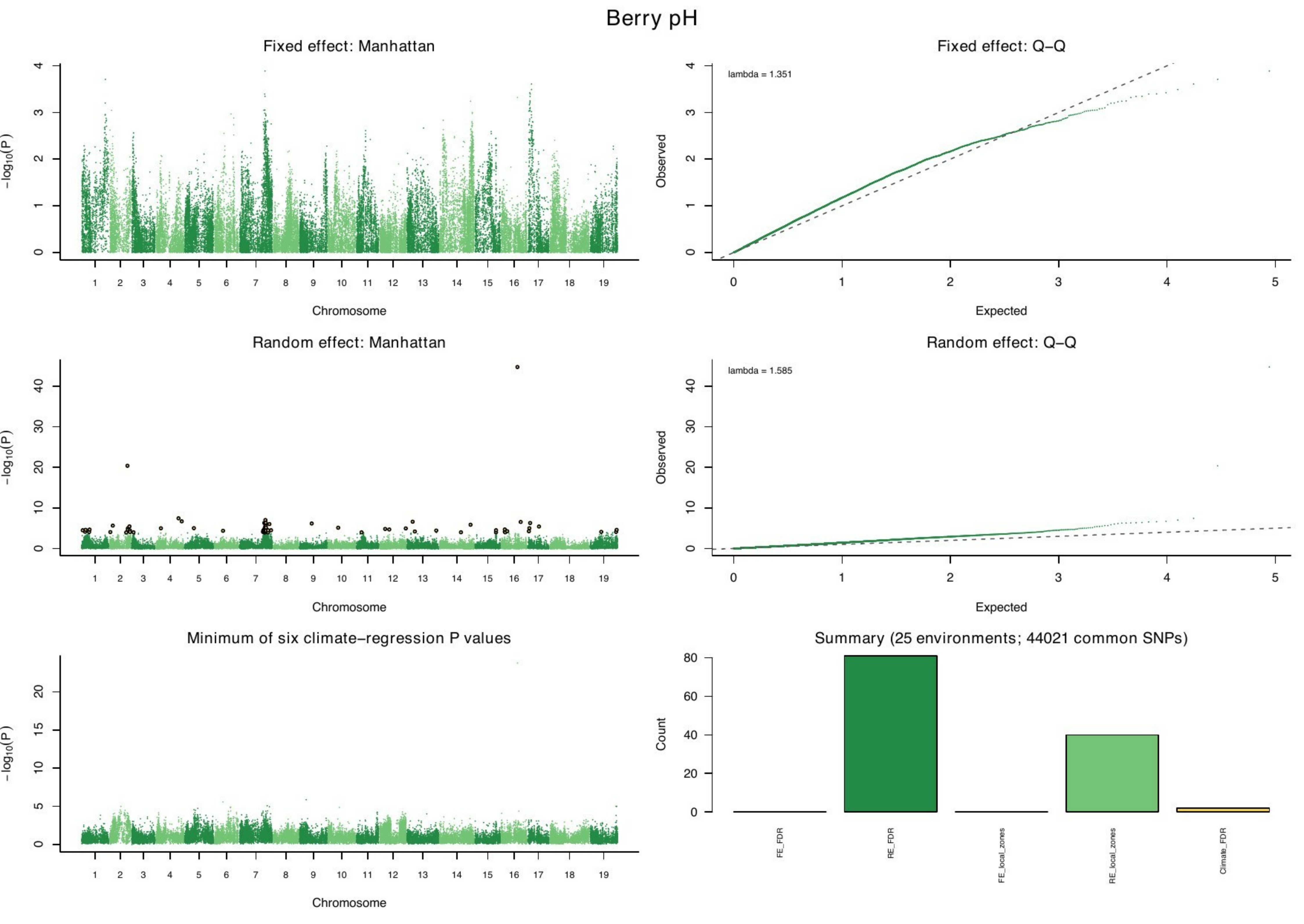
Berry pH
Fixed effect: Manhattan
Fixed effect: Q−Q
lambda = 1.351
Random effect: Manhattan
Random effect: Q−Q
lambda = 1.585
Minimum of six climate−regression P values
Summary (25 environments; 44021 common SNPs)
−log10(P)
Chromosome
Observed
Expected
Count
FE_FDR
RE_FDR
FE_local_zones
RE_local_zones
Climate_FDR

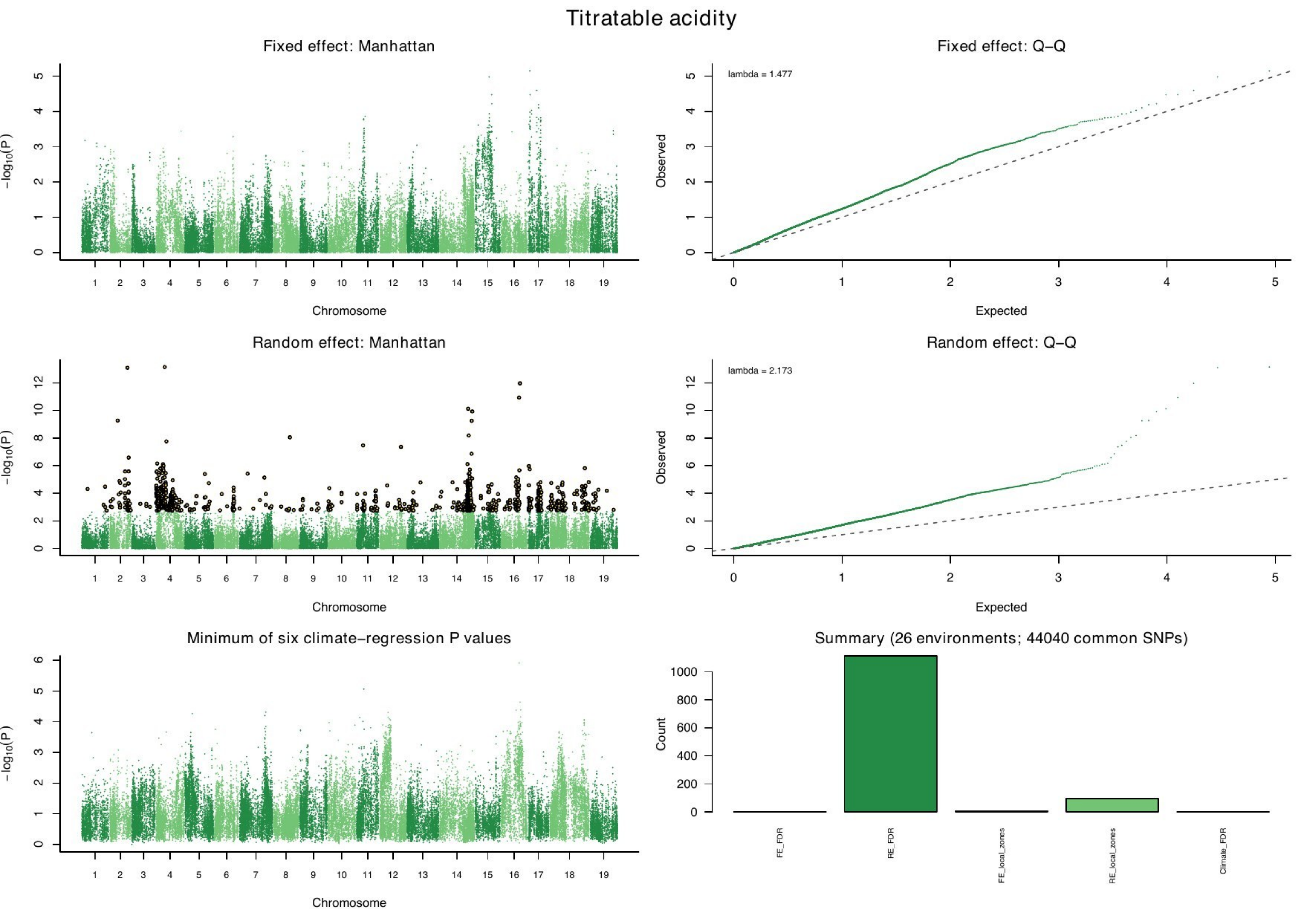

Titratable acidity
Fixed effect: Manhattan
Fixed effect: Q–Q
lambda = 1.477
Random effect: Manhattan
Random effect: Q–Q
lambda = 2.173
Minimum of six climate–regression P values
Summary (26 environments; 44040 common SNPs)
−log10(P)
Chromosome
Observed
Expected
Count
FE_FDR
RE_FDR
FE_local_zones
RE_local_zones
Climate_FDR

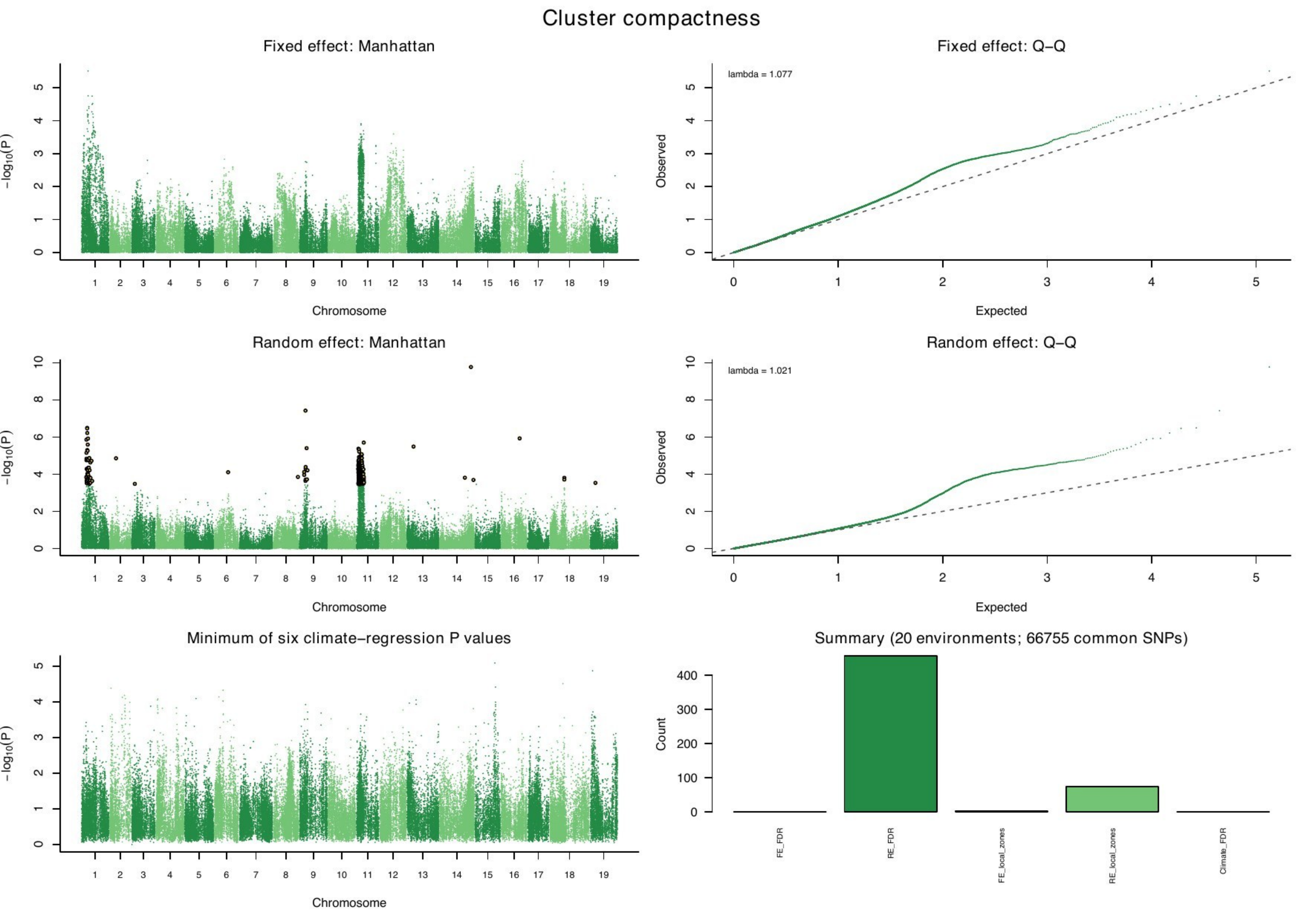

Cluster compactness
Fixed effect: Manhattan
Fixed effect: Q–Q
lambda = 1.077
Random effect: Manhattan
Random effect: Q–Q
lambda = 1.021
Minimum of six climate–regression P values
Summary (20 environments; 66755 common SNPs)
−log10(P)
Chromosome
Observed
Expected
Count
FE_FDR
RE_FDR
FE_local_zones
RE_local_zones
Climate_FDR

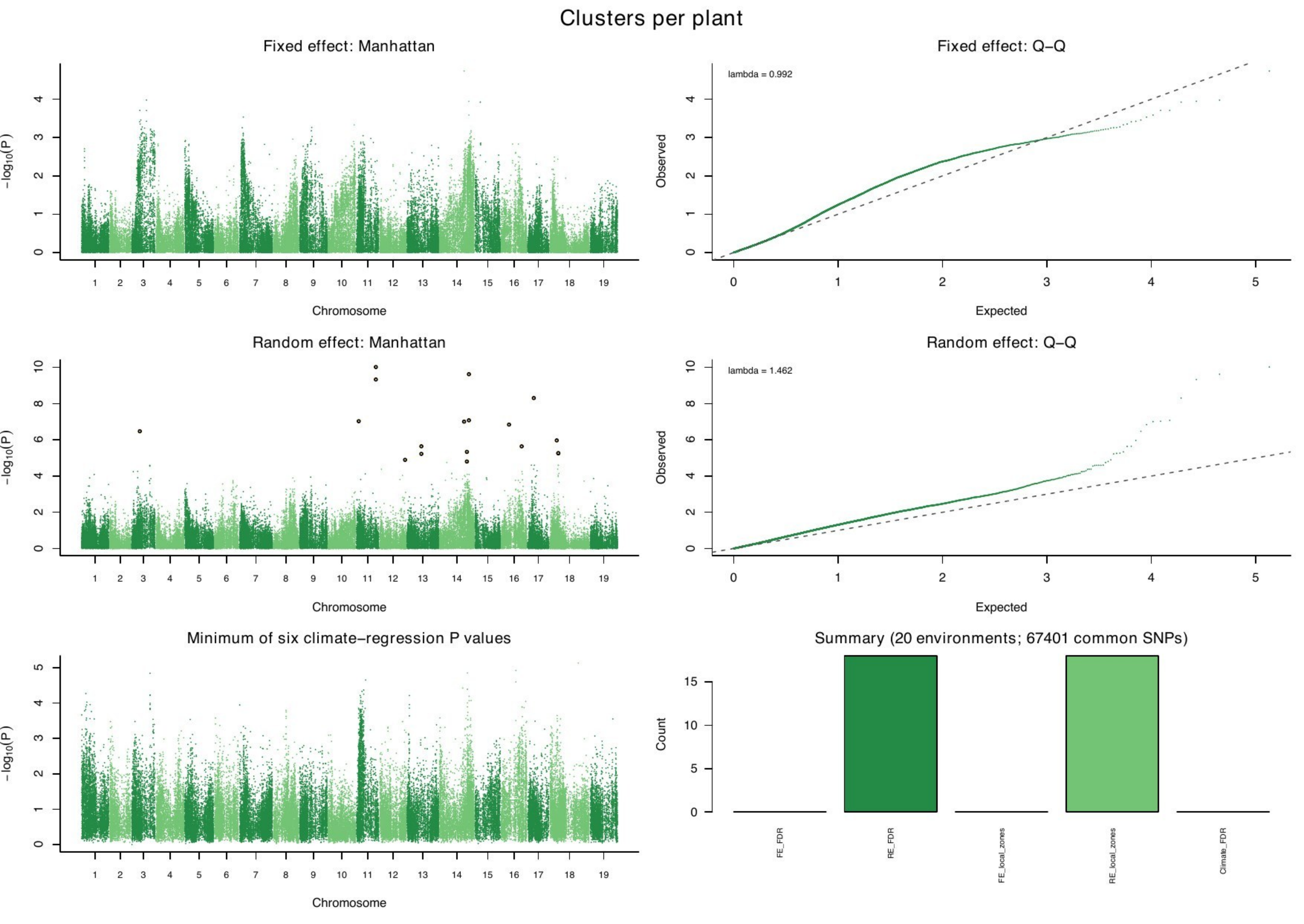

Clusters per plant
Fixed effect: Manhattan
Fixed effect: Q–Q
lambda = 0.992
Random effect: Manhattan
Random effect: Q–Q
lambda = 1.462
Minimum of six climate–regression P values
Summary (20 environments; 67401 common SNPs)
−log10(P)
Chromosome
Observed
Expected
Count
FE_FDR
RE_FDR
FE_local_zones
RE_local_zones
Climate_FDR

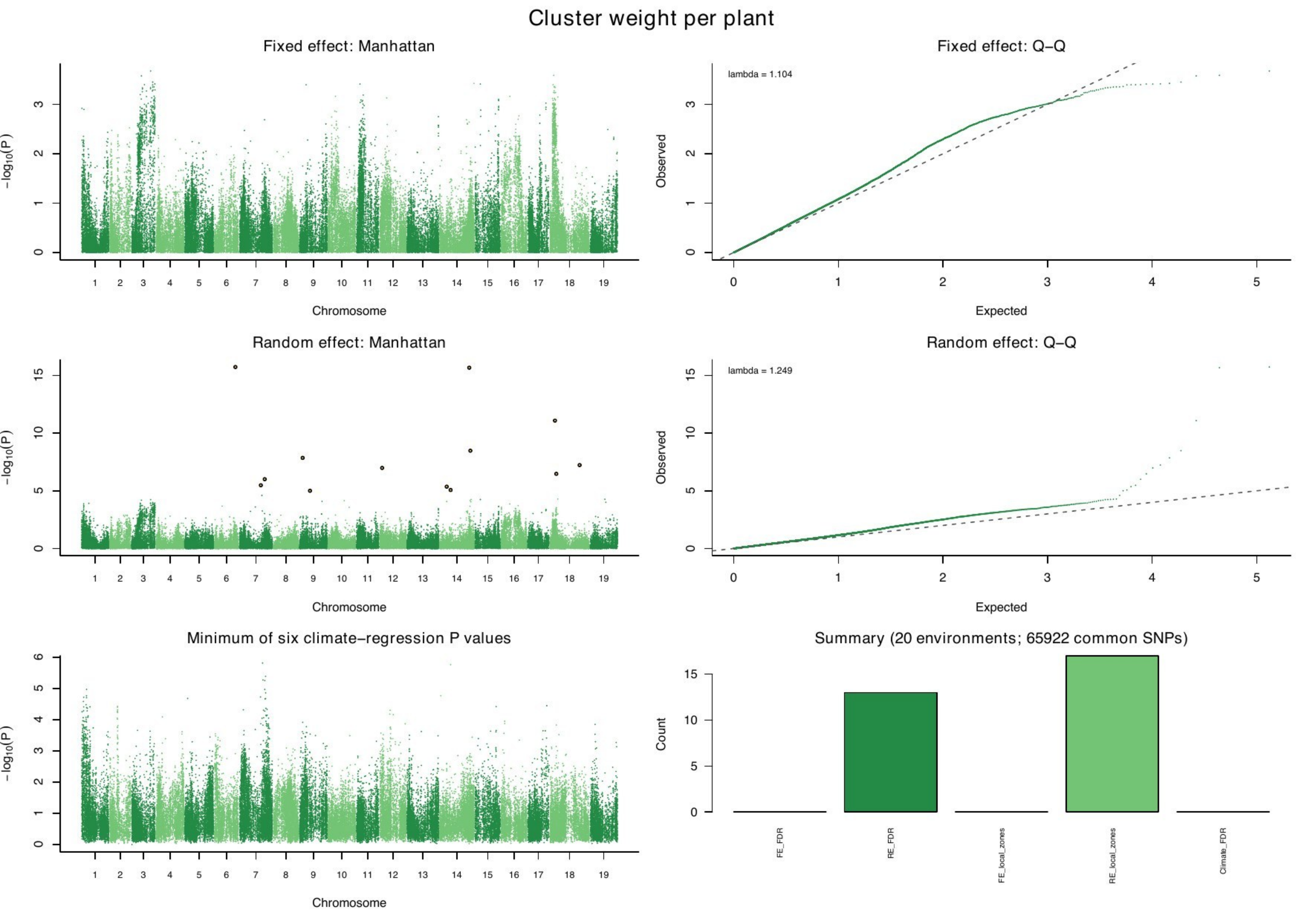
Cluster weight per plant
Fixed effect: Manhattan
Fixed effect: Q–Q
lambda = 1.104
Random effect: Manhattan
Random effect: Q–Q
lambda = 1.249
Minimum of six climate–regression P values
Summary (20 environments; 65922 common SNPs)
−log10(P)
Chromosome
Observed
Expected
Count
FE_FDR
RE_FDR
FE_local_zones
RE_local_zones
Climate_FDR

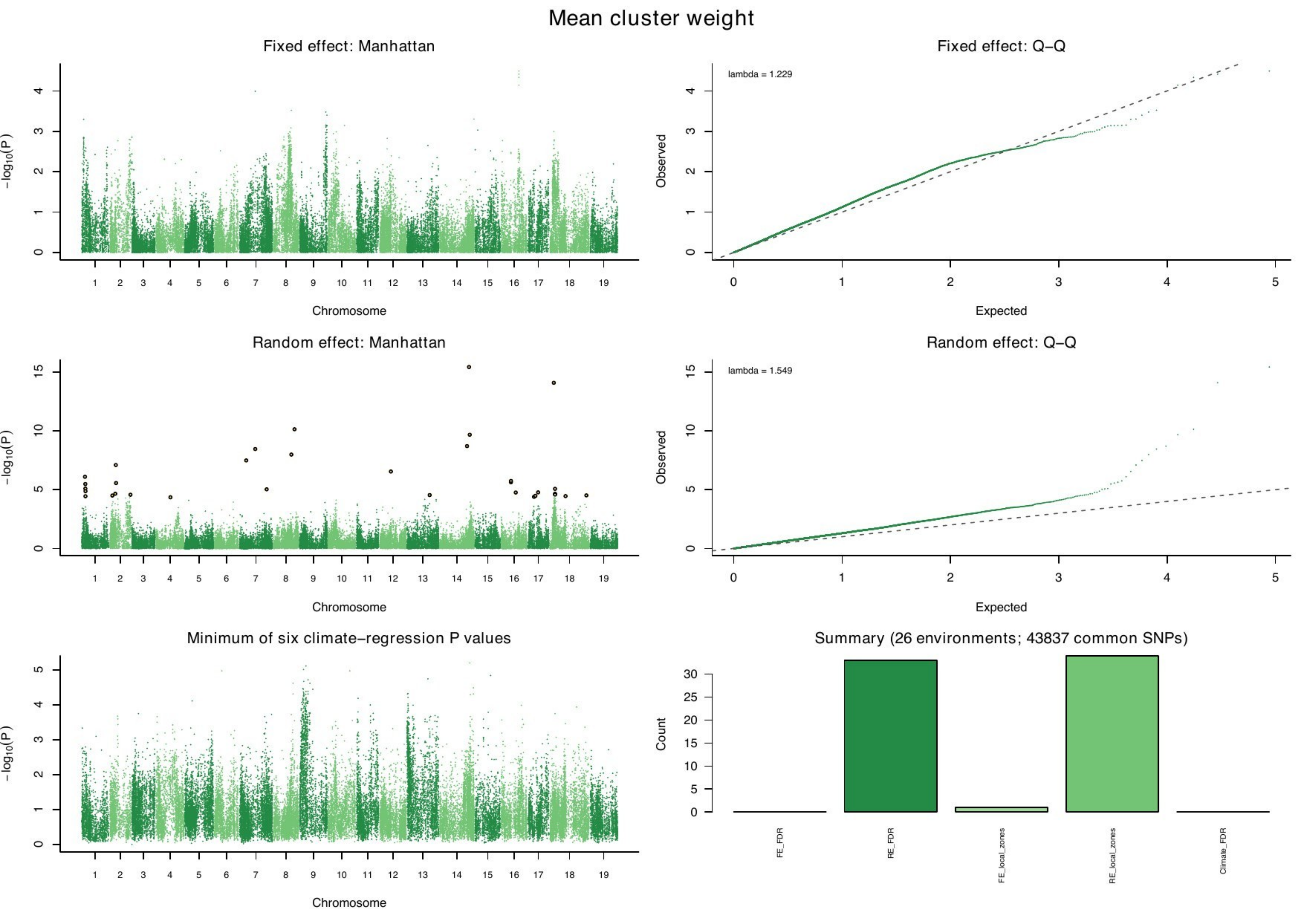

Mean cluster weight
Fixed effect: Manhattan
Fixed effect: Q–Q
lambda = 1.229
Random effect: Manhattan
Random effect: Q–Q
lambda = 1.549
Minimum of six climate–regression P values
Summary (26 environments; 43837 common SNPs)
$-\log_{10}(P)$
Chromosome
Observed
Expected
Count
FE_FDR
RE_FDR
FE_local_zones
RE_local_zones
Climate_FDR

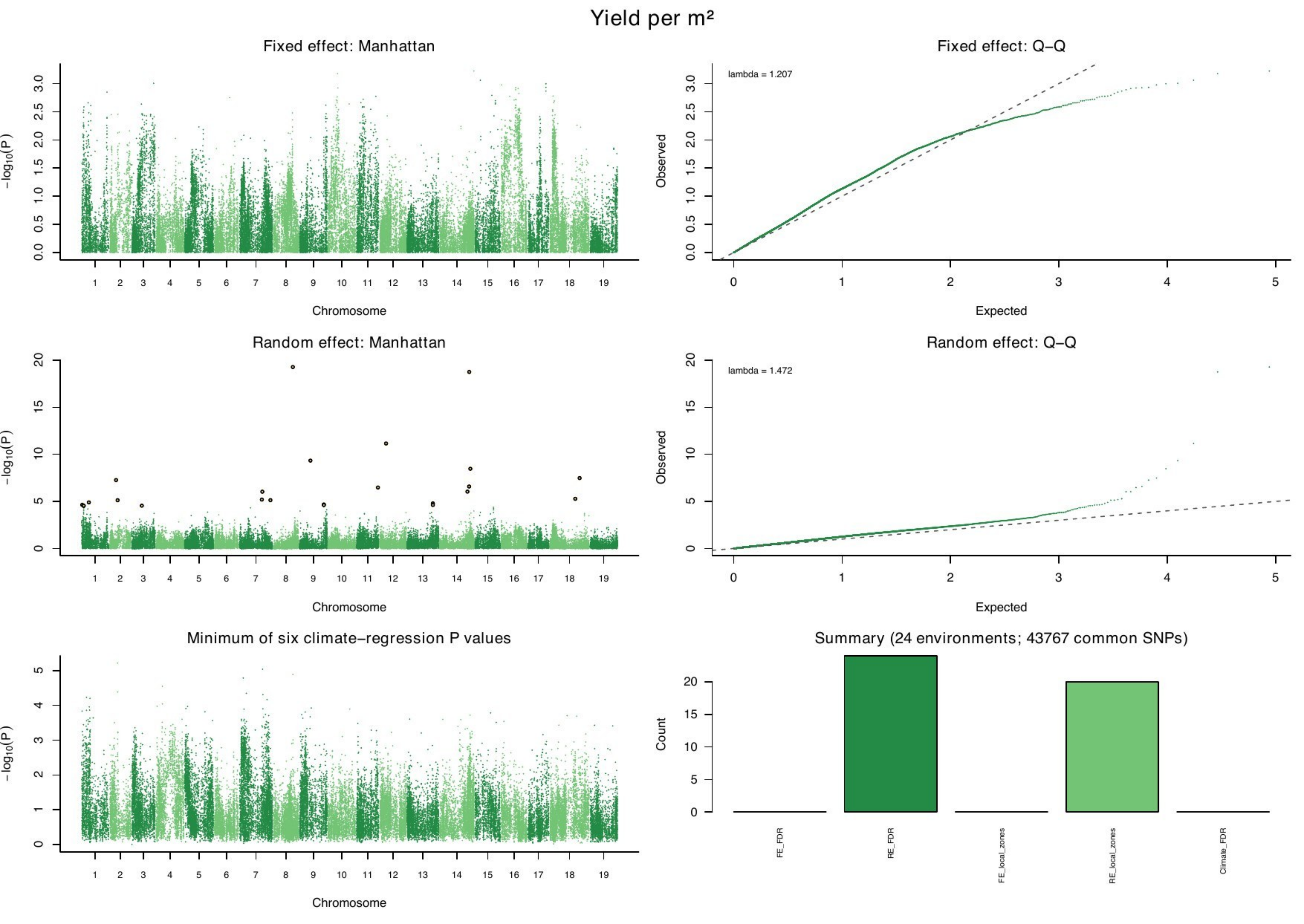
Yield per m²
Fixed effect: Manhattan
Fixed effect: Q−Q
lambda = 1.207
Random effect: Manhattan
Random effect: Q−Q
lambda = 1.472
Minimum of six climate−regression P values
Summary (24 environments; 43767 common SNPs)
−log10(P)
Chromosome
Observed
Expected
Count
FE_FDR
RE_FDR
FE_local_zones
RE_local_zones
Climate_FDR
