## Supplementary File 4 for "Uncovering the genetic basis of agronomic traits in over 1,000 grapevine genotypes derived from a disease resistance breeding program"

### Budbreak

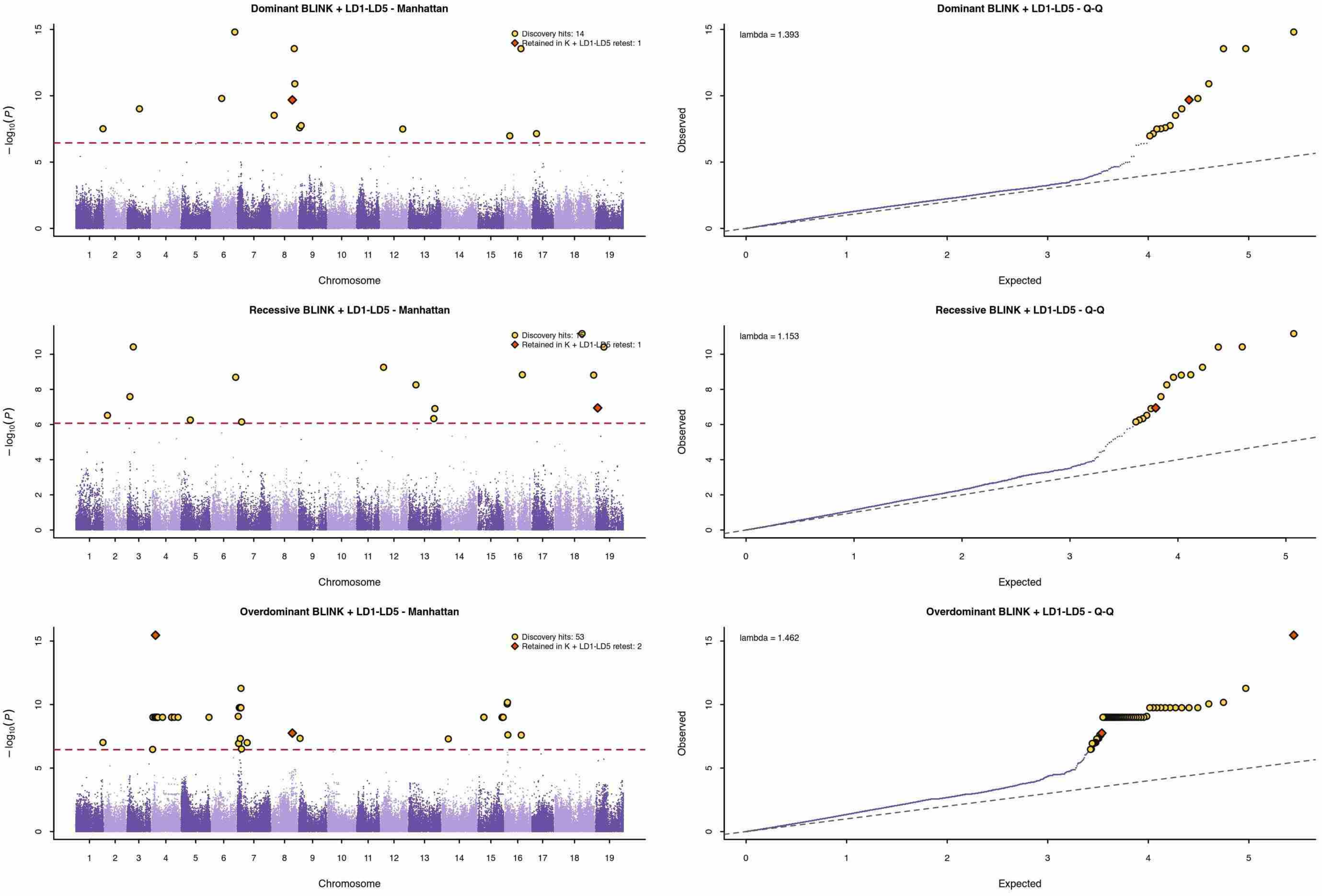


Dashed red line: 5% trait-specific Bonferroni threshold. Yellow circles: discovery hits. Orange diamonds: loci retained in the K + LD1-LD5 selected-candidate retest.

### Flowering

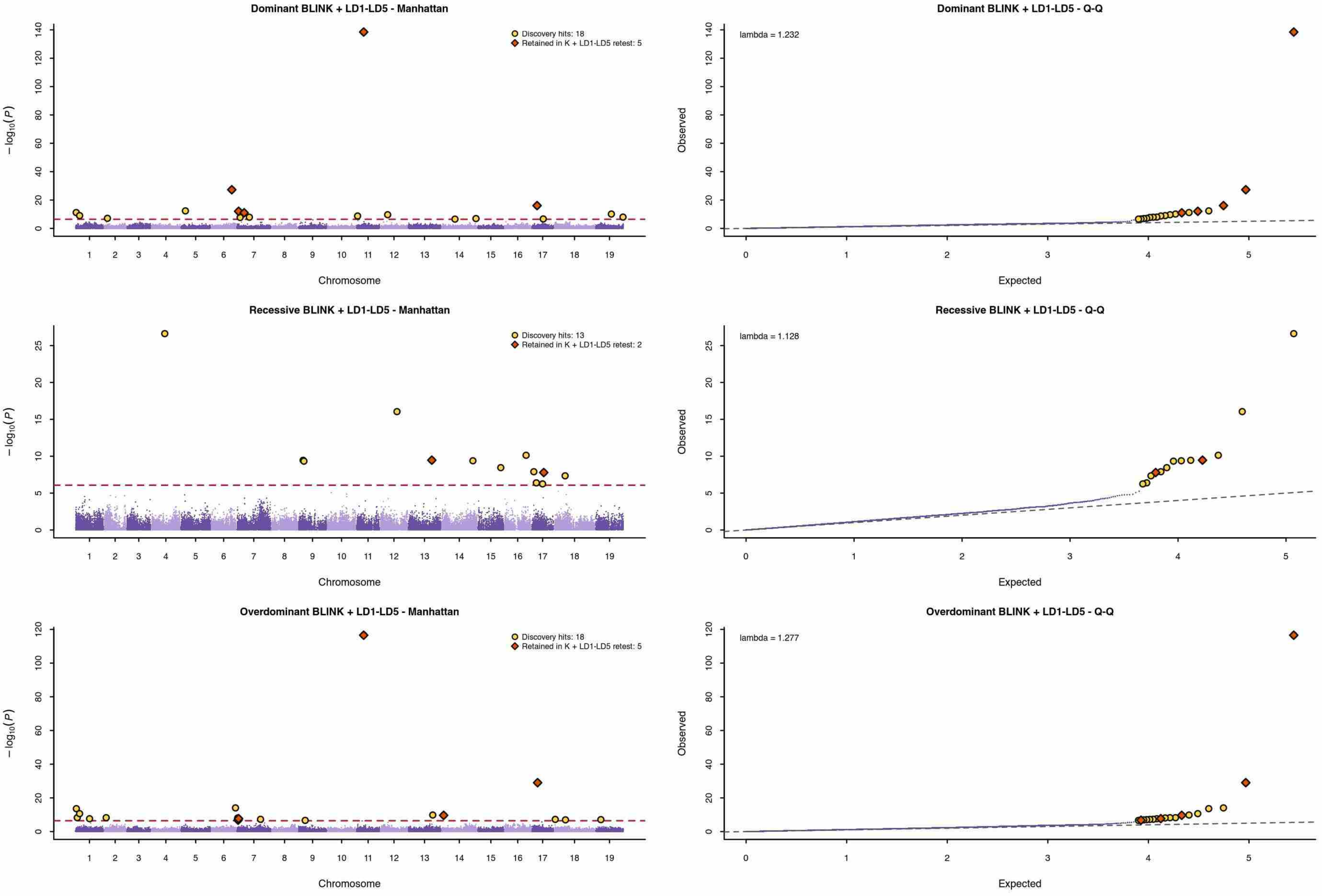


Dashed red line: 5% trait-specific Bonferroni threshold. Yellow circles: discovery hits. Orange diamonds: loci retained in the K + LD1-LD5 selected-candidate retest.

### Veraison

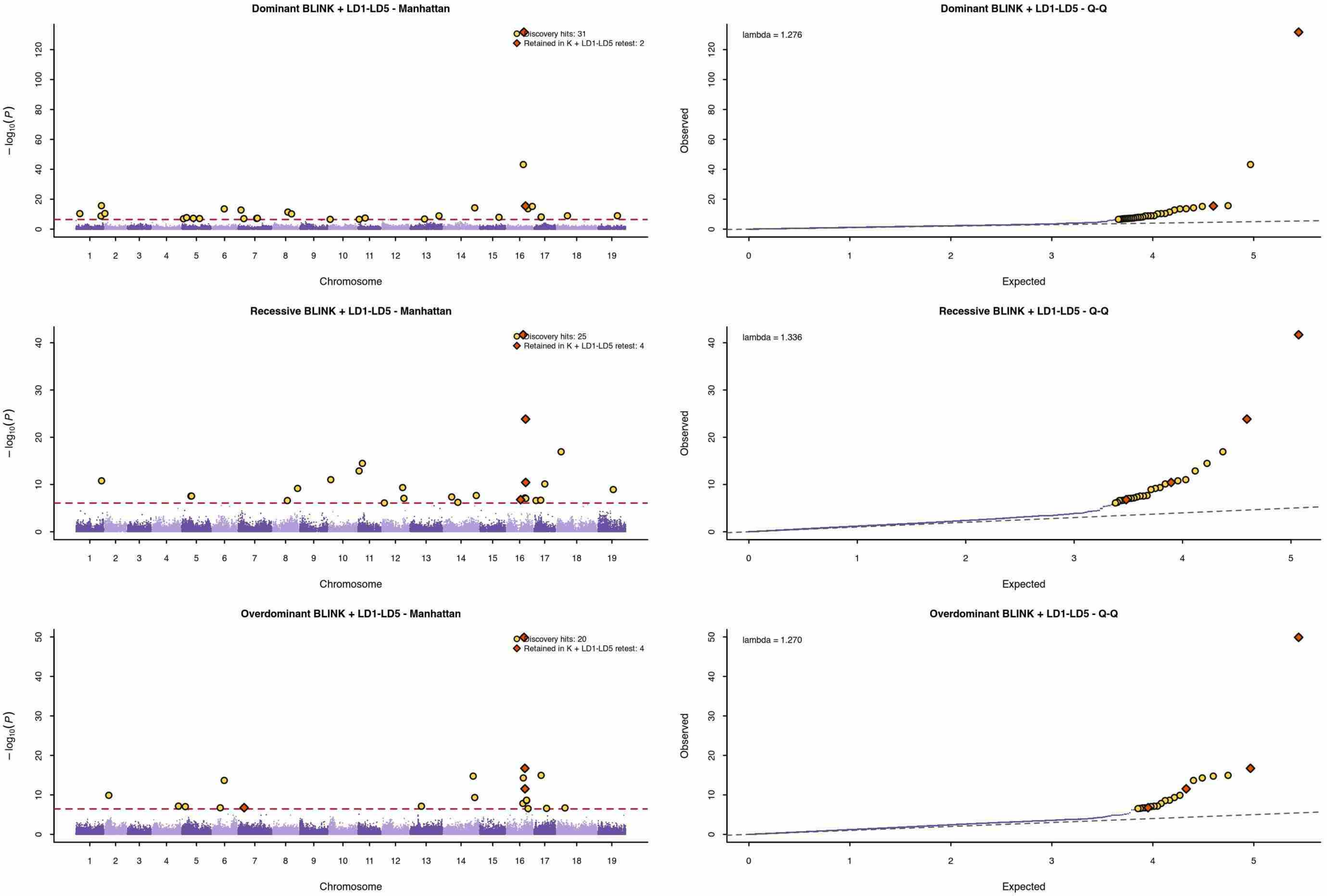


Dashed red line: 5% trait-specific Bonferroni threshold. Yellow circles: discovery hits. Orange diamonds: loci retained in the K + LD1-LD5 selected-candidate retest.

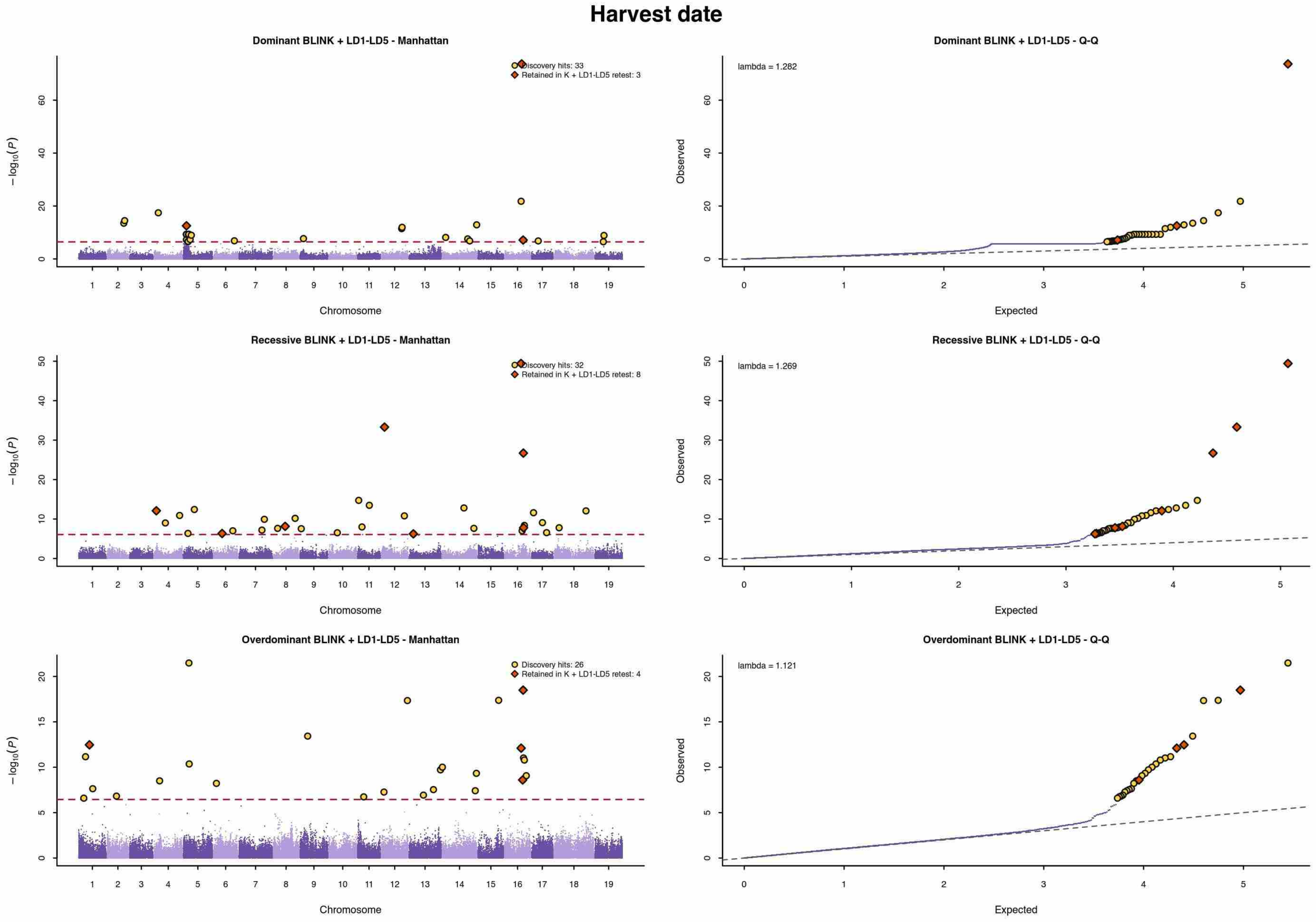


Dashed red line: 5% trait-specific Bonferroni threshold. Yellow circles: discovery hits. Orange diamonds: loci retained in the K + LD1-LD5 selected-candidate retest.

### Mean berry weight

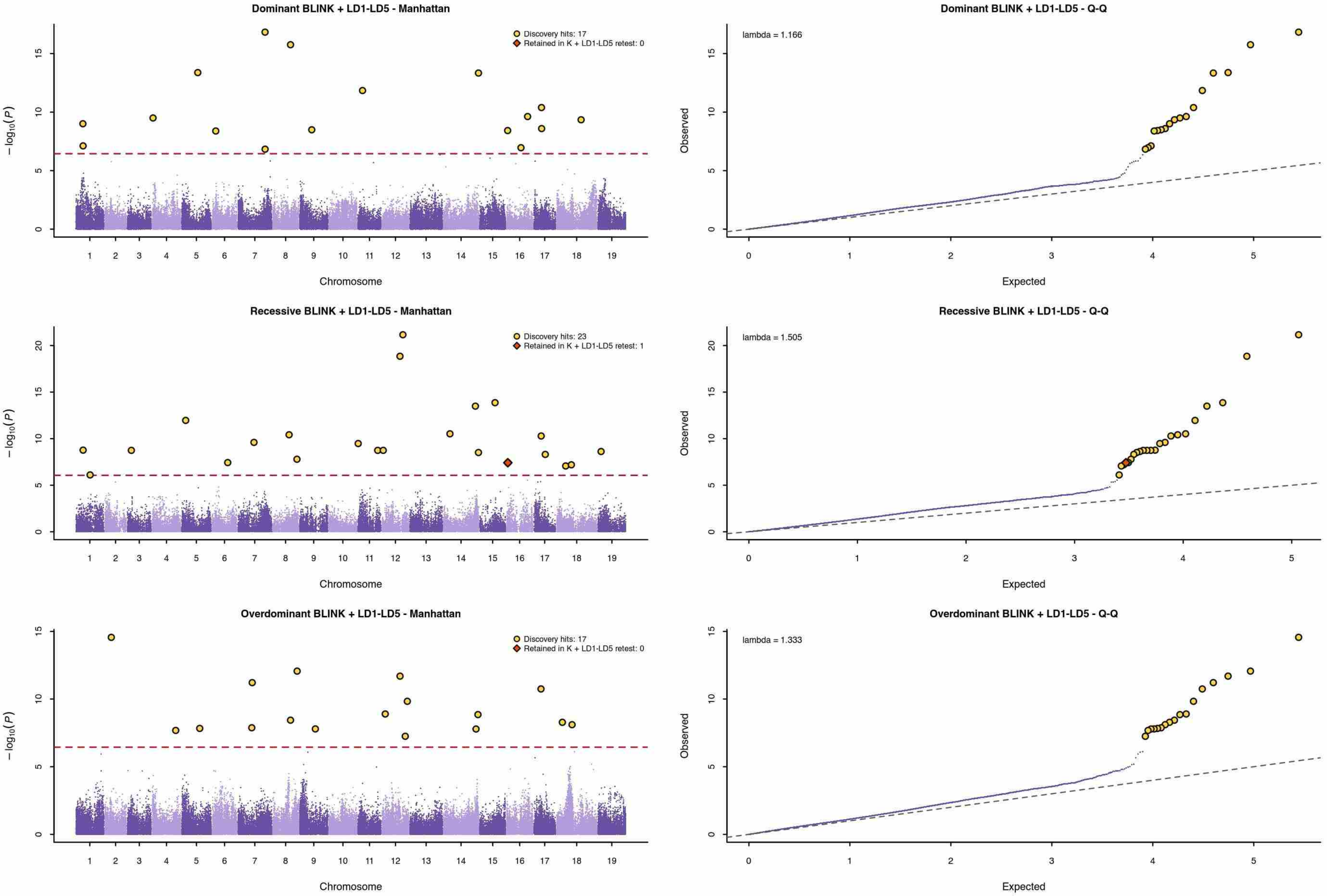


Dashed red line: 5% trait-specific Bonferroni threshold. Yellow circles: discovery hits. Orange diamonds: loci retained in the K + LD1-LD5 selected-candidate retest.

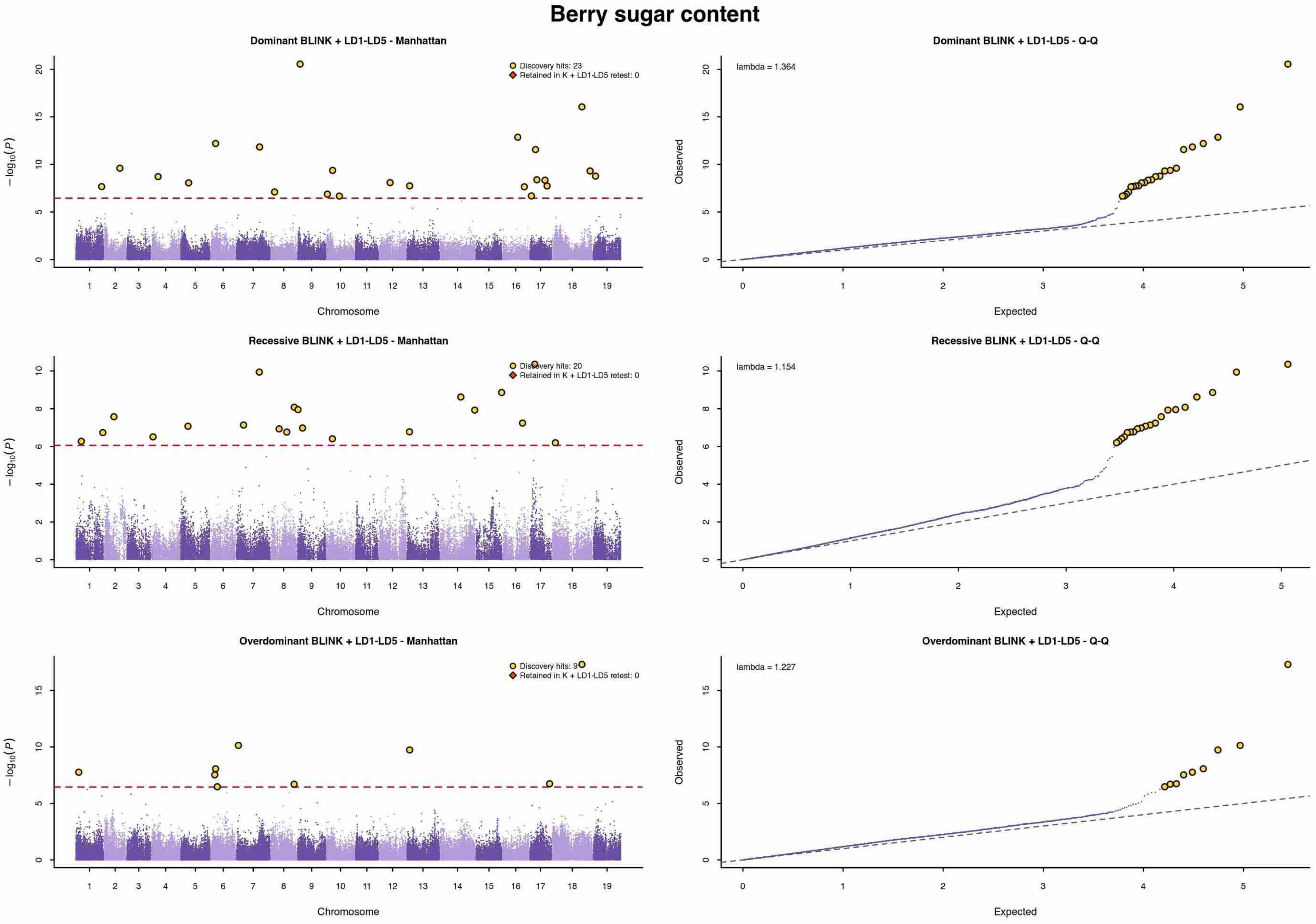


Dashed red line: 5% trait-specific Bonferroni threshold. Yellow circles: discovery hits. Orange diamonds: loci retained in the K + LD1-LD5 selected-candidate retest.

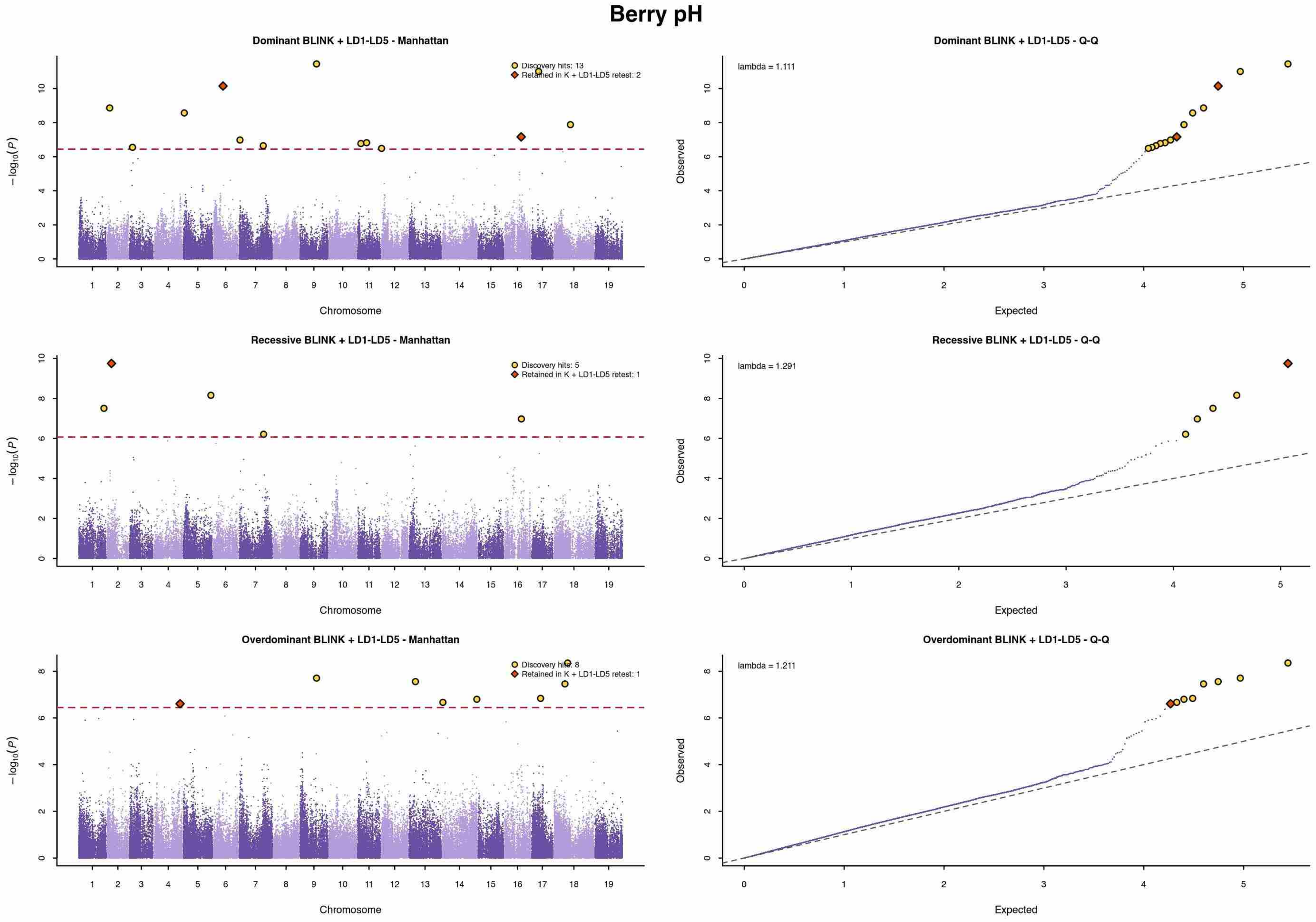


Dashed red line: 5% trait-specific Bonferroni threshold. Yellow circles: discovery hits. Orange diamonds: loci retained in the K + LD1-LD5 selected-candidate retest.

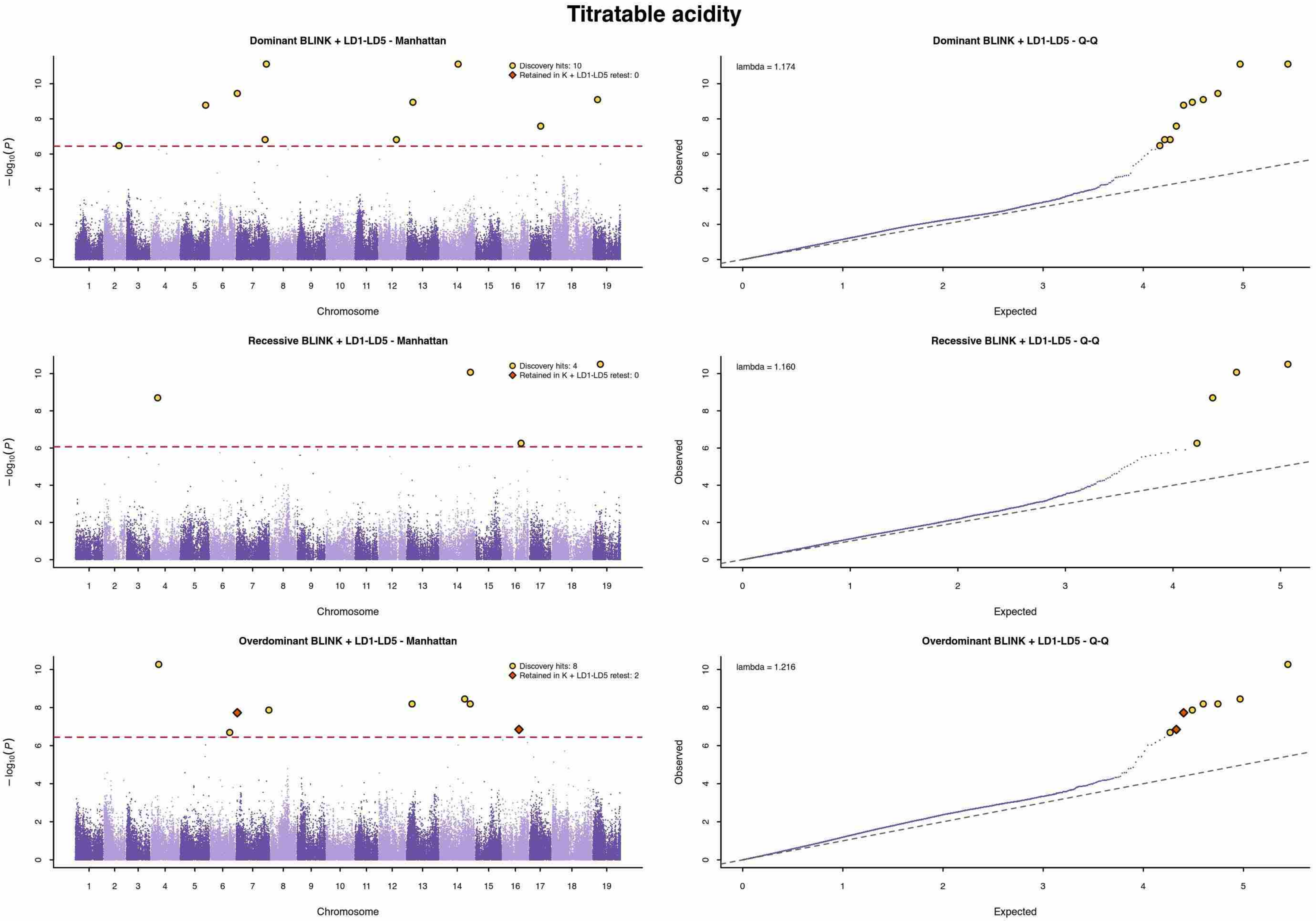

Dashed red line: 5% trait-specific Bonferroni threshold. Yellow circles: discovery hits. Orange diamonds: loci retained in the K + LD1-LD5 selected-candidate retest.

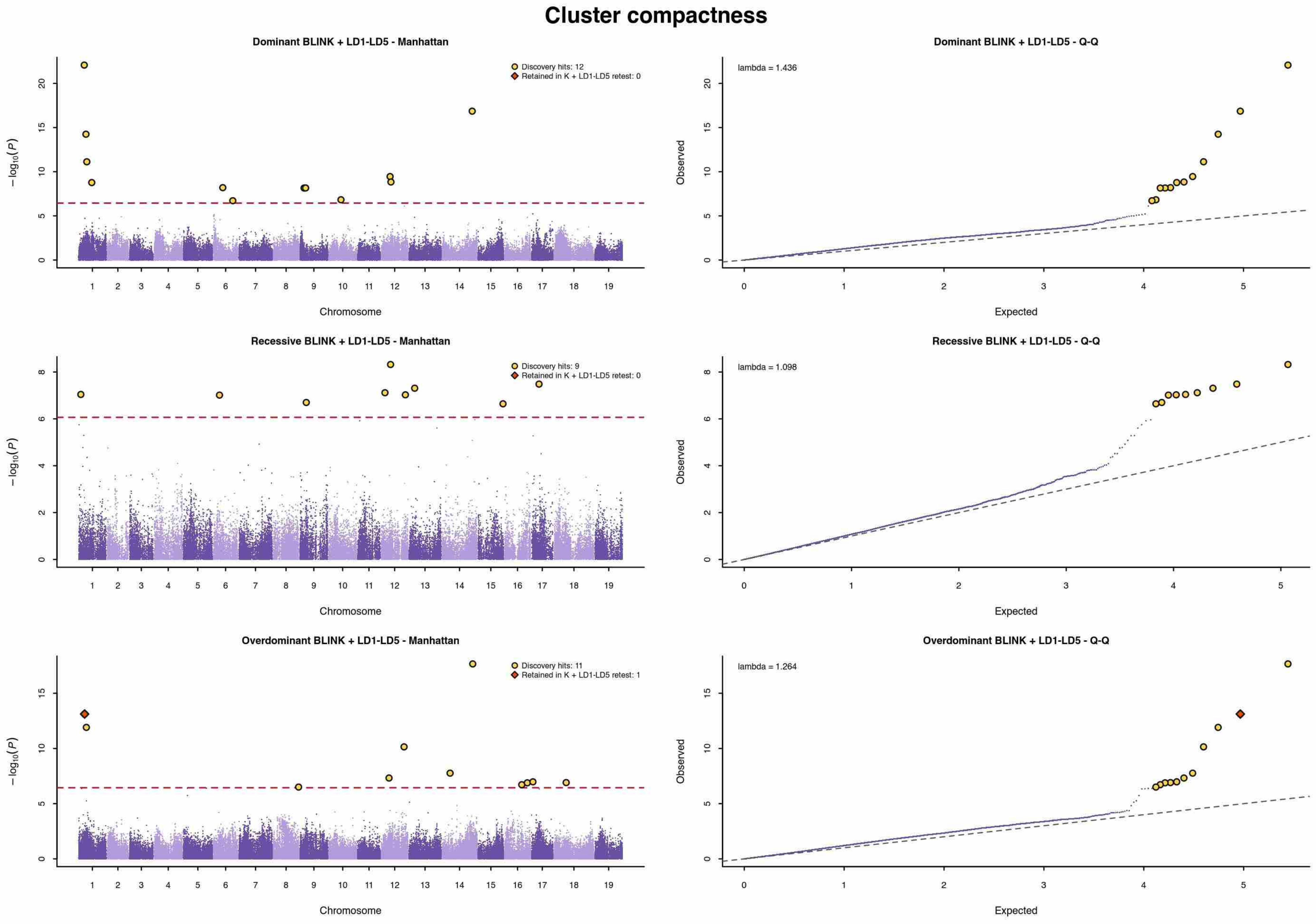


Dashed red line: 5% trait-specific Bonferroni threshold. Yellow circles: discovery hits. Orange diamonds: loci retained in the K + LD1-LD5 selected-candidate retest.

### Clusters per plant

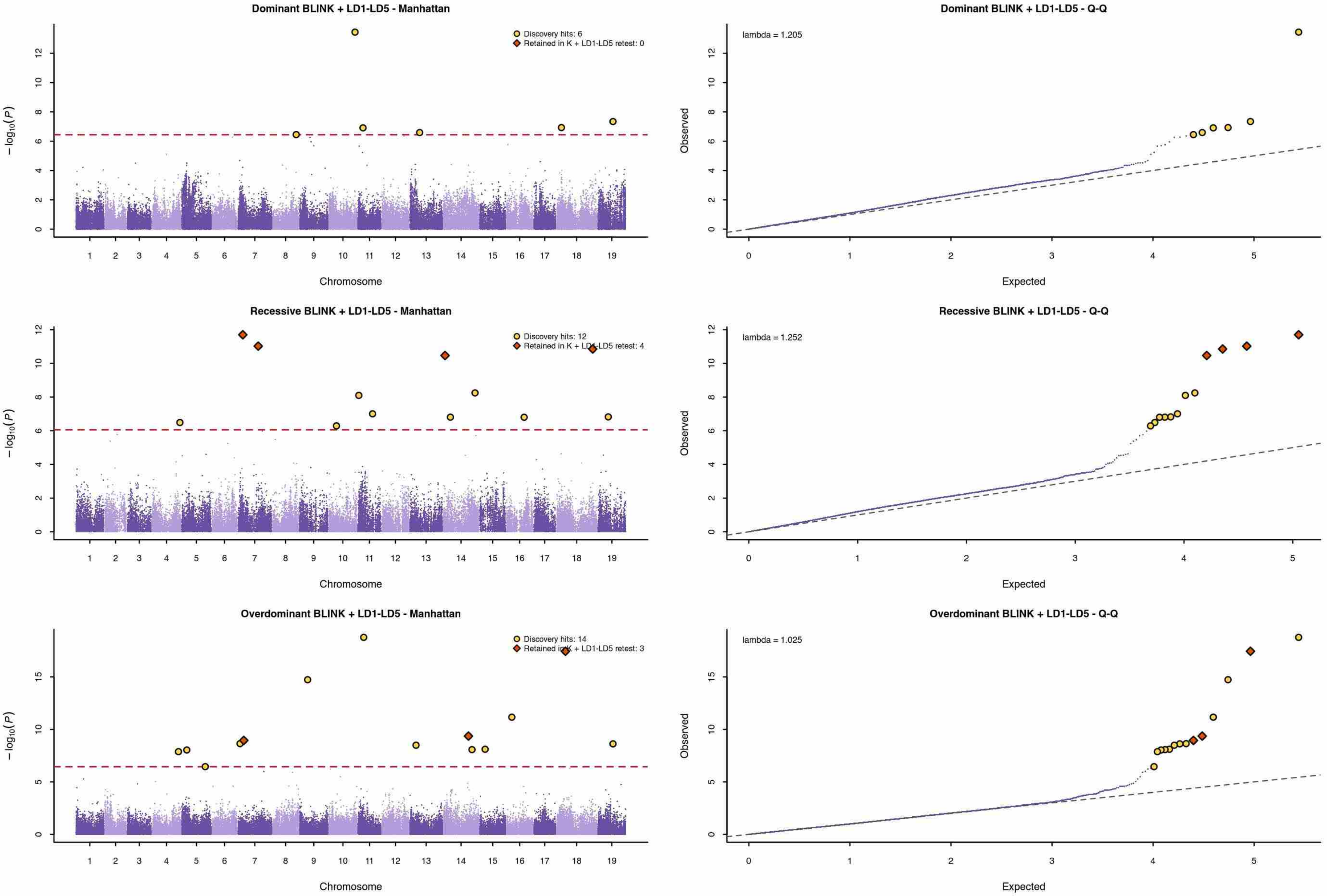


Dashed red line: 5% trait-specific Bonferroni threshold. Yellow circles: discovery hits. Orange diamonds: loci retained in the K + LD1-LD5 selected-candidate retest.

### Cluster weight per plant

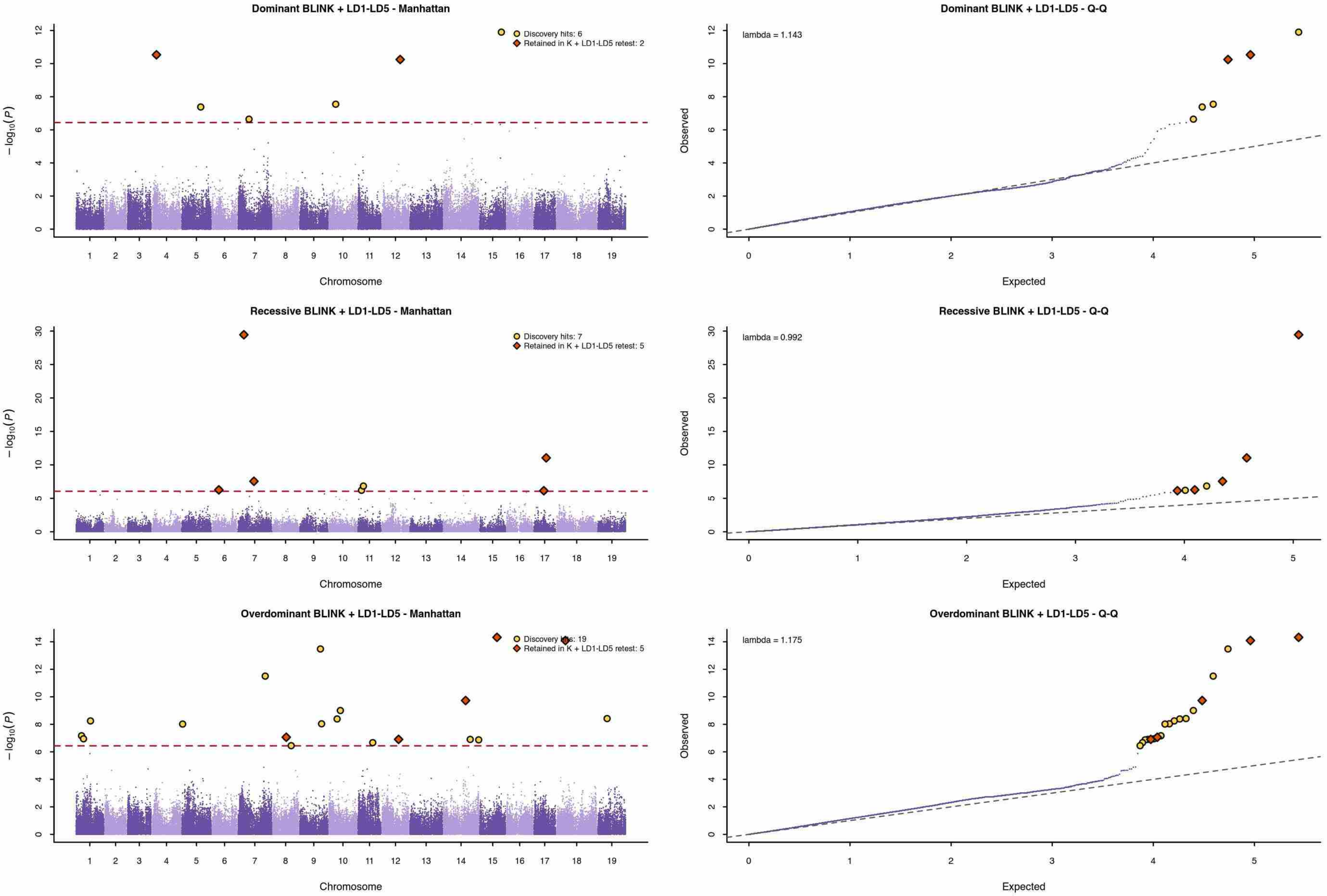


Dashed red line: 5% trait-specific Bonferroni threshold. Yellow circles: discovery hits. Orange diamonds: loci retained in the K + LD1-LD5 selected-candidate retest.

### Mean cluster weight

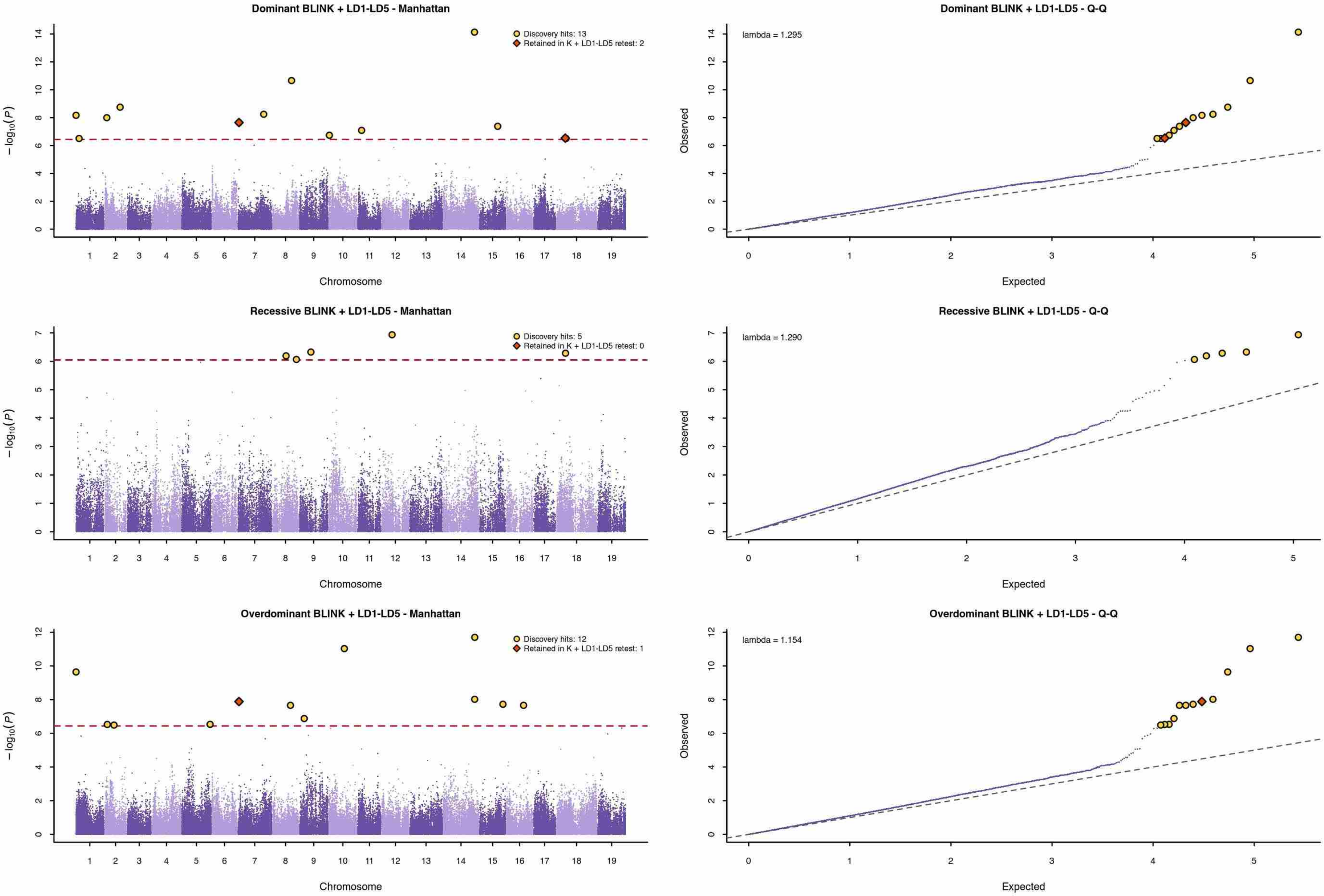


Dashed red line: 5% trait-specific Bonferroni threshold. Yellow circles: discovery hits. Orange diamonds: loci retained in the K + LD1-LD5 selected-candidate retest.

Yield per $m^2$

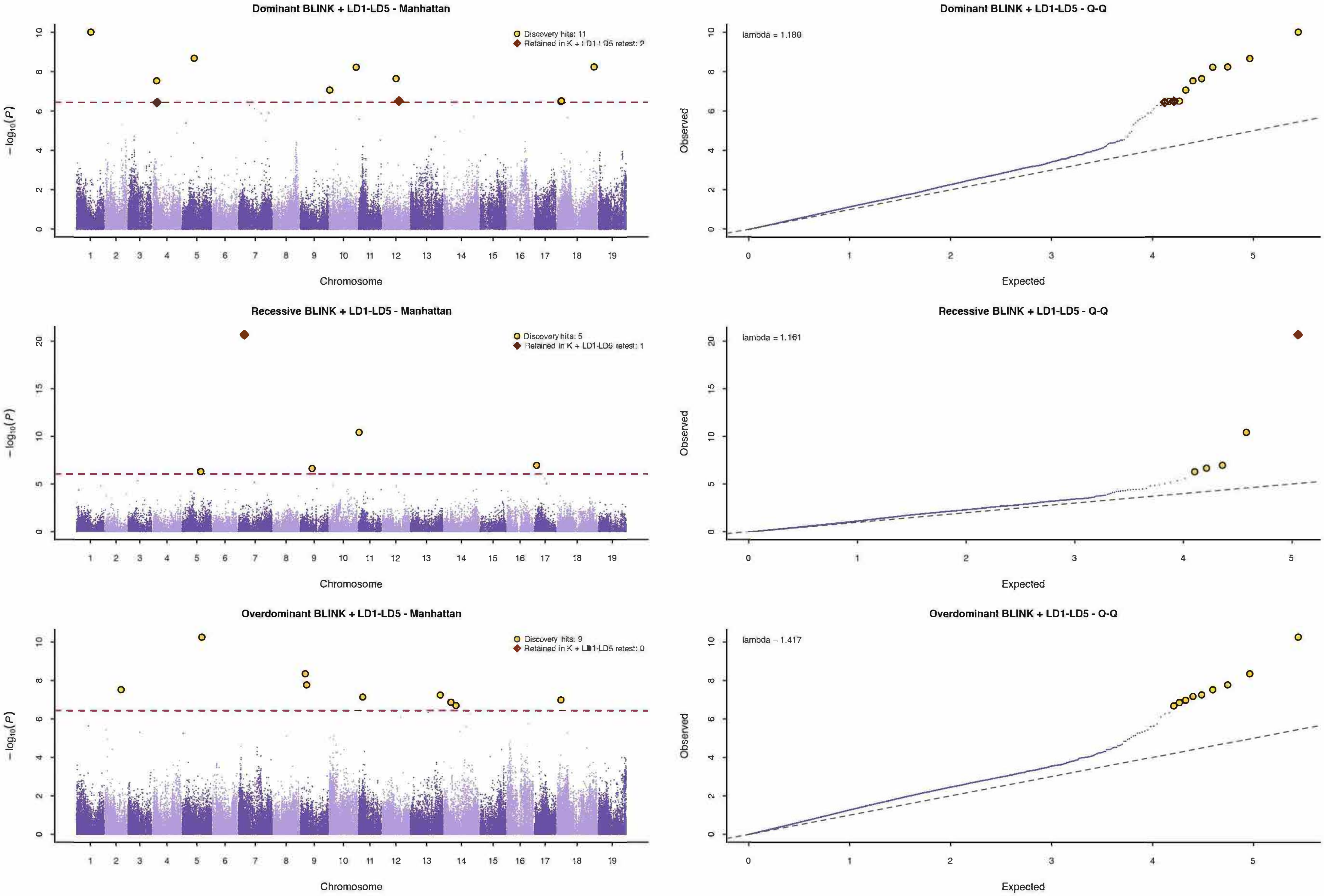


Dashed red line: 5% trait-specific Bonferroni threshold. Yellow circles: discovery hits. Orange diamonds: loci retained in the K + LD1-LD5 selected-candidate retest.
