## Supplementary File 3 for "Uncovering the genetic basis of agronomic traits in over 1,000 grapevine genotypes derived from a disease resistance breeding program"

### Budbreak

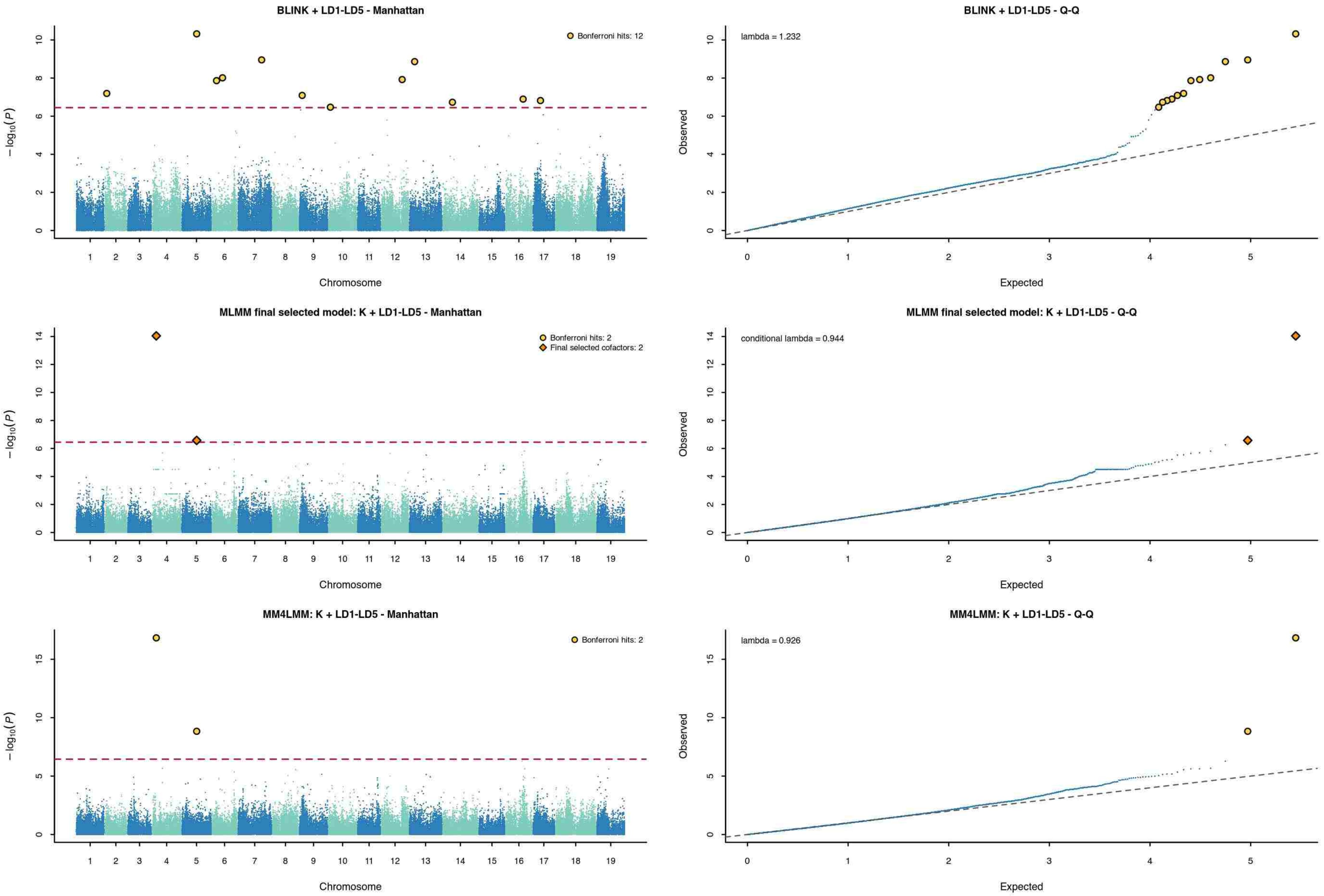


Dashed red line: 5% trait-specific Bonferroni threshold. Yellow circles: significant loci. Orange diamonds (MLMM): final selected cofactors.

### Flowering

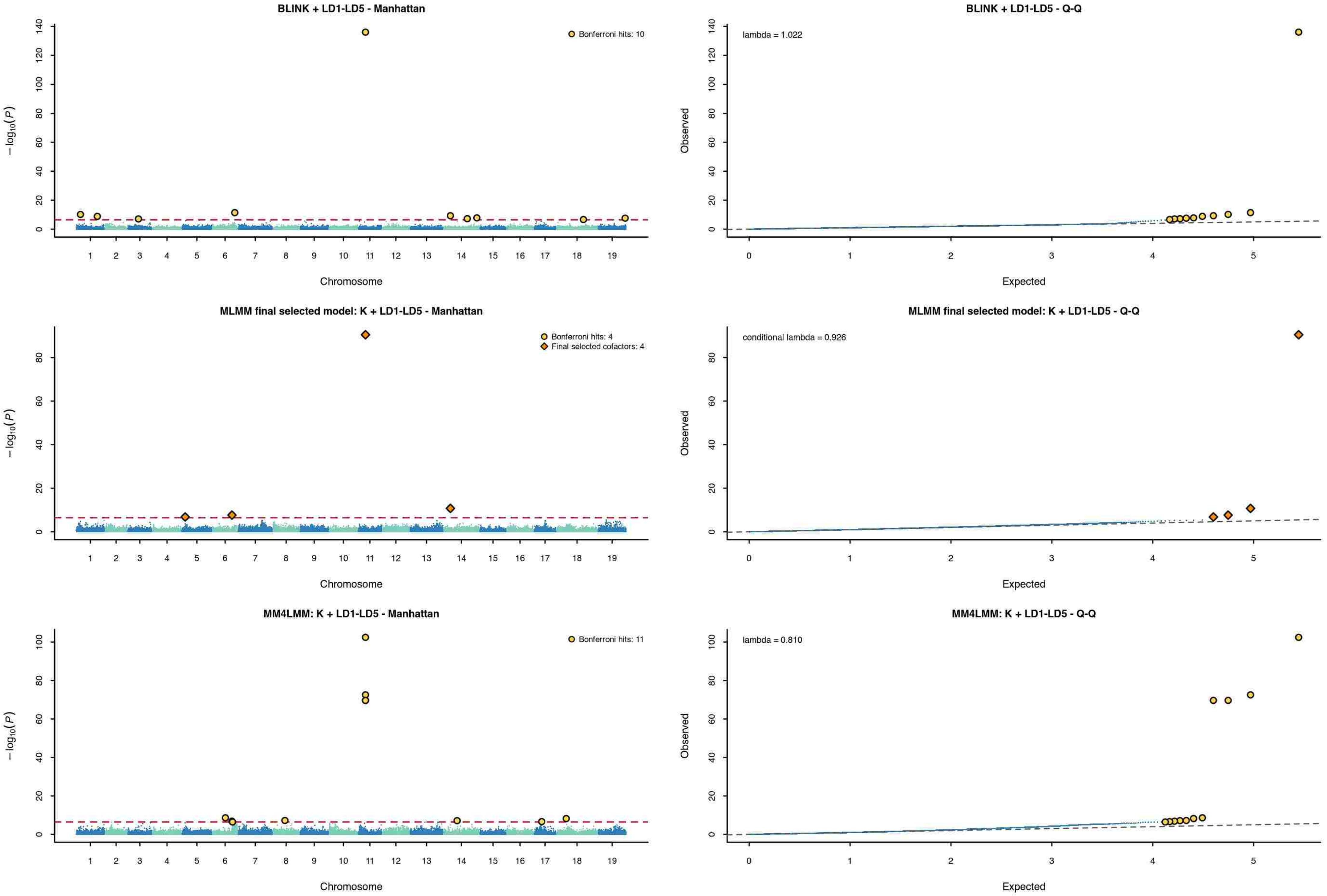


Dashed red line: 5% trait-specific Bonferroni threshold. Yellow circles: significant loci. Orange diamonds (MLMM): final selected cofactors.

### Veraison

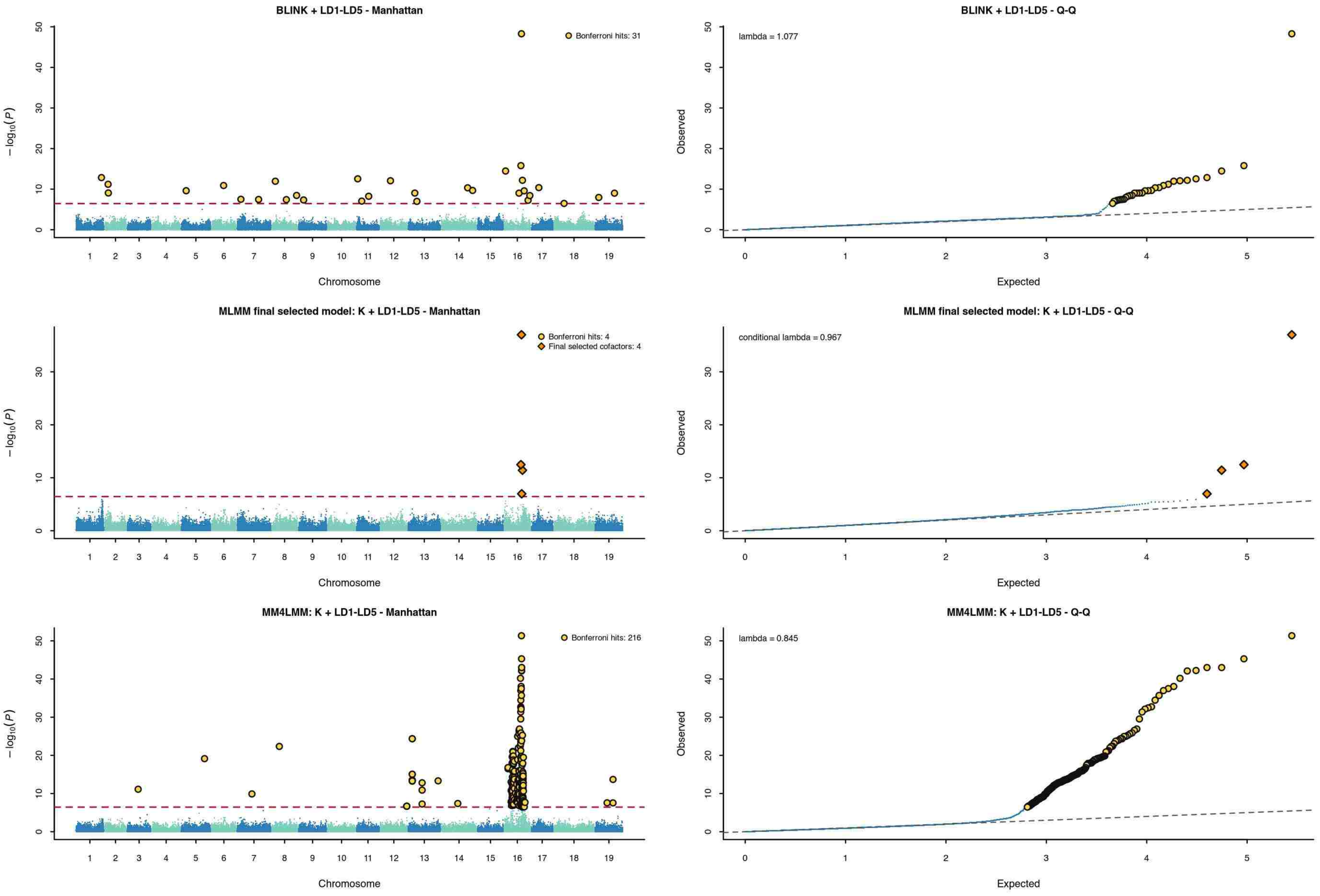


Dashed red line: 5% trait-specific Bonferroni threshold. Yellow circles: significant loci. Orange diamonds (MLMM): final selected cofactors.

### Harvest date

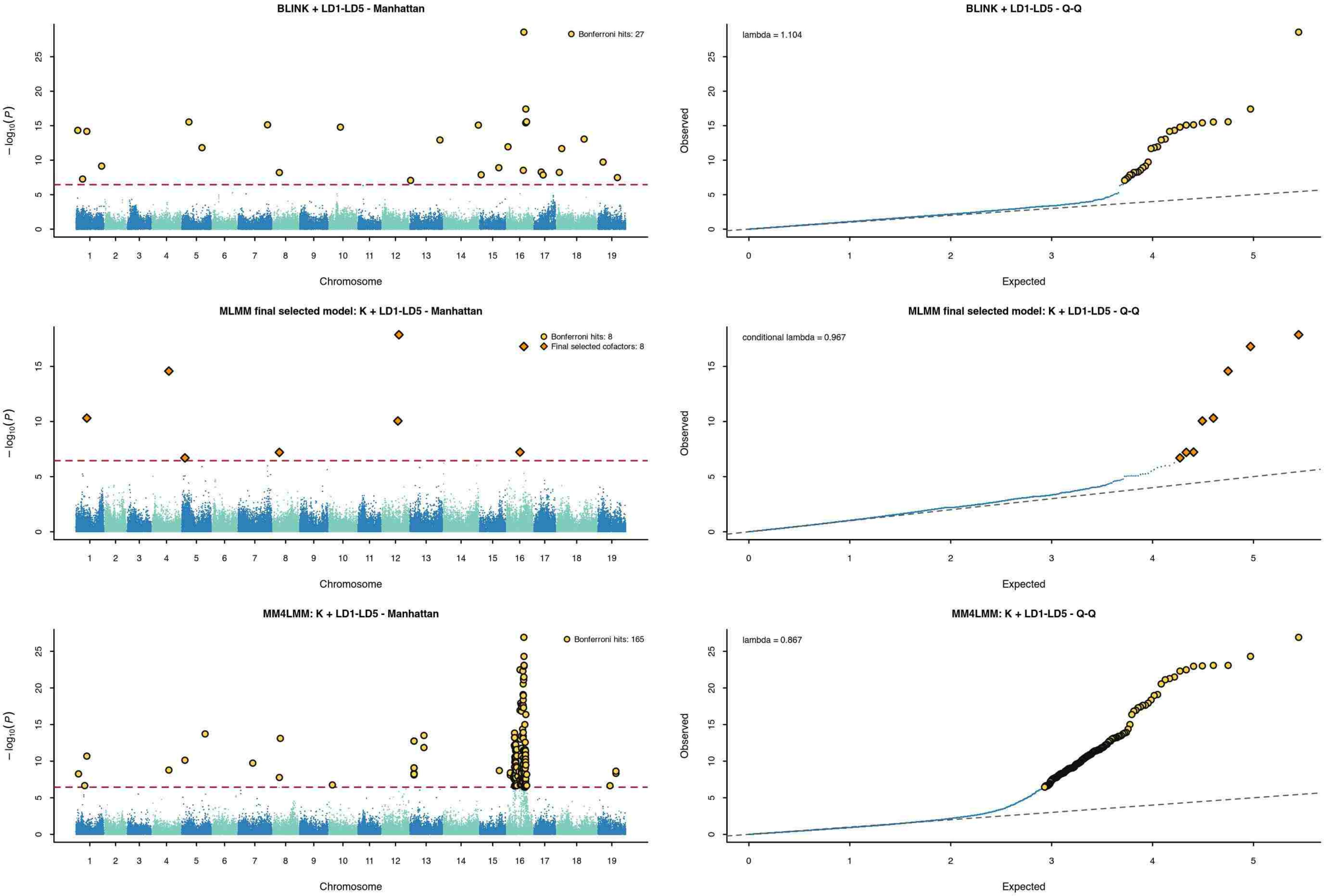


Dashed red line: 5% trait-specific Bonferroni threshold. Yellow circles: significant loci. Orange diamonds (MLMM): final selected cofactors.

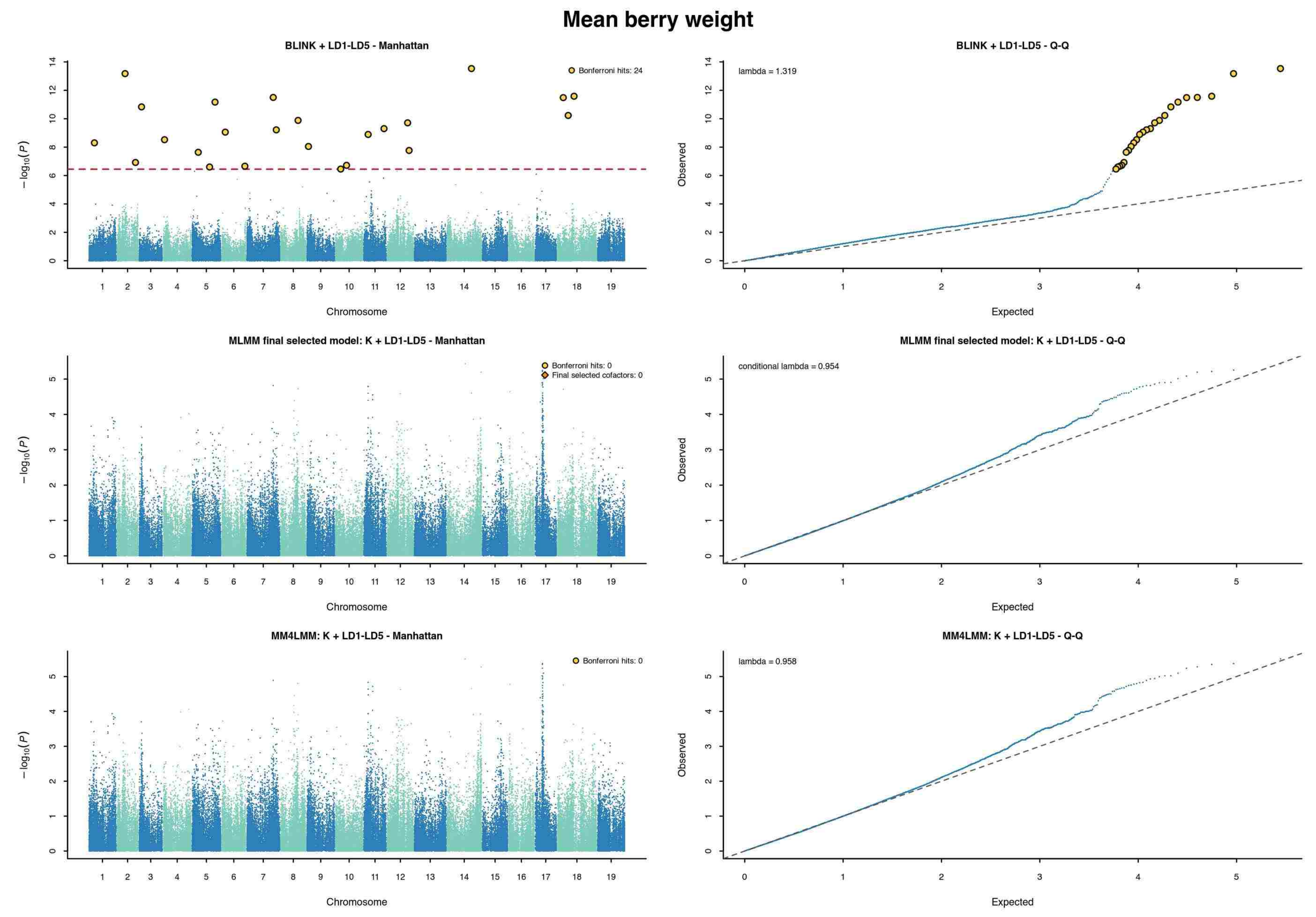


Dashed red line: 5% trait-specific Bonferroni threshold. Yellow circles: significant loci. Orange diamonds (MLMM): final selected cofactors.

### Berry sugar content

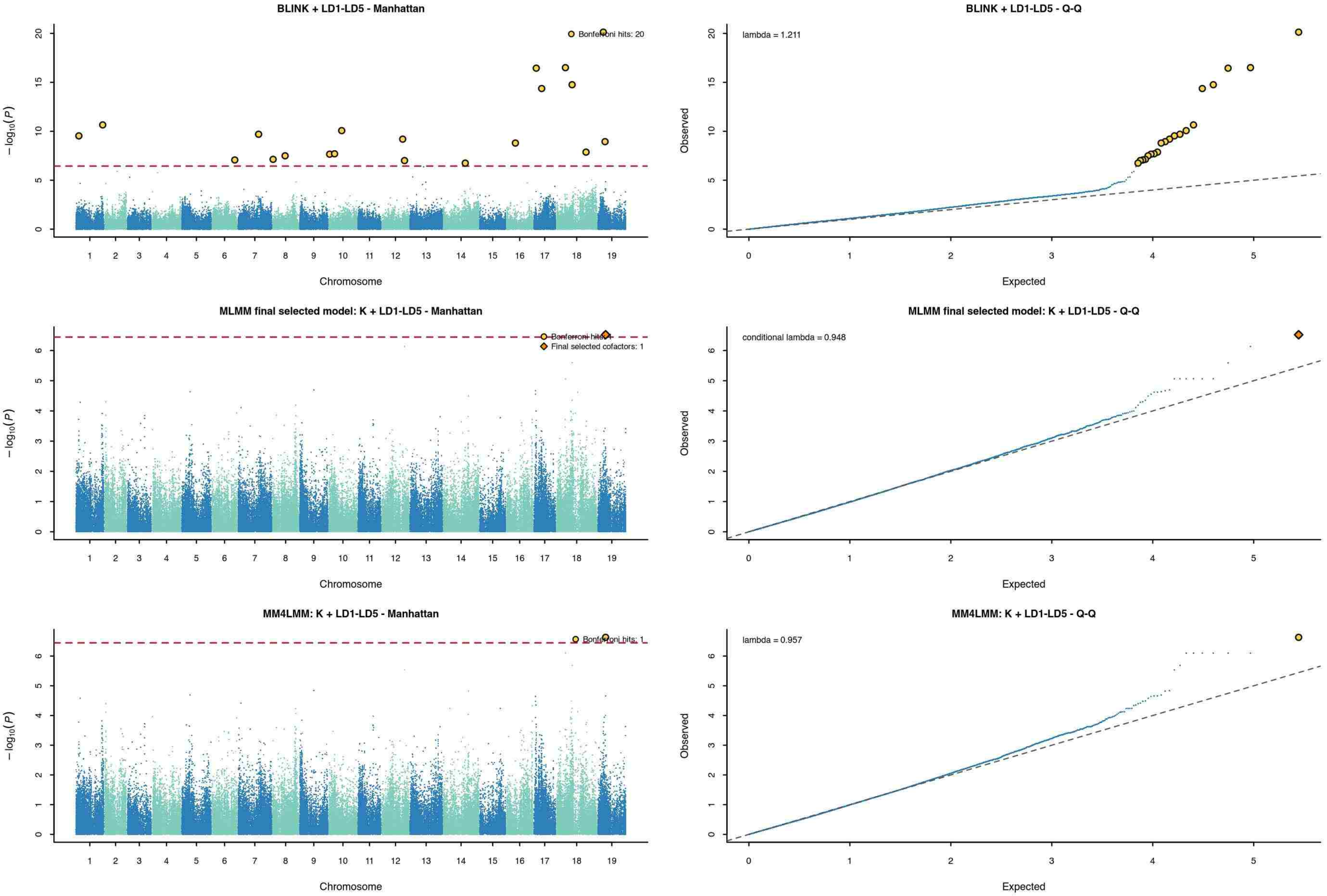


Dashed red line: 5% trait-specific Bonferroni threshold. Yellow circles: significant loci. Orange diamonds (MLMM): final selected cofactors.

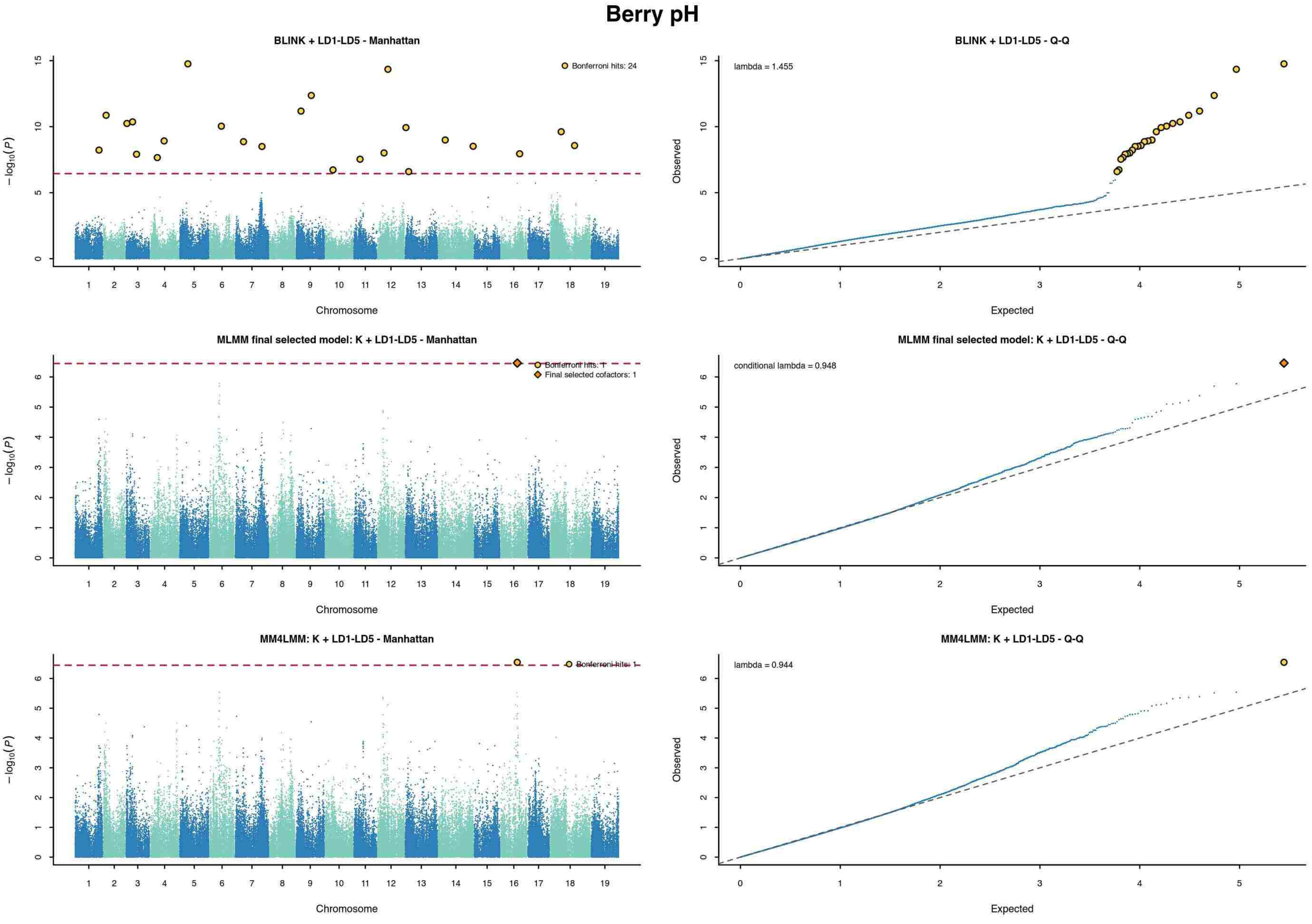


Dashed red line: 5% trait-specific Bonferroni threshold. Yellow circles: significant loci. Orange diamonds (MLMM): final selected cofactors.

### Titratable acidity

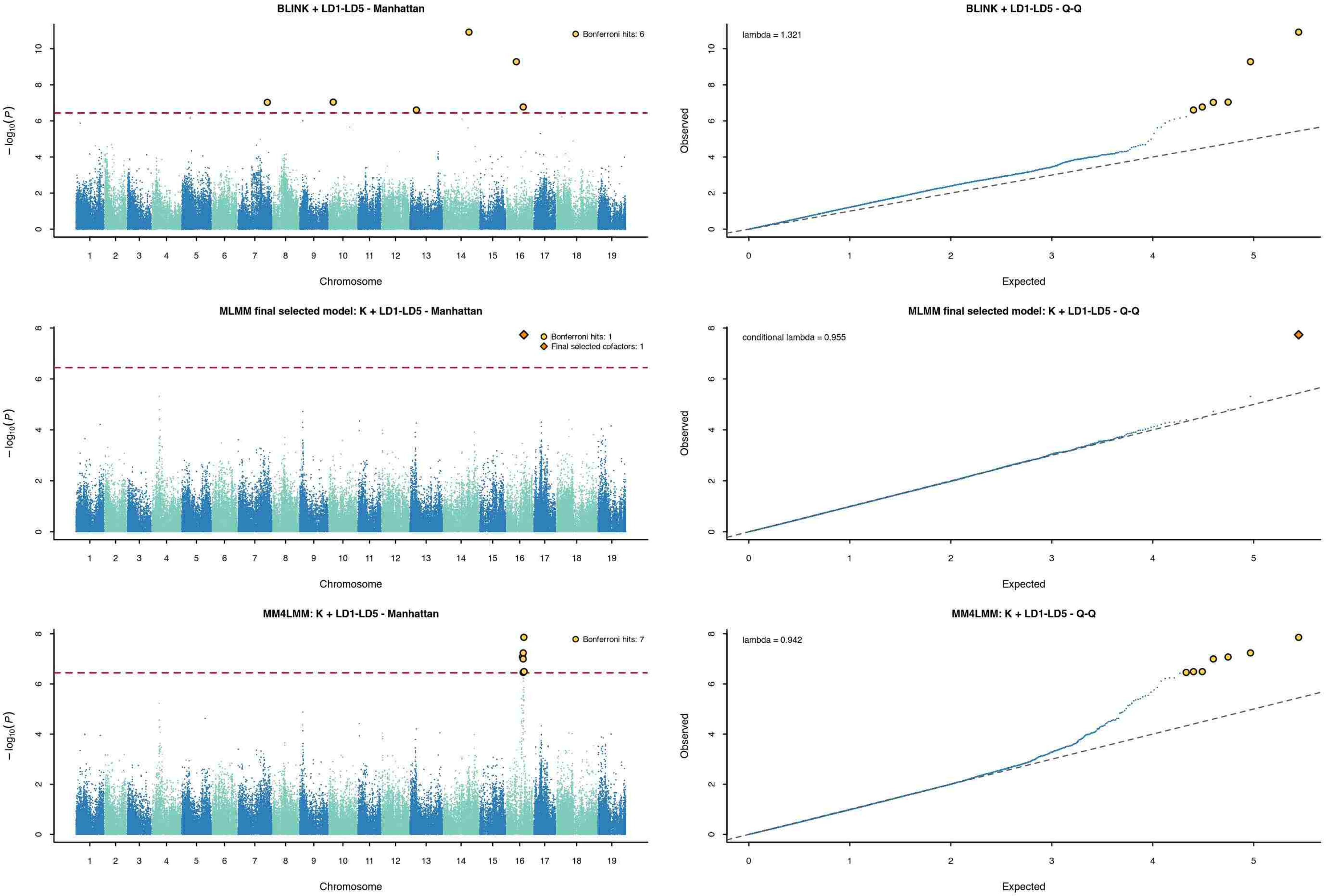


Dashed red line: 5% trait-specific Bonferroni threshold. Yellow circles: significant loci. Orange diamonds (MLMM): final selected cofactors.

### Cluster compactness

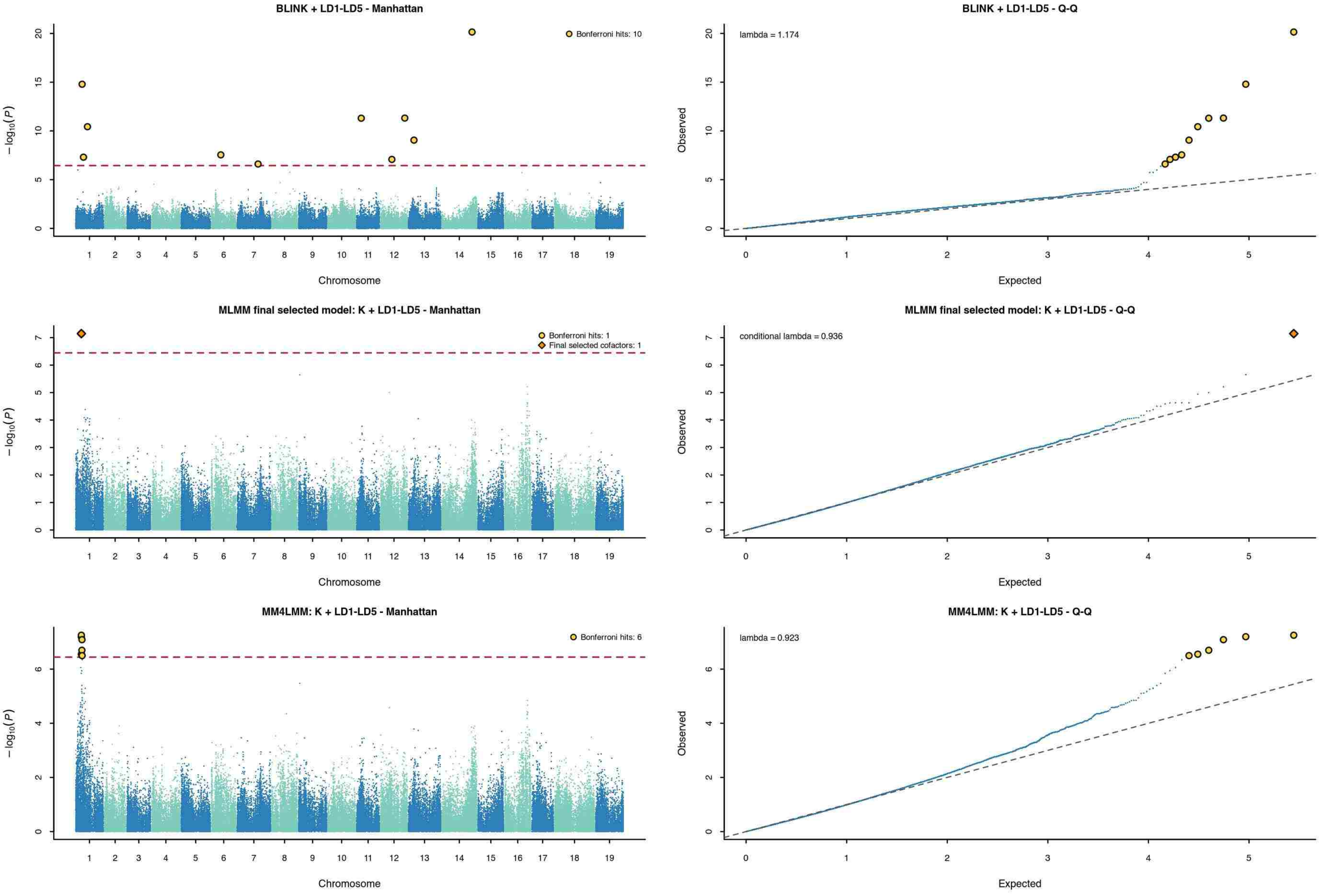


Dashed red line: 5% trait-specific Bonferroni threshold. Yellow circles: significant loci. Orange diamonds (MLMM): final selected cofactors.

### Clusters per plant

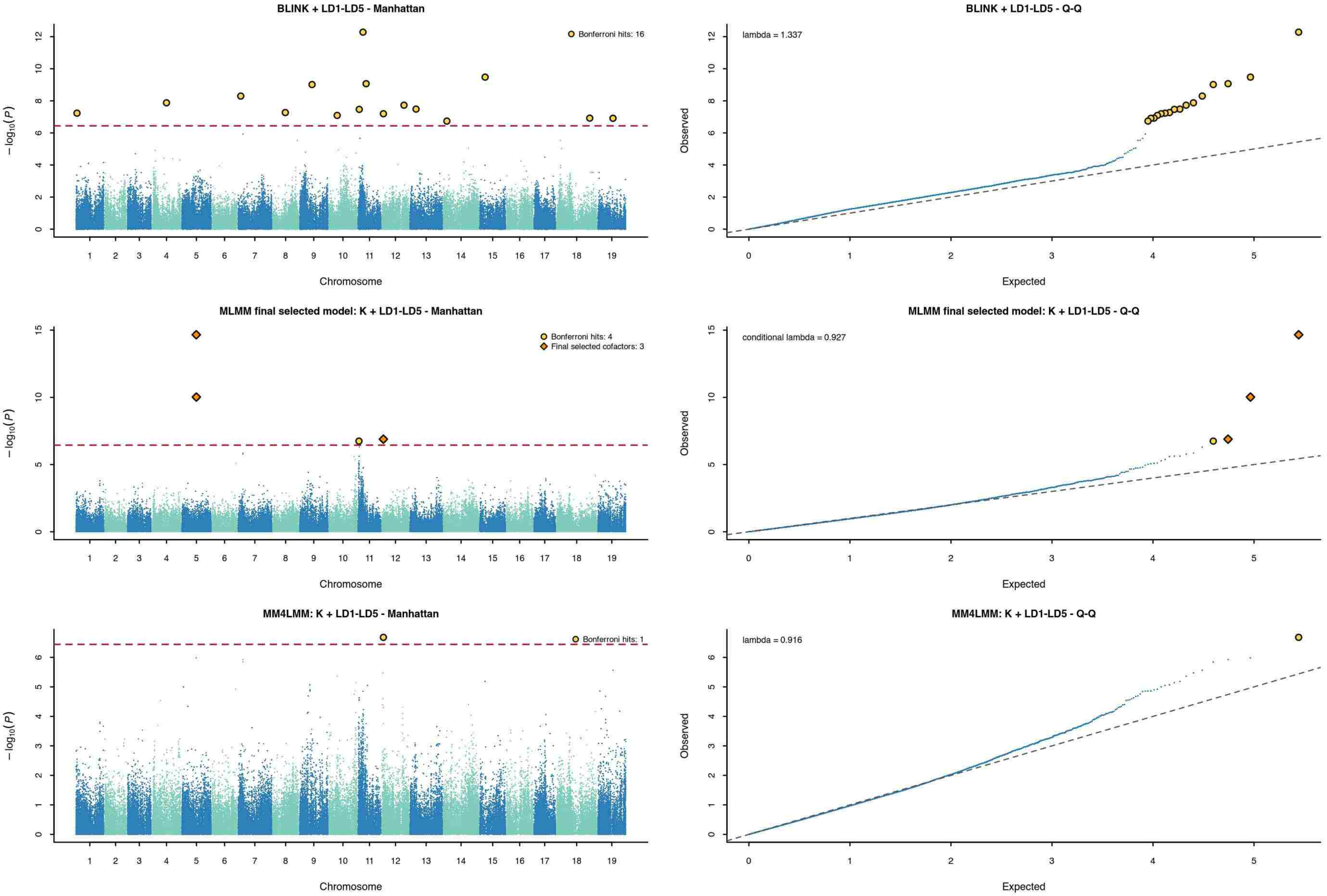


Dashed red line: 5% trait-specific Bonferroni threshold. Yellow circles: significant loci. Orange diamonds (MLMM): final selected cofactors.

### Cluster weight per plant

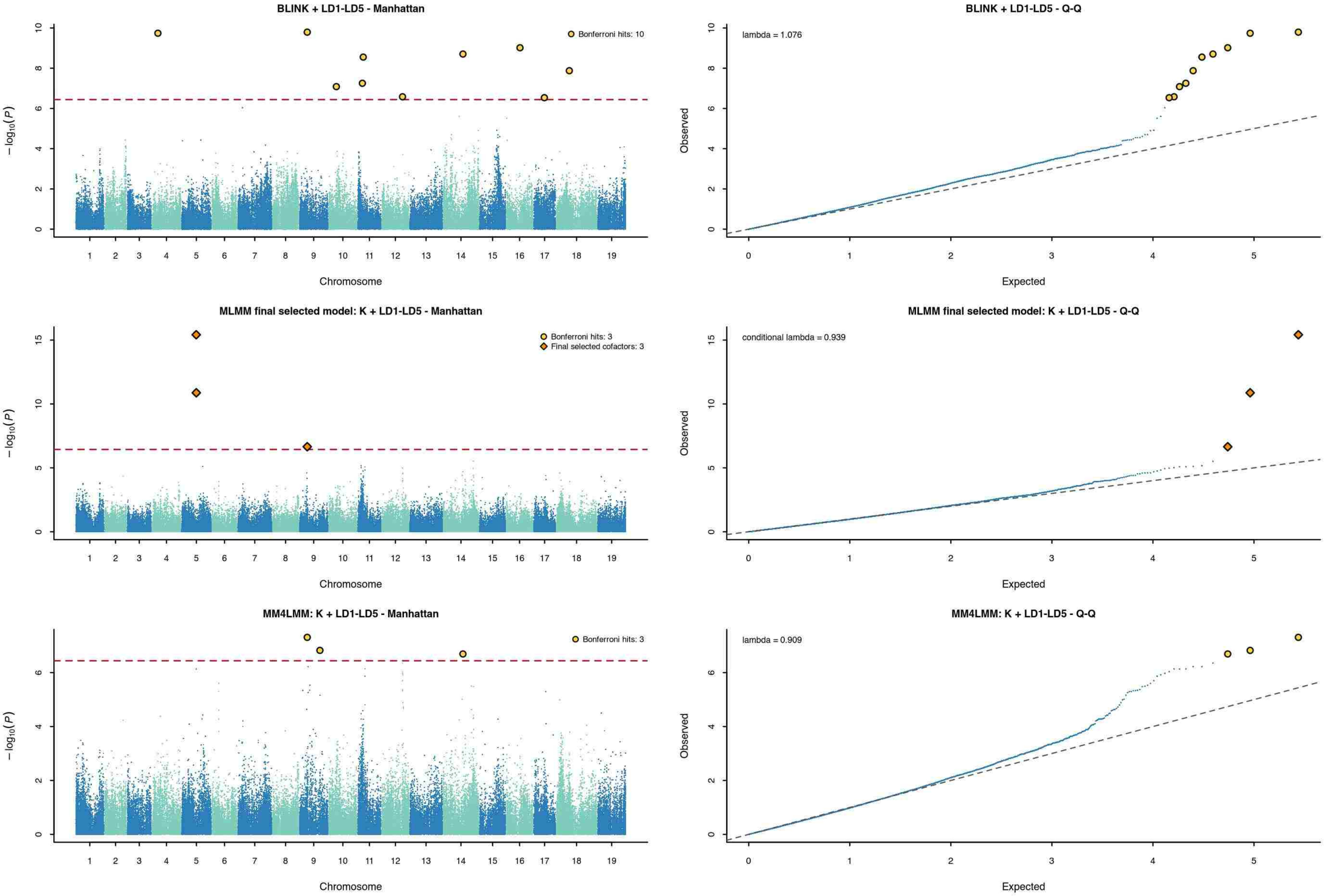


Dashed red line: 5% trait-specific Bonferroni threshold. Yellow circles: significant loci. Orange diamonds (MLMM): final selected cofactors.

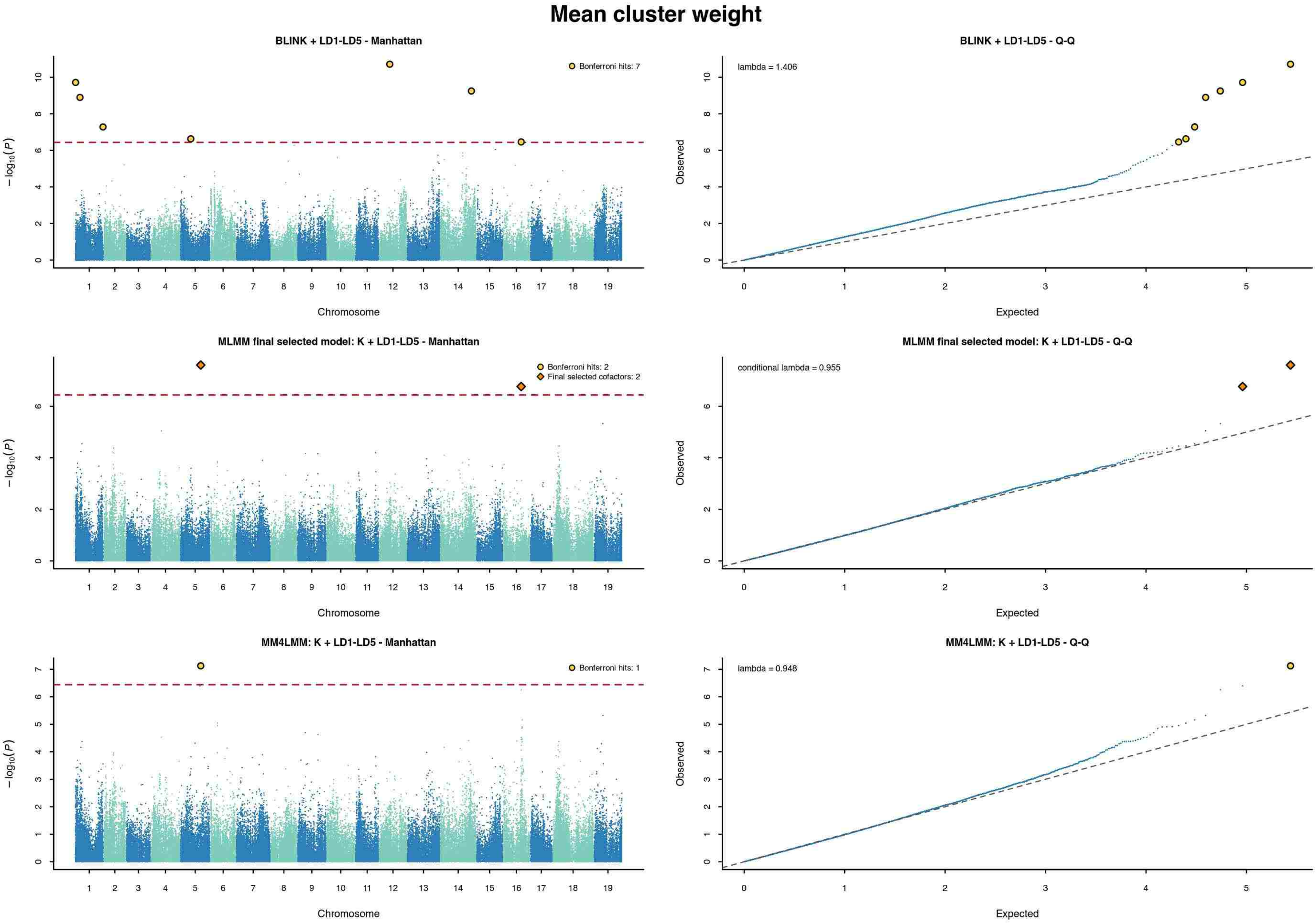


Dashed red line: 5% trait-specific Bonferroni threshold. Yellow circles: significant loci. Orange diamonds (MLMM): final selected cofactors.

Yield per $m^2$

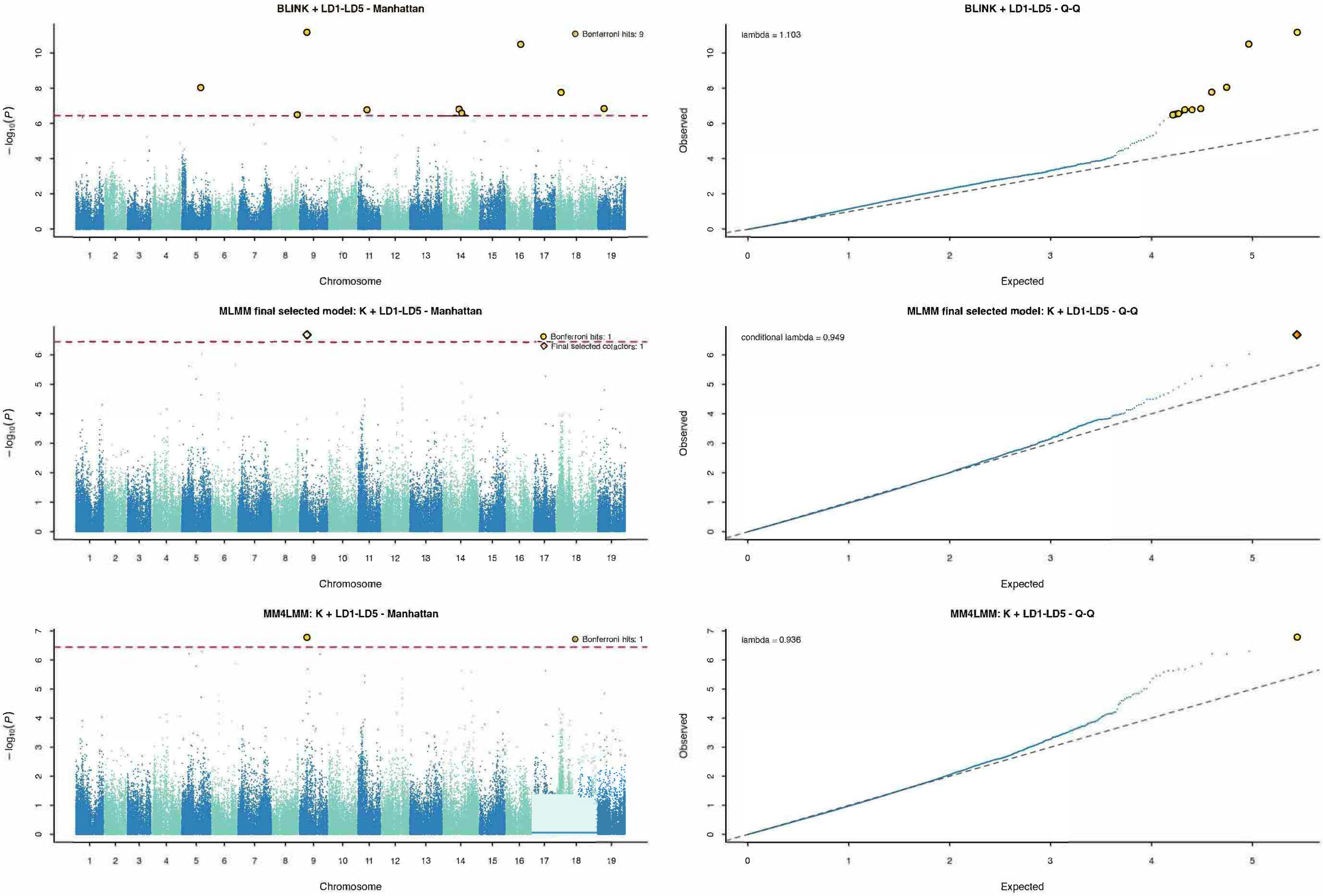


Dashed red line: 5% trait-specific Bonferroni threshold. Yellow circles: significant loci. Orange diamonds (MLMM): final selected cofactors.
