## Supplementary File 2 for "Uncovering the genetic basis of agronomic traits in over 1,000 grapevine genotypes derived from a disease resistance breeding program"

### Population-adjusted mpQTL diagnostics — Budbreak (BUD_DATE)

n tested = 8,054; significant markers = 4; nominal support intervals = 4; lambdaGC = 1.111

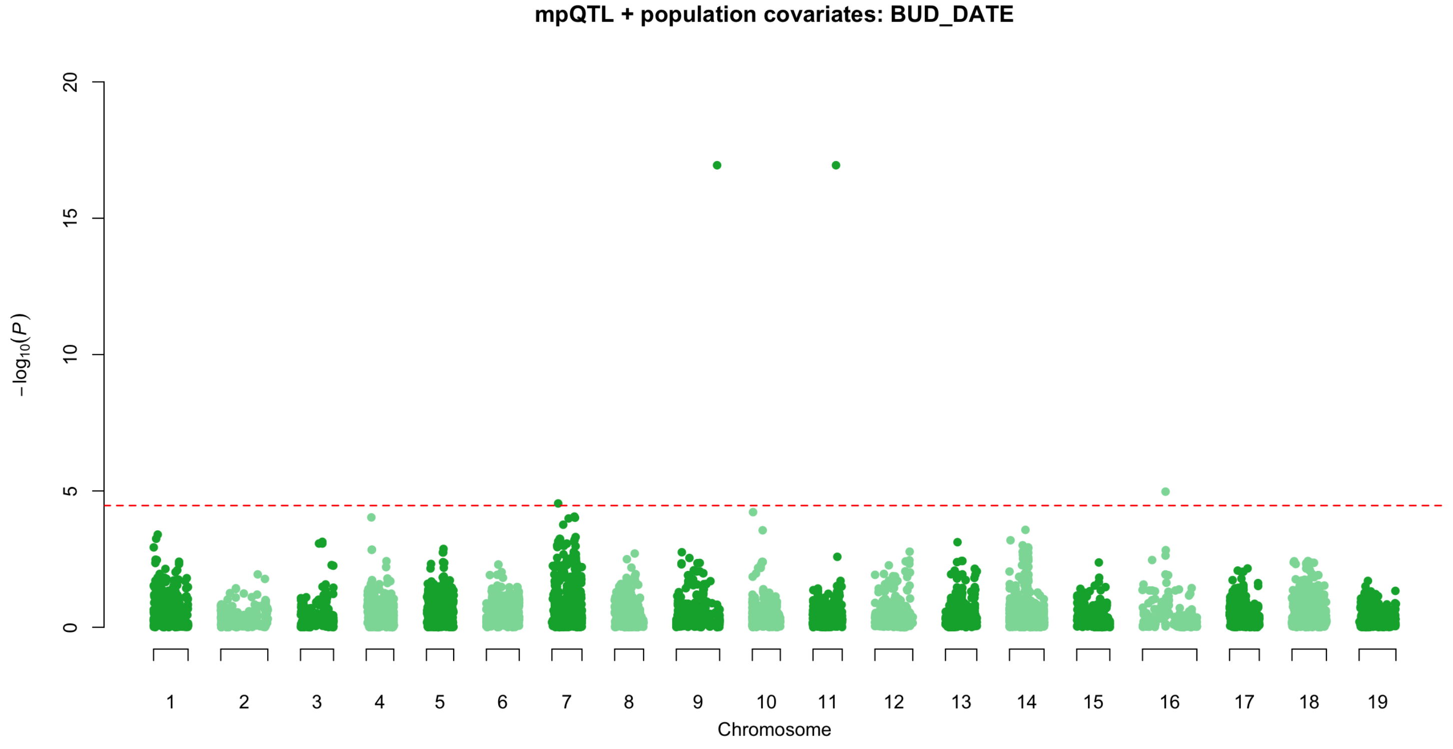


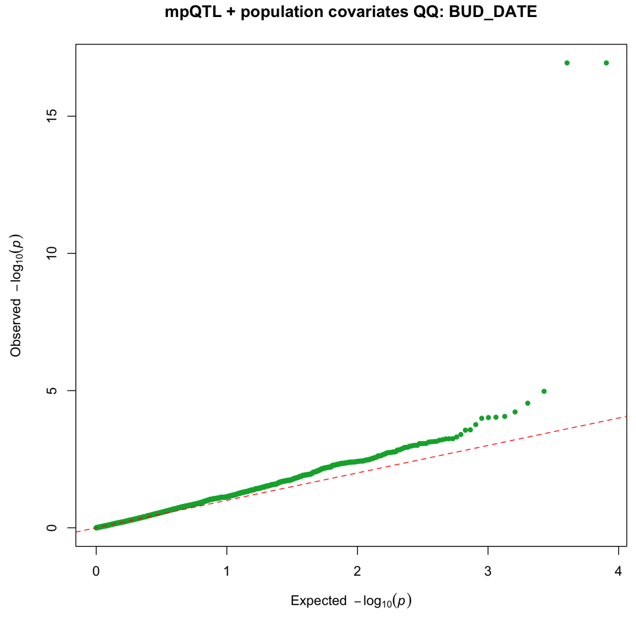


Model: realized kinship + 12 fixed population indicators; mpQTL 0.6.2 approximate P3D/EMMAX; Li–Ji threshold P = 3.4424437e-5.

Nominal positional supports are heuristic 1.5-log10(P)-drop intervals; see Supplementary Tables 10-12 and 18 for complete results and limitations.

### Population-adjusted mpQTL diagnostics — Flowering (FLO_50)

n tested = 8,050; significant markers = 10; nominal support intervals = 5; lambdaGC = 1.084

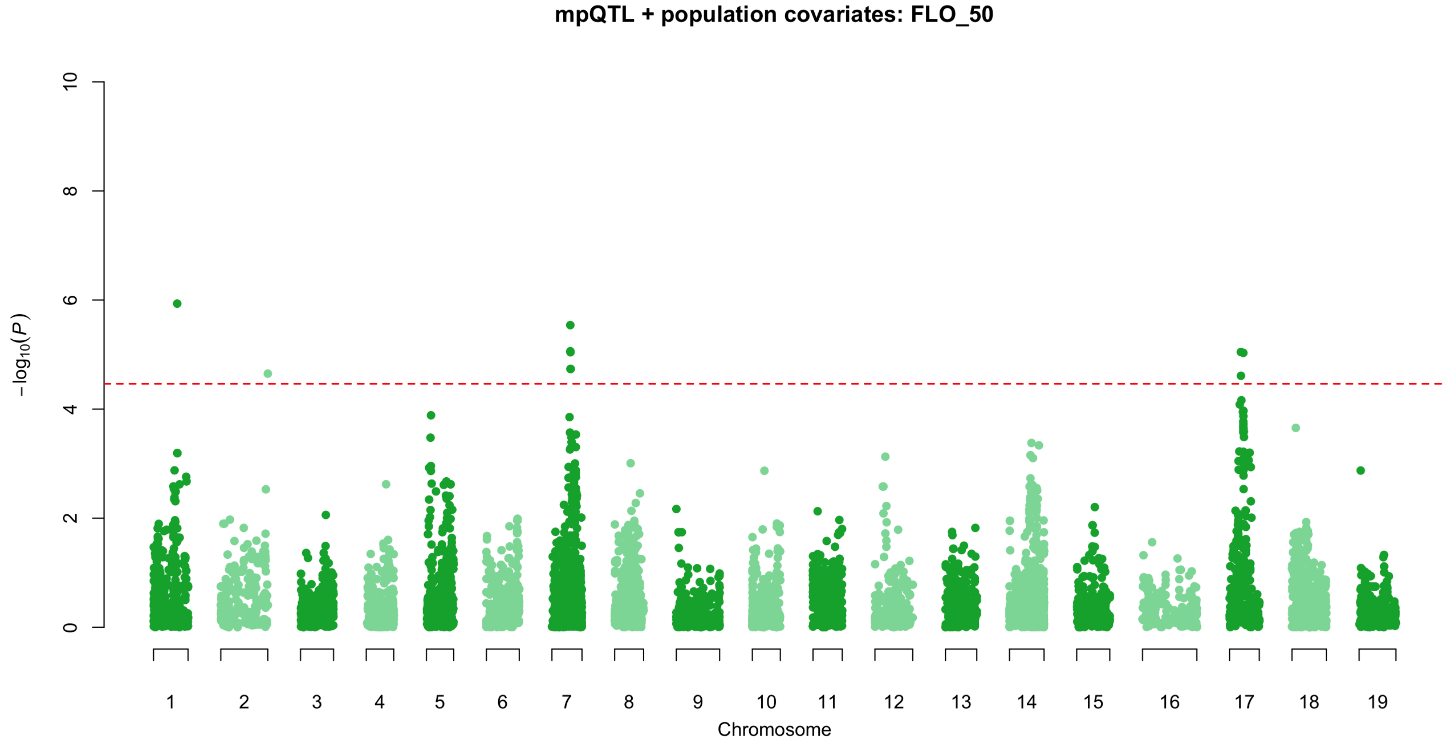


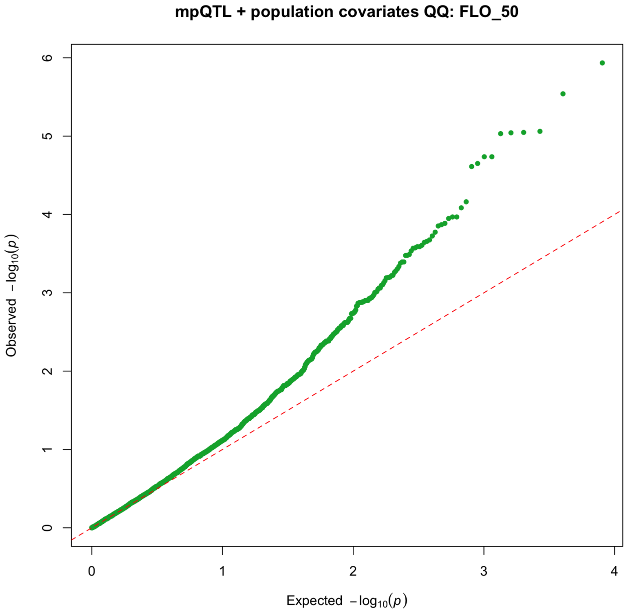


Model: realized kinship + 12 fixed population indicators; mpQTL 0.6.2 approximate P3D/EMMAX; Li–Ji threshold P = 3.4424437e-5.

Nominal positional supports are heuristic 1.5-log10(P)-drop intervals; see Supplementary Tables 10-12 and 18 for complete results and limitations.

### Population-adjusted mpQTL diagnostics — Veraison (VER_50)

n tested = 8,050; significant markers = 6; nominal support intervals = 4; lambdaGC = 1.155

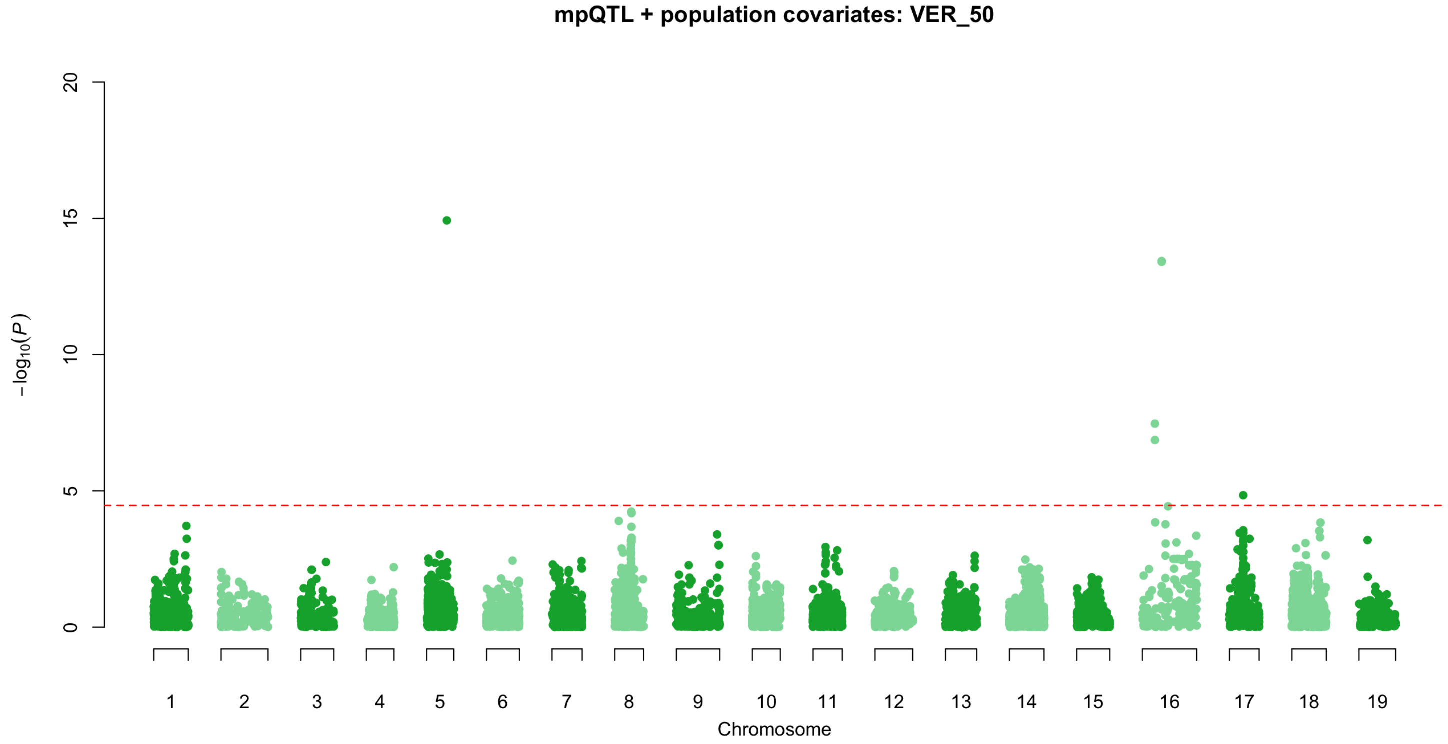


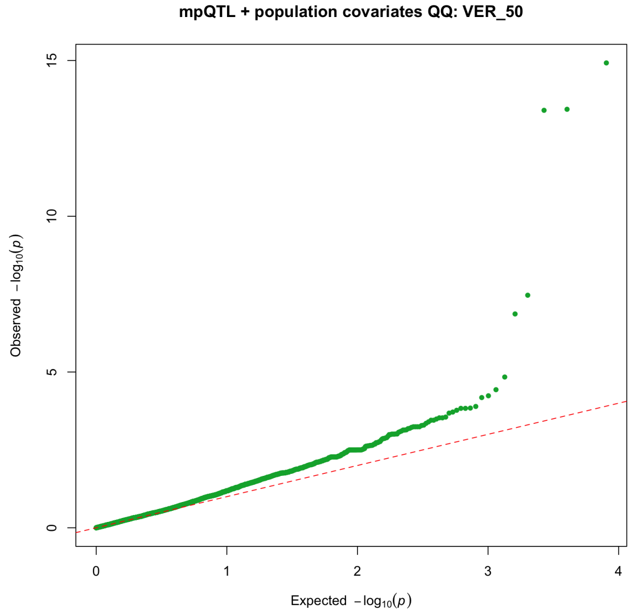


Model: realized kinship + 12 fixed population indicators; mpQTL 0.6.2 approximate P3D/EMMAX; Li–Ji threshold P = 3.4424437e-5.

Nominal positional supports are heuristic 1.5-log10(P)-drop intervals; see Supplementary Tables 10-12 and 18 for complete results and limitations.

### Population-adjusted mpQTL diagnostics — Harvest date (HARVEST_DATE)

n tested = 8,050; significant markers = 6; nominal support intervals = 4; lambdaGC = 1.213

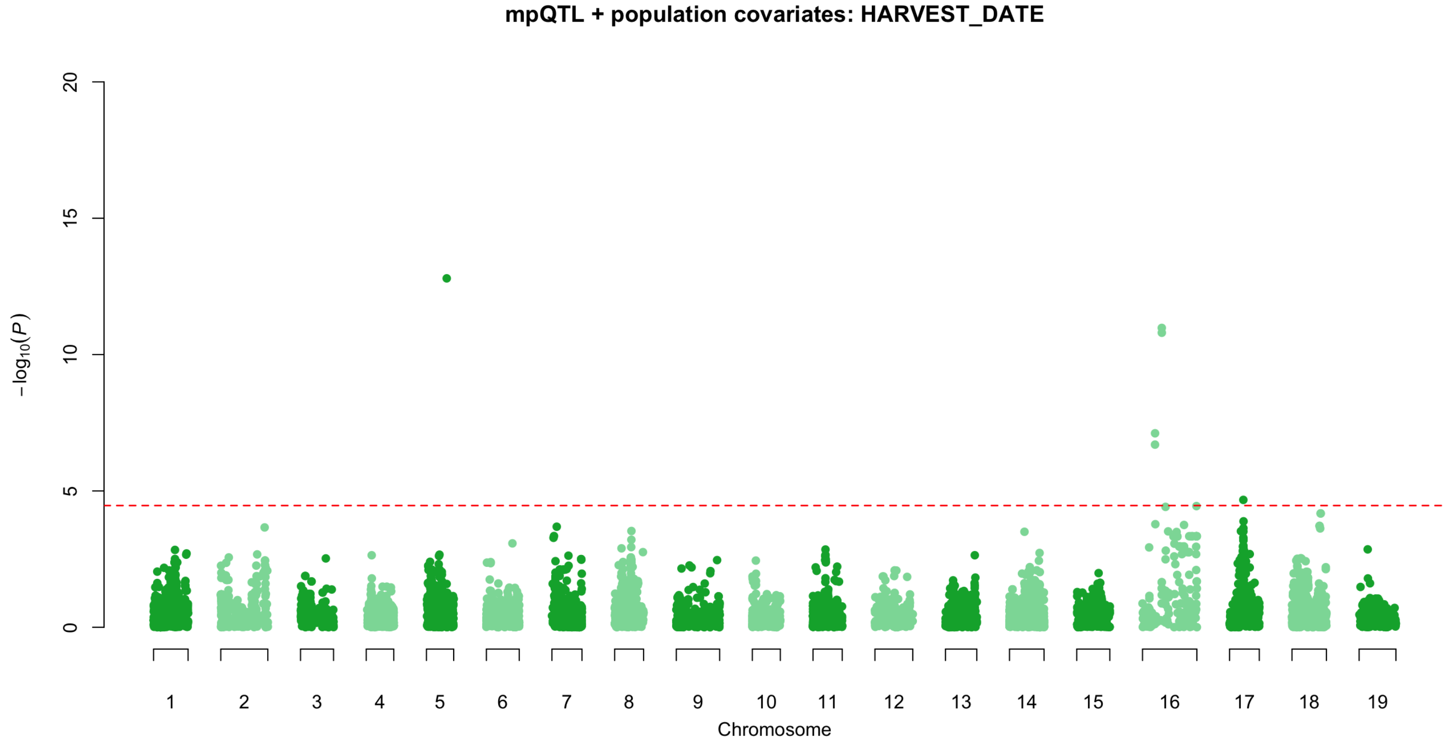


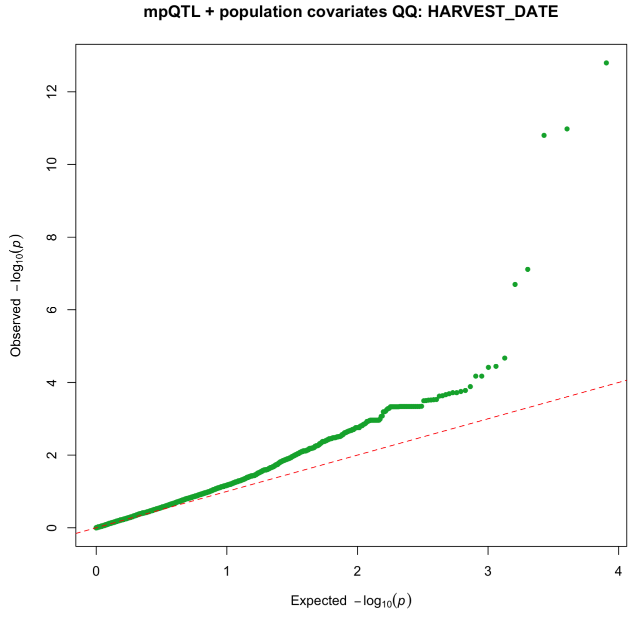


Model: realized kinship + 12 fixed population indicators; mpQTL 0.6.2 approximate P3D/EMMAX; Li–Ji threshold P = 3.4424437e-5.

Nominal positional supports are heuristic 1.5-log10(P)-drop intervals; see Supplementary Tables 10-12 and 18 for complete results and limitations.

### Population-adjusted mpQTL diagnostics — Mean berry weight (SBER_W_g)

n tested = 8,050; significant markers = 12; nominal support intervals = 5; lambdaGC = 1.428

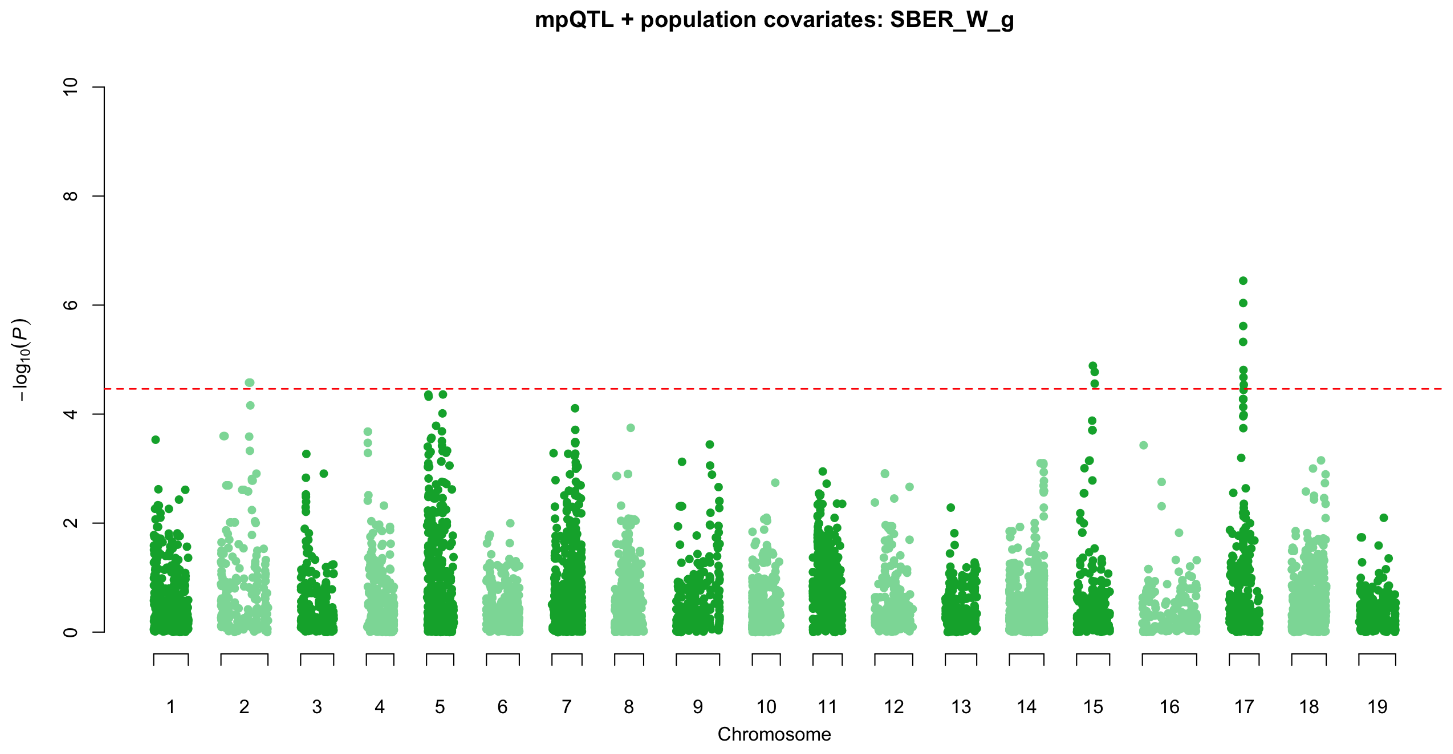


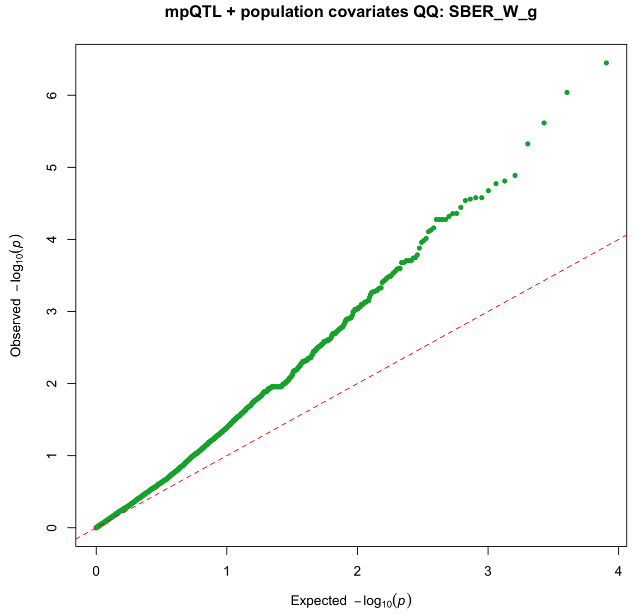


Model: realized kinship + 12 fixed population indicators; mpQTL 0.6.2 approximate P3D/EMMAX; Li–Ji threshold P = 3.4424437e-5.

Nominal positional supports are heuristic 1.5-log10(P)-drop intervals; see Supplementary Tables 10-12 and 18 for complete results and limitations.

### Population-adjusted mpQTL diagnostics — Berry sugar content (TSS)

n tested = 8,050; significant markers = 8; nominal support intervals = 8; lambdaGC = 1.279

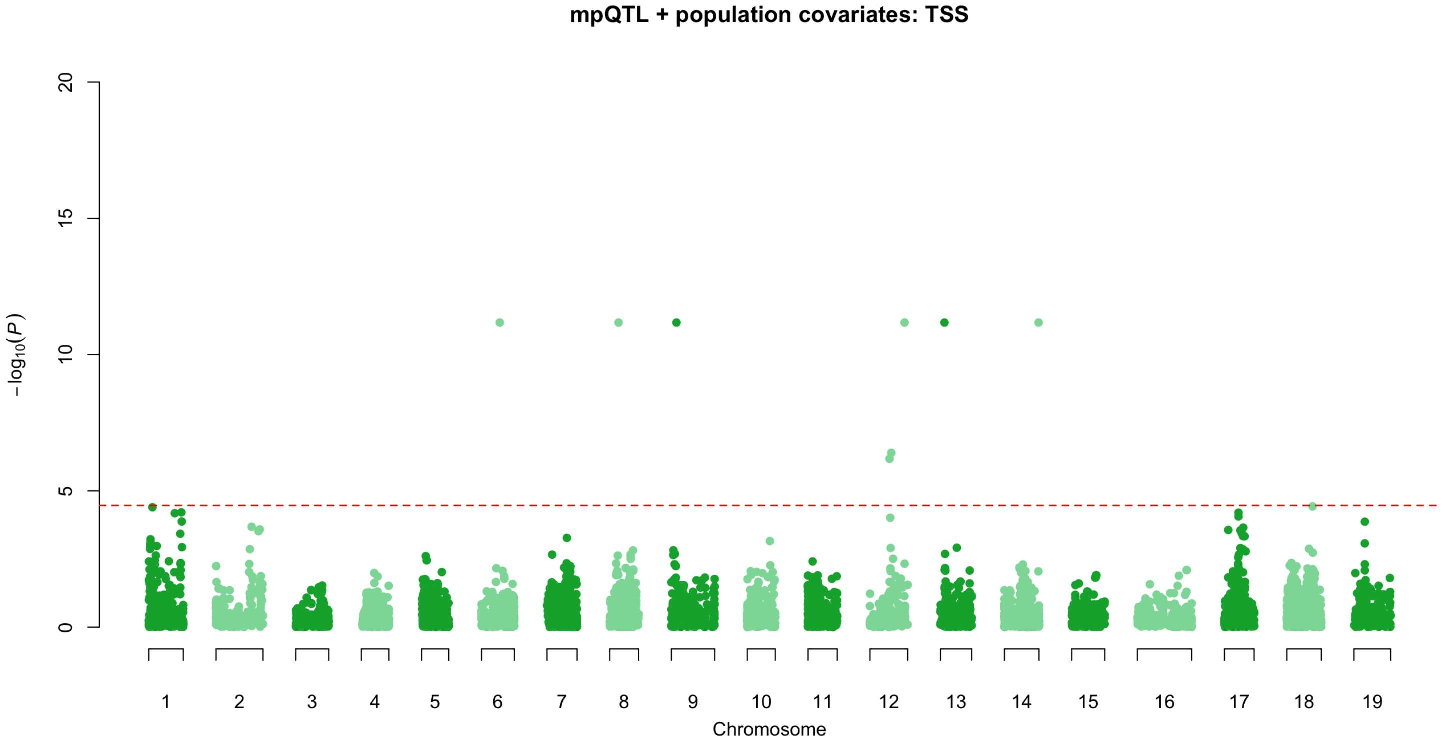


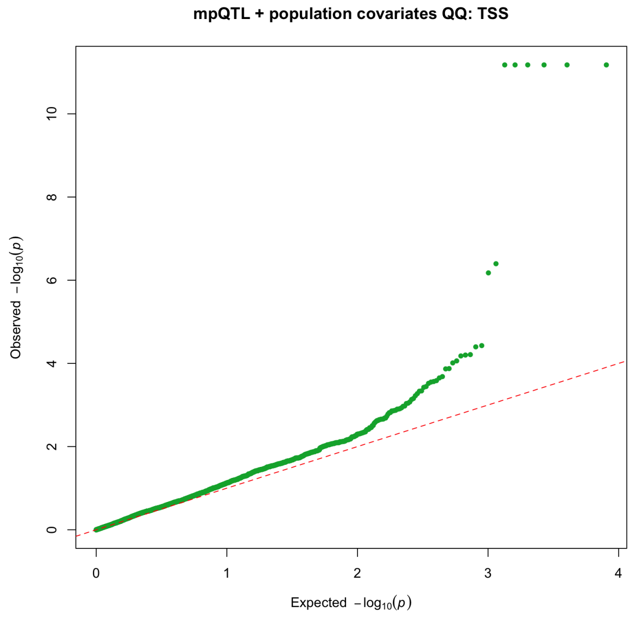


Model: realized kinship + 12 fixed population indicators; mpQTL 0.6.2 approximate P3D/EMMAX; Li–Ji threshold P = 3.4424437e-5.

Nominal positional supports are heuristic 1.5-log10(P)-drop intervals; see Supplementary Tables 10-12 and 18 for complete results and limitations.

### Population-adjusted mpQTL diagnostics — Berry pH (BERRY_pH)

n tested = 8,050; significant markers = 1; nominal support intervals = 1; lambdaGC = 1.132

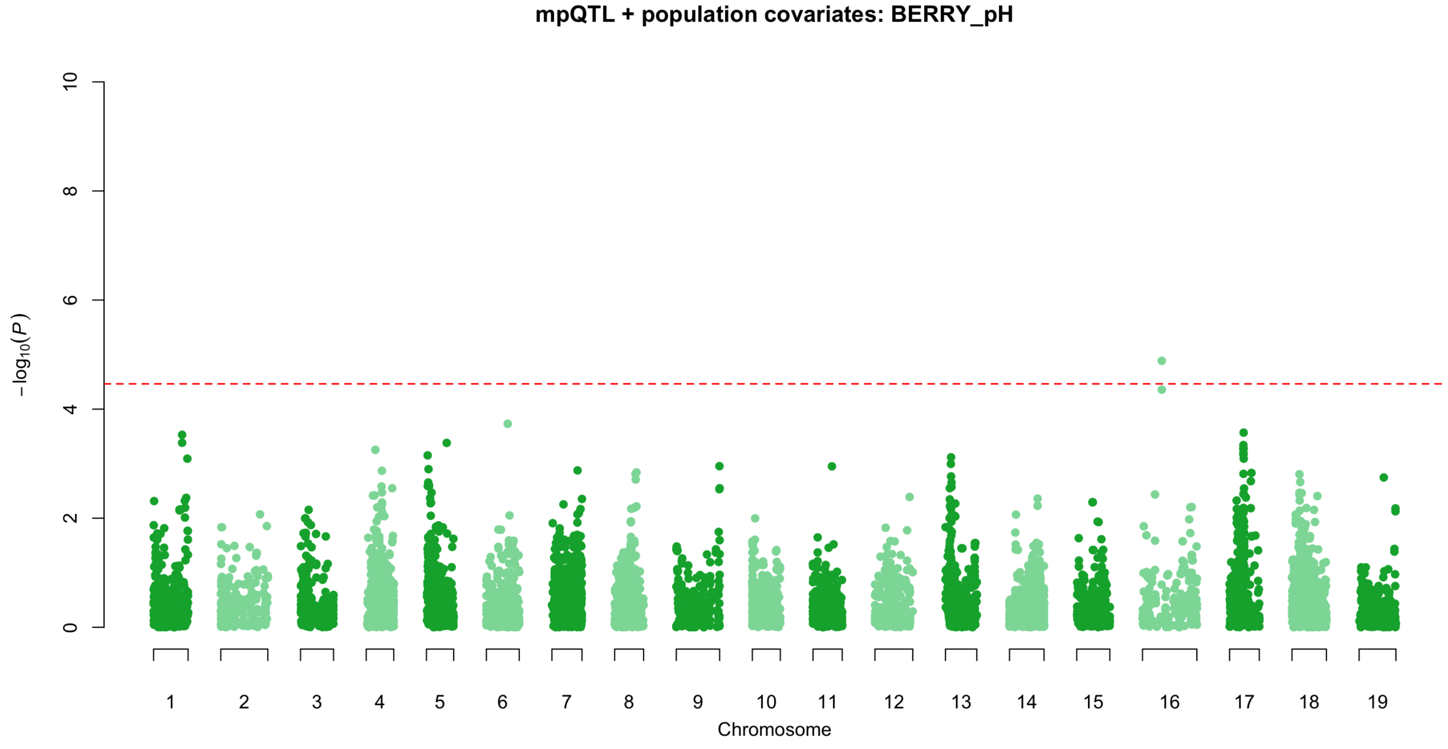


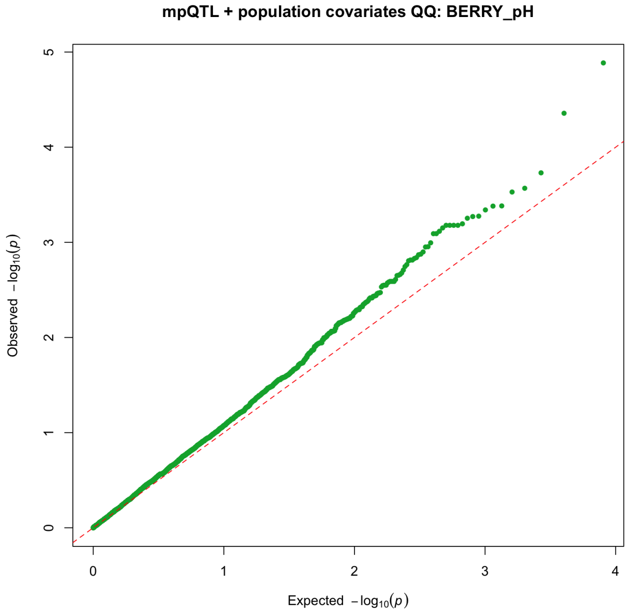


Model: realized kinship + 12 fixed population indicators; mpQTL 0.6.2 approximate P3D/EMMAX; Li–Ji threshold P = 3.4424437e-5.

Nominal positional supports are heuristic 1.5-log10(P)-drop intervals; see Supplementary Tables 10-12 and 18 for complete results and limitations.

### Population-adjusted mpQTL diagnostics — Titratable acidity (BER_TA_g)

n tested = 8,050; significant markers = 1; nominal support intervals = 1; lambdaGC = 1.373

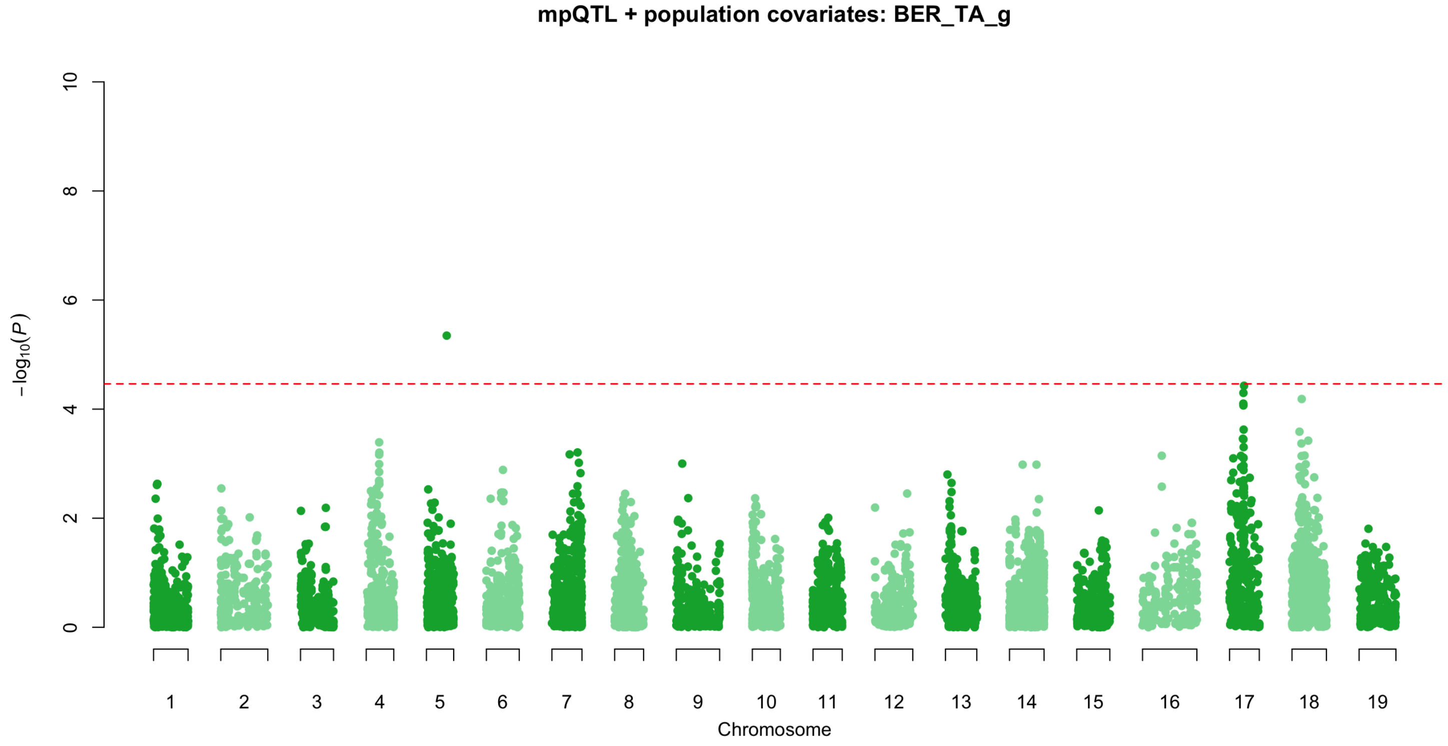


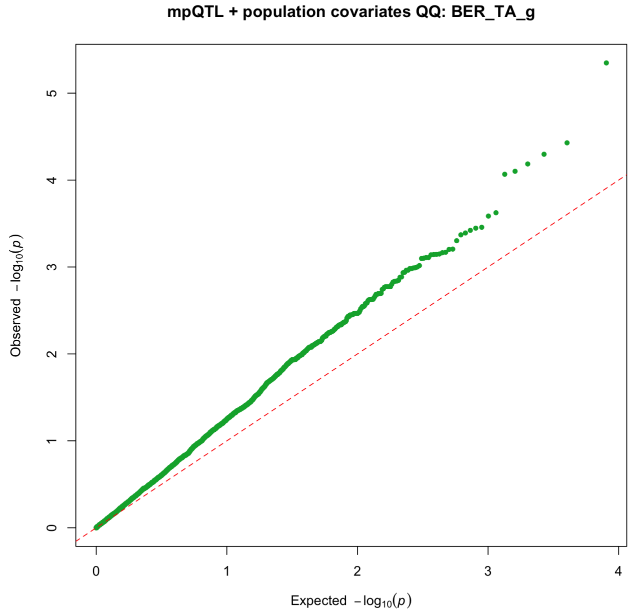


Model: realized kinship + 12 fixed population indicators; mpQTL 0.6.2 approximate P3D/EMMAX; Li–Ji threshold P = 3.4424437e-5.

Nominal positional supports are heuristic 1.5-log10(P)-drop intervals; see Supplementary Tables 10-12 and 18 for complete results and limitations.

### Population-adjusted mpQTL diagnostics — Cluster compactness (MORPHO_OIV_204)

n tested = 8,050; significant markers = 13; nominal support intervals = 5; lambdaGC = 1.075

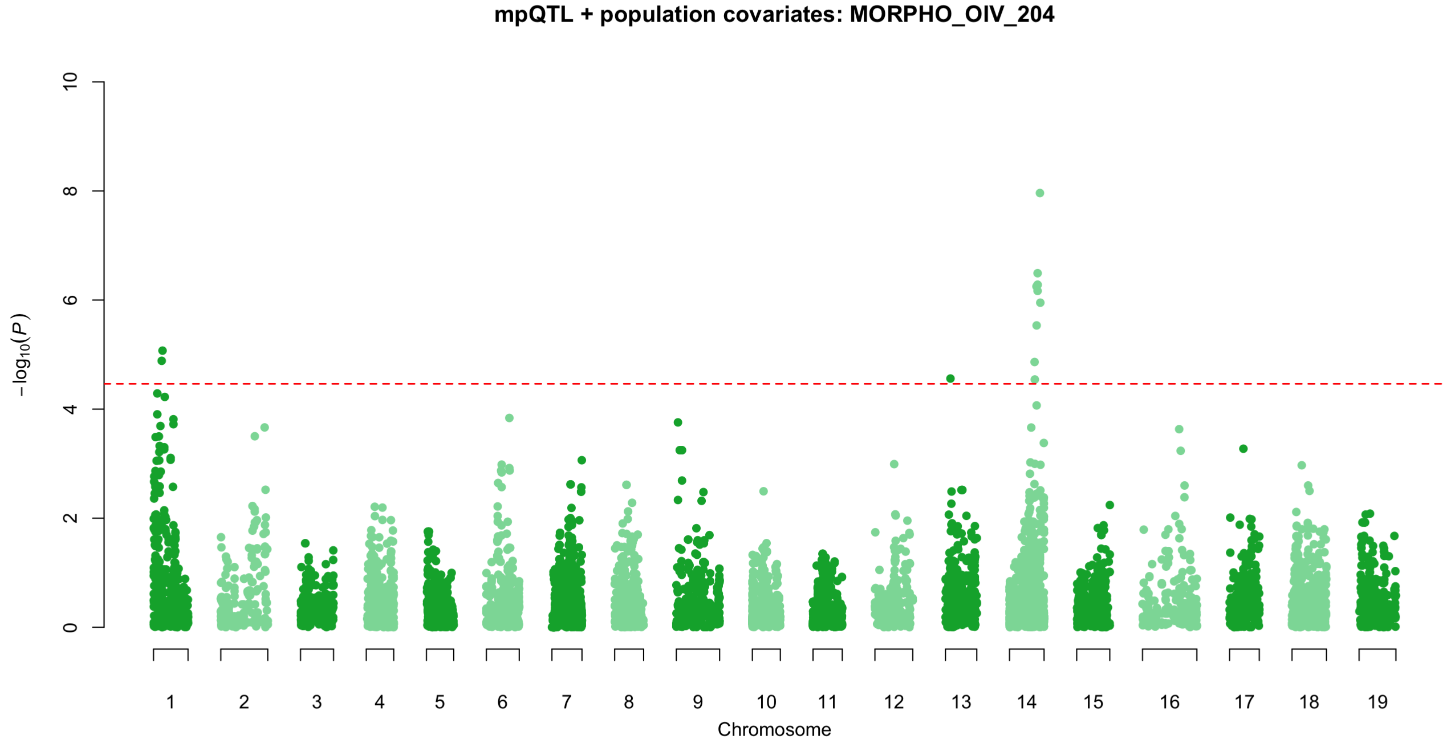


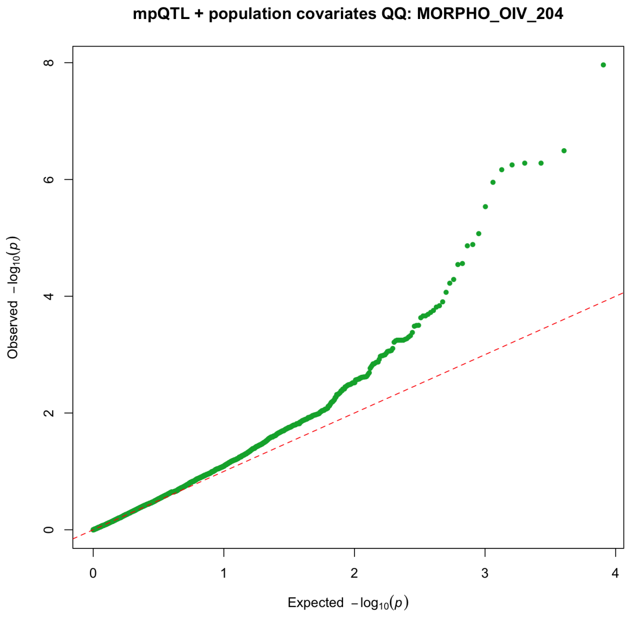


Model: realized kinship + 12 fixed population indicators; mpQTL 0.6.2 approximate P3D/EMMAX; Li–Ji threshold P = 3.4424437e-5.

Nominal positional supports are heuristic 1.5-log10(P)-drop intervals; see Supplementary Tables 10-12 and 18 for complete results and limitations.

### Population-adjusted mpQTL diagnostics — Clusters per plant (NB_CLUST_PLANT)

n tested = 8,050; significant markers = 4; nominal support intervals = 3; lambdaGC = 1.307

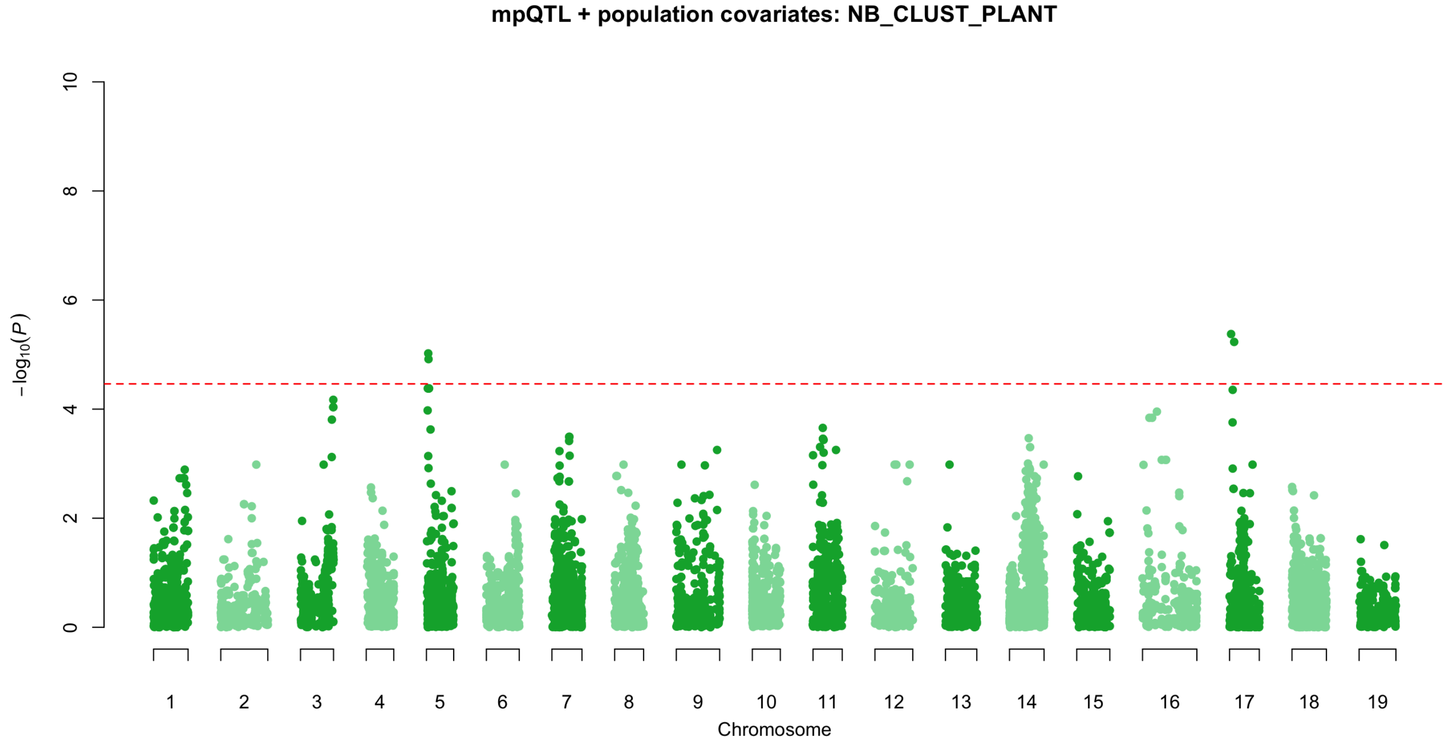


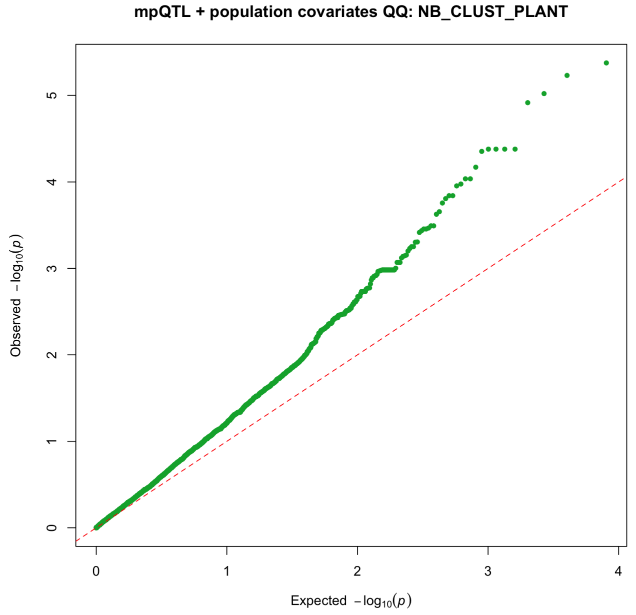


Model: realized kinship + 12 fixed population indicators; mpQTL 0.6.2 approximate P3D/EMMAX; Li–Ji threshold P = 3.4424437e-5.

Nominal positional supports are heuristic 1.5-log10(P)-drop intervals; see Supplementary Tables 10-12 and 18 for complete results and limitations.

### Population-adjusted mpQTL diagnostics — Cluster weight per plant (YIELD_PLANT)

n tested = 8,050; significant markers = 2; nominal support intervals = 2; lambdaGC = 1.102

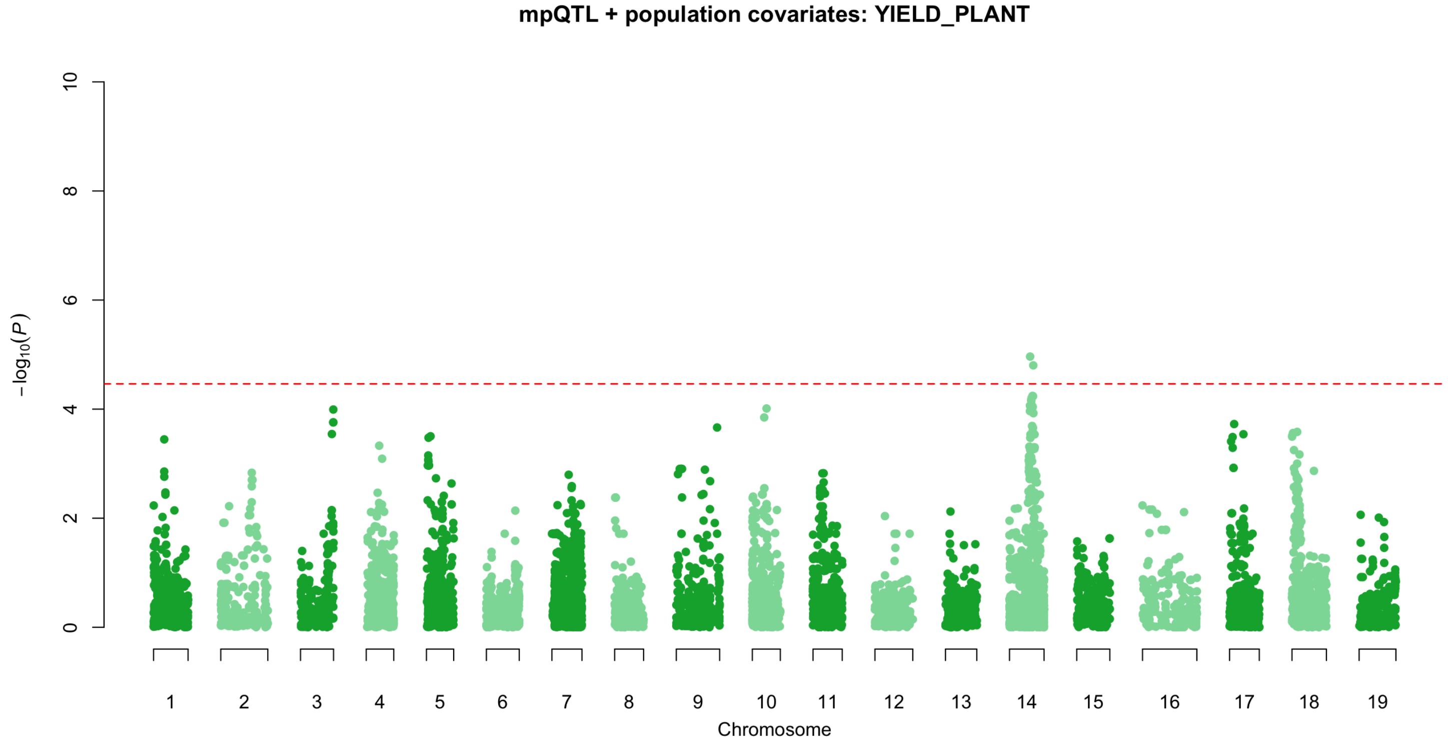


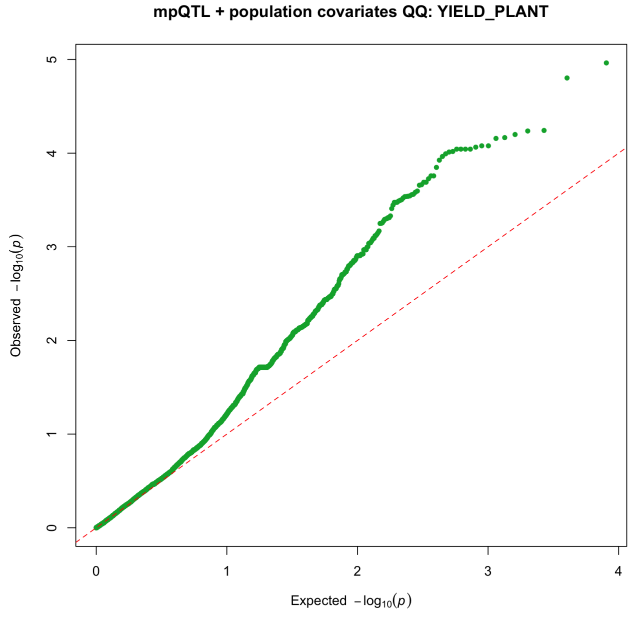


Model: realized kinship + 12 fixed population indicators; mpQTL 0.6.2 approximate P3D/EMMAX; Li–Ji threshold P = 3.4424437e-5.

Nominal positional supports are heuristic 1.5-log10(P)-drop intervals; see Supplementary Tables 10-12 and 18 for complete results and limitations.

### Population-adjusted mpQTL diagnostics — Mean cluster weight (SCLUST_W)

n tested = 7,686; significant markers = 2; nominal support intervals = 2; lambdaGC = 1.191

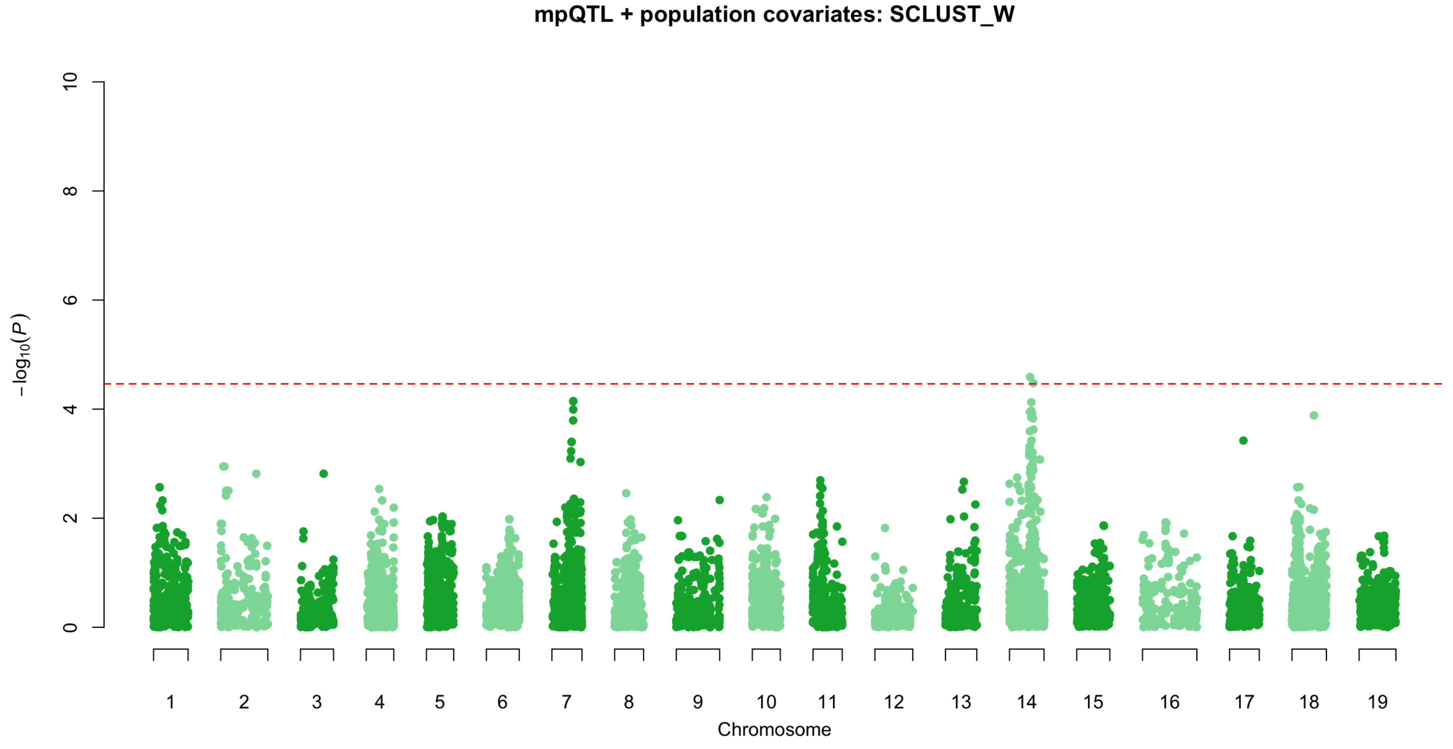


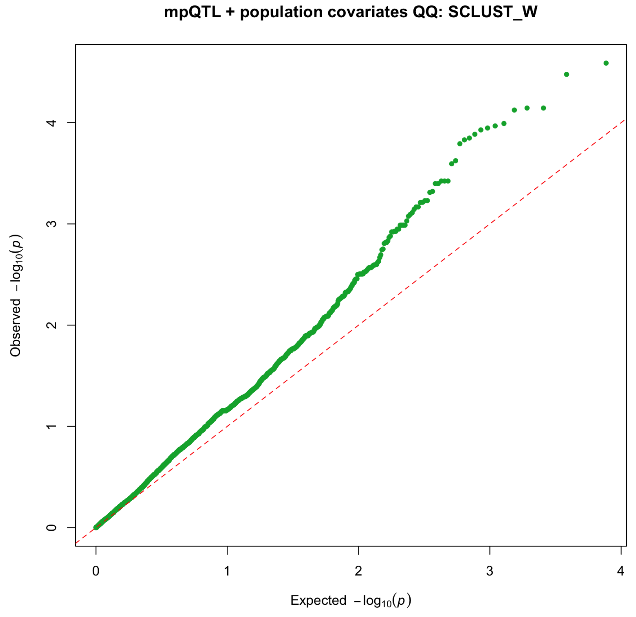


Model: realized kinship + 12 fixed population indicators; mpQTL 0.6.2 approximate P3D/EMMAX; Li–Ji threshold P = 3.4424437e-5.

Nominal positional supports are heuristic 1.5-log10(P)-drop intervals; see Supplementary Tables 10-12 and 18 for complete results and limitations.

### Population-adjusted mpQTL diagnostics — Yield per m² (YIELD_OIV_504)

n tested = 8,050; significant markers = 5; nominal support intervals = 1; lambdaGC = 1.120

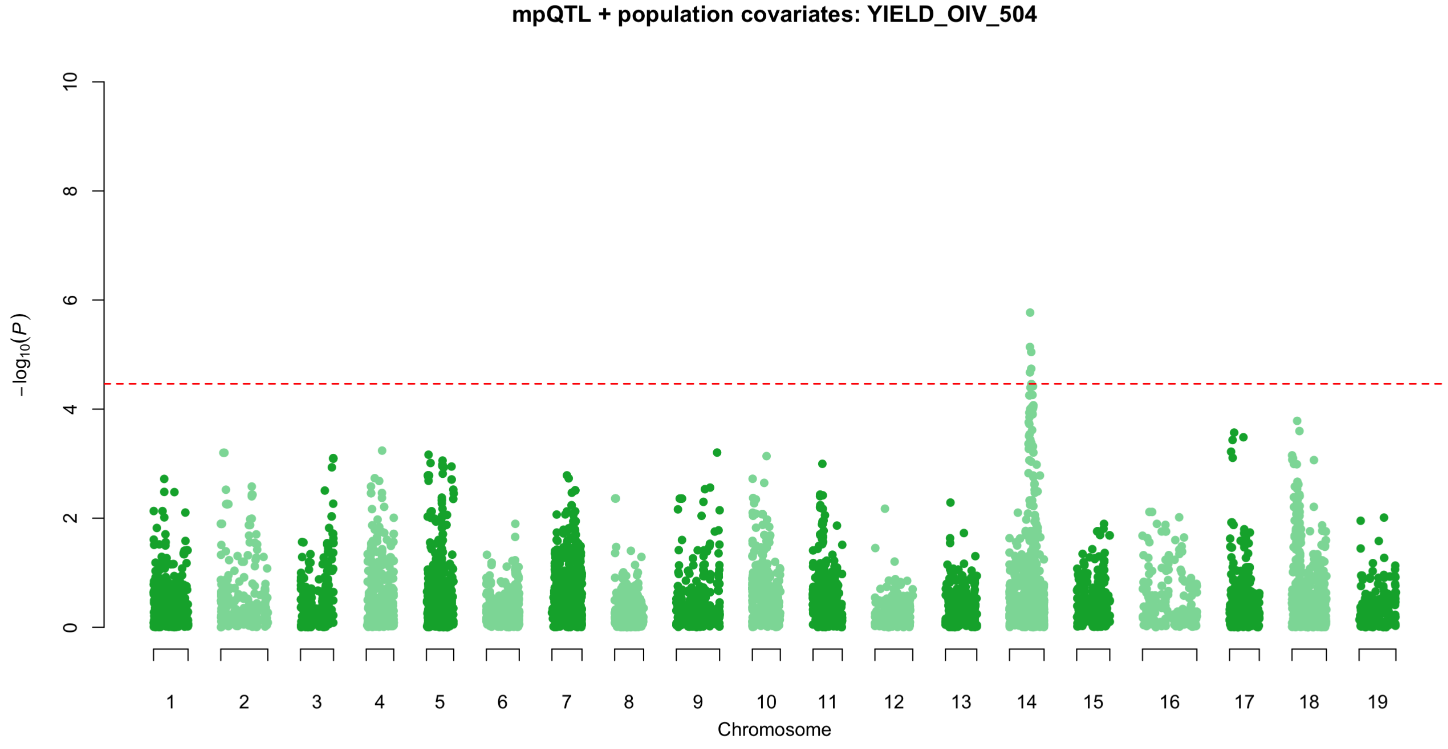


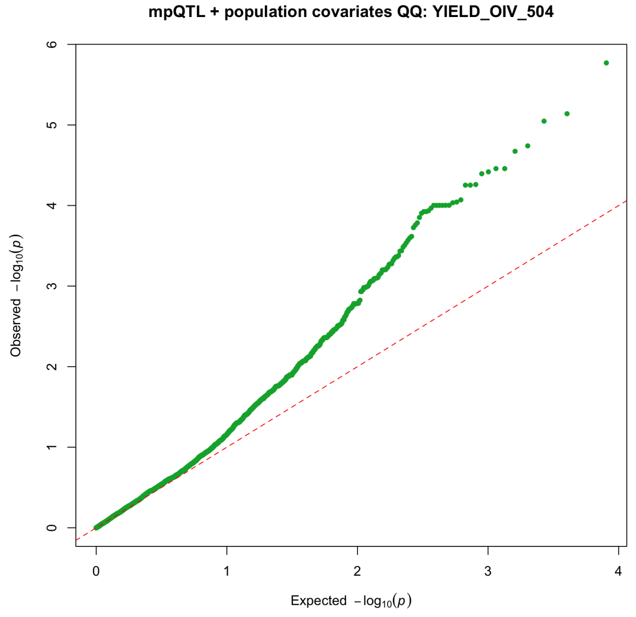


Model: realized kinship + 12 fixed population indicators; mpQTL 0.6.2 approximate P3D/EMMAX; Li–Ji threshold P = 3.4424437e-5.

Nominal positional supports are heuristic 1.5-log10(P)-drop intervals; see Supplementary Tables 10-12 and 18 for complete results and limitations.
